# Analysis of the limited *M. tuberculosis* accessory genome reveals potential pitfalls of pan-genome analysis approaches

**DOI:** 10.1101/2024.03.21.586149

**Authors:** Maximillian G. Marin, Christoph Wippel, Natalia Quinones-Olvera, Mahboobeh Behruznia, Brendan M. Jeffrey, Michael Harris, Brendon C. Mann, Alex Rosenthal, Karen R. Jacobson, Robin M. Warren, Heng Li, Conor J. Meehan, Maha R. Farhat

## Abstract

Pan-genome analysis is a fundamental tool for studying bacterial genome evolution; however, the variety of methods used to define and measure the pan-genome poses challenges to the interpretation and reliability of results. To quantify sources of bias and error related to common pan-genome analysis approaches, we evaluated different approaches applied to curated collection of 151 *Mycobacterium tuberculosis* (*Mtb*) isolates. *Mtb* is characterized by its clonal evolution, absence of horizontal gene transfer, and limited accessory genome, making it an ideal test case for this study. Using a state-of-the-art graph-genome approach, we found that a majority of the structural variation observed in *Mtb* originates from rearrangement, deletion, and duplication of redundant nucleotide sequences. In contrast, we found that pan-genome analyses that focus on comparison of coding sequences (at the amino acid level) can yield surprisingly variable results, driven by differences in assembly quality and the softwares used. Upon closer inspection, we found that coding sequence annotation discrepancies were a major contributor to inflated *Mtb* accessory genome estimates. To address this, we developed panqc, a software that detects annotation discrepancies and collapses nucleotide redundancy in pan-genome estimates. When applied to *Mtb* and *E. coli* pan-genomes, panqc exposed distinct biases influenced by the genomic diversity of the population studied. Our findings underscore the need for careful methodological selection and quality control to accurately map the evolutionary dynamics of a bacterial species.

## Introduction

Even within the boundaries of phenotypically defined bacterial species, gene content can vary substantially. (1, 2). The concept of pan-genome, often defined as the union of all genes found across a population (3), emerged from the necessity to describe this variability in genomic content. Stemming originally from bacterial genomics, this concept now extends throughout the tree of life(4). Genes in the pan-genome are typically divided into two categories: core genes, which are shared by all, or nearly all, members of the population, and accessory genes, which are found only in a subset of the population (3). Multiple mechanisms drive variability in bacterial genomic content, including horizontal gene transfer, sequence deletion, and sequence duplication with diversification (5). With increasing frequency, pan-genome analyses are generating new insights into the genetic diversity and adaptability of bacterial populations, with important implications for fields such as medicine, agriculture, and environmental science (1, 6–10).

A large number of bioinformatic tools have been developed to analyze genome content across and between species, but they often differ in their approach for defining and measuring the pan-genome. For example, there are at least 38 different pan-genome analysis pipelines described in a recent review (3). Critical to any pan-genome analysis are two key choices: 1) the unit of sequence to be compared across genomes (whole genome or coding sequences, nucleotide or amino acid, etc.) and 2) the criteria for evaluating sequence similarity and homology. For example, analysis may focus on nucleotide sequence, k-mer content, annotated coding sequences (CDSs), or COGs (Clusters of Orthologous Genes), to name a few. These choices, with differences in resolution and sensitivity, can significantly change the type of variation detected, ultimately affecting the downstream interpretation of a pan-genome. Furthermore, the specific sample of genomes and whether they are sufficiently representative of intraspecific diversity can also impact the size and content of the predicted pan-genome. In addition to these challenges, the sequencing technology and genome assembly approach can impact the accuracy and completeness of the pan-genomes generated(11). Pan-genome inference is thus highly sensitive to methodological assumptions made during the comparison and clustering of sequences, as well as to the quality and representativeness of the input sequences.

*Mycobacterium tuberculosis (Mtb)* is the primary cause of the disease tuberculosis, which is responsible for an estimated 1.6 million deaths every year (12). *Mtb* is a clonally evolving pathogen with no ongoing horizontal gene transfer or interstrain recombination (13–15), and limited structural variation. Therefore, the *Mtb* pan-genome is believed to be shaped largely by deletions, and some duplications of existing sequence, leading to an overall pattern of reductive genome evolution (16–18). In contrast to this view, some studies have inferred a surprisingly wide range of pan-genome sizes (19, 20) for *Mtb* populations, as well as for other highly similar bacteria within the *Mycobacterium tuberculosis* complex. For instance, one study estimated 7,620 distinct accessory genes in the *Mtb* pan genome, while recent studies have estimated an accessory genome on the order of 500 genes (18, 21).

Due to its conserved genome structure and lack of horizontal gene transfer, *Mtb* is a useful organism to benchmark the specificity of accessory genome prediction in pan-genome analyses. Additionally, genomic clonality and limited structural variation in *Mtb* facilitate complete genome assembly using hybrid long- and short-read sequencing. This allows us to study how sequencing and assembly quality independently influence the pan-genome inference of a highly clonal population.

In this work we use hybrid short- and long-read assemblies from 151 clinical *Mtb* genomes, spanning 7 lineages, to quantify the accessory genome of human derived *Mtb*. We first build a pan-genome graph to characterize structural variation between genomes at the nucleotide level. We find that a majority of the structural variation in the *Mtb* genome involves reconfiguration of existing nucleotide sequence content, instead of true differences in gene content. Then, we benchmark common bacterial pan-genome analysis tools and find that several pipelines are prone to overinflating accessory genome size due to CDS annotation discrepancies, and that this pitfall can be worsened by the use of short-read assemblies as input.

To address this we developed panqc, a software that detects annotation discrepancies and collapses CDSs with redundant nucleotide sequences in pan-genome predictions. Our software takes outputs from existing pan-genome prediction software, such as Panaroo and Roary, and collapses together CDSs that are highly similar at the nucleotide level. This tends to correct inflated pan-genome estimates that are largely driven by discrepancies in CDS annotations. The goal of panqc is to complement existing pan-genome analysis tools, by transparently allowing users to correct for nucleotide redundancy according to the needs of their analysis.

## Results

### Curating a dataset of high quality ***Mtb*** genomes

In order to confidently assess the pan-genome of *Mtb*, we curated a collection of 151 complete assemblies from diverse human adapted isolates, sequenced with both short-, and long-read technologies (Oxford Nanopore and PacBio). For each isolate, we generated both a hybrid genome assembly (long read *de novo* assembly with short-read polishing) and a short-read (SR) *de novo* assembly. This was done using 6 previously published datasets (22–25), as well as 8 isolates sequenced for this study (PacBio HiFi) (**Methods**). The resulting dataset spans the global diversity of the *Mtb* phylogeny by including lineages 1-6 and 8 (**Figure 1A**). The hybrid genome assemblies have high sequence similarity and conserved genome characteristics: estimated pairwise average nucleotide identity (ANI) (99.8-100%), pairwise k-mer jaccard similarity (0.94-0.99), genome size (4.38-4.44 Mb), number of predicted proteins (4020 - 4135 CDSs), and GC content (65.6% - 65.6%) (**Figure 1B, S1, Table 1**). As would be expected, SR and hybrid assemblies predominantly showed differences in continuity and size. SR assemblies were consistently more fragmented, had a slightly lower cumulative length, and had fewer predicted coding sequences compared to the hybrid assemblies (**Supp Results, Table 1, Figure S2**).

**Figure 1.**
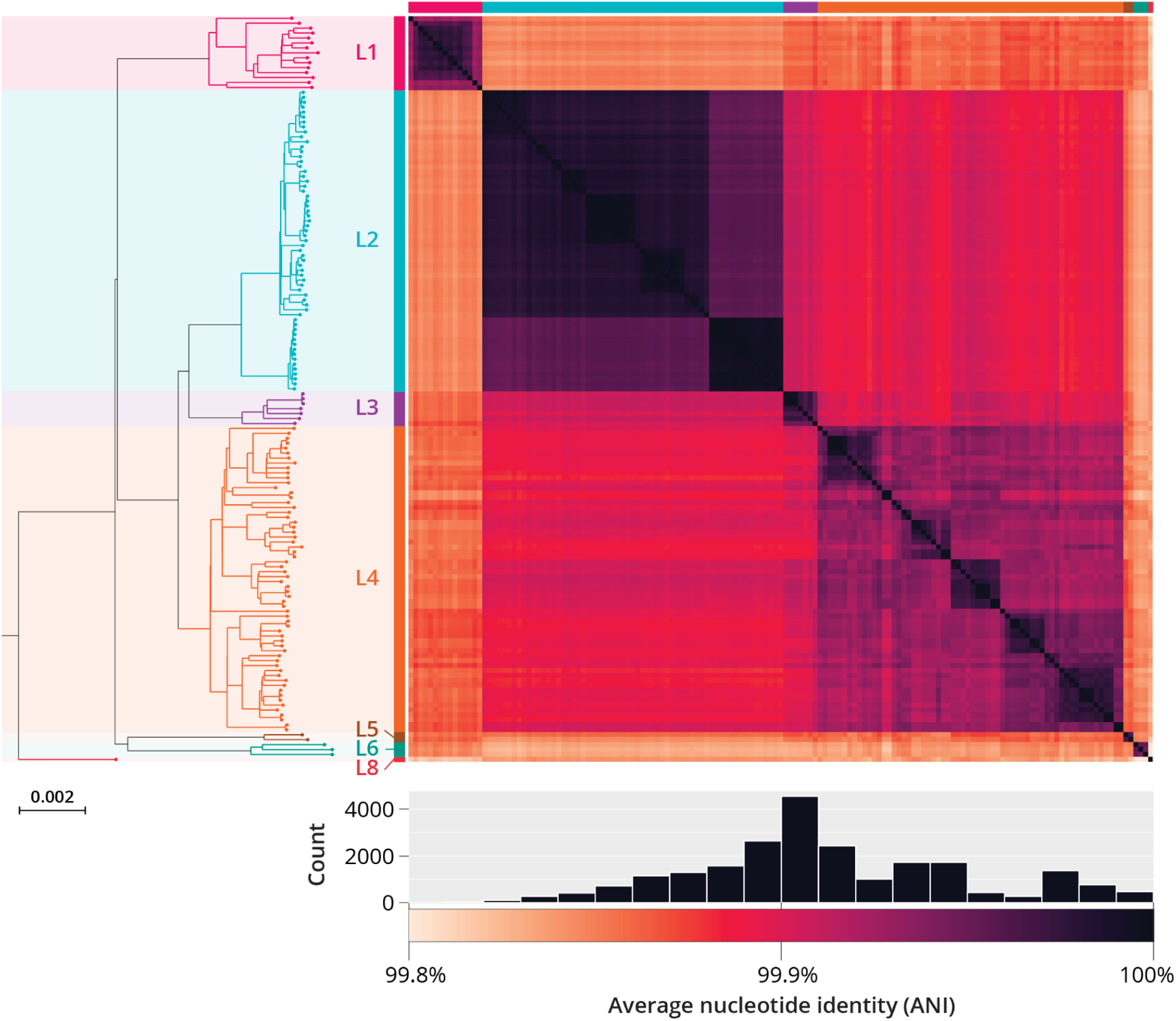
Overview of dataset of 151 complete *Mtb* genomes. Left, maximum likelihood phylogeny of 151 *Mtb* isolates generated from hybrid genome assemblies. Right, heatmap of Average Nucleotide Identity (ANI) predictions between all pairs of complete *Mtb* genomes inferred using FastANI algorithm (27). The clustering based on pairwise ANI values agrees with the known lineage divisions identified by the SNP based ML phylogeny. Below, distribution of ANI values between all pairs of complete genomes. The distribution of estimated pairwise ANI values between all pairs of *Mtb* genomes ranged from 99.8% to 100% (median pairwise ANI: 99.9%).

**Table 1.** Assembly characteristics compared between complete & short read only assemblies (N=151)

| Stat | Assembly Approach | Median w/ IQR |
| --- | --- | --- |
| BUSCO Completeness Score | Hybrid (LR + SR) | 99.4 (IQR: 99.3 - 99.5) |
|  | Short read only | 99.4 (IQR: 99.3 - 99.6) |
| Cumulative length | Hybrid (LR + SR) | 4,413 kb (IQR: 4,407 - 4,421 kb) |
|  | Short read only | 4,314 kb (IQR: 4,289 - 4,336 kb) |
| # of contigs | Hybrid (LR + SR) | 1 contigs (IQR: 1 - 1) |
|  | Short read only | 116 contigs (IQR: 106 - 132) |
| N50 | Hybrid (LR + SR) | 4,413 kb (IQR: 4,407 - 4,421 kb) |
|  | Short read only | 92 kb (IQR: 73 - 107 kb) |
| # of predicted CDSs (Bakta) | Hybrid (LR + SR) | 4074 (IQR: 4065 - 4088) |
|  | Short read only | 4035 (IQR: 4022 - 4046) |
| GC Content | Hybrid (LR + SR) | 65.6% (IQR: 65.6 - 65.6%) |
|  | Short read only | 65.5% (IQR: 65.5 - 65.6%) |

**Table 2.**
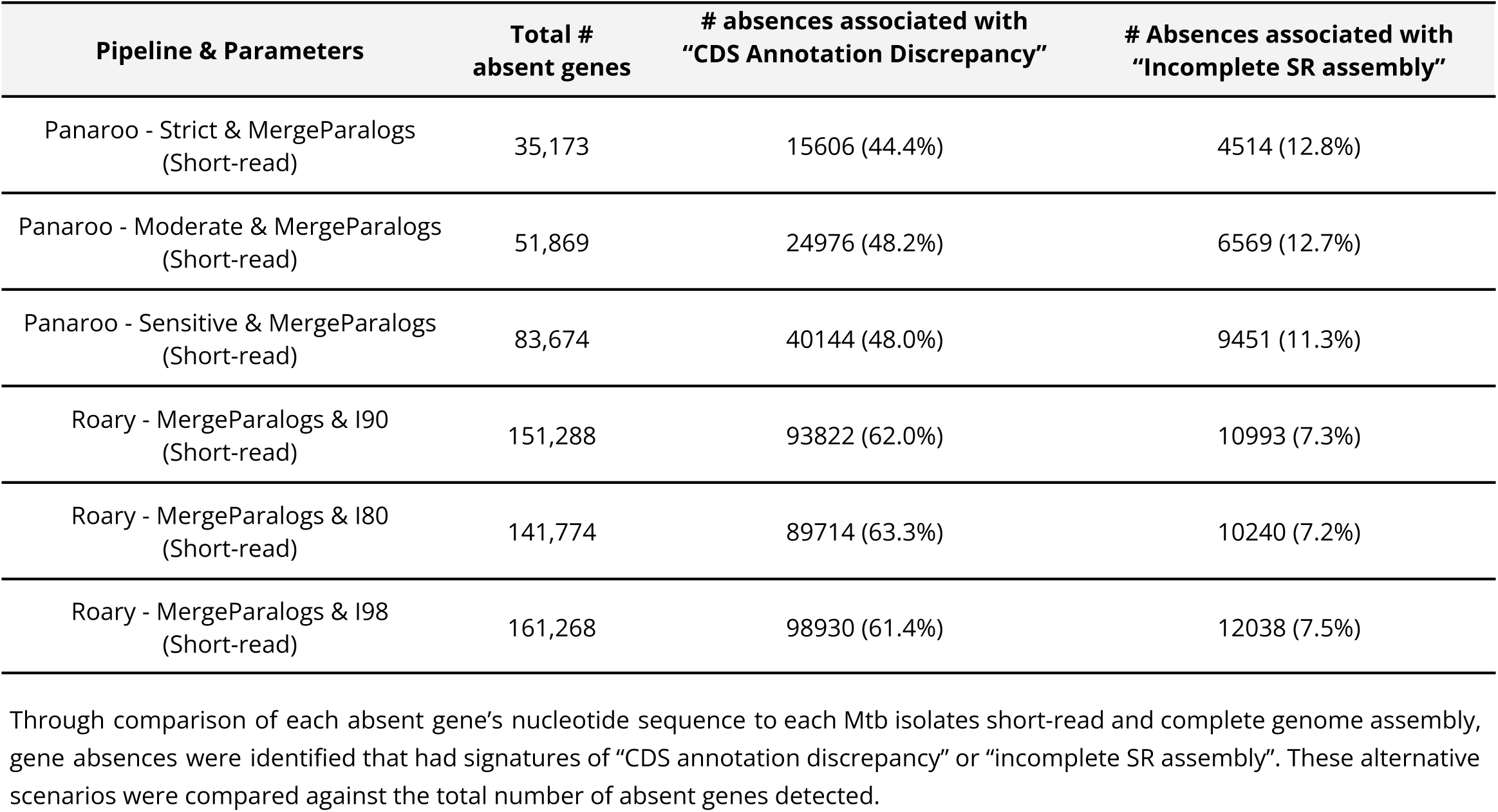
Sources of predicted gene absences with short-read based Mtb pan-genome analyses.

### Most structural variation in ***Mtb*** is attributed to rearrangements of existing sequence, rather than to novel sequence content

We used the minigraph algorithm to construct a pan-genome graph of structural variants (SVs) found across all 151 curated *Mtb* genomes and the H37Rv reference genome (26). minigraph constructs a graph of all SVs (≥ 50bp), while maintaining genomic co-linearity, facilitating comparison of the graph to the coordinates of a reference genome. We classified nodes within the pan-genome graph into core nodes and SV nodes. Core nodes represent a genomic region found across all isolates, while SV nodes represent structural variants found in at least one genome. The resulting pan-genome graph contains 536 core nodes, with a cumulative length of 3.9 Mb, and 2602 SV nodes, with a cumulative length of 1.3 Mb (**Figure 2**). Across the graph, there are 535 distinct clusters of connected SV nodes, henceforth referred to as bubble regions. Examples of two different bubble regions of varying complexity are shown in **Figure 2A**.

**Figure 2.**
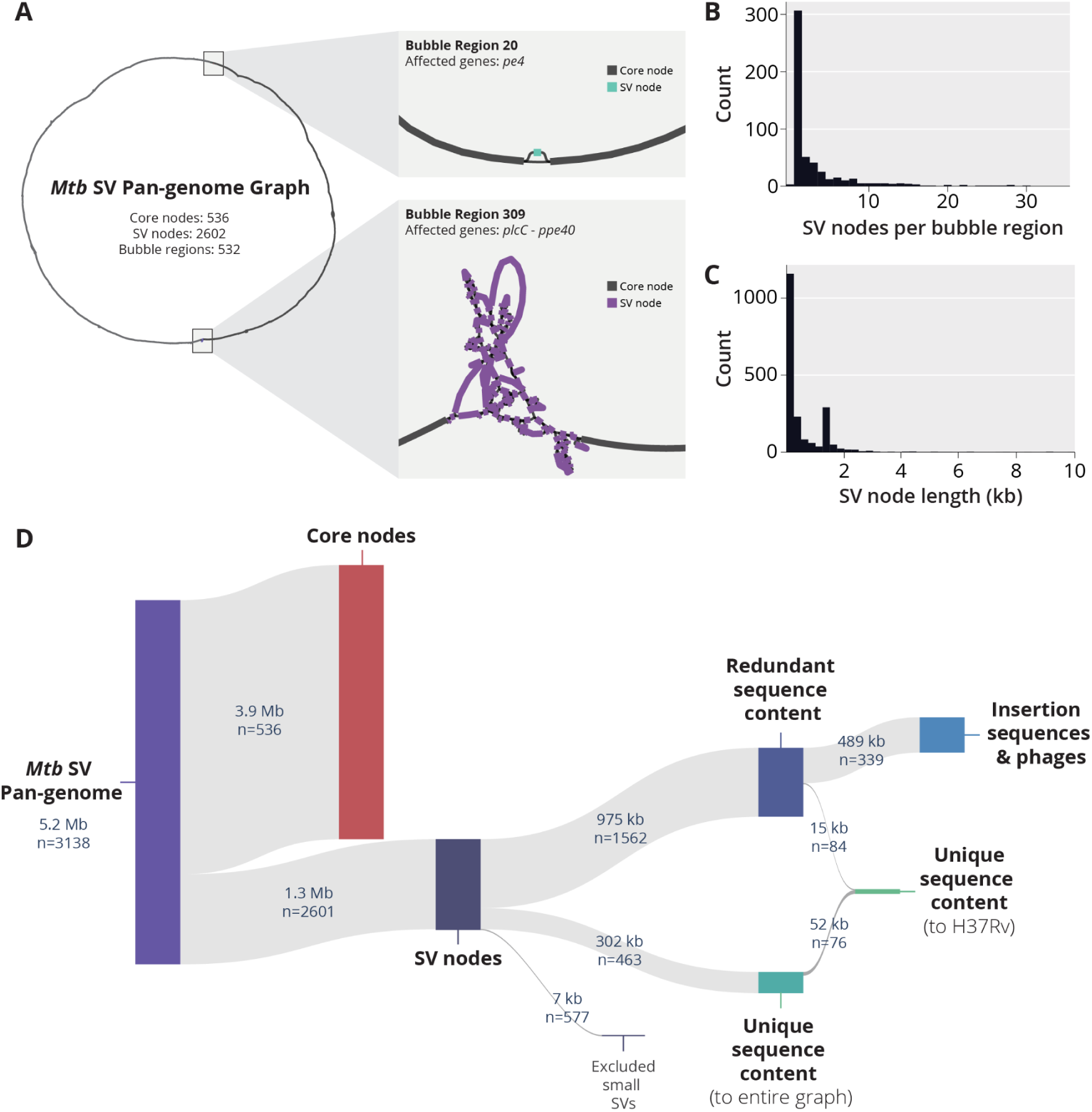
Overview of SV pan-genome analysis approach with minigraph. **A)** A high level view of the *Mtb* SV pan-genome graph generated using the Minigraph algorithm. Nodes within the graph can be classified as either core nodes, representing genome sequences found across all genomes, or SV nodes, representing sequences that are differentially present in the input set of genomes. Clusters of SV nodes within the graph are classified into bubble regions. Bubble Region #20 is an example of a simple region, containing a single SV node (186 bp) in pe4 (Rv0160c). In contrast, Bubble region #309 is a much more complex bubble region, containing a total of 88 SV nodes with a cumulative length of 55,759 bp. Bubble #309 represents structural diversity ranging from plcC (Rv2349c) to ppe40 (Rv2356c) relative to the following coordinates of the H37Rv reference genome: NC_000962.3:2,627,049-2,639,487 **B)** Distribution of the number of SV nodes per identified bubble region. **C)** Distribution of SV node length across the Mtb SV pan-genome. **D)** Hierarchical breakdown of number of nodes and cumulative length of redundant and unique sequence content within the SV pan-genome graph.

Given the large number of SV nodes detected, we aimed to understand if these nodes represented truly novel sequence content differences or if they originated from reconfigurations of existing sequences (through deletions, duplications and translocations, for example). To do this, we developed a computationally efficient k-mer comparison approach to characterize redundant sequence content within the graph (**Methods**). We identified 463 SV nodes with unique k-mer content (**Figure 2D**), indicating that only 23% (302 kb) of the total cumulative length of SV nodes represent novel sequence content. When we investigated the redundant sequence content, we found that more than half (489 kb) had a strong k-mer overlap with known *Mtb* phage and insertion sequences (**Figure S5**). Upon further inspection, a single transposable element, IS6110, was responsible for the vast majority (455 kb) of the sequence content within this category.

We identified that only a minor fraction of SV nodes (67 kb, 5% of the total length) represented sequences absent from the H37Rv reference. These SV nodes were spread across 65 bubble regions in the graph, and contain known deletions unique to specific *Mtb* lineages, such as TbD1 (17, 28)**Figure S3**). The 18 largest bubble regions (≥1 kb novel to H37Rv) are highlighted in **Table S2**.

### The choice of software and specific pipeline parameters can substantially impact pan-genome size estimates

In bacterial pan-genome analysis workflows, genomes are usually first annotated *de novo*, followed by clustering of protein coding sequences (CDS) by homology, and this is finally used to distinguish core and accessory genes. After these steps, post-processing steps vary greatly, depending on the specific pan-genome analysis pipeline used. Given the apparent overestimation of the *Mtb* accessory genome size in previous studies, we evaluated the variability in pan-genome size predictions across commonly used pipelines: Panaroo, Roary, and PPanGGolin. We focused specifically on three variables: 1) CDS clustering parameters (sequence identity threshold, merging of paralogs, and pipeline heuristics), 2) assembly type (*de novo* short-read assembly vs. hybrid genome assembly) and 3) annotation pipeline (Bakta or PGAP).(26–30) (**Methods**).

Across all analysis combinations, pan-genome estimates ranged widely from 220-2912 accessory genes and from 2868-3833 core genes (**Figure 3A, Figure S4, File S8**). Using hybrid assemblies systematically resulted in larger core genome, and smaller accessory genome estimates. Panaroo predictions for accessory genome size were the most robust to changes in assembly type and annotation pipeline, while the estimates of Roary and PPAnGGolin varied more drastically in response to these variables. Notably, using PPanGGOlin produced the most conservative accessory genome prediction when applied to PGAP annotated *Mtb* hybrid assemblies (220 accessory genes), while using Bakta annotations of the identical set of assemblies resulted in an estimate nearly 4 times larger (793 accessory genes).

**Figure 3.**
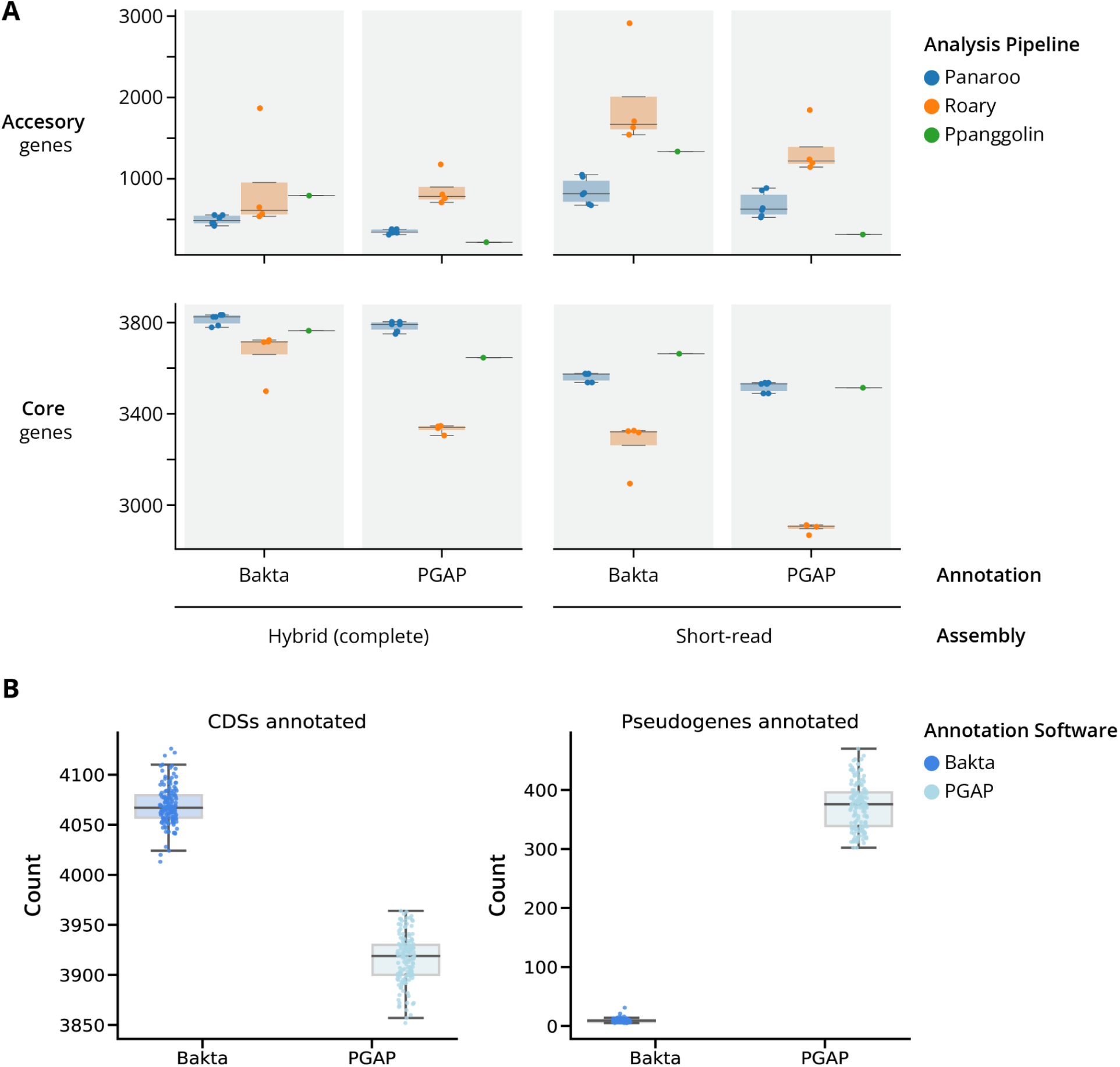
Comparing Mtb pan-genome predictions across different annotation & sequencing approaches. **A)** Comparison of the # of core and accessory genes) estimated for the identical population of 151 *Mtb* isolates across different de novo assembly types (hybrid vs short-read), annotation approaches (Bakta vs PGAP), and different parameters of pan-genome analysis pipelines (Panaroo, Roary, PPanGGolin). **B)** Abundance of CDS and pseudogene feature types annotated by Bakta and PGAP annotation pipelines across all 151 hybrid *Mtb* genomes.

Using PGAP as an annotation tool resulted in a consistently smaller pan-genome size than Bakta (**File S8, Figure 3**). Across all tested parameters, on average, using Bakta estimated 430 more total unique genes compared to using PGAP. To understand the source of this systematic increase, we compared annotation characteristics between Bakta and PGAP. Across all 151 hybrid *Mtb* assemblies, PGAP consistently annotates fewer CDSs and more pseudogenes than Bakta, the latter tending to annotate new CDSs in the case of pseudogenization and/or frameshifts (**Figure 3B, Figure S5, Tables S2-S3, Supplemental Results**).

In pan-genome analysis, the number of absent genes is often overestimated due to incomplete assemblies and annotation discrepancies

We investigated how assembly completeness and continuity affects pan-genome inference when using short-read assemblies for analysis (**Methods**). We evaluated pan-genome predictions produced with Panaroo and Roary, and focused on genes identified as absent in each short-read assembly. We then evaluated three scenarios for a gene’s absence: 1) Absence due to an incomplete SR assembly, where a gene absent in the short-read assembly can be found in the hybrid assembly, and 2) Absence due to CDS annotation discrepancy, where the CDS annotation of the gene is absent, but the gene can be identified at the nucleotide level, and 3) *bona fide* absence, where neither of the two previous criteria were met.

We found that 7%-13% of gene absences were associated with incomplete SR assembly, 44%-63% were associated with CDS annotation discrepancy (**Table 4**), while the minority were *bona fide* absences. Given this high proportion absences due to incomplete assembly or CDS annotation discrepancy, we examined the relationship between the BUSCO completeness scores and each absence scenario. We categorized assemblies based on their BUSCO scores: low (<99%, n=15) or high (>99%, n=136), and looked at its association with absence types, using our most conservative pan-genome estimate (**Figure S6, Methods**). Absences associated with incomplete assembly showed a significant increase in assemblies with low BUSCO scores (Mann-Whitney U test, p = 3.3e-9). However, the frequency of absences due to CDS annotation discrepancy was comparable between low and high BUSCO assemblies (Mann-Whitney U test, p = 0.12).

### Developing a tool to account for nucleotide redundancy within CDS based pan-genome estimates

Motivated by our observation that CDS annotation discrepancies can inflate the estimated pan-genome size, we developed panqc. CDS annotation discrepancies result in high levels of nucleotide redundancy in pan-genome estimates. To address this, panqc takes pan-genome predictions from commonly used pipelines, and collapses CDSs with highly similar nucleotide sequence content. (**Methods**). It is therefore a correction step for applications where paralogs are considered representatives of the same gene. Our algorithm consists of two steps: First, it takes all the CDSs predicted to be absent, and queries the nucleotide sequence against the associated genome assembly. If the nucleotide sequence is found, with a coverage and sequence identity > 90%, the gene is classified as being present at the DNA level, but absent at the CDS level. Second, genes are merged into nucleotide similarity clusters using a k-mer based similarity metric (**Methods**). Finally, pan-genome estimates are readjusted accounting for identified nucleotide redundancy.

We ran panqc on compatible outputs of Roary & Panaroo for our 151 *Mtb* isolates (**Figure 4A & S7, Table 3**). Across all *Mtb* estimates evaluated, panqc reduced the overall accessory genome size by 420 genes (44%) on average. When applied to the most conservative estimate produced by Panaroo (using Complete Assemblies, Strict, and MergeParalogs options), the pan-genome size was reduced by 139 genes. Upon further inspection, CDS annotation discrepancies were responsible for an average of 49% of all absent CDSs initially reported (**Table 5**).

**Figure 4.**
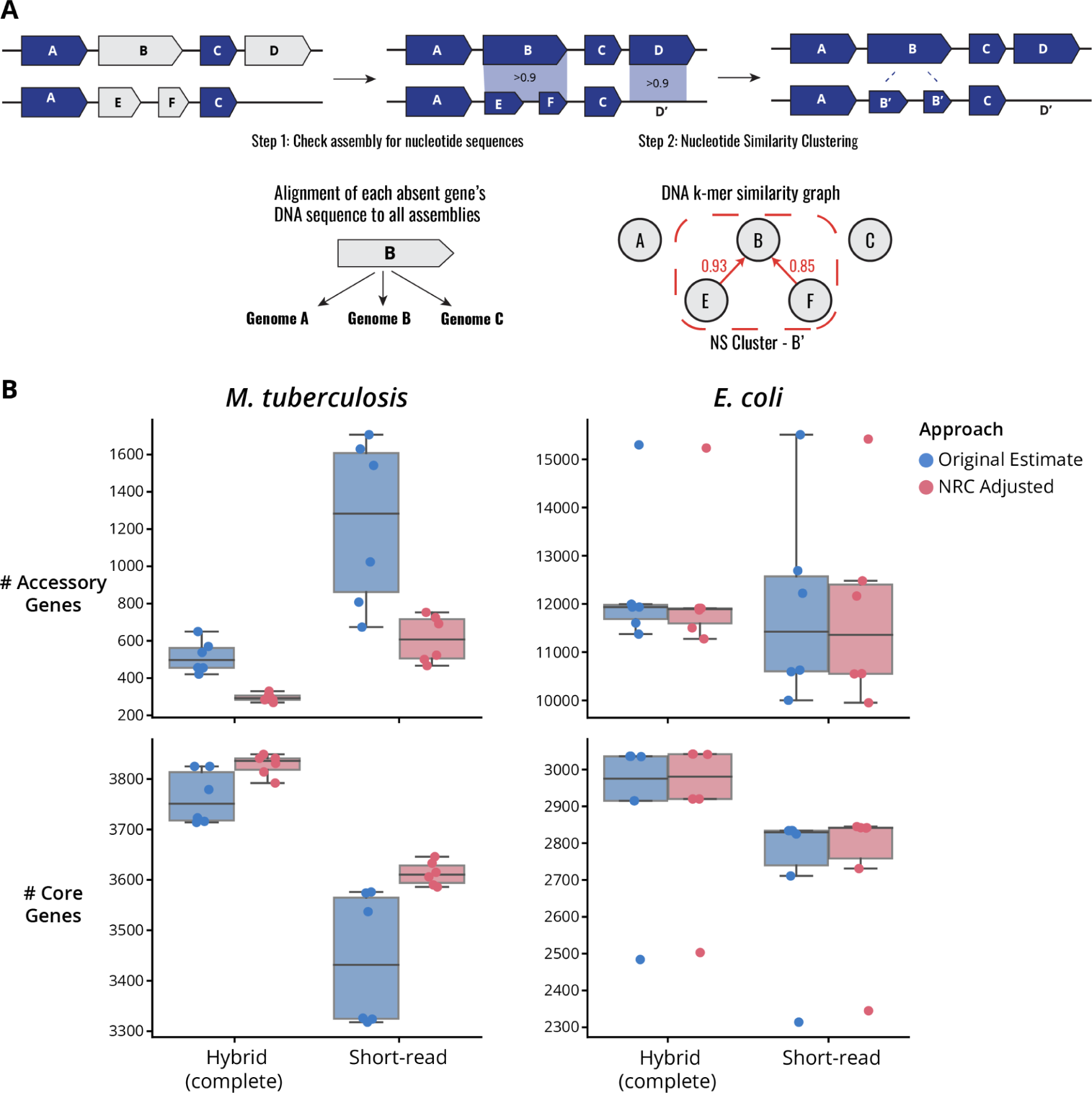
Benchmarking the panqc nucleotide redundancy correction pipeline on pan-genome analyses. **A)** Schematic diagram of the panqc nucleotide redundancy correction pipeline. First, all absent genes are compared to their corresponding genome at the nucleotide level to identify genes which are absent at the CDS-level but present at the nucleotide level. After adjustment for nucleotide presence in each assembly, a nucleotide similarity clustering step is performed to merge CDSs which share highly similar k-mer content. **B)** Comparison of pan-genome size predictions before and after NRC adjustment of Panaroo and Roary estimates for analysis of Mtb (N = 151) and E. coli (N = 50) genomes. All pan-genome size estimates for each data point can be found in **Tables 3 and 4**.

**Table 3.** Pan-genome size estimates for 151 *Mtb* isolates before and after panqc adjustment.

| Assembly type | Pipeline | Parameters | Adjustment | # total size | # core | # accessory | ΔTotal | ΔCore | ΔAccessory |
| --- | --- | --- | --- | --- | --- | --- | --- | --- | --- |
| Hybrid | Panaroo | Strict & MergeParalogs | Original | 4200 | 3779 | 421 | -139 | 13 | -152 |
|  |  |  | panqc | 4061 | 3792 | 269 |  |  |  |
| Hybrid | Panaroo | Moderate & MergeParalogs | Original | 4280 | 3825 | 455 | -155 | 16 | -171 |
|  |  |  | panqc | 4125 | 3841 | 284 |  |  |  |
| Hybrid | Panaroo | Sensitive & MergeParalogs | Original | 4281 | 3825 | 456 | -156 | 16 | -172 |
|  |  |  | panqc | 4125 | 3841 | 284 |  |  |  |
| Hybrid | Roary | MergeParalogs & Identity 98 (Default) | Original | 4366 | 3716 | 650 | -187 | 133 | -320 |
|  |  |  | panqc | 4179 | 3849 | 330 |  |  |  |
| Hybrid | Roary | MergeParalogs & Identity 90 | Original | 4293 | 3723 | 570 | -154 | 108 | -262 |
|  |  |  | panqc | 4139 | 3831 | 308 |  |  |  |
| Hybrid | Roary | MergeParalogs & Identity 80 | Original | 4252 | 3714 | 538 | -138 | 100 | -238 |
|  |  |  | panqc | 4114 | 3814 | 300 |  |  |  |
| Short-read | Panaroo | Strict & MergeParalogs | Original | 4211 | 3537 | 674 | -158 | 49 | -207 |
|  |  |  | panqc | 4053 | 3586 | 467 |  |  |  |
| Short-read | Panaroo | Moderate & MergeParalogs | Original | 4382 | 3574 | 808 | -249 | 59 | -308 |
|  |  |  | panqc | 4133 | 3633 | 500 |  |  |  |
| Short-read | Panaroo | Sensitive & MergeParalogs | Original | 4600 | 3576 | 1024 | -431 | 70 | -501 |
|  |  |  | panqc | 4169 | 3646 | 523 |  |  |  |
| Short-read | Roary | MergeParalogs & Identity 98 (Default) | Original | 5025 | 3318 | 1707 | -657 | 297 | -954 |
|  |  |  | panqc | 4368 | 3615 | 753 |  |  |  |
| Short-read | Roary | MergeParalogs & Identity 90 | Original | 4956 | 3326 | 1630 | -625 | 280 | -905 |
|  |  |  | panqc | 4331 | 3606 | 725 |  |  |  |
| Short-read | Roary | MergeParalogs & Identity 80 | Original | 4866 | 3324 | 1542 | -584 | 266 | -850 |
|  |  |  | panqc | 4282 | 3590 | 692 |  |  |  |

*Mtb* pan-genome statistics (Total, Core, and Accessory genes) were quantified before and after panqc adjustment. The panqc pipeline was run with default settings where the minimum query coverage (90%) and length (90%) for gene rediscovery at the nucleotide level. Additionally, a k-mer similarity cutoff of 0.8 was used for identifying nucleotide similarity clusters.

**Table 4.** Pan-genome size estimates for 50 *E. coli* isolates before and after panqc adjustment.

| Assembly type | Pipeline | Parameters | Adjustment | # total size | # core | # accessory | ΔTotal | ΔCore | ΔAccessory |
| --- | --- | --- | --- | --- | --- | --- | --- | --- | --- |
| Hybrid | Panaroo | Strict & MergeParalogs | Original | 14410 | 3035 | 11375 | -93 | 6 | -99 |
|  |  |  | panqc (Default) | 14317 | 3041 | 11276 |  |  |  |
| Hybrid | Panaroo | Moderate & MergeParalogs | Original | 14641 | 3036 | 11605 | -95 | 6 | -101 |
|  |  |  | panqc (Default) | 14546 | 3042 | 11504 |  |  |  |
| Hybrid | Panaroo | Sensitive & MergeParalogs | Original | 15032 | 3036 | 11996 | -115 | 6 | -121 |
|  |  |  | panqc (Default) | 14917 | 3042 | 11875 |  |  |  |
| Hybrid | Roary | MergeParalogs & Identity 98 (Default) | Original | 17783 | 2484 | 15299 | -46 | 19 | -65 |
|  |  |  | panqc (Default) | 17737 | 2503 | 15234 |  |  |  |
| Hybrid | Roary | MergeParalogs & Identity 90 | Original | 14851 | 2915 | 11936 | -21 | 5 | -26 |
|  |  |  | panqc (Default) | 14830 | 2920 | 11910 |  |  |  |
| Hybrid | Roary | MergeParalogs & Identity 80 | Original | 14851 | 2915 | 11936 | -21 | 5 | -26 |
|  |  |  | panqc (Default) | 14830 | 2920 | 11910 |  |  |  |
| Short-read | Panaroo | Strict & MergeParalogs | Original | 12836 | 2834 | 10002 | -44 | 7 | -51 |
|  |  |  | panqc (Default) | 12792 | 2841 | 9951 |  |  |  |
| Short-read | Panaroo | Moderate & MergeParalogs | Original | 13462 | 2834 | 10628 | -71 | 8 | -79 |
|  |  |  | panqc (Default) | 13391 | 2842 | 10549 |  |  |  |
| Short-read | Panaroo | Sensitive & MergeParalogs | Original | 15524 | 2834 | 12690 | -202 | 8 | -210 |
|  |  |  | panqc (Default) | 15322 | 2842 | 12480 |  |  |  |
| Short-read | Roary | MergeParalogs & Identity 98 (Default) | Original | 17825 | 2314 | 15511 | -60 | 31 | -91 |
|  |  |  | panqc (Default) | 17765 | 2345 | 15420 |  |  |  |
| Short-read | Roary | MergeParalogs & Identity 90 | Original | 14933 | 2711 | 12222 | -37 | 20 | -57 |
|  |  |  | panqc (Default) | 14896 | 2731 | 12165 |  |  |  |
| Short-read | Roary | MergeParalogs & Identity 80 | Original | 13420 | 2825 | 10595 | -20 | 20 | -40 |
|  |  |  | panqc (Default) | 13400 | 2845 | 10555 |  |  |  |

*E. coli* pan-genome statistics (Total, Core, and Accessory genes) were quantified before and after panqc adjustment. The panqc pipeline was run with default settings where the minimum query coverage (90%) and length (90%) for gene rediscovery at the nucleotide level. Additionally, a k-mer similarity cutoff of 0.8 was used for identifying nucleotide similarity clusters.

**Table 5.** Frequency of CDS Annotation discrepancies associated with absent gene calls in *Mtb* pan-genome analyses.

| Assembly type | Pipeline | Parameters | Total #<br>absent genes | # absences associated with<br>“CDS Annotation Discrepancy” |
| --- | --- | --- | --- | --- |
| Hybrid | Panaroo | Strict & MergeParalogs | 33046 | 10784 (32.6%) |
| Hybrid | Panaroo | Moderate & MergeParalogs | 36487 | 12852 (35.2%) |
| Hybrid | Panaroo | Sensitive & MergeParalogs | 36625 | 12976 (35.4%) |
| Hybrid | Roary | MergeParalogs & Identity 98 (Default) | 64856 | 35226 (54.3%) |
| Hybrid | Roary | MergeParalogs & Identity 90 | 54830 | 29292 (53.4%) |
| Hybrid | Roary | MergeParalogs & Identity 80 | 51372 | 28144 (54.8%) |
| Short-read | Panaroo | Strict & MergeParalogs | 35173 | 15606 (44.4%) |
| Short-read | Panaroo | Moderate & MergeParalogs | 51869 | 24976 (48.2%) |
| Short-read | Panaroo | Sensitive & MergeParalogs | 83674 | 40144 (48.0%) |
| Short-read | Roary | MergeParalogs & Identity 98 (Default) | 161268 | 98930 (61.3%) |
| Short-read | Roary | MergeParalogs & Identity 90 | 151288 | 93822 (62.0%) |
| Short-read | Roary | MergeParalogs & Identity 80 | 141774 | 89714 (63.3%) |

Because pan-genome estimates can be used to assess gene gain and loss across the phylogeny, we assessed the effect of using the panqc correction on these predictions. Gene gain and loss events were predicted by ancestral state reconstruction using gene presence/absence information before and after panqc adjustment (**Methods**). The panqc adjustment reduced the number of gene gain events predicted, from 719 to 285, while reducing the number of gene loss events from 1176 to 999 (**Figure S9**). This suggests that redundant sequence content can result in false predictions of gain events. In both cases, gene loss appeared to be the main force driving *Mtb* genomic evolution, but the pattern became more apparent after adjusting for nucleotide redundancy.

To assess the applicability of panqc beyond a hyper-clonal population, we applied it to pan-genome analyses of hybrid and short-read assemblies originating from 50 *E.coli* isolates spanning nine phylogroups (**Methods, Figure S10-11**) (29). The *E. coli* pan-genome estimates ranged from 2314-3036 core genes and 10002-15511 accessory genes. Applying panqc modestly reduced the estimated accessory genome size by 80 genes (0.7%) on average (**Figure 4B & S8, Table 4**). This is similar to the absolute number of genes corrected for *Mtb*, but much smaller in proportion to the overall *E.coli* accessory genome size. In contrast to *Mtb*, CDS annotation discrepancies in *E. coli* were on average responsible for only 1.0% of all absent genes initially reported by the pan-genome estimate (**Table 6**).

**Table 6.** Frequency of CDS Annotation discrepancies associated with absent gene calls in *E. coli* pan-genome analyses.

| Assembly type | Pipeline | Parameters | Total #<br>absent genes | # absences associated with<br>“CDS Annotation Discrepancy” |
| --- | --- | --- | --- | --- |
| Complete | Panaroo | Strict & MergeParalogs | 481901 | 5915 (1.2%) |
| Complete | Panaroo | Moderate & MergeParalogs | 492056 | 6050 (1.2%) |
| Complete | Panaroo | Sensitive & MergeParalogs | 511206 | 6798 (1.3%) |
| Complete | Roary | MergeParalogs & Identity 98 (Default) | 651347 | 9102 (1.4%) |
| Complete | Roary | MergeParalogs & Identity 90 | 506502 | 1565 (0.3%) |
| Complete | Roary | MergeParalogs & Identity 80 | 506502 | 1565 (0.3%) |
| Short-read | Panaroo | Strict & MergeParalogs | 406594 | 4523 (1.1%) |
| Short-read | Panaroo | Moderate & MergeParalogs | 435712 | 5869 (1.3%) |
| Short-read | Panaroo | Sensitive & MergeParalogs | 536646 | 10516 (2.0%) |
| Short-read | Roary | MergeParalogs & Identity 98 (Default) | 654563 | 10412 (1.6%) |
| Short-read | Roary | MergeParalogs & Identity 90 | 511437 | 2905 (0.6%) |
| Short-read | Roary | MergeParalogs & Identity 80 | 437116 | 2419 (0.6%) |

## Discussion

In our work, we comprehensively evaluated different approaches for estimating the *Mtb* pan-genome, providing practical insights regarding the pitfalls of pan-genome analysis. Using a state-of-the-art pan-genome graph approach, we confirmed that *Mtb* genome’s nucleotide content is highly conserved and that genomic reduction is the predominant force driving *Mtb* evolution. We found that using an amino-acid centered approach can yield highly variable pan-genome predictions (even predicting thousands of accessory genes), but nucleotide-based comparison reveals that much of this predicted “accessory” arises from rearrangement of existing redundant sequences, and not from novel sequence content. In particular, we identified that discrepancies in CDS annotation are a key factor that can contribute to the inflation of accessory genome size. Motivated by these observations, we developed panqc, a software to adjust pan-genome predictions for nucleotide redundancy by first identifying CDS annotation discrepancies and then collapsing CDSs with highly similar nucleotide sequence content.

To assess sources of variation in *Mtb* pan-genome estimates, we compared different pan-genome prediction pipelines, where we tested combinations of different genome assembly strategies, annotation pipelines, and pan-genome estimation software. We found a large range of predicted accessory genome sizes, ranging from 220-2951 total genes, raising concerns about the utility of these estimates for studying gene content differences in *Mtb*. We found that using complete hybrid assemblies resulted in more consistent pan-genome predictions relative to using short-read assemblies. This is due to the fact that the short-read assemblies used have inconsistent levels of genome completeness and continuity. These results demonstrate that assessing consistency in completeness prior to pan-genome analysis of short-read assemblies is a crucial quality control step to avoid inflated predictions of the accessory genome, as has been addressed in previous work(11). Furthermore, independent of assembly strategy, the choice of annotation pipeline had an important effect on the pan-genome predictions. We found that using PGAP resulted in a consistently smaller pan-genome than using Bakta. Further exploration revealed that this is due to PGAP annotating pseudogenes with higher frequency. Pseudogenes are by default considered non-coding sequences, and if they are not explicitly accounted for by the pipeline used, they will be considered absent, despite the nucleotide sequence being present. The interaction between the annotation pipeline and how the pan-genome software handles pseudogenes can contribute to systematic biases in pan genome predictions.

Each pan-genome software tested demonstrated distinct strengths and weaknesses. Panaroo produced the most consistent prediction across all approaches evaluated, and was robust to differences in CDS annotation, pseudogenes, and variability of assembly quality and completeness. This indicates that Panaroo’s heuristics that account for confounding factors, such as pseudogenization, assembly errors, and/or contamination are quite effective (30). Despite this, we still observe a substantial proportion of nucleotide redundancy in even Panaroo’s most conservative *Mtb* pan-genome estimates. This may reflect the difficulty of accounting for the partícularly high amount of variation in the repetitive and homologous regions of the *Mtb* genome.

Interestingly, PPanGGolin produced much more conservative estimates of accessory genome size when using PGAP annotations instead of Bakta. From investigation of this results, it appears that PPanGGolin is prone to inflated accessory gene estimates when using annotation approaches that are to more likely to annotate multiple CDSs instead of pseudogenes. Roary was the most sensitive to CDS annotation discrepancies, rendering it less suitable for detection of complete loss or gain of gene sequences. Nevertheless, in cases where an analysis needs to be highly sensitive to differences in coding potential, Roary’s increased sensitivity can be beneficial.

We developed a software to address the pitfalls we observed in pan-genome estimates, specifically those related to CDS annotation discrepancies and nucleotide redundancy. Panqc is a software we designed to complement, instead of replace, existing pan-genome software outputs by giving the user transparent control over how nucleotide redundancy is accounted for in their analysis. We found that even the most conservative pan-genome estimates for *Mtb* still had a surprising amount of nucleotide redundancy driving the size of the predicted accessory genome. Using panqc allowed us to focus our analysis on differential presence of complete gene sequences, instead of differences in CDS annotations created by small genetic variants or errors in assembly.

To explore the broader utility of panqc, we ran it on pan-genome predictions of a diverse set of *E. coli* isolates. These isolates have high species-level diversity, putting them at the opposite end of the spectrum from the clonal *Mtb*. In contrast to *Mtb*, we detected relatively minimal amounts of nucleotide redundancy in our *E. coli* pan-genome estimates. This indicates that the pitfalls we encountered are particularly relevant for comparison of highly similar genomes, like our *Mtb* dataset, but become less of a problem when the population of genomes compared drastically differs in terms of sequence content. We suspect that as the population of genomes analyzed becomes more similar in gene content, CDS annotation discrepancies will increase in relative proportion of the estimated pan-genome. Even in cases where there is little nucleotide redundancy detected, panqc can give users confidence that their pan-genome estimates are not being primarily driven by CDS annotation discrepancies. We envision that panqc can be used in conjunction with other tools available for quality control and validation of pan-genome estimates, such as Panaroo’s suite of post-processing scripts or Panstripe(30, 31).

Overall, this work emphasizes the need to compare genomes at both the nucleotide and amino-acid levels to provide a comprehensive view of a population’s genome evolution and diversity. While certain research questions can be addressed by focusing primarily on either protein-level or nucleotide-level differences, a comprehensive grasp of the evolutionary dynamics influencing protein variation will require methods that smartly integrate and leverage both levels of sequence information. Steps in this direction are exemplified by two softwares examined in this work, Panaroo and PPanGGolin(30, 32), along with the recent development of new tools like ggcaller and pangene(33, 34), all of which enhance their CDS analysis through the use of pan-genome graphs. We anticipate that improvements in the next wave of pan-genome analysis methods will continue to come from approaches that innovate on the integration of nucleotide and amino acid level information in biological meaningful ways.

## Methods

### Dataset of clinical ***Mtb*** isolates with long- and short-read WGS

We compiled a dataset of 151 *Mycobacterium tuberculosis* (*Mtb*) isolates with both short-read (Illumina) and long-read (Oxford Nanopore, PacBio) sequencing data, including both published data (n = 143) and newly sequenced isolates (n = 8, PacBio HiFi and Illumina WGS). **File S2** details all relevant ENA/SRA run accessions and metadata. Due to significant variations in sequencing depth and read length within the published Oxford Nanopore (ONT) datasets, we employed stringent selection criteria for inclusion in analysis. Specifically, we selected only isolates with ONT sequencing that could be assembled into a single, circular contig when using the Flye assembler (v2.6). This selection was crucial to ensure that the hybrid assemblies reflect complete *Mtb* genomes.

### H37Rv reference genome and annotations

The H37Rv (NCBI Accession: `NC_000962.3`) genome sequence and annotations was used as the standard reference genome for all analyses involving *Mtb*. Functional category annotations for all genes of H37Rv were downloaded from Release 3 (2018-06-05) of MycoBrowser (https://mycobrowser.epfl.ch/releases). The H37Rv reference sequence was also annotated with the `Bakta` (v4.8) and `PGAP` (v6.4) pipelines for comparison with the official H37Rv annotations. The DNA Features Viewer python library was used to generate programmatic visualizations of the NCBI, PGAP, and Bakta H37Rv annotations (35) shown in **Figure S5** and in **Files S9-S10**.

### Selection of a diverse dataset of E. coli genome assemblies

From a published analysis of *E. coli* genomic diversity, 50 published genomes were selected from Shaw et. al. 2021(29). In this study, all genomes were assembled using an hybrid approach using both long and short-read genome sequencing data. In order to assure a diverse set of genomes, representative isolates from the following 9 *e. coli* phylotypes were selected from published metadata: A, B1, B2, C, D, E, F, G, and clade V. To complement the available hybrid assemblies, the paired-end short-read genome sequencing data for each isolate was downloaded from the NCBI Sequence Read Archive for *de novo* short-read assembly. Metadata for all evaluated *E. coli* isolates, including assembly and sequencing run accessions, are provided in **Supplemental File S2**.

### Hybrid genome assembly with long and short read sequencing

The hybrid genome assembly and polishing process was tailored to the specific requirements of various long-read WGS platform and chemistry versions used for analysis (PacBio subreads [RSII & Sequel II], ONT v9.4.1, PacBio CCS/HiFi [Sequel II[reads), as well as taking into account the software versions available at the time of data processing. Refer to the supplemental methods for the exact combination of softwares used.

### Short read ***de novo*** assembly

The following assembly approach was applied to all paired-end Illumina WGS data from *M. tuberculosis* and *E. coli* isolates. First, the paired-end Illumina WGS reads were trimmed with `Trimmomatic` (v0.39). After read processing, *de novo* short-read assemblies were then generated using `Unicycler` (v0.4.8), which serves as an assembly optimizer for `SPAdes` (v3.13). Prior to assembly of the *Mtb* isolates, the trimmed reads were additionally filtered using `Kraken2` to keep only reads that were confidently classified as Mycobacterium tuberculosis complex (MTBC, TaxID: 77643)(36). After assembly of the *Mtb* isolates, `Kraken2` was used to select only contigs that were classified as MTBC (TaxID: 77643). This Kraken2 filtering was performed to minimize chances of contaminating contigs from other species being included in the pan-genome analysis using short-reads. The standard complete `Kraken2` RefSeq database was used for all sequence classification. All relevant code, with exact run parameters, can be found in the `4.Mtb.Generate.SRAsms.smk` and `11.Ecoli.SRAsms.PGAnalysis.smk` Snakemake workflows.

### Phylogeny inference of ***Mtb*** isolates

Genetic variants relative to the H37Rv reference genome were inferred for each hybrid genome assembly using `minimap2` & `paftools.js`(37). A concatenated SNP alignment was then generated by identifying and extracting single nucleotide polymorphisms (SNPs) from each genome assembly using bcftools(38). From the SNP alignment a maximum likelihood phylogeny was inferred using IQ-Tree with the general time reversible model and a SNP ascertainment bias correction(39). The inferred phylogeny can be found on Zenodo (10.5281/zenodo.10846276).

### High-level assessment of genome sequence similarity

For both the *Mtb* (n=151) and E. coli (n= 50) datasets, FastANI version (v1.3) run with default parameters was used to estimate Average Nucleotide Identity (ANI) between complete genomes(27). For both the *Mtb* (n=151) and E. coli (n= 50) datasets, SourMash version (v4.8.2) was used to calculate the Jaccard Similarity of all unique 31 bp k-mers between each pair of complete genomes(40). To calculate the profile of all canonical 31 bp k-mers for each genome, the `sourmash sketch dna` command was run with the -p scaled=1 parameter. The scaled=1 parameter means that the comparison will be performed with the complete k-mer set (no downsampling). All k-mer signatures were then input into the `sourmash compare` command with default parameters. The Seaborn library was used to visualize heatmaps of estimates ANI and k-mer Jaccard Similarity across each bacterial population(41).

### Construction of the ***Mtb*** SV pan-genome graph

The *Mtb* SV pan-genome graph was built with `Minigraph` (v0.19) using H37Rv as the initial reference and with all 151 complete genome assemblies as input. `GFAtools` was used for all graph manipulations and reformatting of bubble region and node information (26). The Bandage software was used for visualization of the resulting *Mtb* SV pan-genome graph (42). All relevant code, with exact run parameters, can be found in the `7.Mtb.HybridAsms.Minigraph.smk` script (Runnable as a Snakemake workflow).

### Annotation of assemblies

All hybrid and short-read assemblies (*M. tuberculosis* and *E. coli*) were annotated with the `Bakta` (v4.8) and `PGAP` (v6.4) annotation pipelines. The GFF annotation files output by each annotation pipeline were used as input to all pan-genome analysis pipelines evaluated. All genome assemblies (Hybrid & Short-read) and their respective annotations used in this study are available on Zenodo (10.5281/zenodo.10846276)

### Pan-genome analysis tools & benchmarking

For the *Mtb* and *E. coli* datasets of hybrid and short-read *de novo* assemblies, we tested three commonly used pan-genome analysis pipelines, Panaroo, Roary, and PPanGGolin, and tested them in combination with two different genome annotation pipelines - Bakta and PGAP(30, 32, 43–45). For the Panaroo pipeline, we evaluated 6 different parameter combinations by varying both i) how strictly the accessory genome was filtered and ii) whether paralogs were considered separate genes or merged (--merge_paralogs). We evaluated Roary with 4 parameter combinations by varying i) whether paralogs are merged (-s), and ii) the amino acid minimum sequence identity threshold (-i) used to cluster CDS sequences. Only the default settings of the PPanGGOLiN pipeline were evaluated.

For each tool, we looked at the predicted number of i) total genes in the pan-genome, ii) core genes, and iii) accessory genes. All tools classified accessory genes as a gene which is found in less than 99% of all assemblies within the population, while a core gene was defined as any gene found in 99% or more of assemblies**. File S8** contains a summary of all pan-genome statistics produced during this analysis.

### Adjusting for nucleotide redundancy within CDS-based pan-genome estimates using the panqc software

The panqc nucleotide redundancy correction pipeline adjusts for CDS annotation discrepancies and adjusts for nucleotide redundancy within an estimated pan-genome with two steps. In step one, all genes absent at the CDS level are aligned to each corresponding assembly at the nucleotide level. Next, all genes are re-clustered and merged using a nucleotide k-mer based metric of nucleotide similarity. Cases where two or more genes are divergent at the protein level but highly similar at the nucleotide level will be merged into a single “nucleotide similarity gene cluster”. The panqc pipeline is compatible with both Roary and Panaroo when run with settings that merges paralogs. This is important to note because collapsing highly similar nucleotide sequences would only make sense if the user has already made the decision to collapse highly similar CDS sequences, regardless of genomic context.

### Phylogeny inference of ***E. coli*** isolates

A core genome alignment was generated for all 50 hybrid E. coli genomes using Panaroo run on assemblies with the following settings: --merge_paralogs, --clean-mode strict, --remove-invalid-genes, --alignment core, --aligner mafft. From the core gene alignment a maximum likelihood phylogeny was inferred using IQ-Tree with the general time reversible model(39). Relevant code can be found in the 6.Mtb.HybridAsms.BuildPhylogeny.smk script. The inferred phylogeny can be found on Zenodo (10.5281/zenodo.10846276).

### Inferring gene gain and loss events in the ***Mtb*** phylogeny

Ancestral state reconstruction of gene presence was performed for each node of the Mtb phylogeny using the ape R package’s Ancestral Character Estimation (ace) function(46) and modified code from the Panstripe package(31). After ancestral character estimation, the number of inferred gene gain and loss events were summarized for each branch. The described approach was then applied to pan-genome predictions of Panaroo (--merge_paralogs, --clean-mode strict, complete assemblies) before and after adjustment with the NRC pipeline (Default settings). To visualize the frequency of gene gain and loss events per branch of the tree separately, modified code from the Panstripe package was used. The RunPhyloAnalysis.WiPanstripe.V1.R script contains all code and modified functions to quantify gene gain and loss events based gene presence information (gene_presence_absence.csv) and the phylogeny of 151 Mtb isolates.

## Acknowledgements

We thank the members of the Farhat lab for helpful discussions and comments on the research project and manuscript. We acknowledge the International Science and Technology Center for their support in establishing the TB Portal agreement with Georgia and CRDF Global for their support in establishing the TB Portal agreements with Azerbaijan and Moldova. Portions of this research were conducted on the O2 High Performance Compute Cluster, supported by the Research Computing Group at Harvard Medical School.

## Data, Materials, and Software Availability

All SRA/ENA run accessions and associated metadata for all *M. tuberculosis* and *E*. coli isolates used in this study can be found in **File S2**. Code for data processing and analysis is available from the following GitHub repository, https://github.com/farhat-lab/mtb-pg-benchmarking-2024paper/. The Snakemake workflow engine was used for data processing(47). Data analysis was performed using python (v3.7) based Jupyter notebooks(48). The panqc software is available in the following GitHub repository, https://github.com/maxgmarin/panqc.

## Supplemental material

**File S1**. Supplemental Text, Figures, and Tables:

- **Figure S1.** Overview of k-mer jaccard similarity across *Mtb* genomes.
- **Figure S2.** Comparison of assembly characteristics between complete assemblies and short-read de novo assemblies.
- **Figure S3.** Evaluation of structural variation in TbD1 locus using a pan-genome graph.
- **Figure S4.** *Mtb* Pan-genome estimates for all pipeline-parameter combinations evaluated.
- **Figure S5.** Example of pseudogenes detected by PGAP in place of two CDSs by Bakta.
- **Figure S6.** Evaluating scenarios contributing to inferred absences in short-read assemblies between low and high BUSCO completeness scores.
- **Figure S7.** *Mtb* pan-genome estimates before and after NRC correction.
- **Figure S8.** *E. coli* pan-genome estimates before and after NRC correction.
- **Figure S9.** Overview of gene gain and loss events from *Mtb* gene ancestral state reconstruction.
- **Figure S10.** Overview of selected E. coli isolates from Shaw-2021.
- **Figure S11.** Overview of k-mer jaccard similarity across 50 E. coli genomes.
- **Table S1**. 18 bubble regions identified with at least 1 kb of novel sequence relative to H37Rv.
- **Table S2.** Annotation type summary for H37Rv reference genome annotations.
- **Table S3.** Summary table of feature types annotated by Bakta & PGAP across all 151 complete *Mtb* genome assemblies.
- **Table S4.** Summary of panqc nucleotide similarity clustering of *Mtb* pan-genome estimates.
- **Table S5.** Summary of panqc nucleotide similarity clustering of *E. coli* pan-genome estimates.

**File S2.** SRA/ENA genome assembly and sequencing accessions for long-read and short-read sequencing used in this study

**File S3.** Summary of *Mtb* isolate information and genome assembly (Short & hybrid) characteristics

**File S4.** Pairwise ANI estimates and k-mer jaccard similarities within datasets of hybrid assemblies (Mtb and E. coli)

**File S5.** Phylogeny of 151 *Mtb* isolates in newick format

**File S6.** *Mtb* SV pan-genome graph (GFA). Built from H37Rv and 151 hybrid assemblies using the minigraph (v0.19) algorithm.

**File S7.** k-mer based analysis of SV node sequence composition and uniqueness within *Mtb* SV pan-genome graph

**File S8.** *Mtb* & *E. coli* pan-genome estimates produced by pipeline-parameter combinations tested

**File S9.** Annotated feature type frequencies compared between PGAP & Bakta for 151 *Mtb* hybrid assemblies

**File S10.** Table of manually identified differences in H37Rv pseudogene annotation between PGAP & Bakta

**File S11.** Pseudogene annotation comparison visualization between the standard NCBI annotation, PGAP, and Bakta for the H37Rv reference sequence

**File S12.** Phylogeny of 50 *E. coli* isolates in newick format

**File S13.** Collection of all *Mtb* and *E. coli* assemblies and their respective annotations used in this work. This includes both hybrid (long + short read) and de novo short-read assemblies.

Files S5, S6, S12, S13 are hosted on Zenodo due to their non-standard file type or large file size, at https://zenodo.org/records/10846276

