## Supplementary material for "Analysis of the limited *M. tuberculosis* accessory genome reveals potential pitfalls of pan-genome analysis approaches": File S1 - Supplemental text and figures

**This PDF file includes:**

Summary of supplemental figures and tables
Summary of supplemental files
Supplemental Text (Methods & Results)
Supplemental References
Figures S1 to S11
Tables S1 to S5

#### Summary of supplemental figures & tables

**Figure S1.** Overview of k-mer jaccard similarity across *Mtb* genomes.

**Figure S2.** Comparison of assembly characteristics between complete assemblies and short-read de novo assemblies.

**Figure S3.** Evaluation of structural variation in TbD1 locus using a pan-genome graph.

**Figure S4.** *Mtb* Pan-genome estimates for all pipeline-parameter combinations evaluated.

**Figure S5.** Example of pseudogenes detected by PGAP in place of two CDSs by Bakta.

**Figure S6.** Evaluating scenarios contributing to inferred absences in short-read assemblies between low and high BUSCO completeness scores.

**Figure S7.** *Mtb* pan-genome estimates before and after NRC correction.

**Figure S8.** *E. coli* pan-genome estimates before and after NRC correction.

**Figure S9.** Overview of gene gain and loss events from *Mtb* gene ancestral state reconstruction.

**Figure S10.** Overview of selected E. coli isolates from Shaw-2021.

**Figure S11.** Overview of k-mer jaccard similarity across 50 E. coli genomes.

**Table S1**. 18 bubble regions identified with at least 1 kb of novel sequence relative to H37Rv.

**Table S2.** Annotation type summary for H37Rv reference genome annotations.

**Table S3.** Summary table of feature types annotated by Bakta & PGAP across all 151 complete *Mtb* genome assemblies.

**Table S4.** Summary of panqc nucleotide similarity clustering of *Mtb* pan-genome estimates.

**Table S5.** Summary of panqc nucleotide similarity clustering of *E. coli* pan-genome estimates.

#### Summary of supplemental files

**File S1**.  Supplemental Text, Figures, and Tables

**File S2.** SRA/ENA genome assembly and sequencing accessions for long-read and short-read sequencing used in this study

**File S3.** Summary of *Mtb* isolate information and genome assembly (Short & hybrid) characteristics

**File S4.** Pairwise ANI estimates and k-mer jaccard similarities within datasets of hybrid assemblies (Mtb and E. coli)

**File S5.** Phylogeny of 151 *Mtb* isolates in newick format

**File S6.** *Mtb* SV pan-genome graph (GFA). Built from H37Rv and 151 hybrid assemblies using the minigraph (v0.19) algorithm.

**File S7.** k-mer based analysis of SV node sequence composition and uniqueness within *Mtb* SV pan-genome graph

**File S8.** *Mtb* & *E. coli* pan-genome estimates produced by pipeline-parameter combinations tested

**File S9.** Annotated feature type frequencies compared between PGAP & Bakta for 151 *Mtb* hybrid assemblies

**File S10.** Table of manually identified differences in H37Rv pseudogene annotation between PGAP & Bakta

**File S11.** Pseudogene annotation comparison visualization between the standard NCBI annotation, PGAP, and Bakta for the H37Rv reference sequence

**File S12.** Phylogeny of 50 *E. coli* isolates in newick format

**File S13.** Collection of all *Mtb* and *E. coli* assemblies and their respective annotations used in this work. This includes both hybrid (long + short read) and de novo short-read assemblies.

**Files S5, S6, S12, S13** are hosted on Zenodo due to their non-standard file type or large file size, at https://zenodo.org/records/10846276

### Supplemental Results

**SR1 - Comparison of short and long-read characteristics**

We compared the assembly quality characteristics between the short-read and long read assemblies (**Table 1, Figure S1**). All complete assemblies consist of a single circular contig, while the short-read assemblies are consistently more fragmented with a median number of contigs of 116 (IQR: 106 - 132). The median genome size of the complete assemblies is 4,413 kb (4,407 kb - 4,421 kb). The cumulative length of the short-read assemblies are slightly lower than the complete genome assemblies, with a median length of 4,314 kb (IQR: 4,289 kb - 4,336 kb). Both technologies produced assemblies with a highly similar GC content, but the short-read assemblies varied slightly more: median GC content of the hybrid assemblies was 65.6%, IQR: 65.6%-65.6%, and for short-read assemblies the median GC content was 65.5%, IQR 65.5% - 65.6%. We found very high conservation in terms of the # of predicted ORFs for both short and complete assemblies. The median number of ORFs predicted for complete assemblies was 4074 (IQR: 4065 - 4088), while for short-read assemblies it was 4035 (IQR: 4022 - 4046). In summary, hybrid and short-read assembly characteristics were similar except that short-read assemblies systematically were less continuous, had lower cumulative length, and contained less predicted CDSs than complete assemblies.

**SR2 - Identifying novel sequence relative to the H37Rv reference genome**

Next, we wanted to evaluate what proportion of SV nodes in the graph represented sequences not present in the H37Rv reference genome. These SVs are of interest as they represent sequences that were lost in the ancestor to H37Rv but are present in other *Mtb* isolates. We found that only 160/2025 of evaluated SV nodes contained novel nucleotide content relative to H37Rv, with a cumulative length of 66,621 bp. Thus, only 5.5% of evaluated SV nodes within the graph represent novel sequence content relative to H37Rv. Thus, a majority of evaluated SV nodes in the graph are composed of either reconfiguration of already existing nucleotide content or deletions of regions that are present in the H37Rv reference genome.

The 160 SV nodes with novel sequence content relative to H37Rv were spread across 65 different bubble regions in the Mtb pan-genome. To focus our attention on larger SVs with novel sequences relative to H37Rv. From this list we were motivated to quantify the number of potentially large deletions (1 > kb) that occurred in the ancestor of the H37Rv strain. We generated a short list of 18 bubble regions that contained at least 1 kb of novel sequence relative to H37Rv (**Table S1**). The length of cumulative sequence that was novel relative to H37Rv ranged from 1153 bp to 7,900 bp per bubble region.

Notably, Bubble 193 matched the exact coordinates of a known structural variant within the Mtb population, Tbd1. The Tbd1 region is a deletion specific to lineages 2-4 of the *Mtb* population, and implicated in resistance to oxidative stress and hypoxia(1, 2). Our accurate detection of TbD1 serves as a positive control for the ability of our pan-genome graph approach to detect large structural differences between *Mtb* isolates. We further evaluated the graph representation of the Tbd1 deletion in terms of its genomic and phylogenetic context (**Figure S3**). Bubble 193 contains 3 different nodes and has three different observed configurations. The first observed configuration found within the graph is a complete deletion of the expected Tbd1 sequence (ΔTbD1), and this allele was exclusively found in all lineage 2-4 isolates. The second observed configuration was a completely intact TbD1 sequence, found in lineages 1,5, & 6. This distribution of ΔTbD1 and +TbD1 alleles exactly matches prior knowledge of the phylogenetic distribution of this deletion across lineages 1-6. Interestingly, a third configuration of bubble 193 was observed for the single lineage 8 isolate within our collection. The lineage 8 isolate was found to have an IS6110 insertion element disrupting the *mmpL6* gene, presumably resulting in a loss of function. This supports likely convergent disruption of the *mmpL6* gene in the *Mtb* phylogeny, in the first case through deletion and second case through transposon insertion.

**SR3 - Investigating differences between two bacterial genome annotation pipelines on H37Rv**

Motivated by the consistent decrease in the predicted pan-genome size when using PGAP, we explored the differences between the PGAP and Bakta annotation pipelines. To compare these softwares on an identical genome sequence, we first annotated the H37Rv reference genome with both Bakta and PGAP and then compared these annotation results with the official annotations (**Table X**). Although both tools predicted approximately the same number of total genes, we found that PGAP annotated 138 less protein coding sequences (CDSs) than Bakta, but annotated 150 more pseudogenes. PGAP only predicted 3,941 CDSs in the H37Rv reference, while Bakta predicted 4,079 total CDSs.

We manually inspected all pseudogenes in both annotations and found a high frequency of cases where a pseudogene identified by PGAP, is called as multiple CDSs by the Bakta software. In many cases a single pseudo will be annotated as multiple CDSs, resulting in a split CDS scenario. PGAP overall is more likely to call a sequence a pseudogene if it finds a suspect frameshift mutation is present. In contrast, Bakta is much more likely to call new CDSs in the position of a gene with a frameshift mutation. Notably, all the pan-genome analysis pipelines evaluated ignore pseudogenes, meaning any region annotated as a pseudogene will be ignored in downstream analysis and be treated as non-coding sequence. Thus, these annotation differences have consequences for pan-genome analyses that are based primarily on annotated CDS sequences.

### Supplemental Methods

#### Detailed *Mtb* hybrid assembly methods

##### A) PacBio Subreads Assembly & Polishing Pipeline

For all assemblies that were done with PacBio subreads, either from the RS II and Sequel II platforms, the following assembly processing was taken: All long reads (PacBio subreads) were assembled using Flye (v2.6). After assembly, Flye performed three rounds of iterative polishing of the genome assembly with long reads, producing a polished *de novo* long read assembly. If Flye identified the presence of a complete circular contig, Circlator (v1.5.5) was used to standardize the start of each assembly at the *DnaA* (*Rv0001*) locus. The paired-end Illumina WGS reads were trimmed with Trimmomatic (v0.39). Trimmed reads were aligned to the associated de novo long read assembly with BWA-MEM (v0.7.17). Duplicate reads were removed from the resulting alignments using PICARD (v2.22.5). Using the deduplicated alignments, Pilon (v1.23) was then used to correct SNSs and small INDELs in the de novo PacBio assembly. All relevant code, with exact run parameters, can be found in the `1.Mtb.Generate.HybridAsm.PacBioRSII.smk` script (Runnable as a Snakemake workflow).

##### B) ONT Assembly & Polishing Pipeline

For all assemblies made with Oxford Nanopore (Pore v9.4.1) the following assembly processing was taken: All ONT (v9.4.1) long reads ONT were assembled using Flye (v2.6). After the initial assembly, Flye performed three rounds of iterative polishing of the genome assembly with long reads, producing a polished de novo long read assembly. Next, the Medaka software was used to polish the de novo assembly with the ONT reads. If Flye identified the presence of a complete circular contig, Circlator (v1.5.5) was used to standardize the start of each assembly at the *DnaA* (*Rv0001*) locus. The paired-end Illumina WGS reads were trimmed with Trimmomatic (v0.39). Trimmed reads were aligned to the associated de novo long read assembly with BWA-MEM (v0.7.17). Duplicate reads were removed from the resulting alignments using PICARD (v2.22.5). Using the deduplicated alignments, Pilon (v1.23) was then used to correct SNSs and small INDELs in the de novo PacBio assembly. Next, the assembly was further polished with short reads using the PolyPolish (v0.5) pipeline. All relevant code, with exact run parameters, can be found in the `2.Mtb.Generate.HybridAsm.PacBioHiFi.smk` script (Runnable as a Snakemake workflow).

##### C) PacBio CCS (HiFi) Assembly & Polishing Pipeline:

For all assemblies generated from Sequel II CCS/HiFi reads the following assembly processing was taken: All PacBio HiFi reads were assembled using Flye (v2.9). After assembly, Flye performed three rounds of iterative polishing of the genome assembly with long reads, producing a polished de novo long read assembly. If Flye identified the presence of a complete circular contig, Circlator (v1.5.5) was used to standardize the start of each assembly at the *DnaA* (*Rv0001*) locus. The paired-end Illumina WGS reads were trimmed with Trimmomatic (v0.39). Trimmed reads were aligned to the associated de novo long read  assembly with BWA-MEM (v0.7.17). Duplicate reads were removed from the resulting alignments using PICARD (v2.22.5). Using the deduplicated alignments, Pilon (v1.23) was then used to correct SNSs and small INDELs in the de novo PacBio assembly. All relevant code, with exact run parameters, can be found in the `2.Mtb.Generate.HybridAsm.PacBioHiFi.smk` Snakemake workflow.

**Evaluating sequence content of SV nodes within *Mtb* SV pan-genome graph**

For the k-mer analysis of the graph, each node's sequence was broken down into all 31 bp subsequences. The mmh3 python library was used to generate the MurmurHash for all 31-mers within the dataset. Additionally, all k-mers were converted to their canonical k-mer such that a k-mer and its reverse complement would be given the same hash. For measurement of k-mer overlap between nodes and the rest of the graph the Jaccard Containment metric was used. Jaccard Containment is defined as the size of the union of two k-mer sets A and B divided by the length of set A. Put simply, the Jaccard containment is measured as the proportion of k-mers found within A that are also present in a second sequence, B.

For identification of SV nodes with unique sequence content, low k-mer overlap was defined as a k-mer Jaccard Containment of < 0.05. If a node was smaller than 31 bp it was excluded from this analysis. The ~500 SV nodes with length < 31 bp only have a cumulative length 6149 bp out of the 1,301,211 bp of SV nodes within the graph. As these short SV regions accounted for a small proportion (0.5%) of total SV node sequence, we expect that excluding them is unlikely to change our description of the Mtbc accessory genome.

To access the insertion sequence (IS) elements and phage sequences due to the fact that those classifications are the recognized MGEs in the Mtb genome. We compared the k-mer profile of each SV-node to all sequences annotated as related to "Insertion sequence elements & phages" in the H37Rv reference genome (Kapopoulou et al. 2011).

#### Supplemental Figures & Tables

**Figure S1**

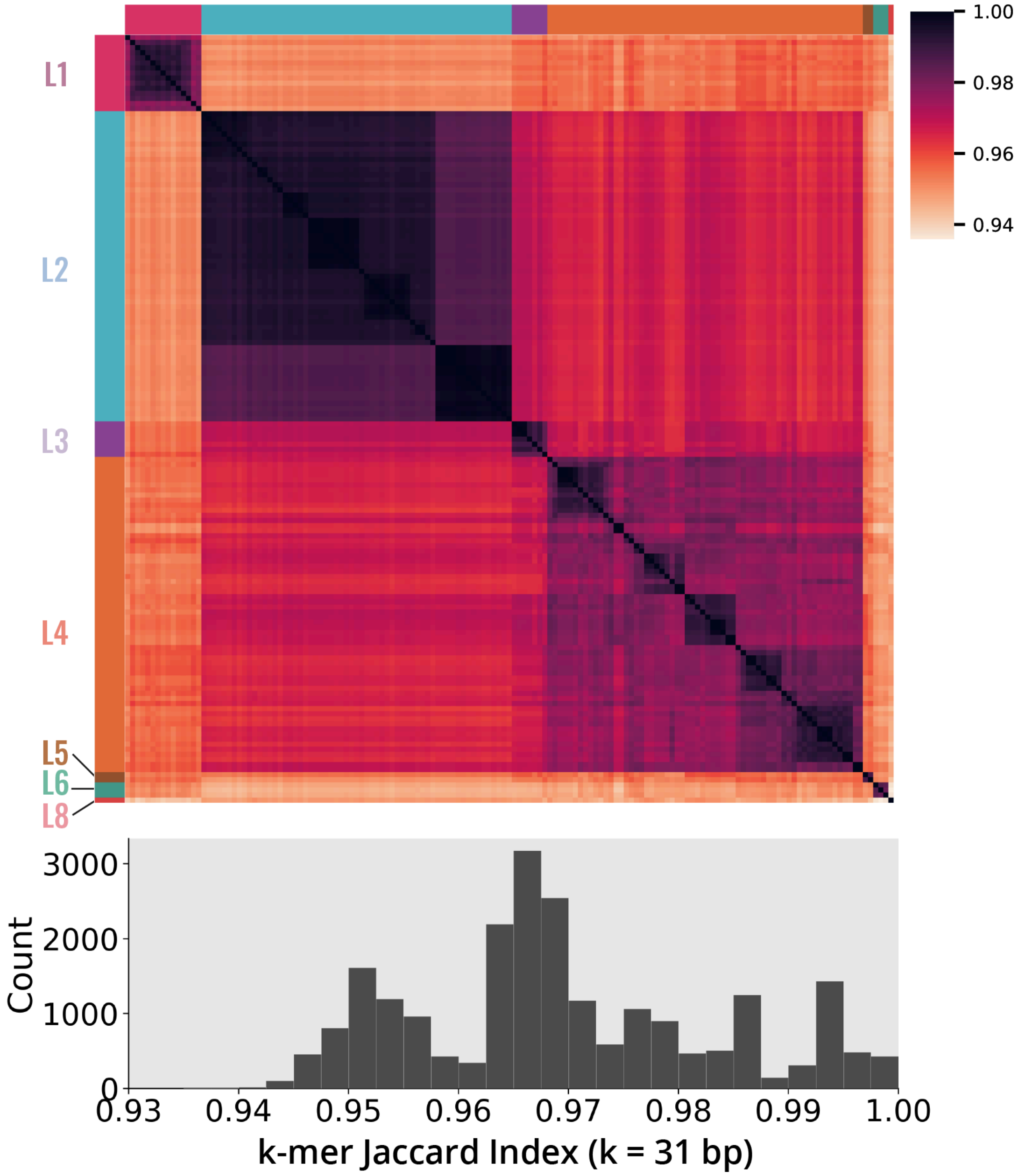


**Figure S1. Overview of k-mer jaccard similarity across *Mtb* genomes.** Heatmap and histogram of Jaccard Similarity of k-mers (k = 31 bp) across all pairs of complete *Mtb* genomes. The median k-mer jaccard similarity between all pairs was 0.97 (IQR: 0.96 - 0.98), while the most distant pair of genomes had a k-mer jaccard similarity of 0.94.

**Figure S2**

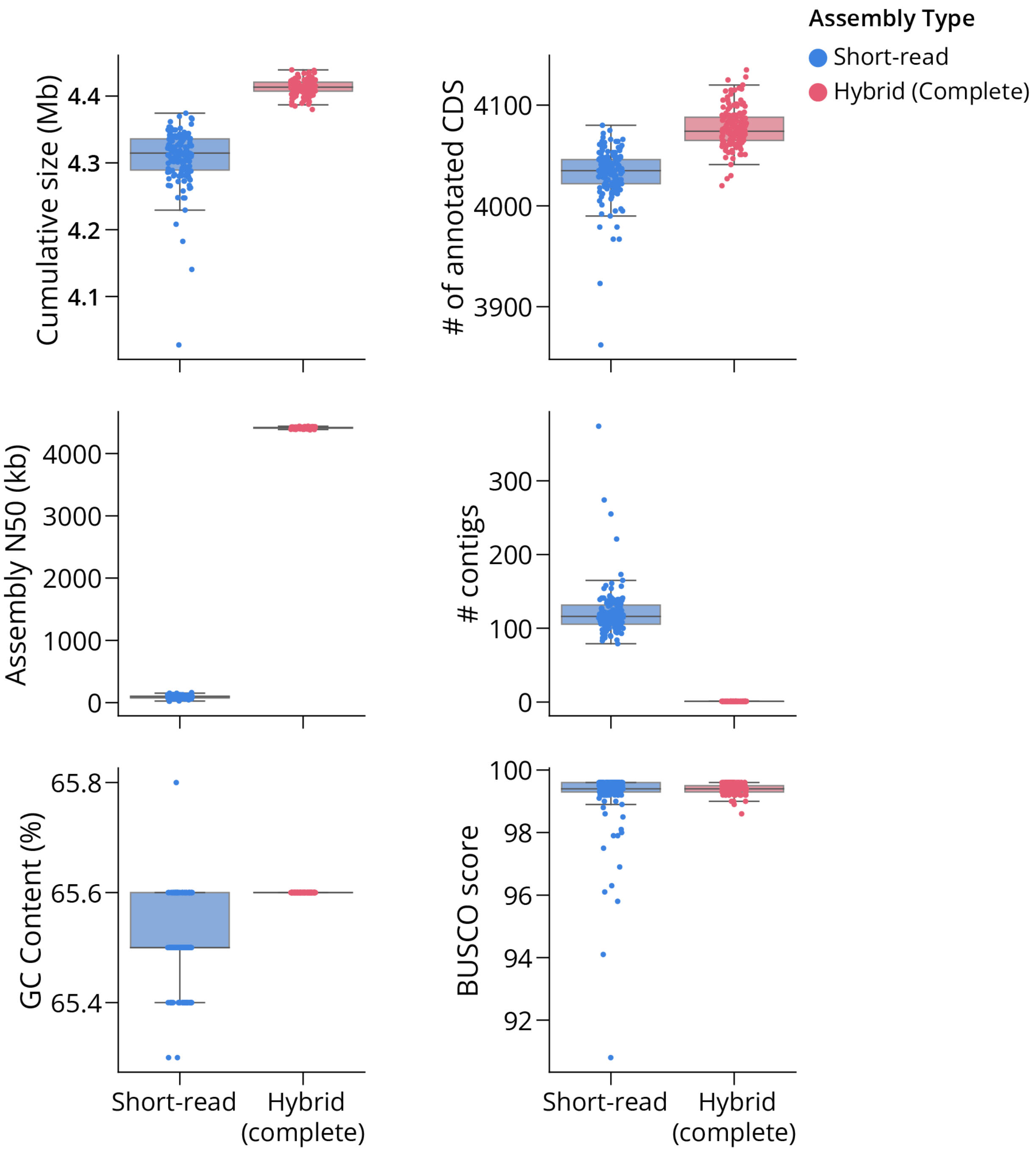


**Figure S2. Comparison of assembly characteristics between complete assemblies and short-read *de novo* assemblies.** Each of the 151 clinical *Mtb* isolates has both a complete hybrid assembly and a de novo short-read assembly available. The following characteristics were compared between the two assembly types: Cumulative assembly length, N50, GC Content, # of annotated CDSs with Bakta, # of total contigs, and BUSCO Completeness Score.

**Table S1**

| BubbleID | H37Rv coordinates | Overlapping H37Rv Genes | # SV Nodes | Cumulative SV Node Length | Cumulative length w/ LOW k-mer match to H37Rv |
|---|---|---|---|---|---|
| Bubble #31 | 334,243 - 338,669 | pe-pgrs3, pe-pgrs4 | 70 | 12,630 bp | 3,211 bp |
| Bubble #60 | 560425 - 561498 | umaA, pcaA | 2 | 2,074 bp | 1,001 bp |
| Bubble #102 | 917629 918744 | Rv0823c, desA1 | 4 | 2,210 bp | 1,095 bp |
| Bubble #116 | 1,053,645 - 1,053,787 | Rv0943c, Rv0944 | 2 | 4,541 bp | 4,399 bp |
| Bubble #129 | 1181855 - 1215202 | fadD14 - pe10 | 48 | 67,323 bp | 1,697 bp |
| Bubble #193 | 1761789 - 1761789 | mmpL6 | 3 | 3,511 bp | 2,153 bp |
| Bubble #221 | 1986625 - 1999613 | Rv1754c - Rv1765A | 116 | 53,885 bp | 7,500 bp |
| Bubble #255 | 2208005 - 2223187 | yrbE3A - Rv1979c | 26 | 29,094 bp | 4,496 bp |
| Bubble #265 | 2268723 - 2268723 | Rv2023A, Rv2024c | 17 | 8,232 bp | 5,518 bp |
| Bubble #362 | 3114093 - 3157869 | Rv2807 - cobO | 224 | 135,873 bp | 1,232 bp |
| Bubble #380 | 3251646 - 3252957 | ppsB | 2 | 2,441 bp | 1,130 bp |
| Bubble #396 | 3403272 - 3404591 | ctaD | 2 | 2,725 bp | 1,406 bp |
| Bubble #404 | 3486249- 3487460 | cyp141 | 4 | 3,722 bp | 1,153 bp |
| Bubble #410 | 3501223 - 3501723 | PPE50 | 13 | 1,859 bp | 1,310 bp |
| Bubble #412 | 3528674 - 3529255 | PPE53 | 6 | 5,517 bp | 1,642 bp |
| Bubble #426 | 3709332 - 3711855 | moaX - Rv3327 | 53 | 29,317 bp | 4,495 bp |
| Bubble #430 | 3730530 - 3735813 | PPE54 | 22 | 11,289 bp | 1,342 bp |
| Bubble #476 | 3937189 - 3938416 | ilvX | 2 | 2,470 bp | 1,243 bp |

**Table S1. 18 bubble regions identified with at least 1 kb of novel sequence relative to H37Rv.** Each bubble identified has at least 1 kb of sequence from SV nodes that have low k-mer match to the H37Rv reference genome. Coordinates of each bubble are given relative to the H37Rv genome assembly. Bubble #193 overlaps exactly with the known coordinates of the 2.1 kb TbD1 deletion, which is implicated in increased resistance to oxidative stress and hypoxia(2). Bubble #116 overlaps with a 4.4 kb sequence identified to be exclusive to lineage 8 of the MTBC(3).

**Figure S3**

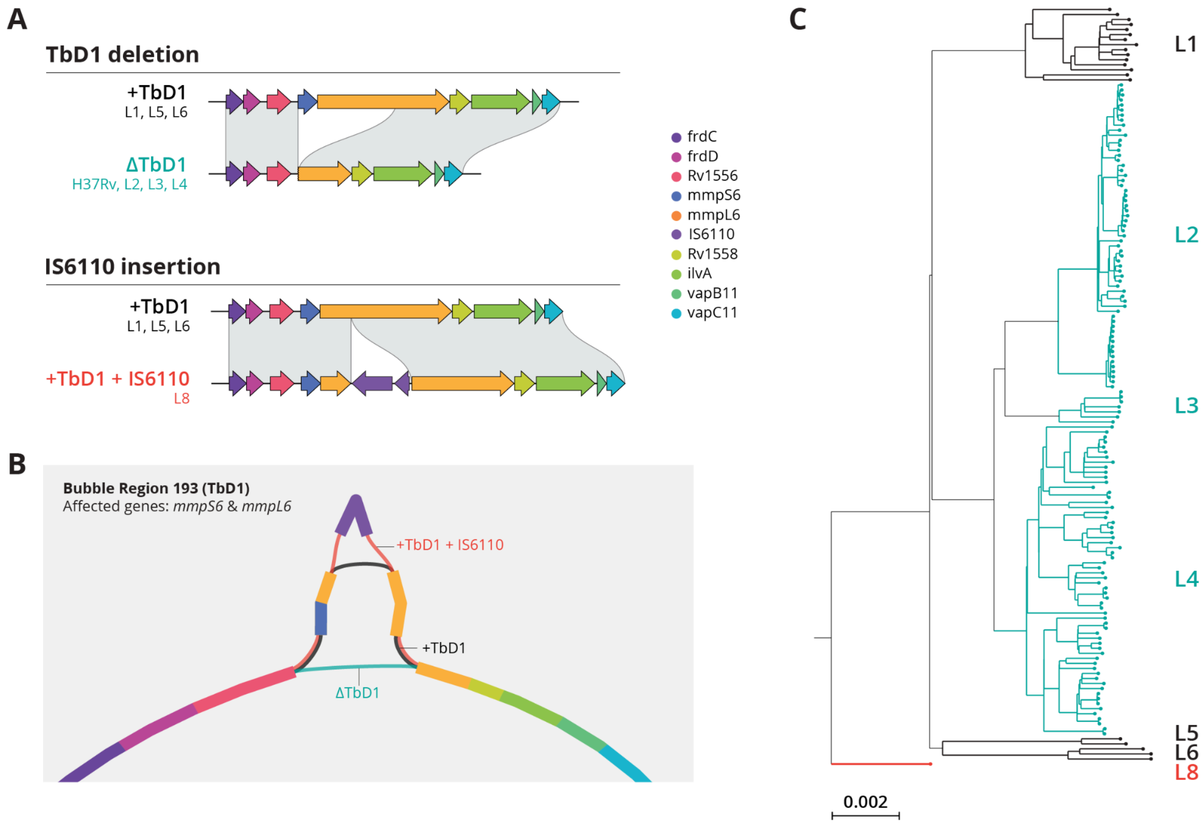


**Figure S3. Evaluation of structural variation in TbD1 locus using a pan-genome graph. A)** The TbD1 deletion refers to a deletion of the entire *mmpS6* gene and part of the *mmpL6* gene observed in lineages 2, 3 and 4 of the MTBC. Lineages 1, 5, and 6 contain the completely intact TbD1 locus (+TbD1). A third configuration of the locus is observed where an IS6110 transposable element has inserted in the mmpL6 gene (+TbD1- 6110), likely disrupting the protein function. **B)** Within the SV pan-genome graph, bubble region #193 represents these three configurations TbD1 locus. Bubble region #193 matches the exact coordinates of the known TbD1 deletion site in the H37Rv genome (`NC_000962.3:1,761,789`). **(C)** Mapping the presence of the three different configurations at the TbD1 locus onto the phylogeny of all isolates. There appears to be convergent loss of function of the mmpL6 gene in the *Mtb* phylogeny, one by deletion and the other by IS6110 insertion.

**Table S2**

| | Bakta | PGAP | Mycobrowser | NCBI annotation (NC_000962.3) |
|---|---|---|---|---|
| **Protein coding** | 4,073 | 3,967 | 4,018 | 3,906 |
| **Pseudogenes** | 6 | 296 | 13 | 30 |
| **RNA genes** | 70 | 49 | 142 | 68 |
| **Total** | 4,149 | 4,312 | 4,173 | 4,004 |

**Table S2. Annotation type summary for H37Rv reference genome annotations.**

**Table S3**

| | Bakta | PGAP |
|---|---|---|
| **Protein coding (CDS)** | 4,067 (IQR: 4,057 - 4,080) | 3,919 (IQR: 3,900 - 3,930) |
| **Pseudogenes** | 9 (IQR: 7 - 10) | 376 (IQR: 339 - 396) |
| **RNA genes** | 72 (IQR: 71 - 72) | 49 (IQR: 0) |
| **Total** | 4,147 (IQR: 4,138 - 4,160) | 4,340 (IQR: 4,320 - 4,356) |

**Table S3. Summary table of feature types annotated by Bakta & PGAP across all 151 complete Mtb genome assemblies.**

**Figure S4**

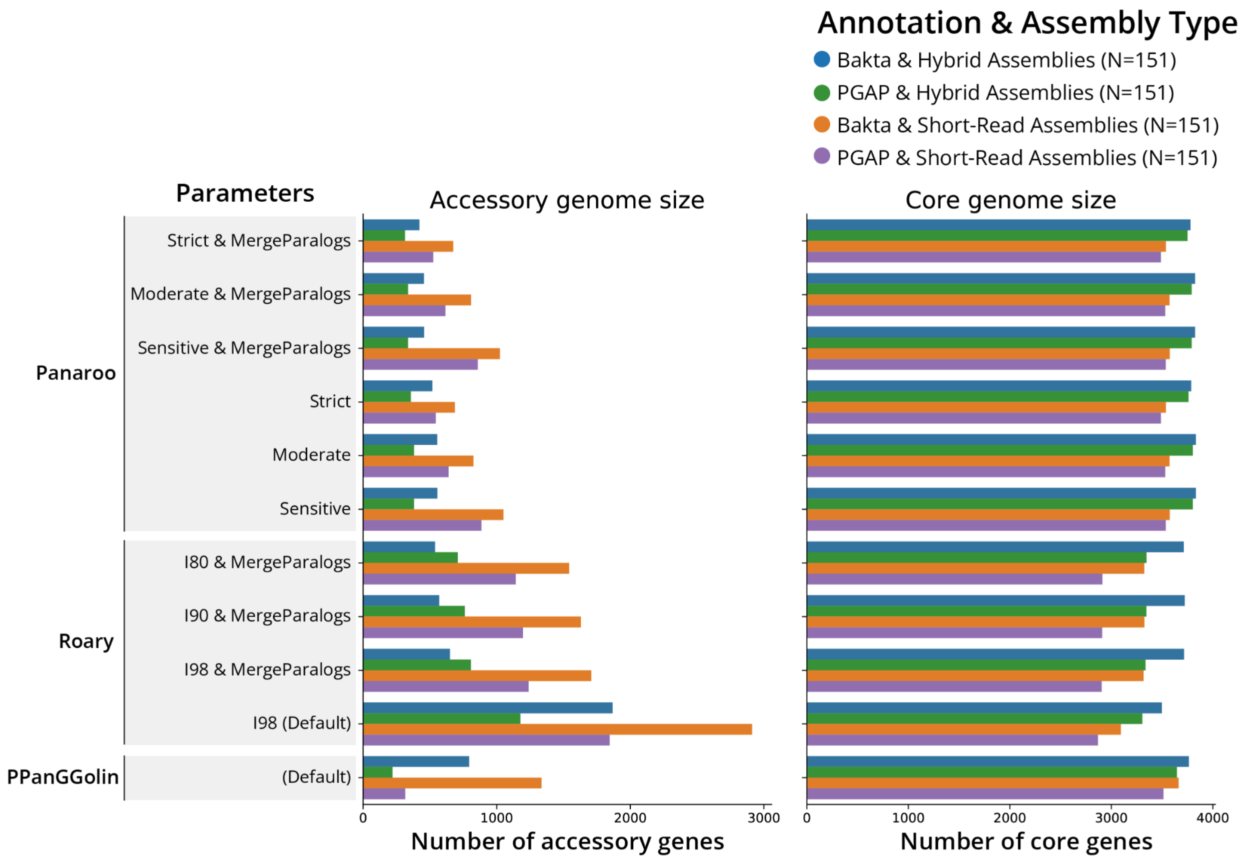


**FIgure S4.** ***Mtb*** **Pan-genome estimates for all pipeline-parameter combinations evaluated.** For each of the three softwares evaluated (Panaroo, Roary, PPanGGolin), different parameters were run in combination with varying annotation pipeline (PGAP, Bakta) and the assembly type used (Hybrid, Short-read). Panaroo was run with three different levels of sensitivity (Strict, Moderate, Sensitive) and with and without merging paralogs. Roary was run with variable amino acid clustering thresholds (80%, 90%, 98%) and with –mergeparalogs. PPanGGolin was run with default settings. The core gene cutoff used for all softwares was >= 99%.

**Figure S5**

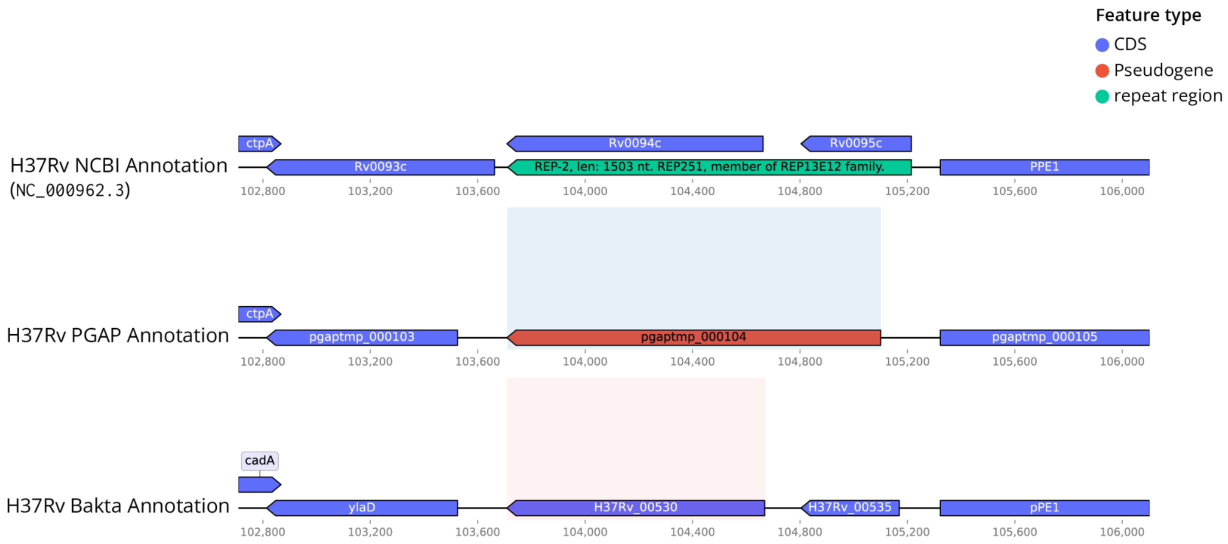


**FIgure S5. Example of pseudogenes detected by PGAP in place of two CDSs by Bakta.** In place of Rv0094c and Rv0095c, we observed that PGAP infers these two ORFs as a single pseudogene. The official NCBI H37Rv reference and Bakta both infer this sequence to contain two separate CDSs. Supplemental file X contains information regarding all 37 split CDS vs pseudogene discrepancies manually identified between H37Rv's Bakta and PGAP annotations.

**Figure S6**

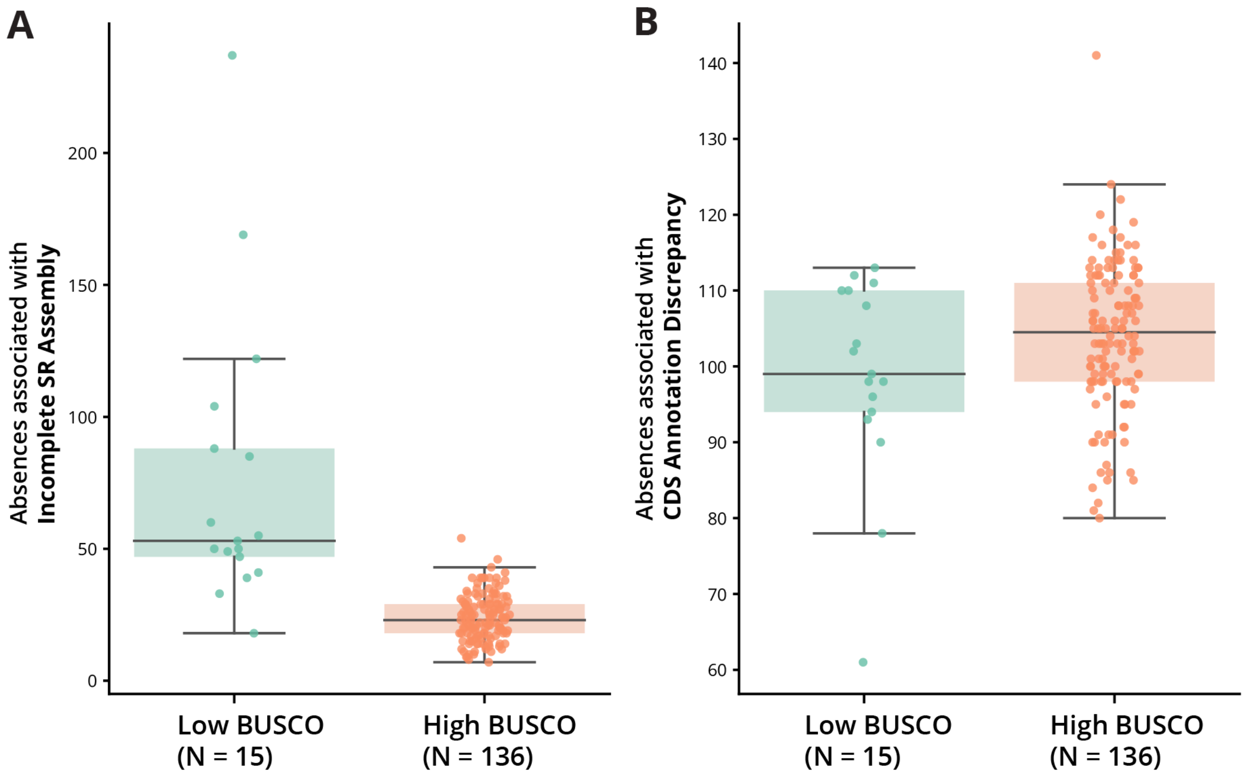


**Figure S6. Evaluating scenarios contributing to inferred absences in short-read assemblies between low and high BUSCO completeness scores.** All SR assemblies were classified as either low BUSCO assemblies (BUSCO < 0.99, N = 15) or high BUSCO assemblies (BUSCO >= 0.99, N = 136). Analysis of the frequency of absences related to "incomplete SR assembly" and "CDS annotation discrepancy" between low and high BUSCO assemblies was performed using the estimates generated by using Bakta annotation & Panaroo (`--clean-mode strict --merge_paralogs`). **A)** Boxplot of absences associated with "incomplete SR assembly" for both low BUSCO and high BUSCO SR assemblies. A low BUSCO SR assemblies were associated with a significant increase in inferred absences related to incomplete assembly (Mann-Whitney U test, p = 3.3e-9) **B)** Boxplot of absences associated with "CDS annotation discrepancies" for both low BUSCO and high BUSCO SR assemblies. No significant difference in the frequency of absences associated with "CDS annotation discrepancy" was observed between low and high BUSCO assemblies (Mann-Whitney U test, p = 0.12).

**Figure S7**

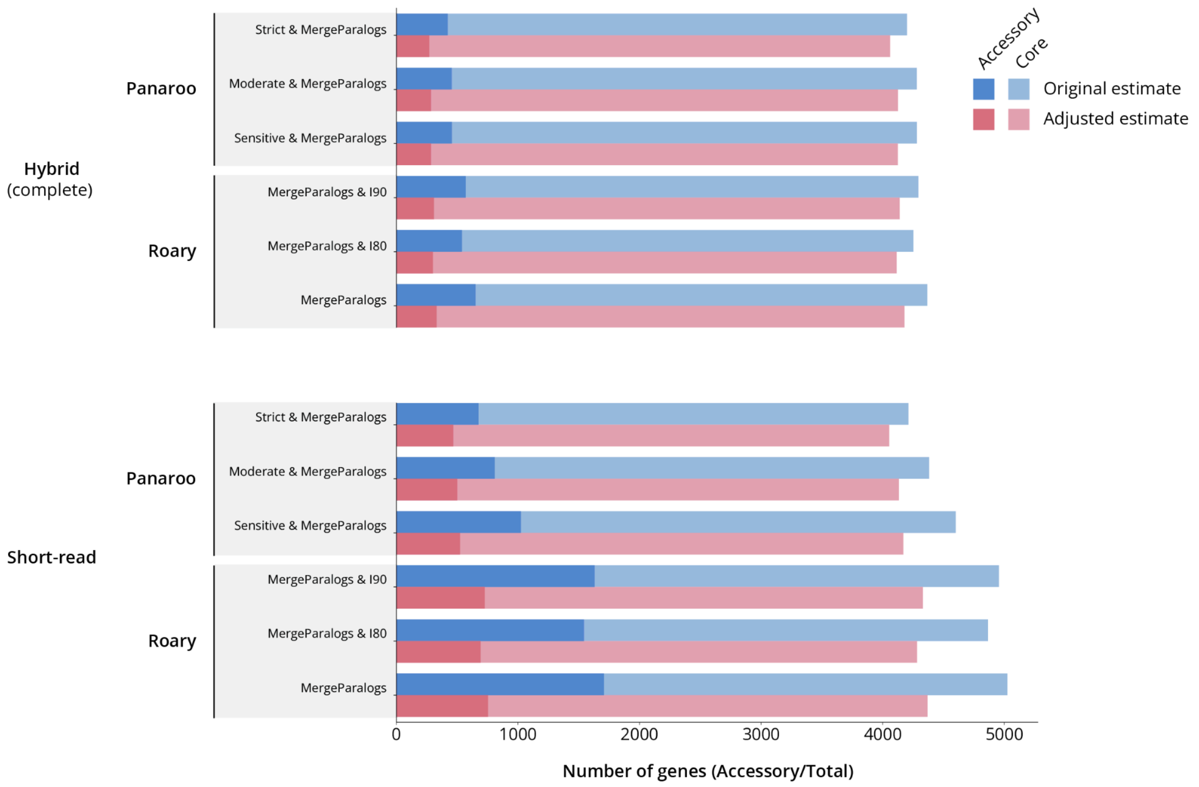


**Figure S7. *Mtb* pan-genome estimates before and after NRC correction.** The NRC pipeline was run on the pan-genome results for all Panaroo and Roary analyses that merged paralogs (--mergeparalogs). Core and accessory gene counts before and after NRC correction are shown in a stacked barplot. The solid color represents the accessory genome, while the transparent section of the bar represents the number of core genes inferred. The absolute bar height represents the total pan-genome size.

**Figure S8**

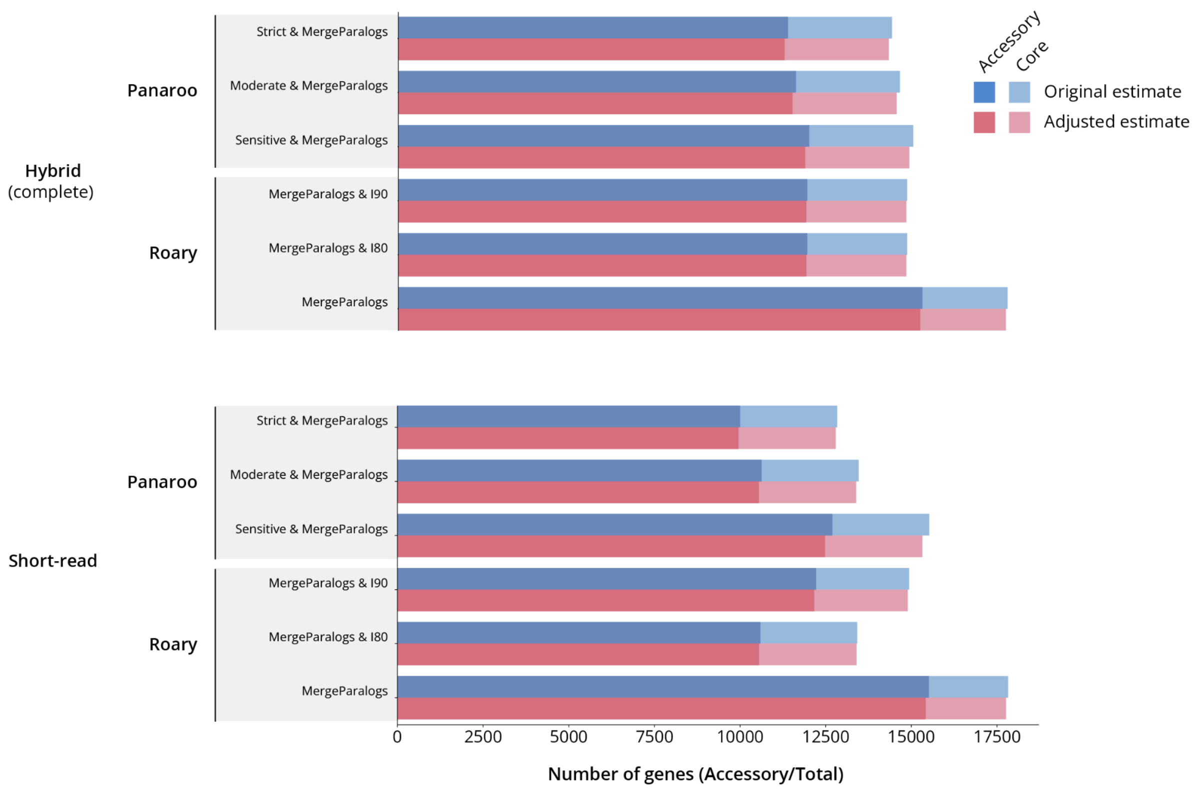


**Figure S8. *E. coli* pan-genome estimates before and after NRC correction.** The NRC pipeline was run on the pan-genome results for all Panaroo and Roary analyses that merged paralogs (--mergeparalogs). Core and accessory gene counts before and after NRC correction are shown in a stacked barplot. The solid color represents the accessory genome, while the transparent section of the bar represents the number of core genes inferred. The absolute bar height represents the total pan-genome size.

#### Figure S9

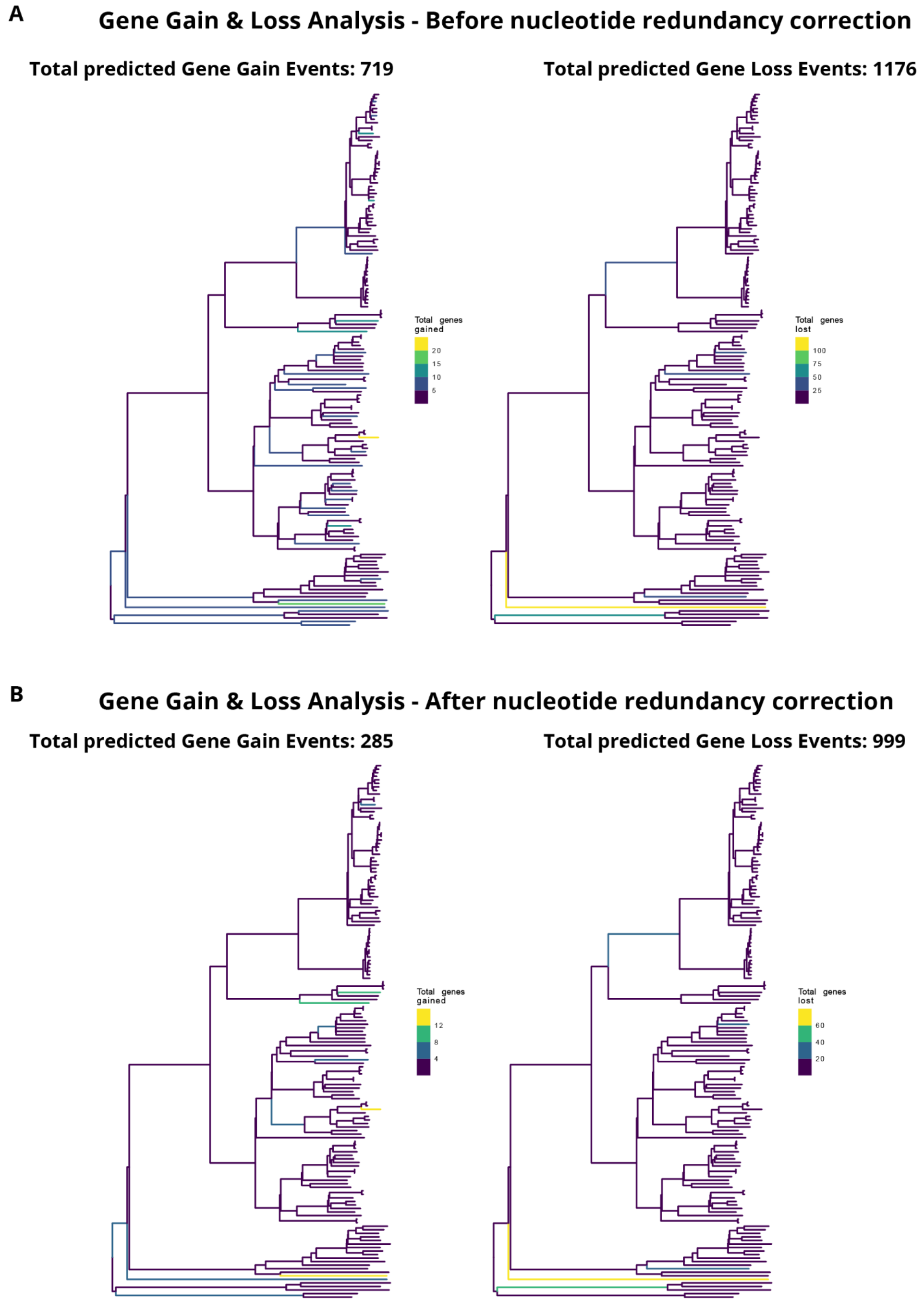


**Figure S9. Overview of gene gain and loss events from *Mtb* gene ancestral state reconstruction.** Inferred phylogenetic distribution of gene gain and loss events across 151 *Mtb* genomes before **A)** and after **B)** adjusting for nucleotide redundancy is shown. Ancestral character estimation (using the `ape` R package) was used to identify the frequency of gene gain and gene loss events across the Mtb phylogeny. Ancestral character estimation was based on gene presence information estimated by Panaroo run on all complete assemblies (`--merge_paralogs, --clean-mode strict`).

**Figure S10**

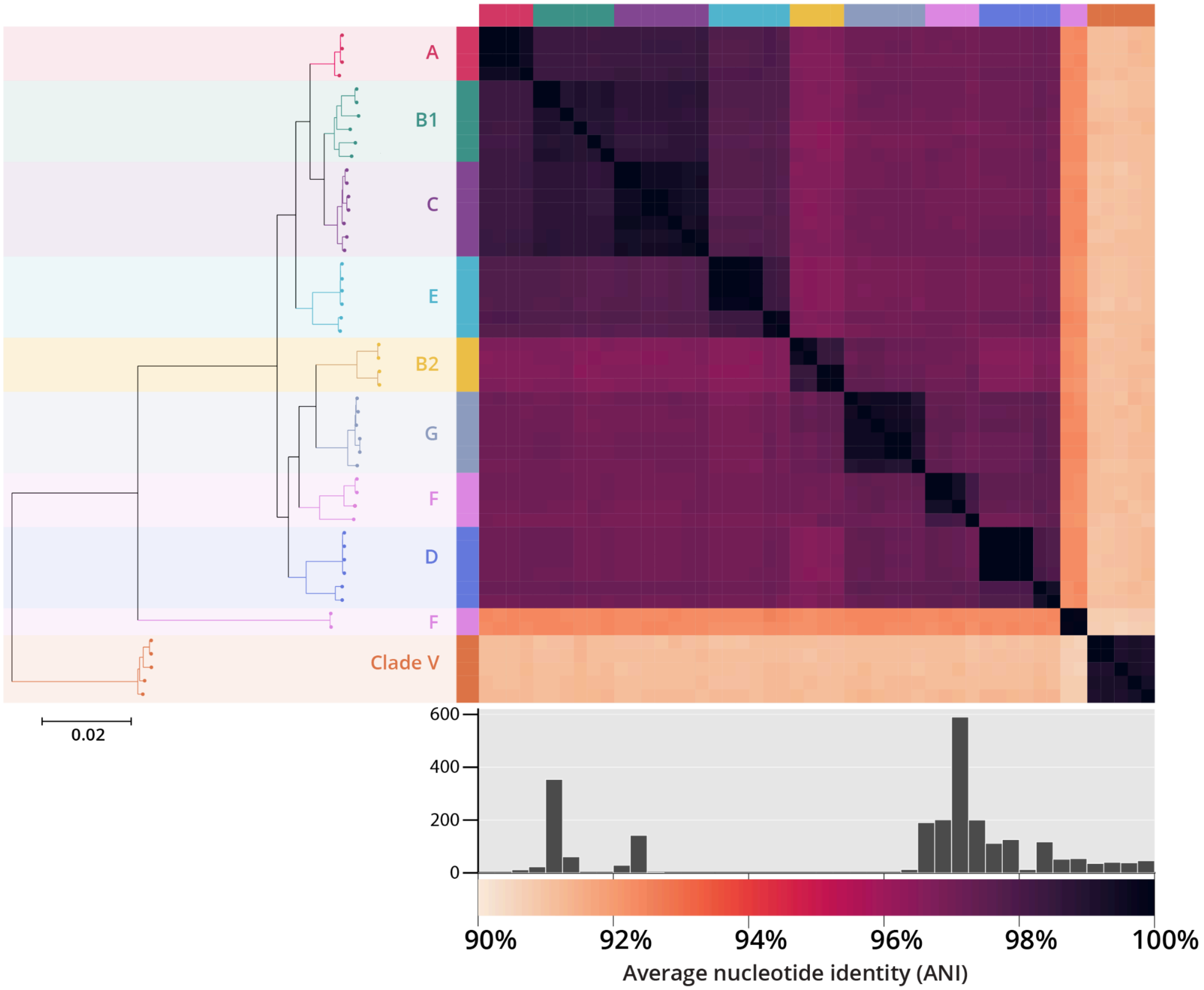


**Figure S10. Overview of selected E. coli isolates from Shaw-2021.** 50 E. coli isolates from across 9 phylogroup classifications were selected as a representative dataset from Shaw et. al. 2021. A heatmap of estimated ANI values within the population is displayed with the core genome phylogeny displayed to the left. Each genome is colored by *E. coli* phylotype classifications identified in Shaw et. al. FastANI was used to estimate ANI between all pairs of genomes. The histogram of all pairwise ANI estimates is shown below. The median ANI across all pairs was 97.1% (IQR: 92.4% - 97.6%).

**Figure S11**

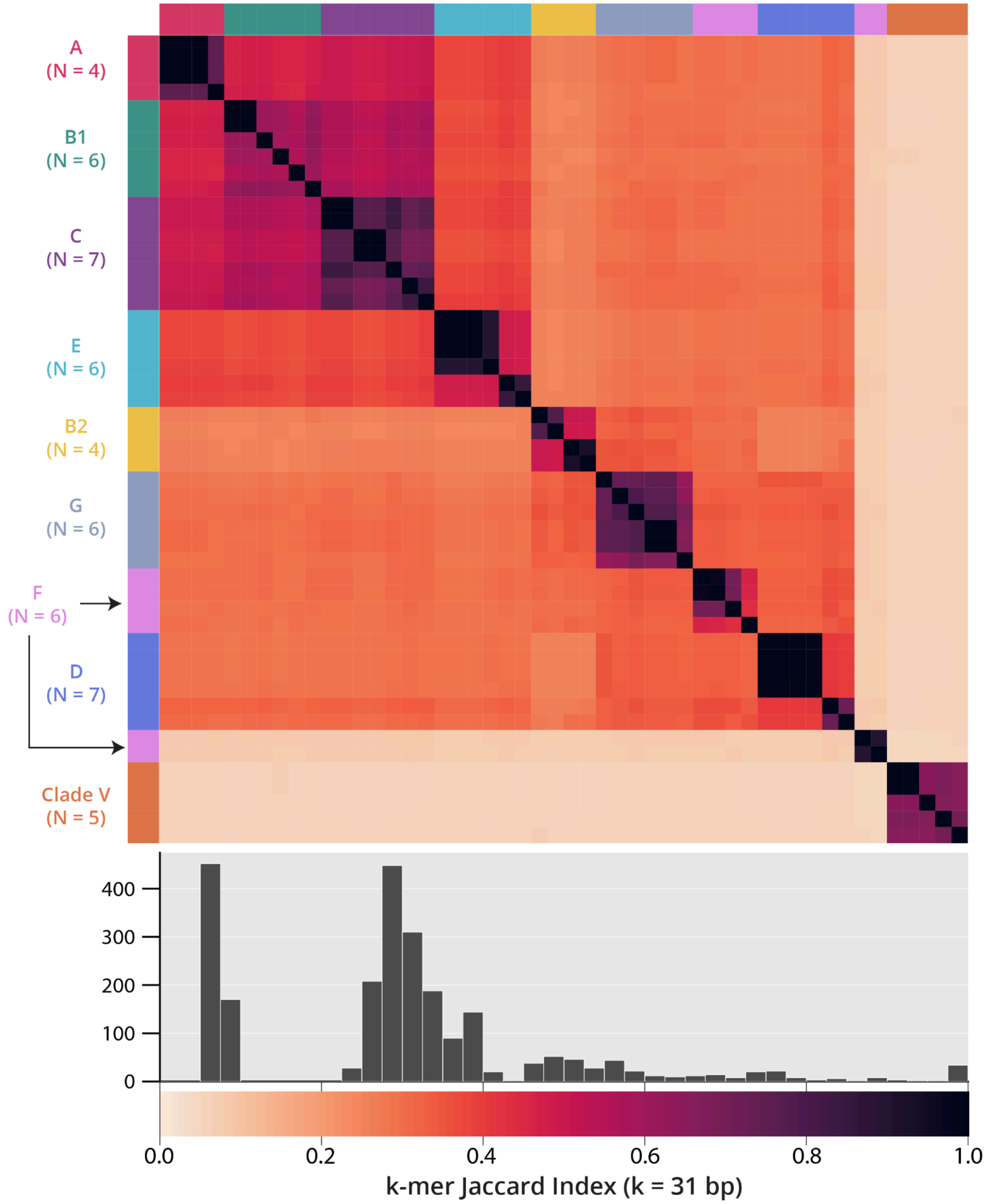


**Figure S11. Overview of k-mer jaccard similarity across 50 *E. coli* genomes.** Heatmap and histogram of Jaccard Similarity of k-mers (k = 31 bp) across all pairs of hybrid *E. coli* genomes. The median k-mer jaccard similarity between all pairs was 0.29 (IQR: 0.09 - 0.36), while the most distant pair of genomes had a k-mer jaccard similarity of 0.05. Each genome is colored by *E. coli* phylotype classifications identified in Shaw et. al.

**Table S4**

| Assembly type | Pipeline | Parameters | # nucleotide similarity clusters | # genes merged | Mean # of genes per cluster |
|---|---|---|---|---|---|
| Complete | Panaroo | Strict & MergeParalogs | 111 | 250 | 2.25 |
| Complete | Panaroo | Moderate & MergeParalogs | 120 | 275 | 2.29 |
| Complete | Panaroo | Sensitive & MergeParalogs | 120 | 276 | 2.3 |
| Complete | Roary | MergeParalogs & Identity 98 (Default) | 139 | 347 | 2.5 |
| Complete | Roary | MergeParalogs & Identity 90 | 107 | 272 | 2.54 |
| Complete | Roary | MergeParalogs & Identity 80 | 91 | 235 | 2.58 |
| Short-read | Panaroo | Strict & MergeParalogs | 75 | 233 | 3.11 |
| Short-read | Panaroo | Moderate & MergeParalogs | 102 | 351 | 3.44 |
| Short-read | Panaroo | Sensitive & MergeParalogs | 151 | 582 | 3.85 |
| Short-read | Roary | MergeParalogs & Identity 98 (Default) | 393 | 1074 | 2.73 |
| Short-read | Roary | MergeParalogs & Identity 90 | 372 | 1017 | 2.73 |
| Short-read | Roary | MergeParalogs & Identity 80 | 352 | 953 | 2.71 |

**Table S4. Summary of panqc nucleotide similarity clustering of *Mtb* pan-genome estimates.** For each estimate analyzed with panqc, the following relevant statistics are shown after nucleotide similarity clustering using a minimum 31-mer overlap (maximum Jaccard Similarity observed) of 0.8: 1) The number of total nucleotide similarity (NS) clusters identified, 2) total number of genes merged into NS clusters, and 3) the mean number of genes belonging to each NS cluster.

**Table S5**

| Assembly type | Pipeline | Parameters | # nucleotide similarity clusters | # genes merged | Mean # of genes per cluster |
|---|---|---|---|---|---|
| Complete | Panaroo | Strict & MergeParalogs | 78 | 171 | 2.19 |
| Complete | Panaroo | Moderate & MergeParalogs | 79 | 174 | 2.2 |
| Complete | Panaroo | Sensitive & MergeParalogs | 90 | 205 | 2.28 |
| Complete | Roary | MergeParalogs & Identity 98 (Default) | 49 | 104 | 2.12 |
| Complete | Roary | MergeParalogs & Identity 90 | 24 | 49 | 2.04 |
| Complete | Roary | MergeParalogs & Identity 80 | 24 | 49 | 2.04 |
| Short-read | Panaroo | Strict & MergeParalogs | 35 | 79 | 2.26 |
| Short-read | Panaroo | Moderate & MergeParalogs | 57 | 128 | 2.25 |
| Short-read | Panaroo | Sensitive & MergeParalogs | 146 | 348 | 2.38 |
| Short-read | Roary | MergeParalogs & Identity 98 (Default) | 60 | 129 | 2.15 |
| Short-read | Roary | MergeParalogs & Identity 90 | 38 | 80 | 2.11 |
| Short-read | Roary | MergeParalogs & Identity 80 | 22 | 45 | 2.05 |

**Table S5. Summary of NRC nucleotide similarity clustering of *E. coli* pan-genome estimates.** For each estimate analyzed with panqc, the following relevant statistics are shown after nucleotide similarity clustering using a minimum 31-mer overlap (maximum Jaccard Similarity observed) of 0.8: 1) The number of total nucleotide similarity (NS) clusters identified, 2) total number of genes merged into NS clusters, and 3) the mean number of genes belonging to each NS cluster.
