## Supplementary material for "Analysis of the limited *M. tuberculosis* accessory genome reveals potential pitfalls of pan-genome analysis approaches": File S11

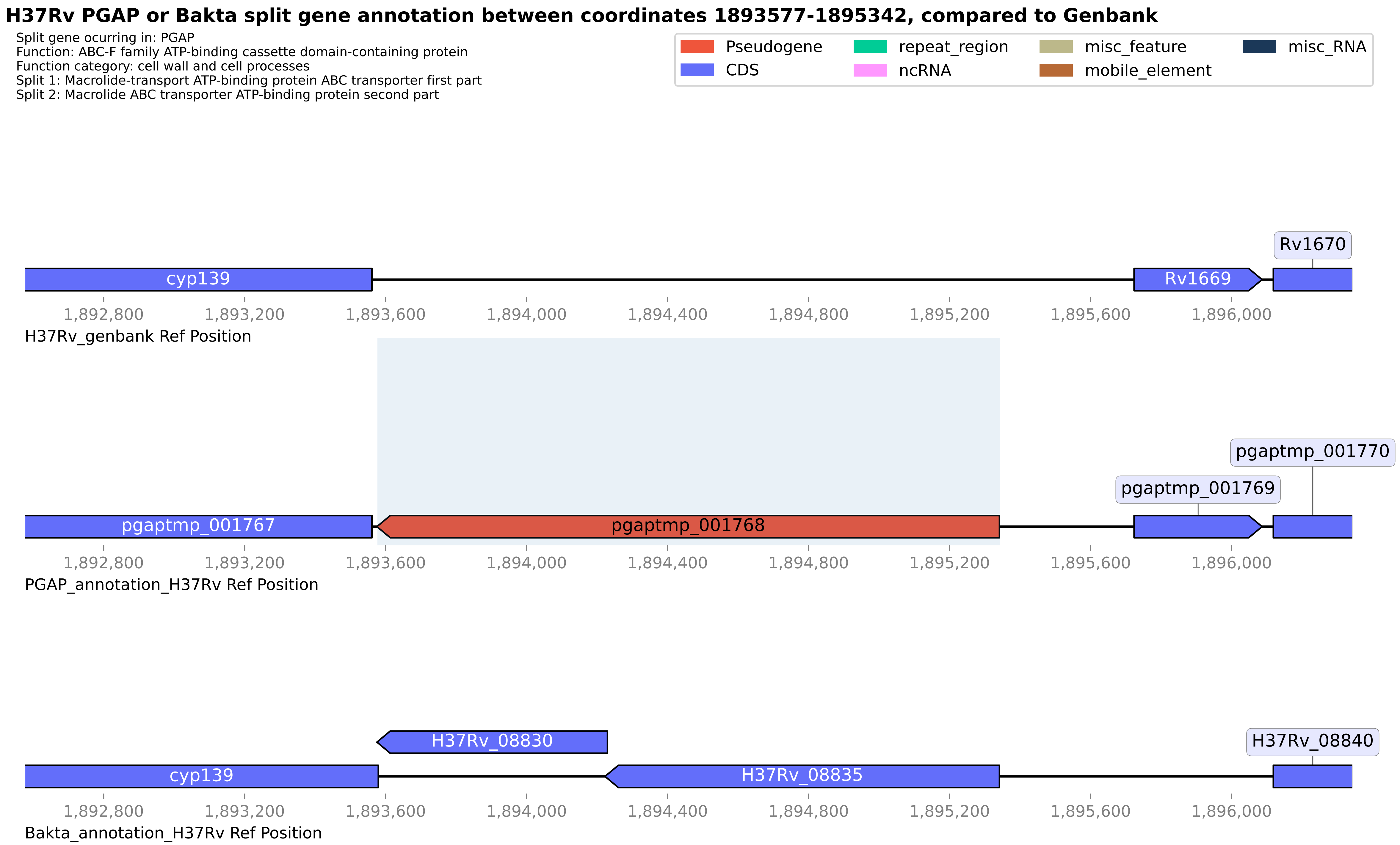

H37Rv PGAP or Bakta split gene annotation between coordinates 1893577-1895342, compared to Genbank
Split gene ocurring in: PGAP
Function: ABC-F family ATP-binding cassette domain-containing protein
Function category: cell wall and cell processes
Split 1: Macrolide-transport ATP-binding protein ABC transporter first part
Split 2: Macrolide ABC transporter ATP-binding protein second part
Pseudogene
CDS
repeat_region
ncRNA
misc_feature
mobile_element
misc_RNA
Rv1670
cyp139
Rv1669
1,892,800
1,893,200
1,893,600
1,894,000
1,894,400
1,894,800
1,895,200
1,895,600
1,896,000
H37Rv_genbank Ref Position
pgaptmp_001770
pgaptmp_001769
pgaptmp_001767
pgaptmp_001768
PGAP_annotation_H37Rv Ref Position
H37Rv_08830
H37Rv_08840
cyp139
H37Rv_08835
Bakta_annotation_H37Rv Ref Position

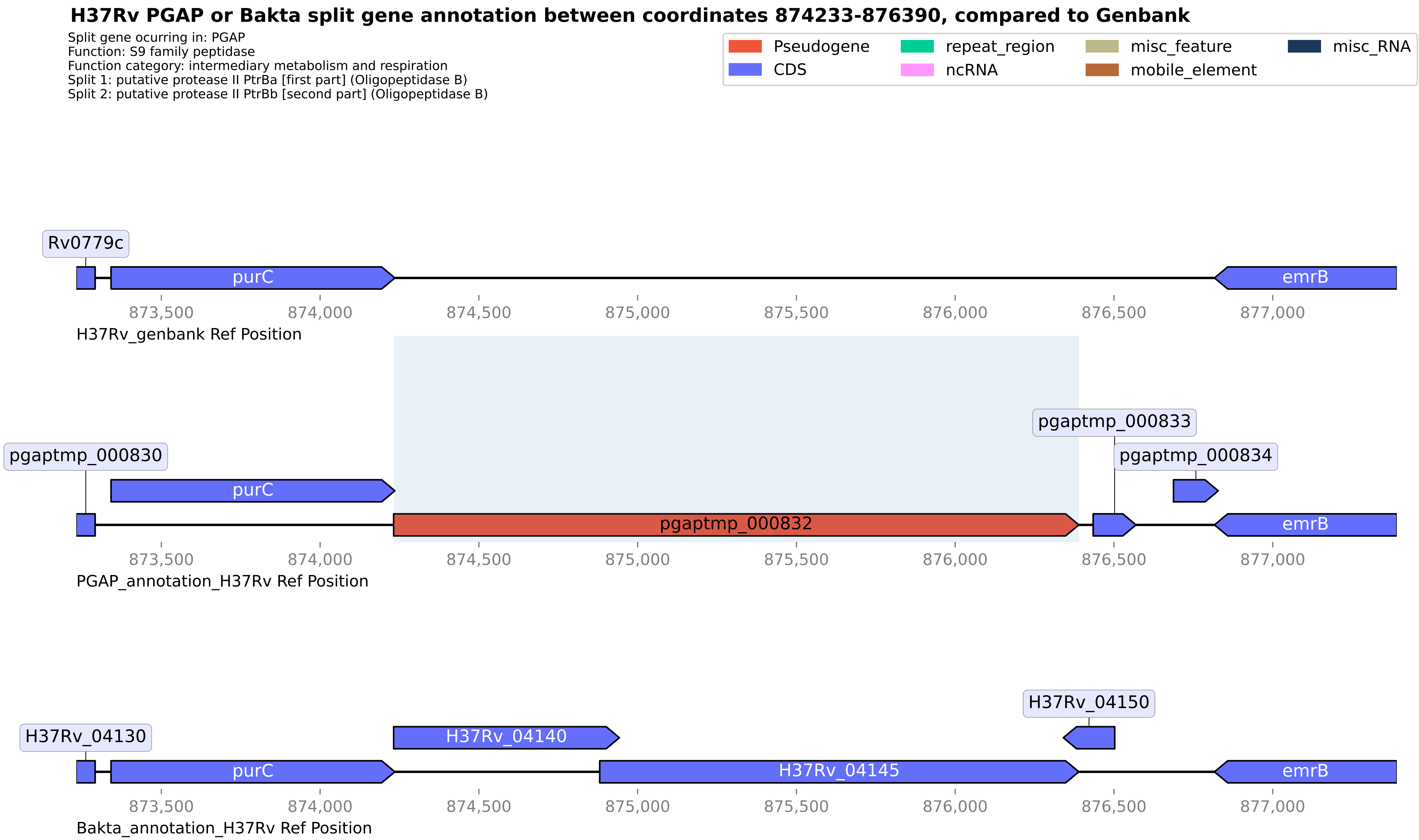

H37Rv PGAP or Bakta split gene annotation between coordinates 874233-876390, compared to Genbank
Split gene ocurring in: PGAP
Function: S9 family peptidase
Function category: intermediary metabolism and respiration
Split 1: putative protease II PtrBa [first part] (Oligopeptidase B)
Split 2: putative protease II PtrBb [second part] (Oligopeptidase B)
Pseudogene
CDS
repeat_region
ncRNA
misc_feature
mobile_element
misc_RNA
Rv0779c
purC
emrB
873,500
874,000
874,500
875,000
875,500
876,000
876,500
877,000
H37Rv_genbank Ref Position
pgaptmp_000830
pgaptmp_000833
pgaptmp_000834
pgaptmp_000832
PGAP_annotation_H37Rv Ref Position
H37Rv_04130
H37Rv_04140
H37Rv_04145
H37Rv_04150
Bakta_annotation_H37Rv Ref Position

**H37Rv PGAP or Bakta split gene annotation between coordinates 2881409-2882147, compared to Genbank**

Split gene ocurring in: PGAP
Function: DUF2652 domain-containing protein
Function category: conserved hypotheticals
Split 1: DUF2652 domain-containing protein
Split 2: Uncharacterized protein Rv2561/Rv2562

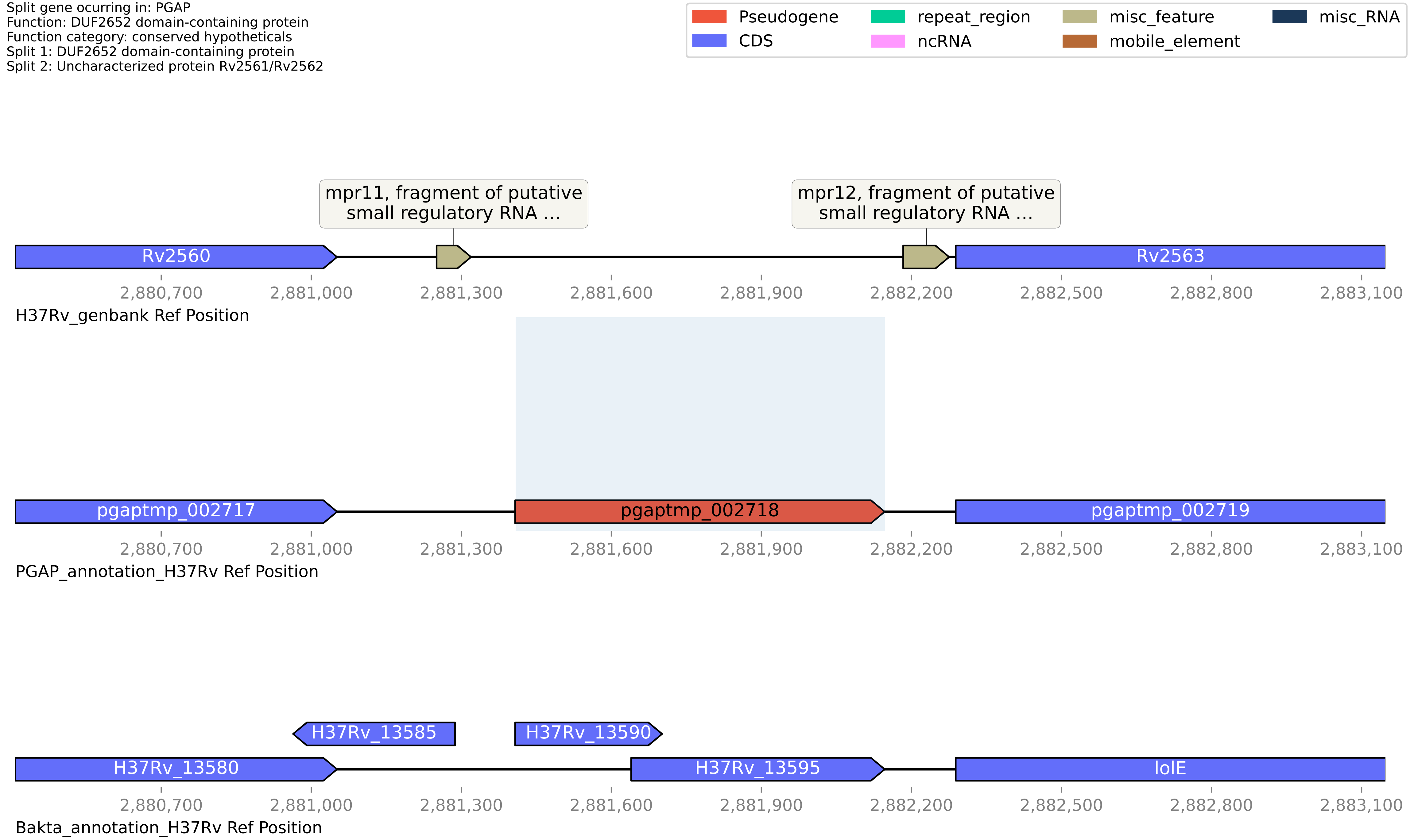

**H37Rv PGAP or Bakta split gene annotation between coordinates 3435718-3436295, compared to Genbank**

Split gene ocurring in: Bakta
Function: LLM class flavin-dependent oxidoreductase ssud
Function category: conserved hypotheticals
Split 1: LLM class flavin-dependent oxidoreductase
Split 2: LLM class flavin-dependent oxidoreductase

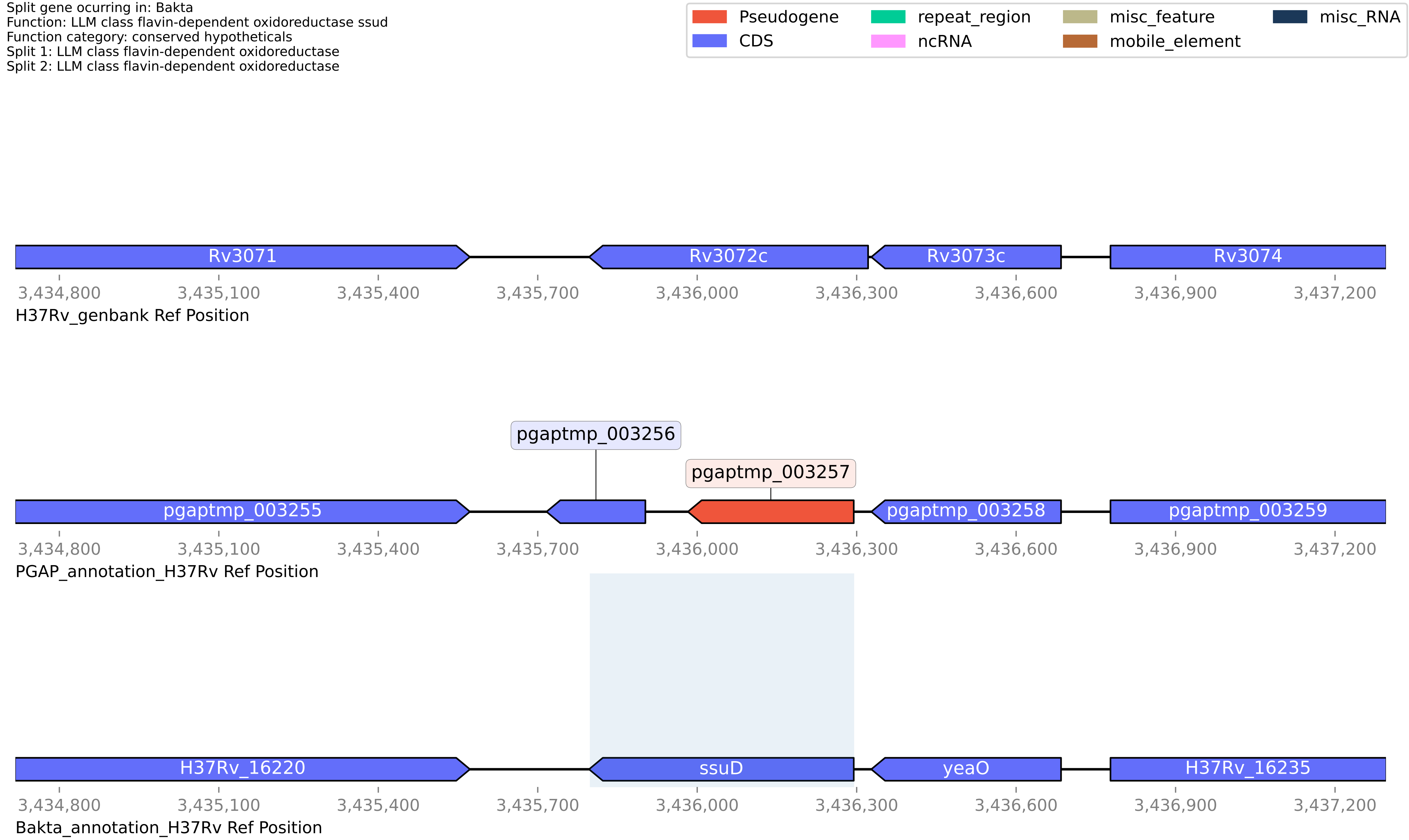

#### H37Rv PGAP or Bakta split gene annotation between coordinates 3800017-3801463, compared to Genbank

Split gene ocurring in: PGAP
Function: ISNCY family transposase
Function category: insertion seqs and phages
Split 1: Transposase and inactivated derivatives, IS5 family
Split 2: Transposase

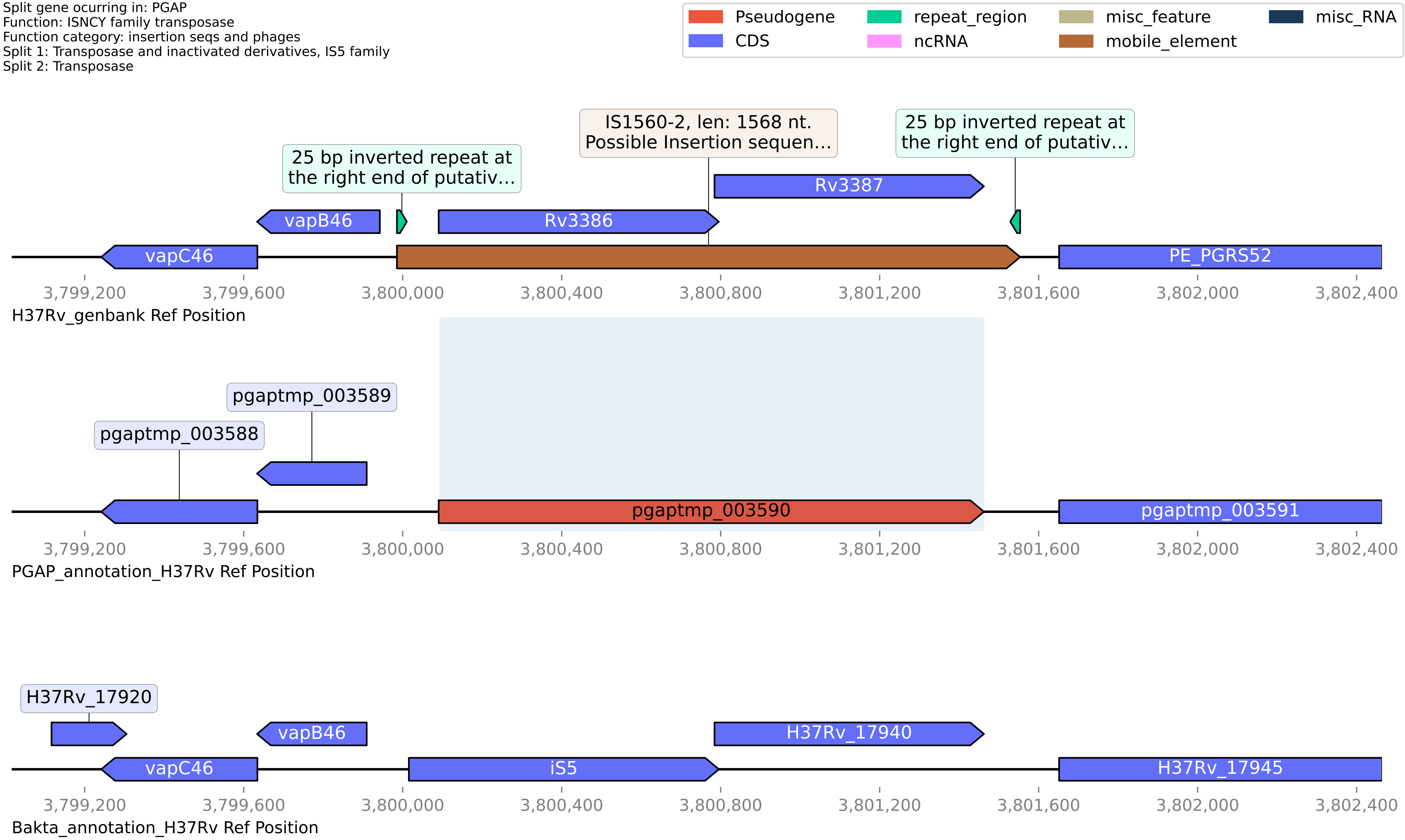

**H37Rv PGAP or Bakta split gene annotation between coordinates 711536-712719, compared to Genbank**

Split gene ocurring in: PGAP
Function: galT
Function category: intermediary metabolism and respiration
Split 1: galactose-1-phosphate uridylyltransferase
Split 2: Galactose-1-phosphate uridylyltransferase

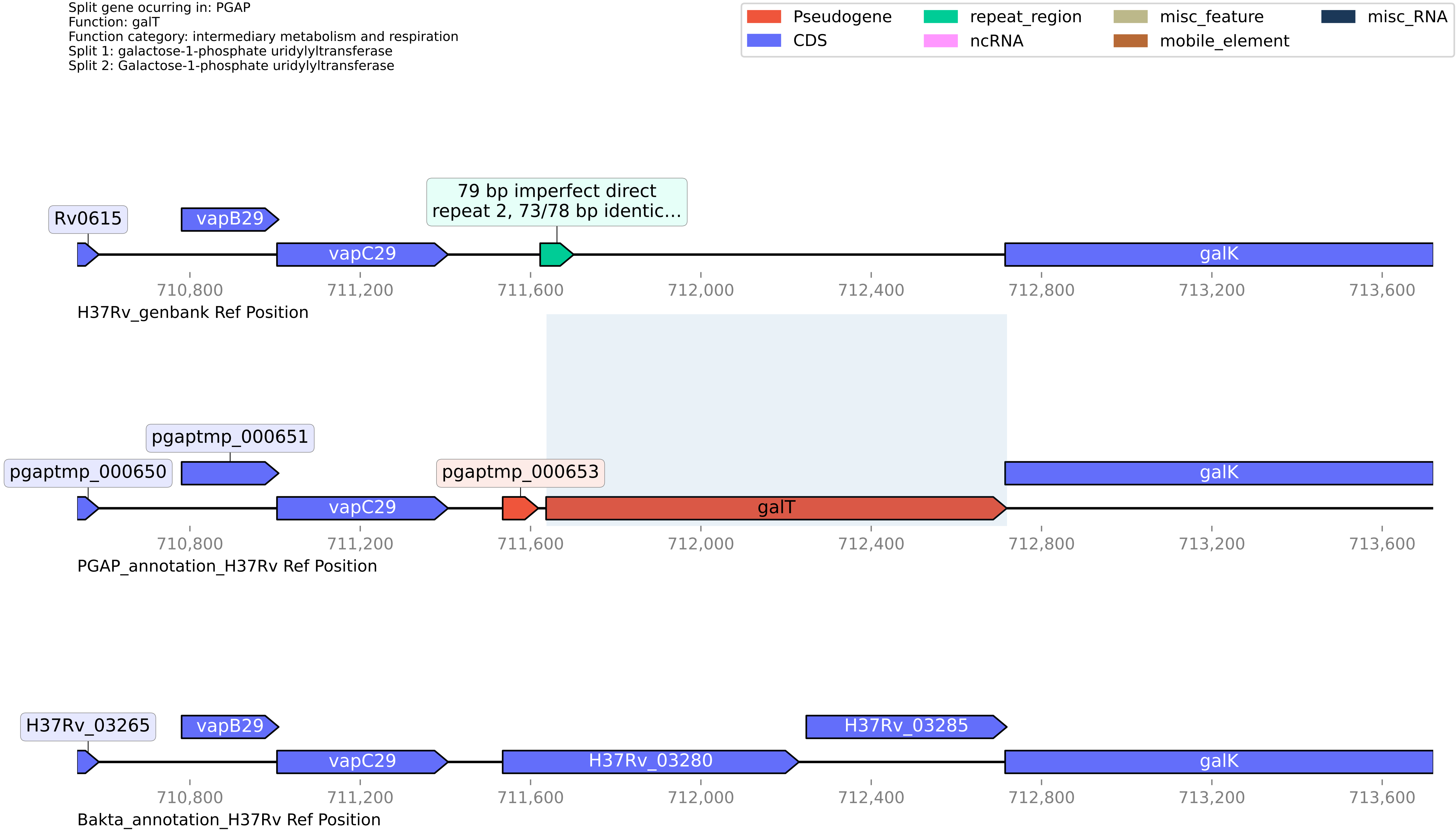

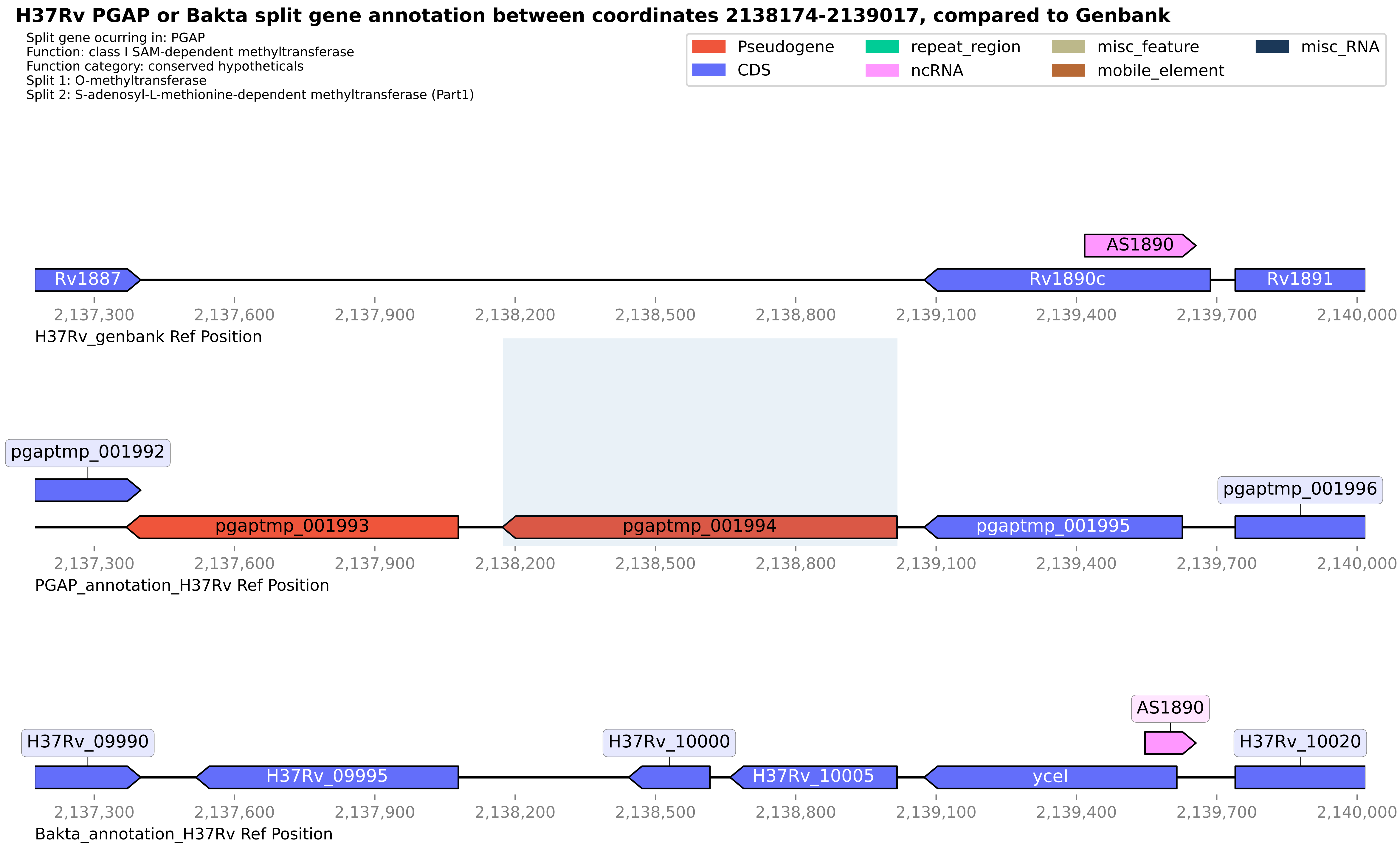

H37Rv PGAP or Bakta split gene annotation between coordinates 2138174-2139017, compared to Genbank
Split gene ocurring in: PGAP
Function: class I SAM-dependent methyltransferase
Function category: conserved hypotheticals
Split 1: O-methyltransferase
Split 2: S-adenosyl-L-methionine-dependent methyltransferase (Part1)
Pseudogene
CDS
repeat_region
ncRNA
misc_feature
mobile_element
misc_RNA
AS1890
Rv1887
Rv1890c
Rv1891
2,137,300
2,137,600
2,137,900
2,138,200
2,138,500
2,138,800
2,139,100
2,139,400
2,139,700
2,140,000
H37Rv_genbank Ref Position
pgaptmp_001992
pgaptmp_001993
pgaptmp_001994
pgaptmp_001995
pgaptmp_001996
PGAP_annotation_H37Rv Ref Position
H37Rv_09990
H37Rv_09995
H37Rv_10000
H37Rv_10005
ycel
AS1890
H37Rv_10020
Bakta_annotation_H37Rv Ref Position

**H37Rv PGAP or Bakta split gene annotation between coordinates 179319-181029, compared to Genbank**

Split gene ocurring in: PGAP
Function: PE-PPE domain-containing protein
Function category: PE/PPE
Split 1: PE family protein
Split 2: PE-PGRS family protein

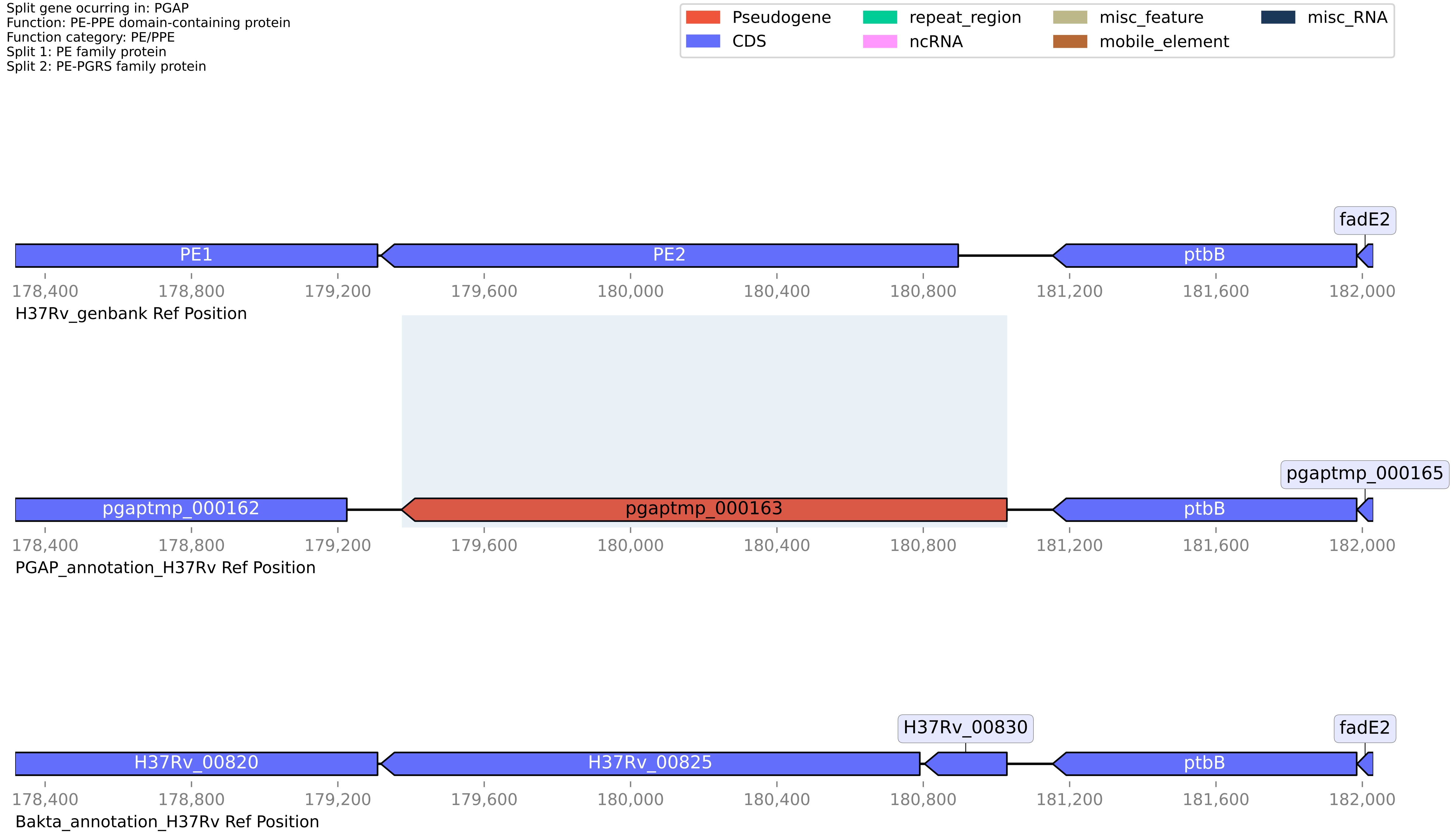

#### H37Rv PGAP or Bakta split gene annotation between coordinates 2500923-2501632, compared to Genbank

Split gene ocurring in: PGAP
Function: 2OG-Fe(II) oxygenase
Function category: conserved hypotheticals
Split 1: proline hydroxylase
Split 2: DUF2086 domain-containing protein

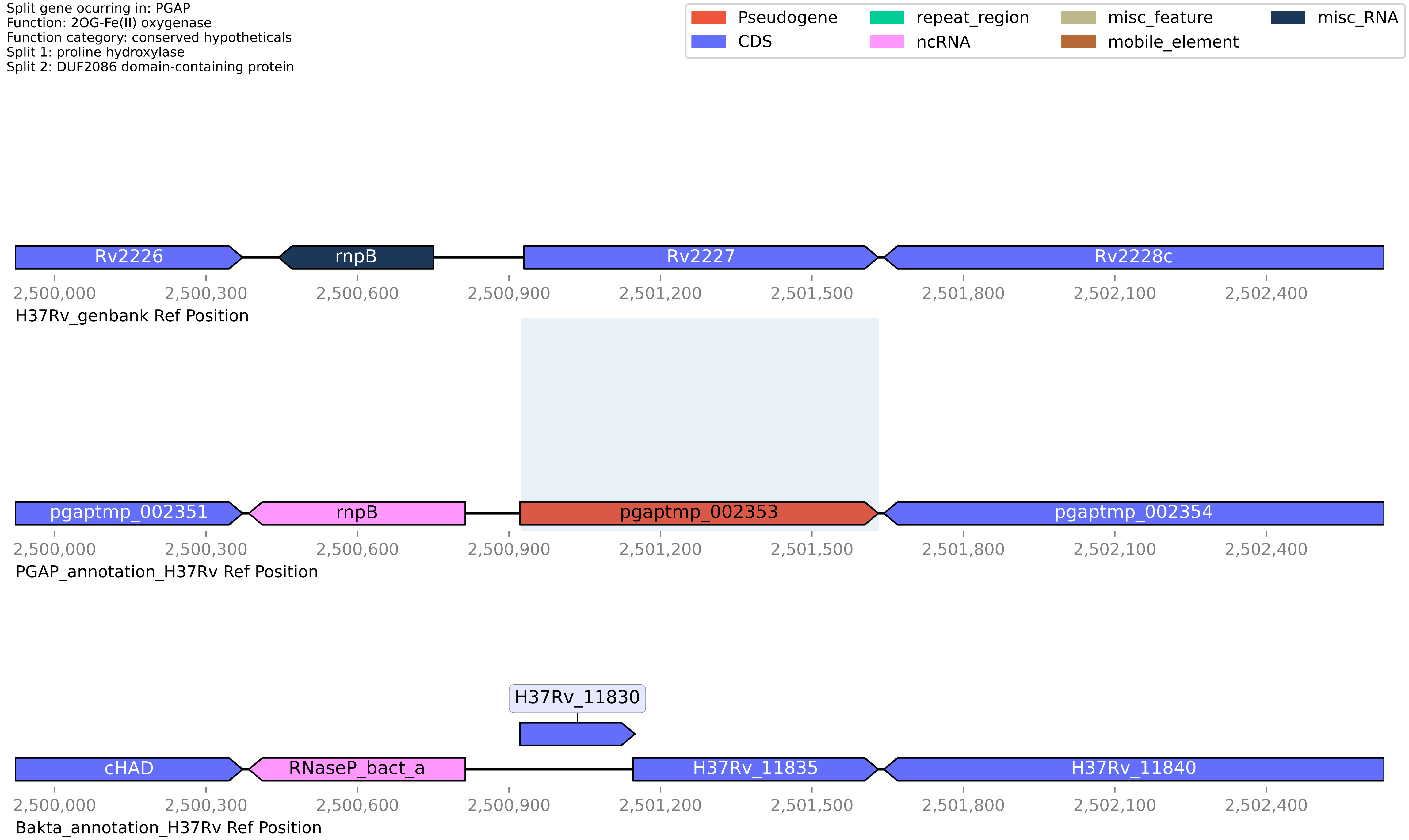

#### H37Rv PGAP or Bakta split gene annotation between coordinates 4192179-4193245, compared to Genbank

Split gene ocurring in: PGAP
Function: NAD(P)/FAD-dependent oxidoreductase
Function category: intermediary metabolism and respiration
Split 1: NAD(P)/FAD-dependent oxidoreductase
Split 2: Oxidoreductase

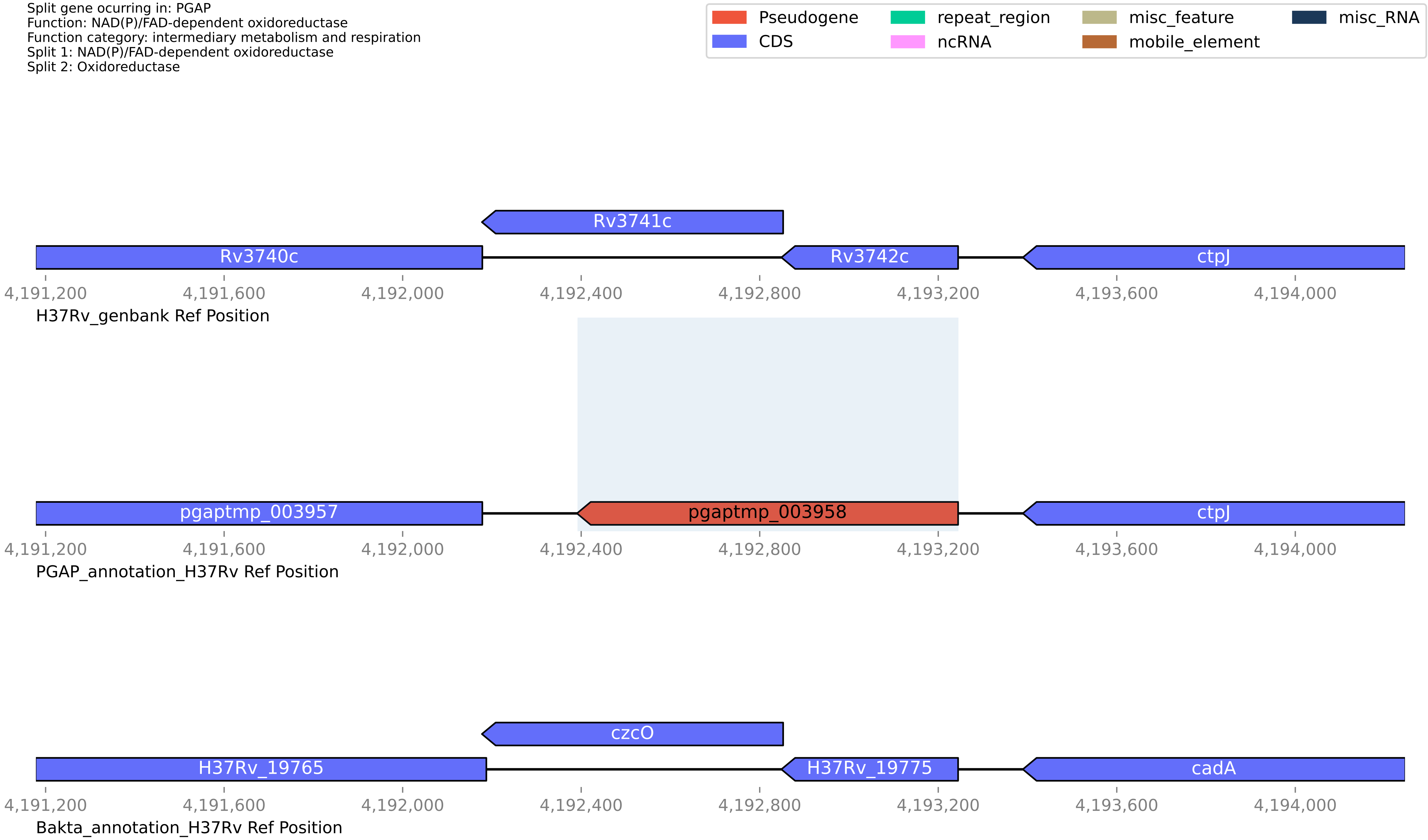

**H37Rv PGAP or Bakta split gene annotation between coordinates 2525402-2526992, compared to Genbank**

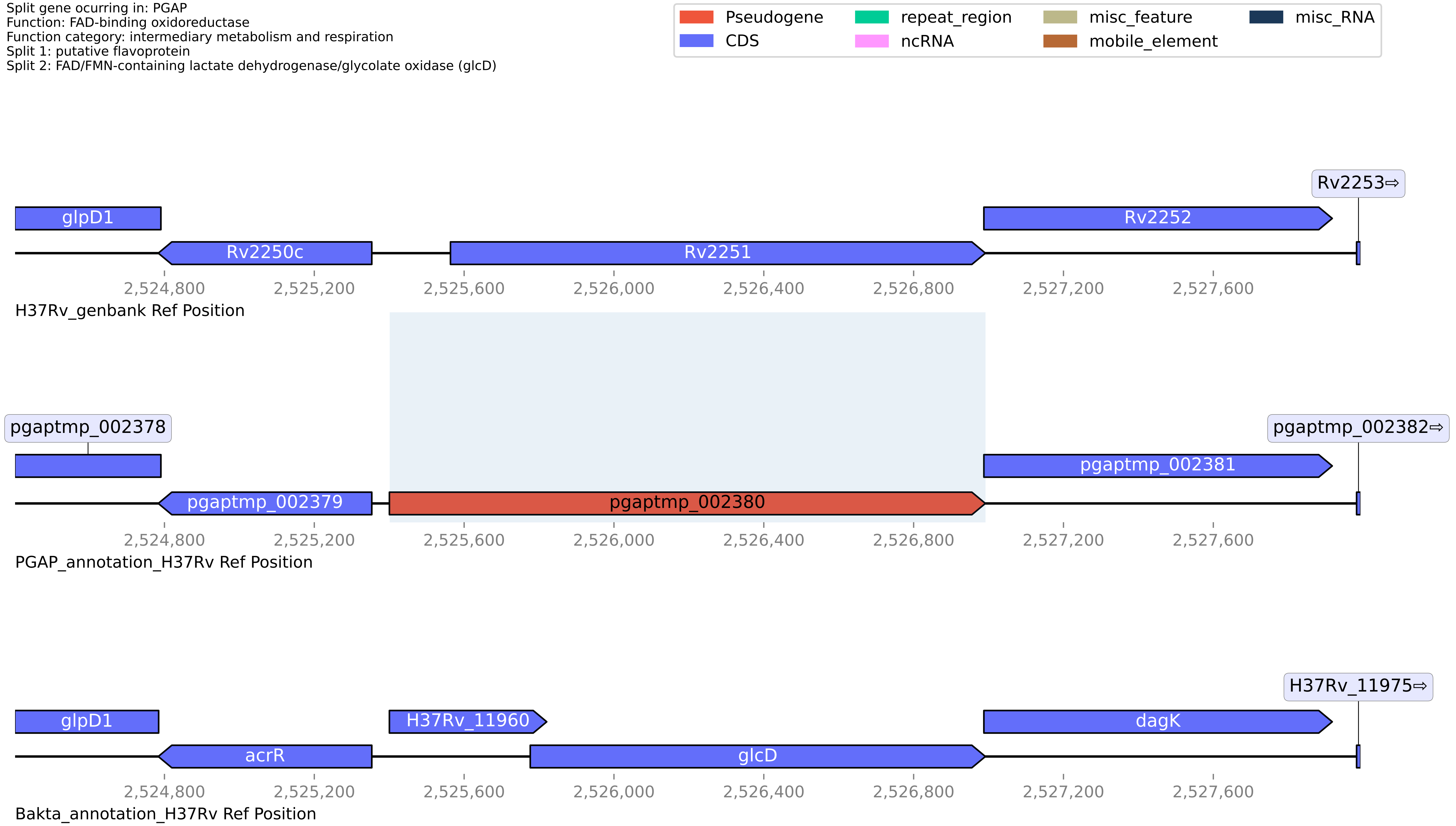

**H37Rv PGAP or Bakta split gene annotation between coordinates 472890-474106, compared to Genbank**

Split gene ocurring in: PGAP
Function: pseudogene
Function category: insertion seqs and phages
Split 1: 13E12 repeat family protein
Split 2: 13E12 repeat family protein

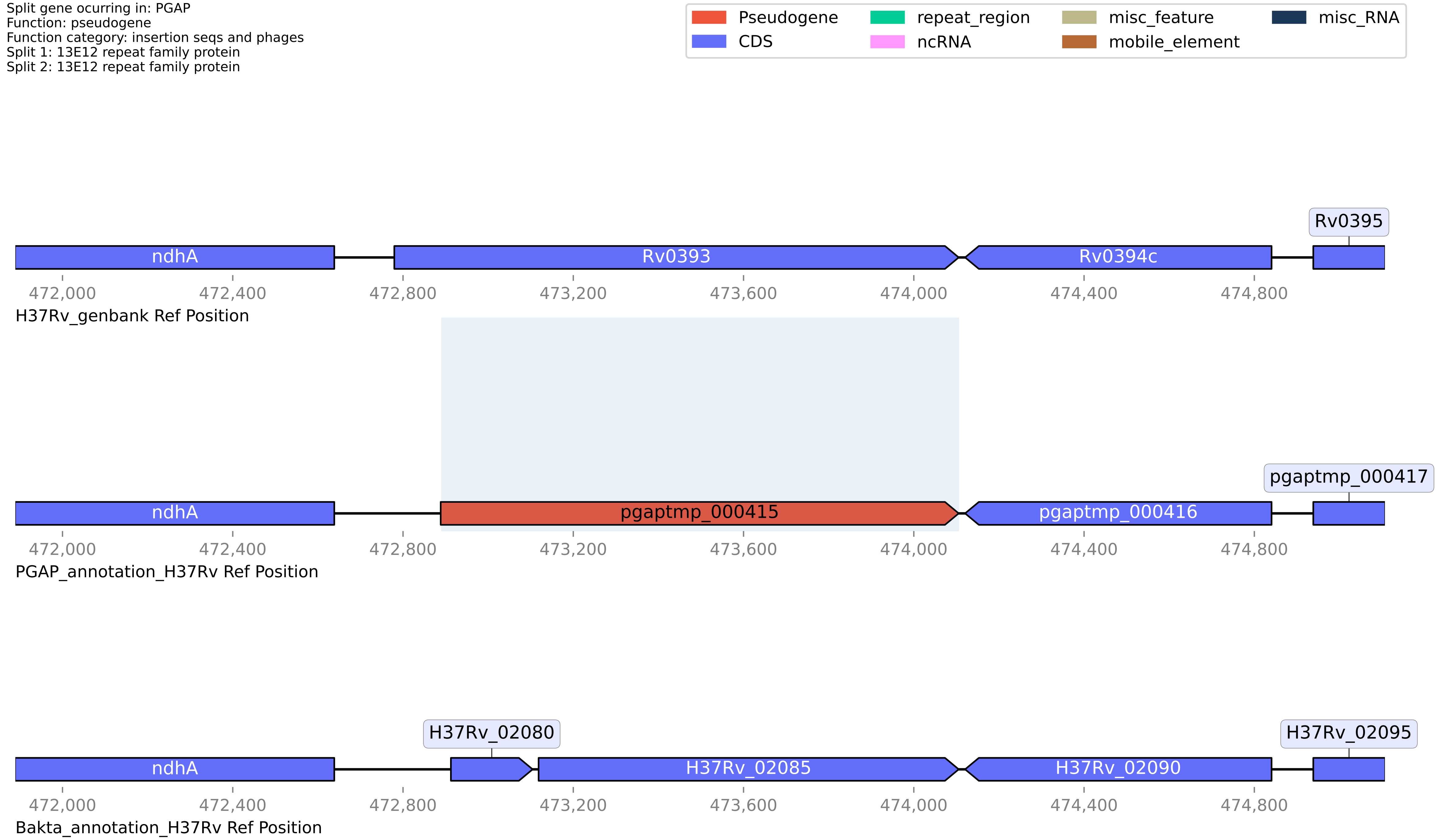

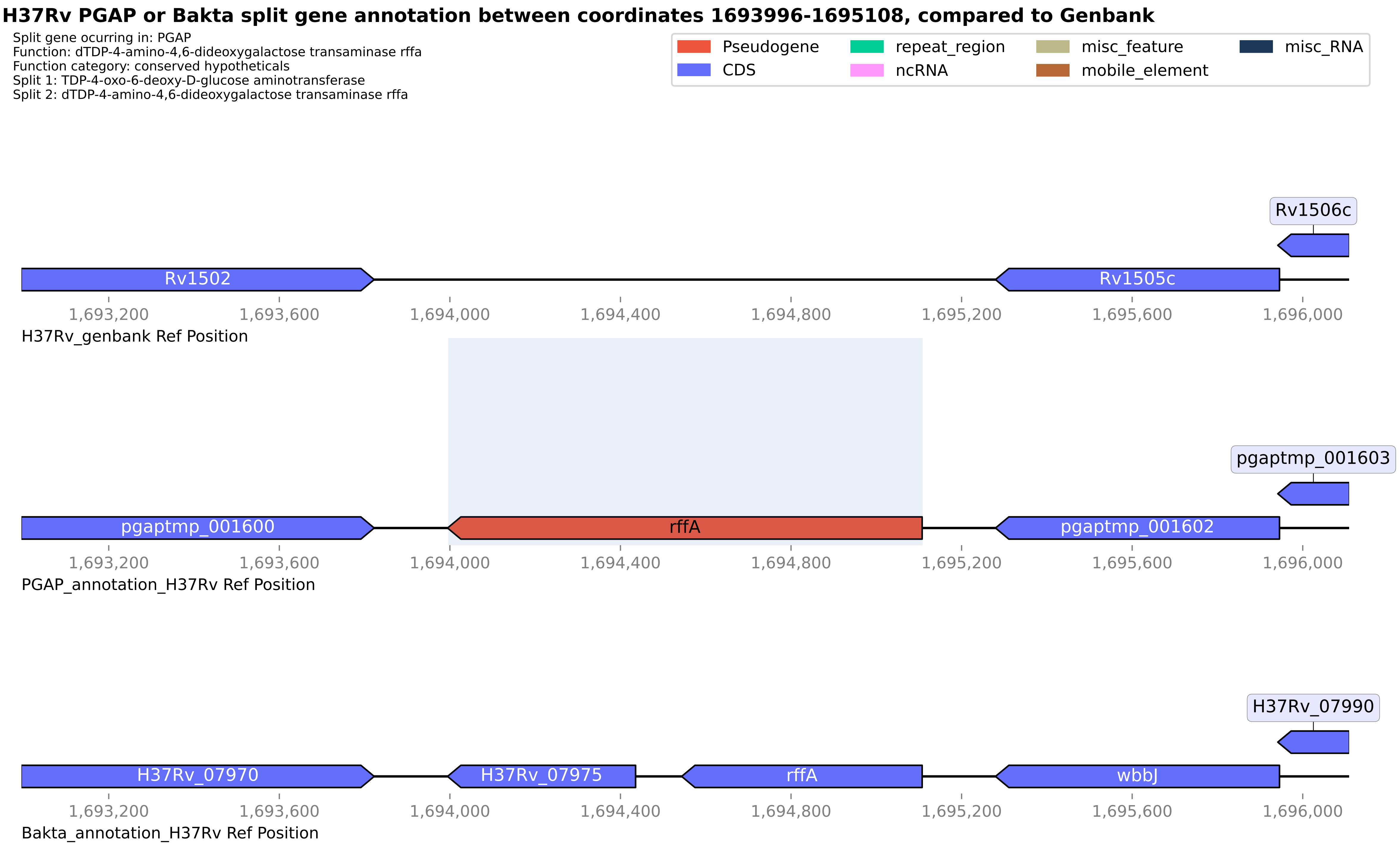

H37Rv PGAP or Bakta split gene annotation between coordinates 1693996-1695108, compared to Genbank
Split gene ocurring in: PGAP
Function: dTDP-4-amino-4,6-dideoxygalactose transaminase rffa
Function category: conserved hypotheticals
Split 1: TDP-4-oxo-6-deoxy-D-glucose aminotransferase
Split 2: dTDP-4-amino-4,6-dideoxygalactose transaminase rffa
Pseudogene
CDS
repeat_region
ncRNA
misc_feature
mobile_element
misc_RNA
Rv1506c
Rv1502
Rv1505c
1,693,200
1,693,600
1,694,000
1,694,400
1,694,800
1,695,200
1,695,600
1,696,000
H37Rv_genbank Ref Position
pgaptmp_001603
pgaptmp_001600
rffA
pgaptmp_001602
PGAP_annotation_H37Rv Ref Position
H37Rv_07990
H37Rv_07970
H37Rv_07975
rffA
wbbJ
Bakta_annotation_H37Rv Ref Position

**H37Rv PGAP or Bakta split gene annotation between coordinates 1893577-1895342, compared to Genbank**

Split gene ocurring in: PGAP
Function: ABC-F family ATP-binding cassette domain-containing protein
Function category: cell wall and cell processes
Split 1: Macrolide-transport ATP-binding protein ABC transporter first part
Split 2: Macrolide ABC transporter ATP-binding protein second part

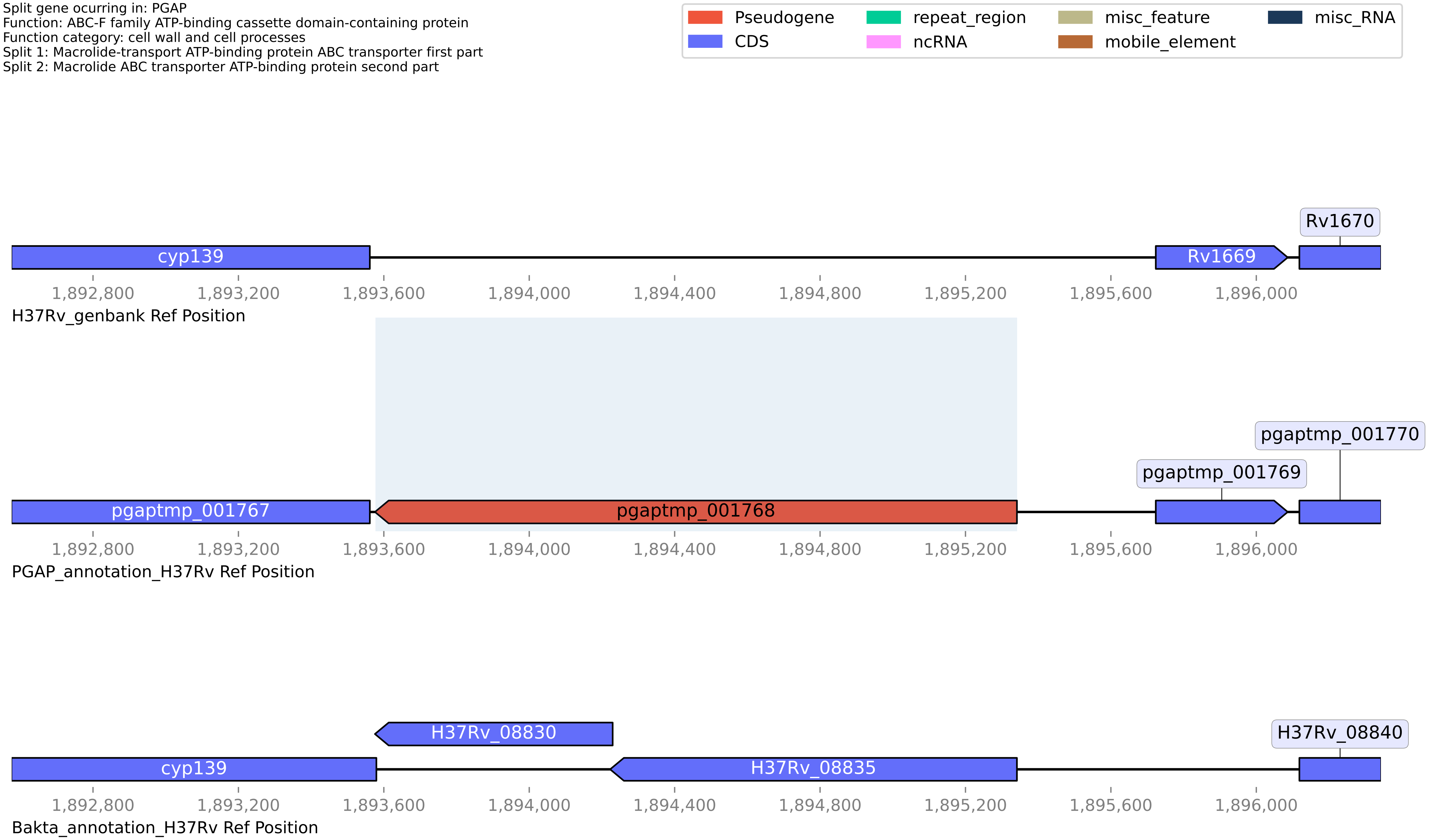

**H37Rv PGAP or Bakta split gene annotation between coordinates 874233-876390, compared to Genbank**

Split gene ocurring in: PGAP
Function: S9 family peptidase
Function category: intermediary metabolism and respiration
Split 1: putative protease II PtrBa [first part] (Oligopeptidase B)
Split 2: putative protease II PtrBb [second part] (Oligopeptidase B)

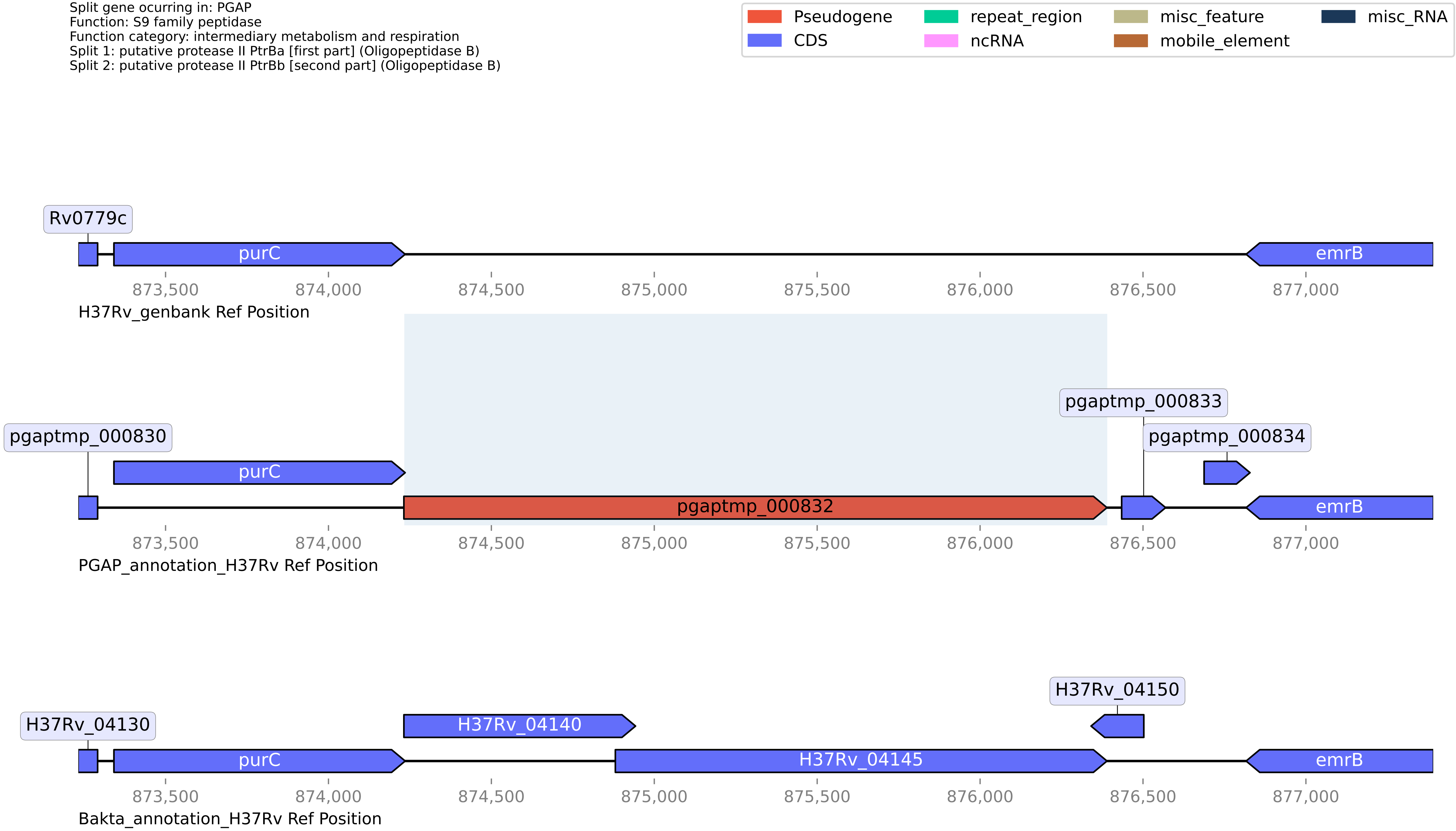

**H37Rv PGAP or Bakta split gene annotation between coordinates 2881409-2882147, compared to Genbank**

Split gene ocurring in: PGAP
Function: DUF2652 domain-containing protein
Function category: conserved hypotheticals
Split 1: DUF2652 domain-containing protein
Split 2: Uncharacterized protein Rv2561/Rv2562

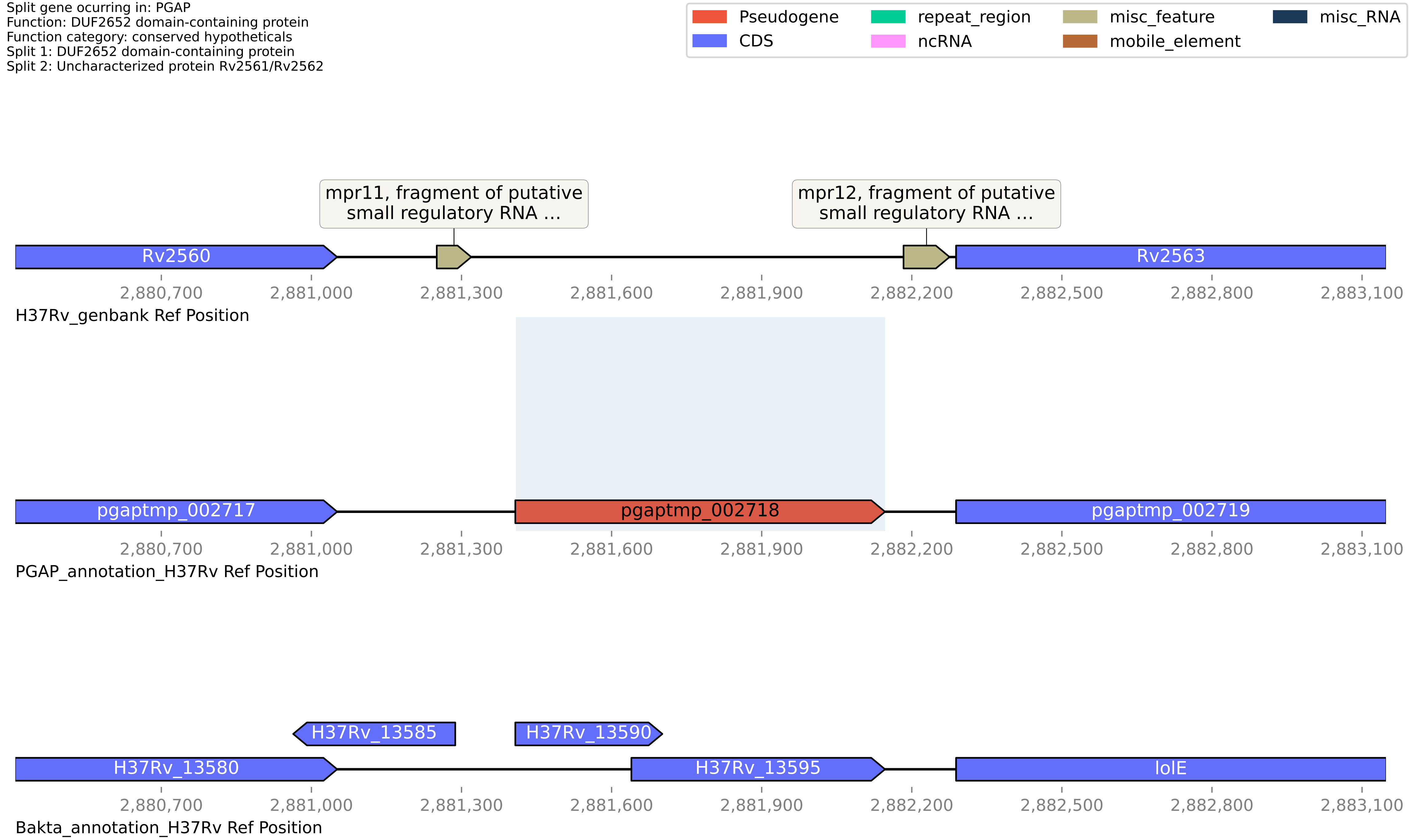

#### H37Rv PGAP or Bakta split gene annotation between coordinates 3435718-3436295, compared to Genbank

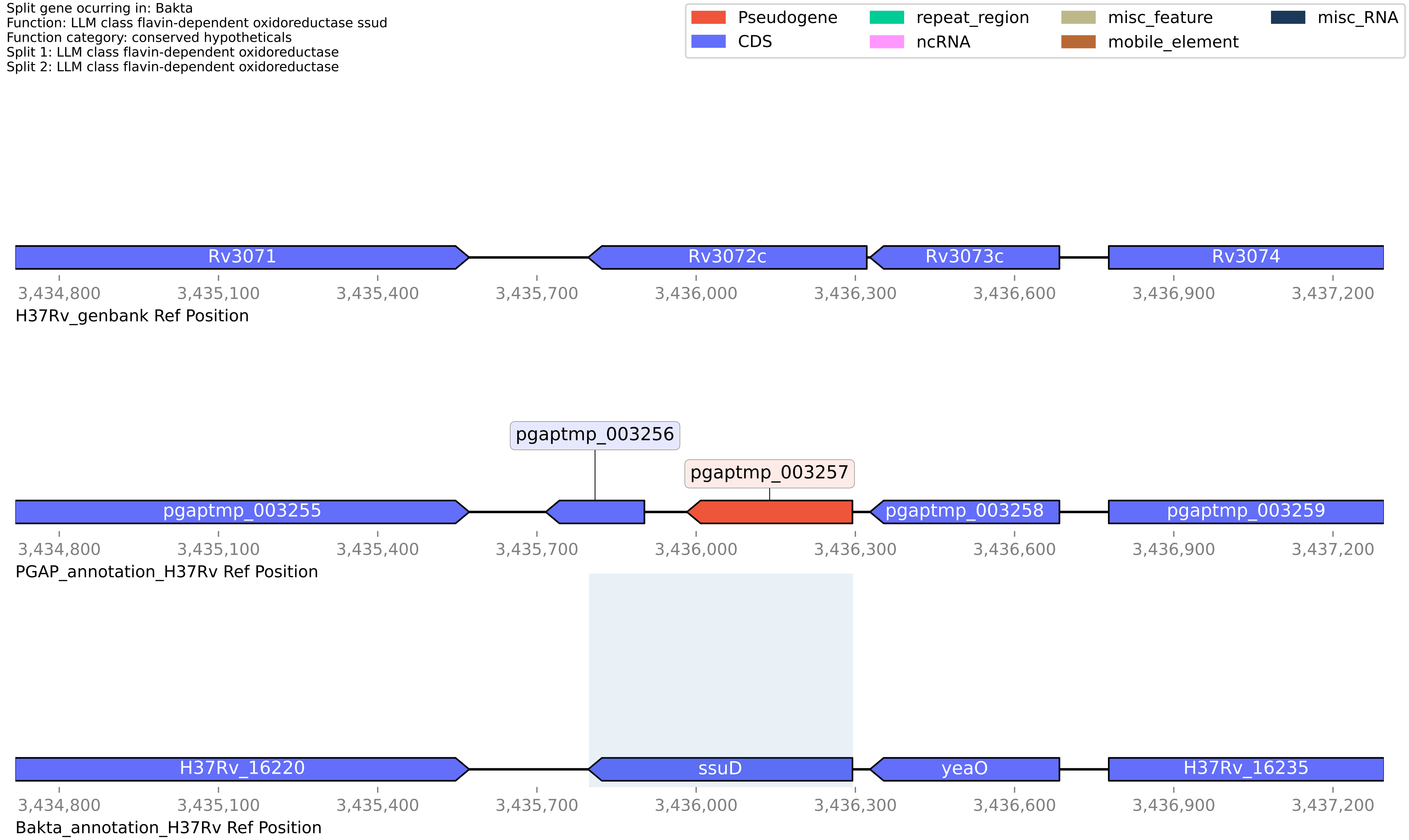

**H37Rv PGAP or Bakta split gene annotation between coordinates 3800017-3801463, compared to Genbank**

Split gene ocurring in: PGAP
Function: ISNCY family transposase
Function category: insertion seqs and phages
Split 1: Transposase and inactivated derivatives, IS5 family
Split 2: Transposase

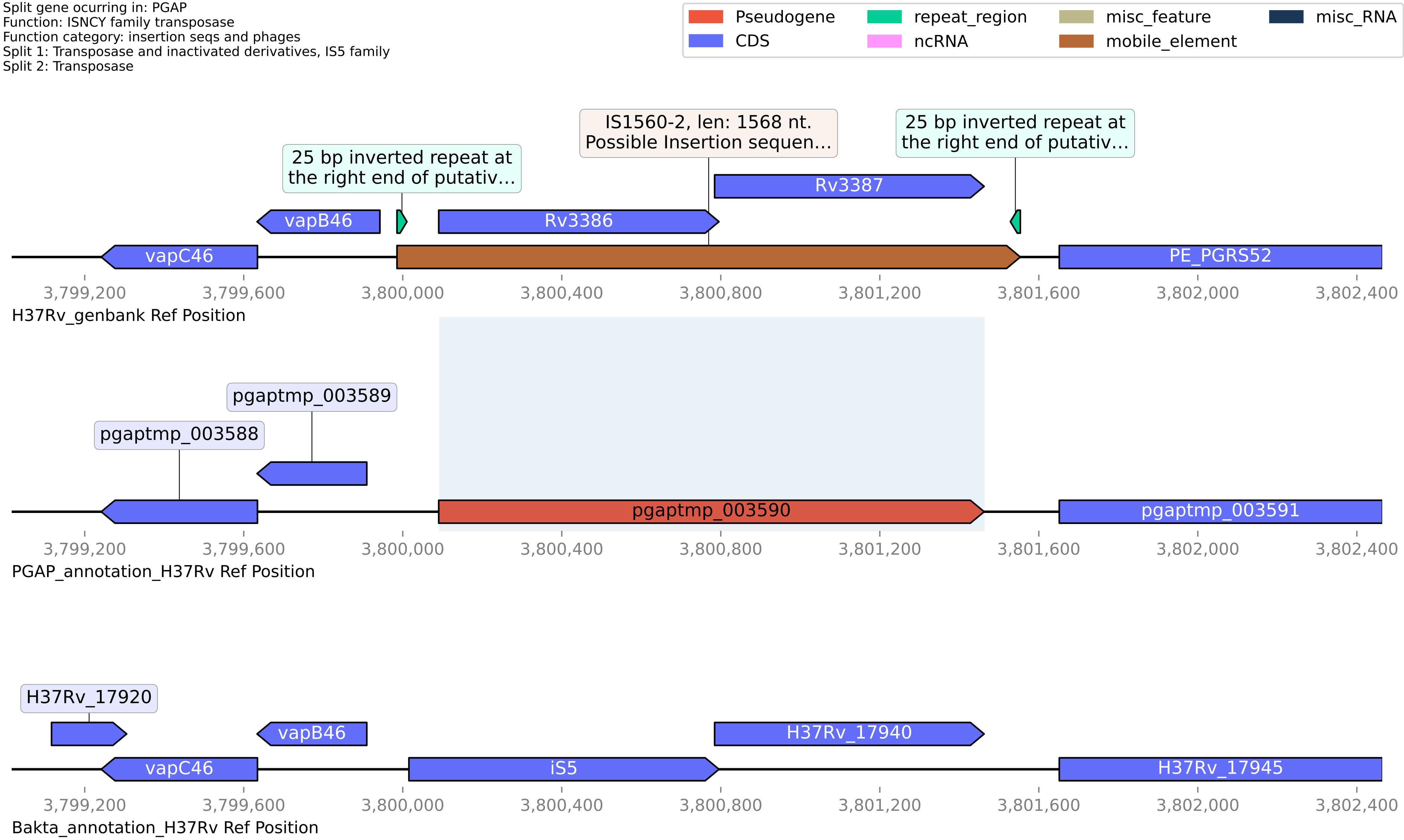

#### H37Rv PGAP or Bakta split gene annotation between coordinates 711536-712719, compared to Genbank

Split gene ocurring in: PGAP
Function: galT
Function category: intermediary metabolism and respiration
Split 1: galactose-1-phosphate uridylyltransferase
Split 2: Galactose-1-phosphate uridylyltransferase

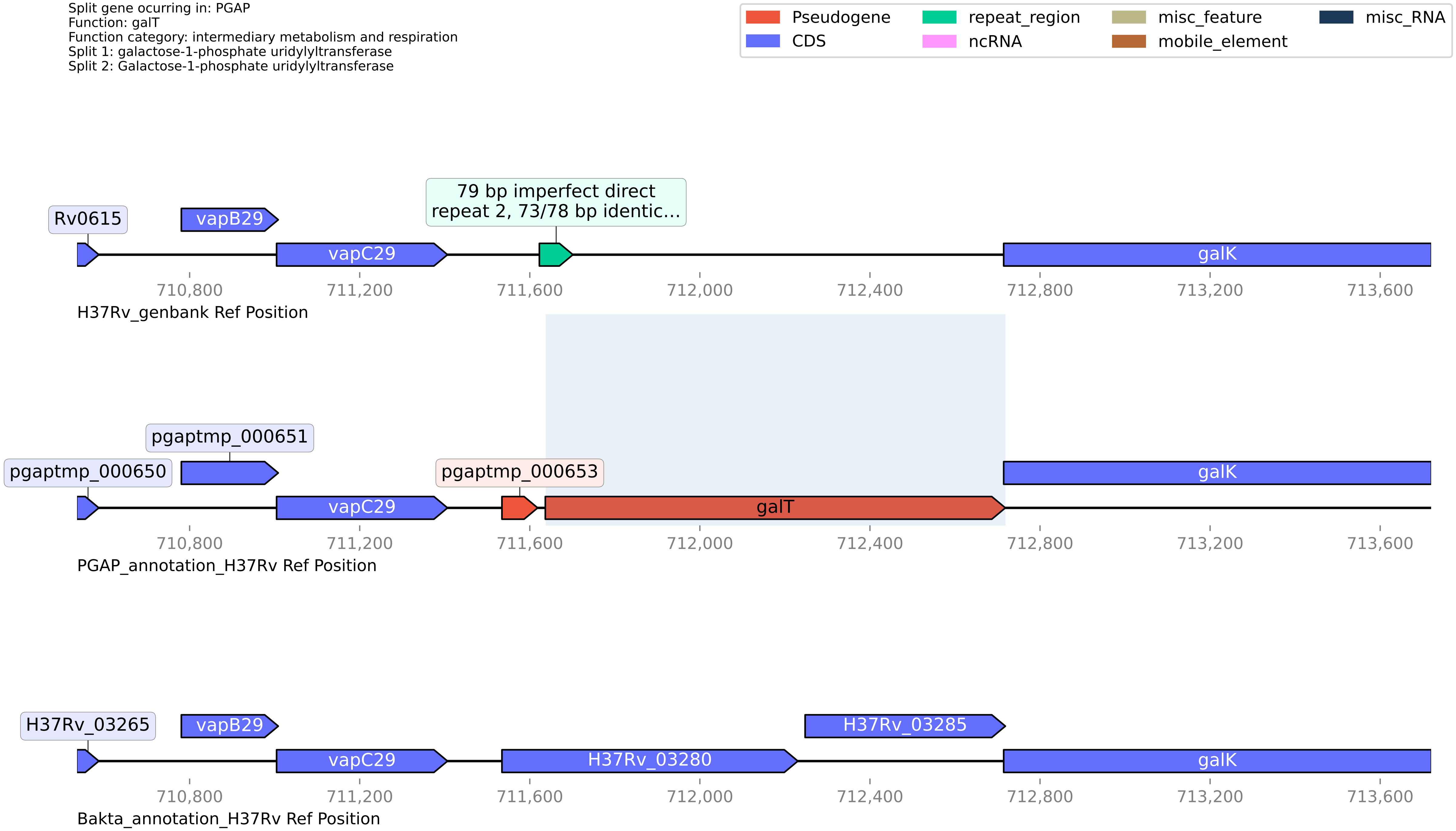

**H37Rv PGAP or Bakta split gene annotation between coordinates 2138174-2139017, compared to Genbank**

Split gene ocurring in: PGAP
Function: class I SAM-dependent methyltransferase
Function category: conserved hypotheticals
Split 1: O-methyltransferase
Split 2: S-adenosyl-L-methionine-dependent methyltransferase (Part1)

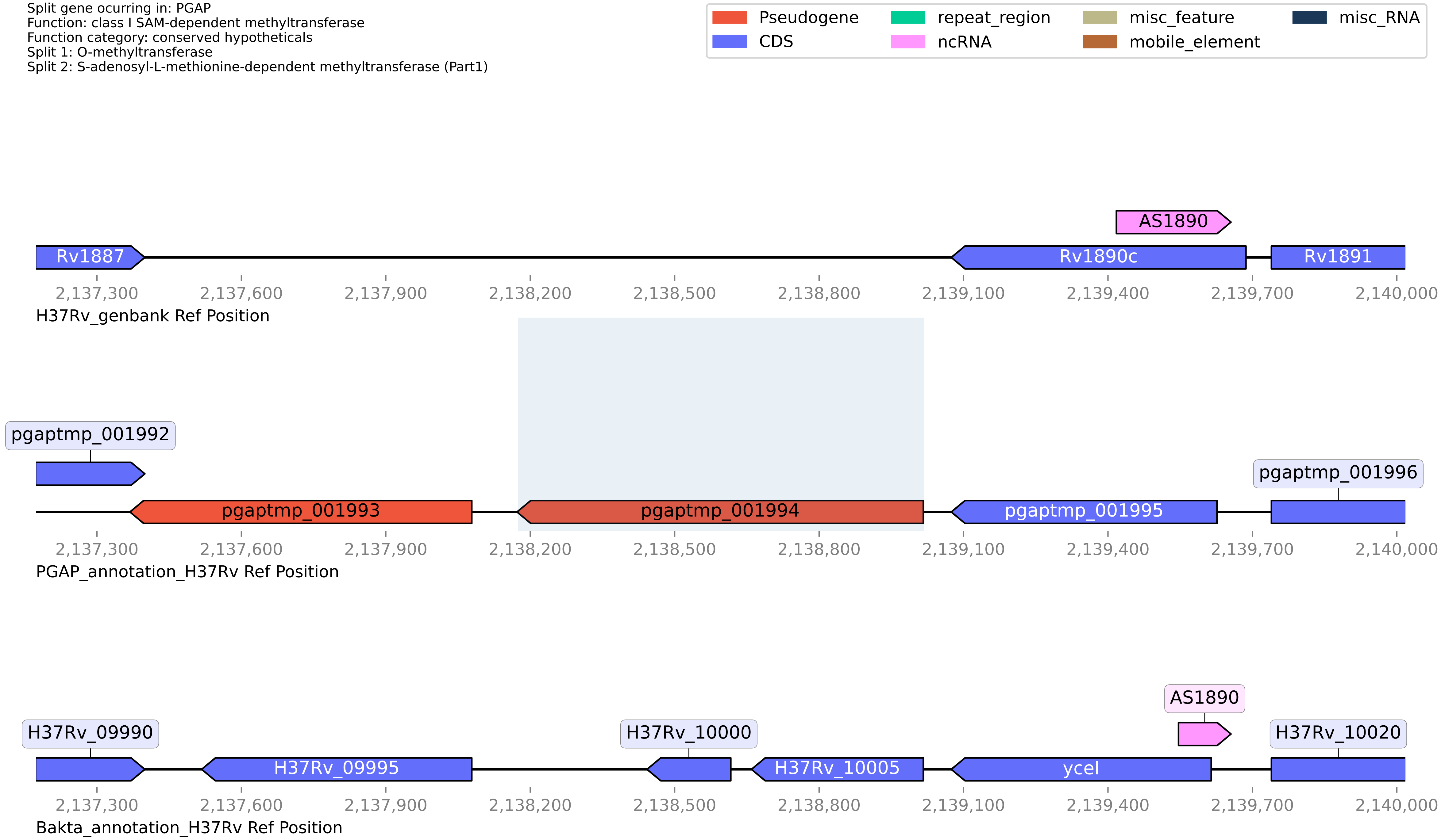

#### H37Rv PGAP or Bakta split gene annotation between coordinates 179319-181029, compared to Genbank

Split gene ocurring in: PGAP
Function: PE-PPE domain-containing protein
Function category: PE/PPE
Split 1: PE family protein
Split 2: PE-PGRS family protein

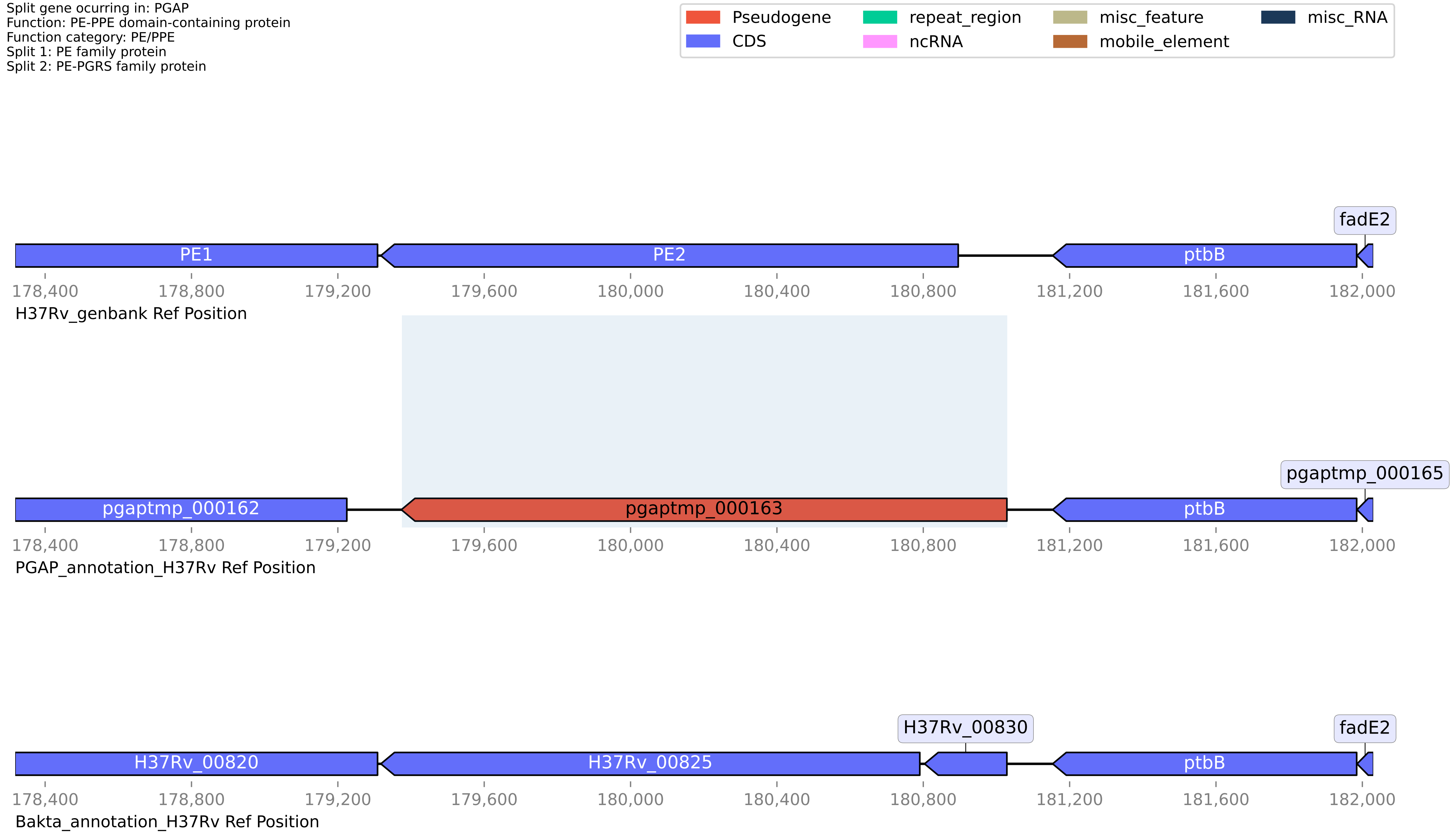

#### H37Rv PGAP or Bakta split gene annotation between coordinates 2500923-2501632, compared to Genbank

Split gene ocurring in: PGAP
Function: 2OG-Fe(II) oxygenase
Function category: conserved hypotheticals
Split 1: proline hydroxylase
Split 2: DUF2086 domain-containing protein

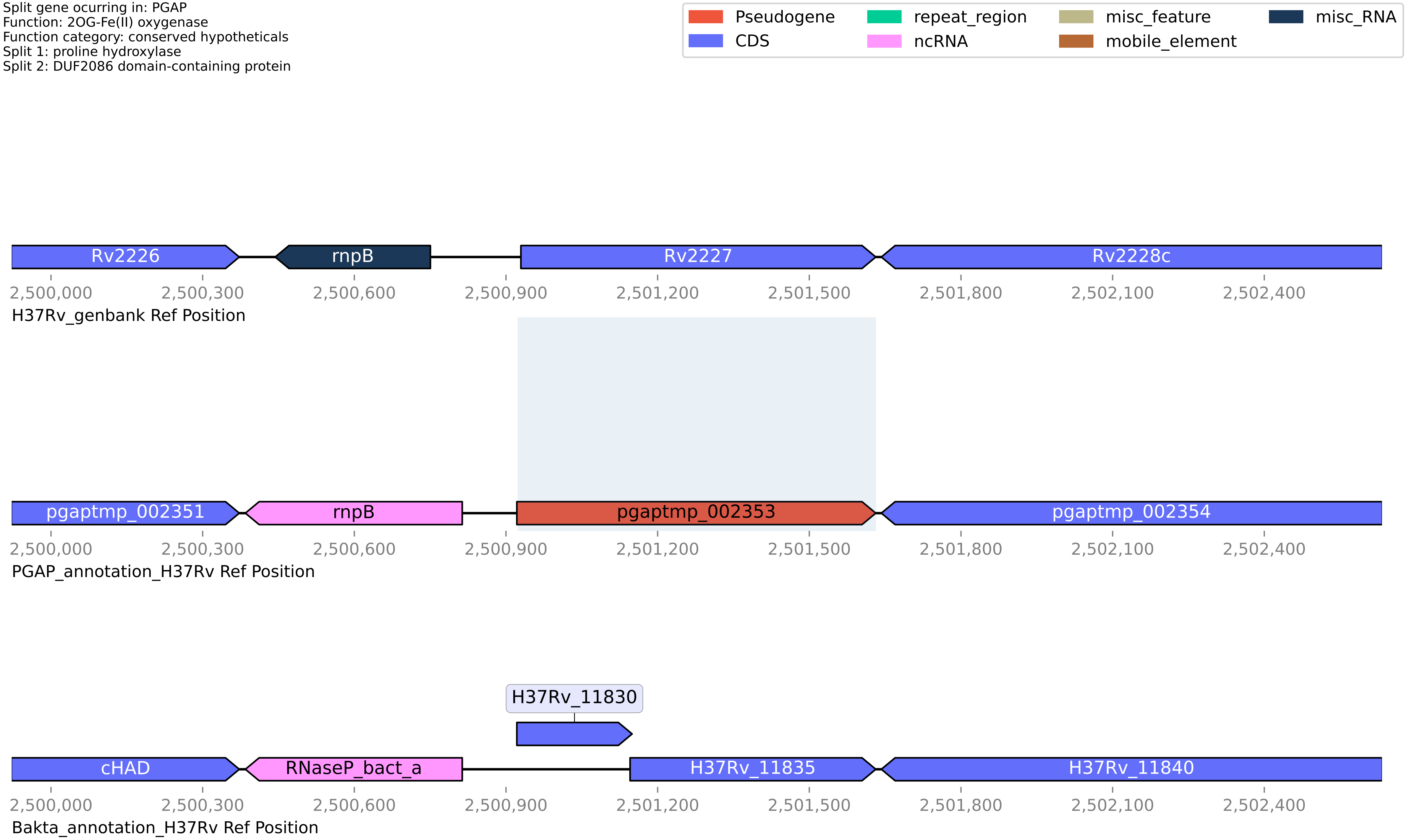

#### H37Rv PGAP or Bakta split gene annotation between coordinates 4192179-4193245, compared to Genbank

Split gene ocurring in: PGAP
Function: NAD(P)/FAD-dependent oxidoreductase
Function category: intermediary metabolism and respiration
Split 1: NAD(P)/FAD-dependent oxidoreductase
Split 2: Oxidoreductase

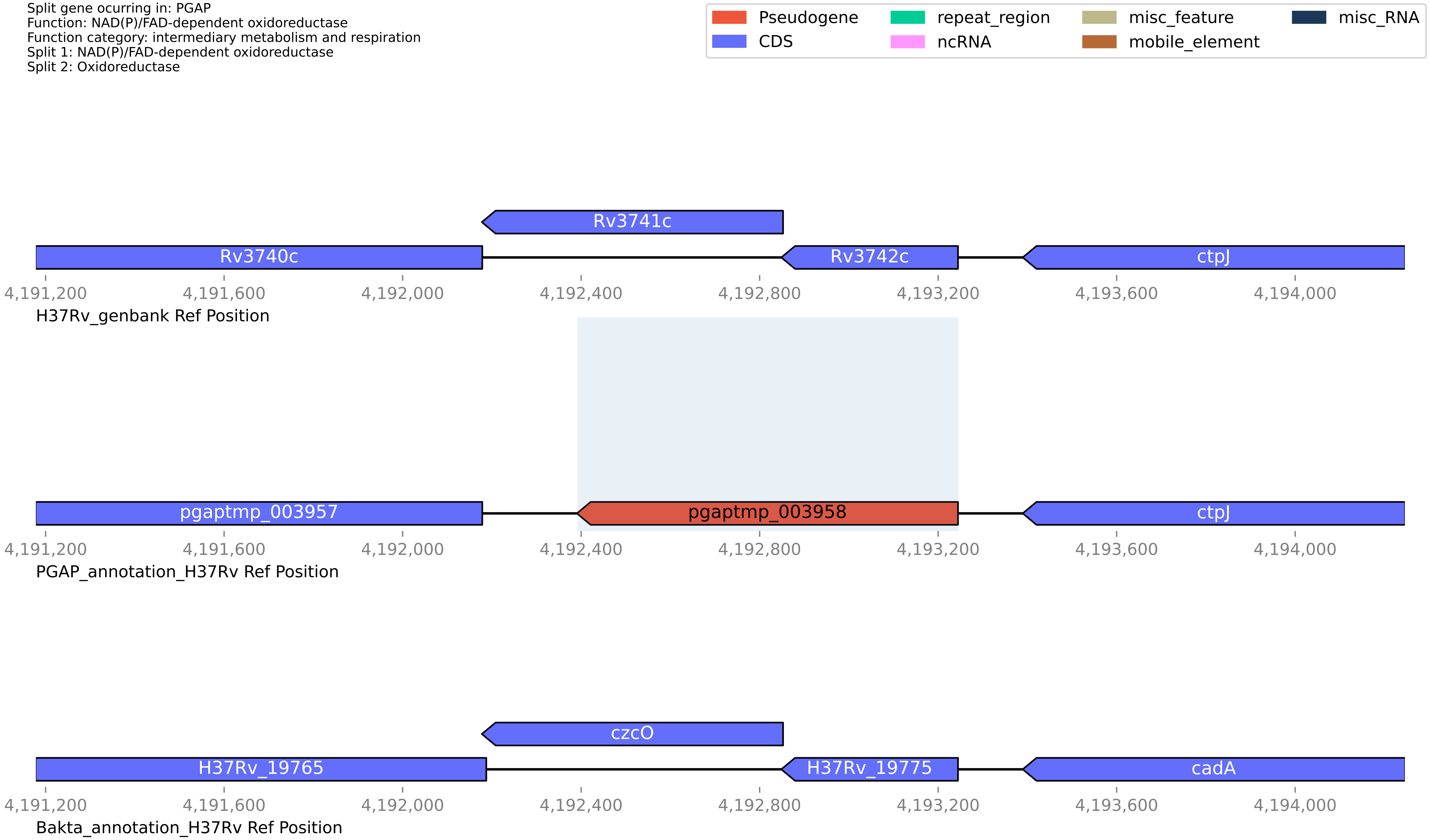

**H37Rv PGAP or Bakta split gene annotation between coordinates 2525402-2526992, compared to Genbank**

Split gene ocurring in: PGAP
Function: FAD-binding oxidoreductase
Function category: intermediary metabolism and respiration
Split 1: putative flavoprotein
Split 2: FAD/FMN-containing lactate dehydrogenase/glycolate oxidase (glcD)

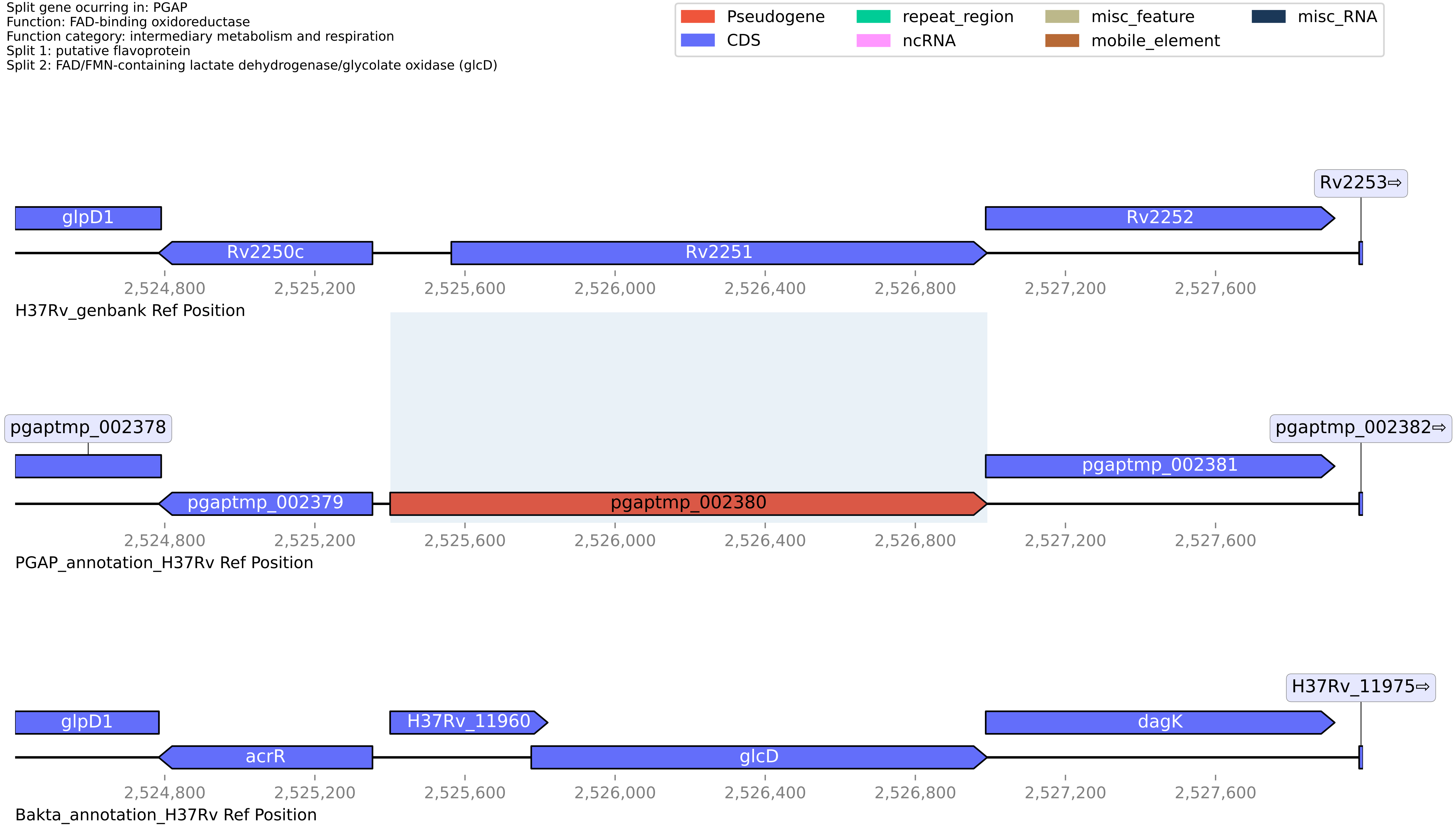

**H37Rv PGAP or Bakta split gene annotation between coordinates 472890-474106, compared to Genbank**

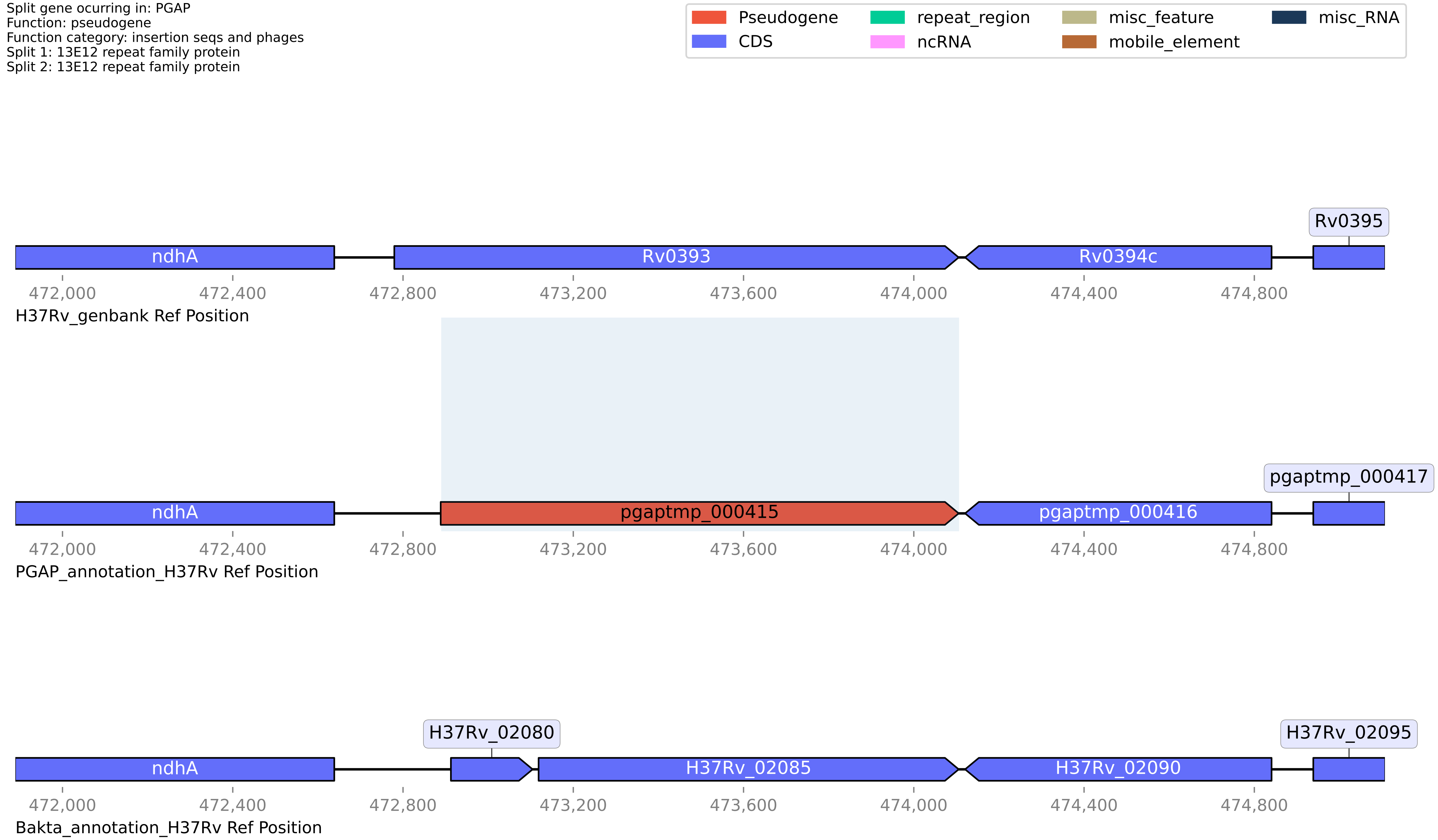

**H37Rv PGAP or Bakta split gene annotation between coordinates 1693996-1695108, compared to Genbank**

Split gene ocurring in: PGAP
Function: dTDP-4-amino-4,6-dideoxygalactose transaminase rffa
Function category: conserved hypotheticals
Split 1: TDP-4-oxo-6-deoxy-D-glucose aminotransferase
Split 2: dTDP-4-amino-4,6-dideoxygalactose transaminase rffa

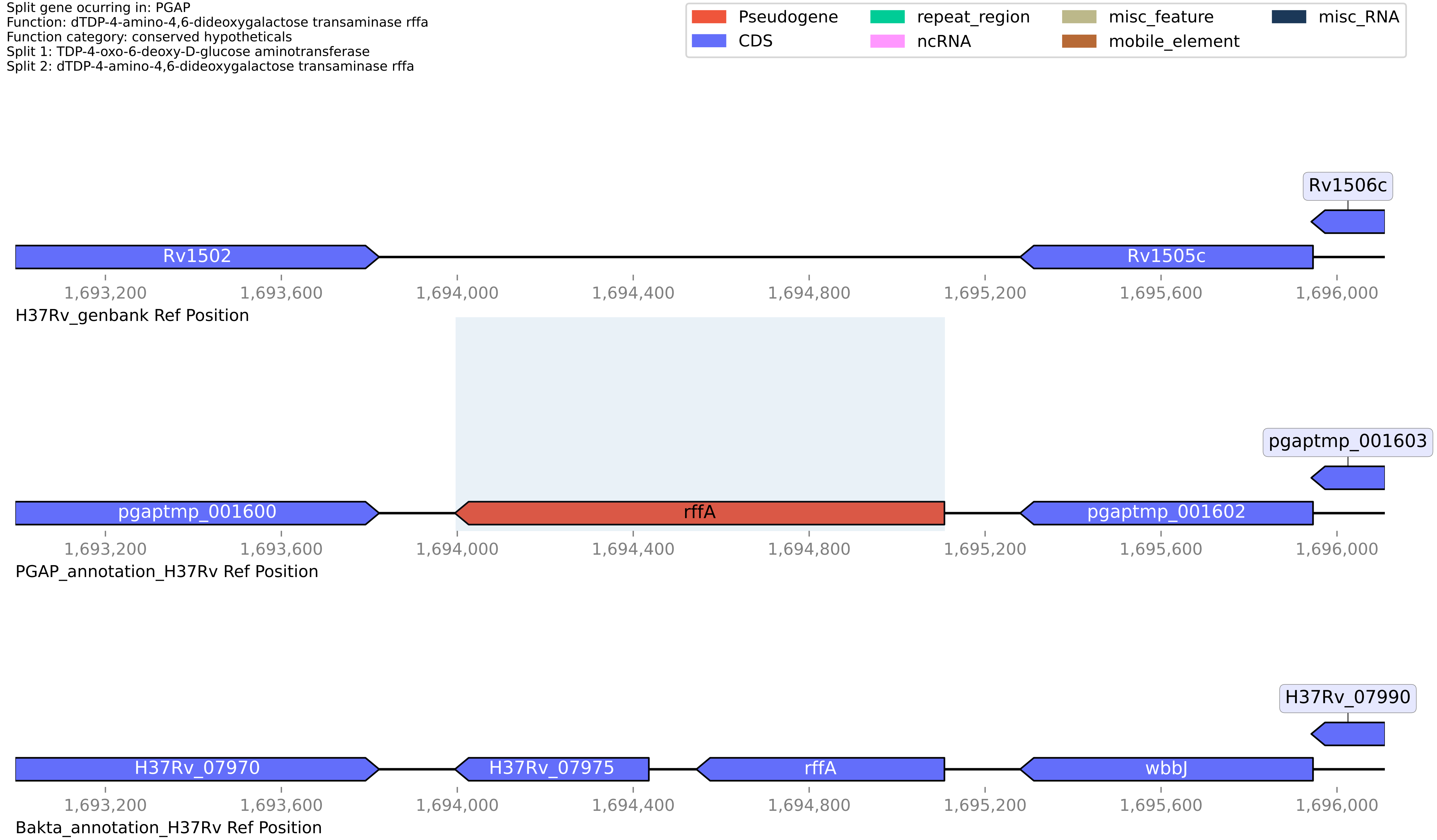

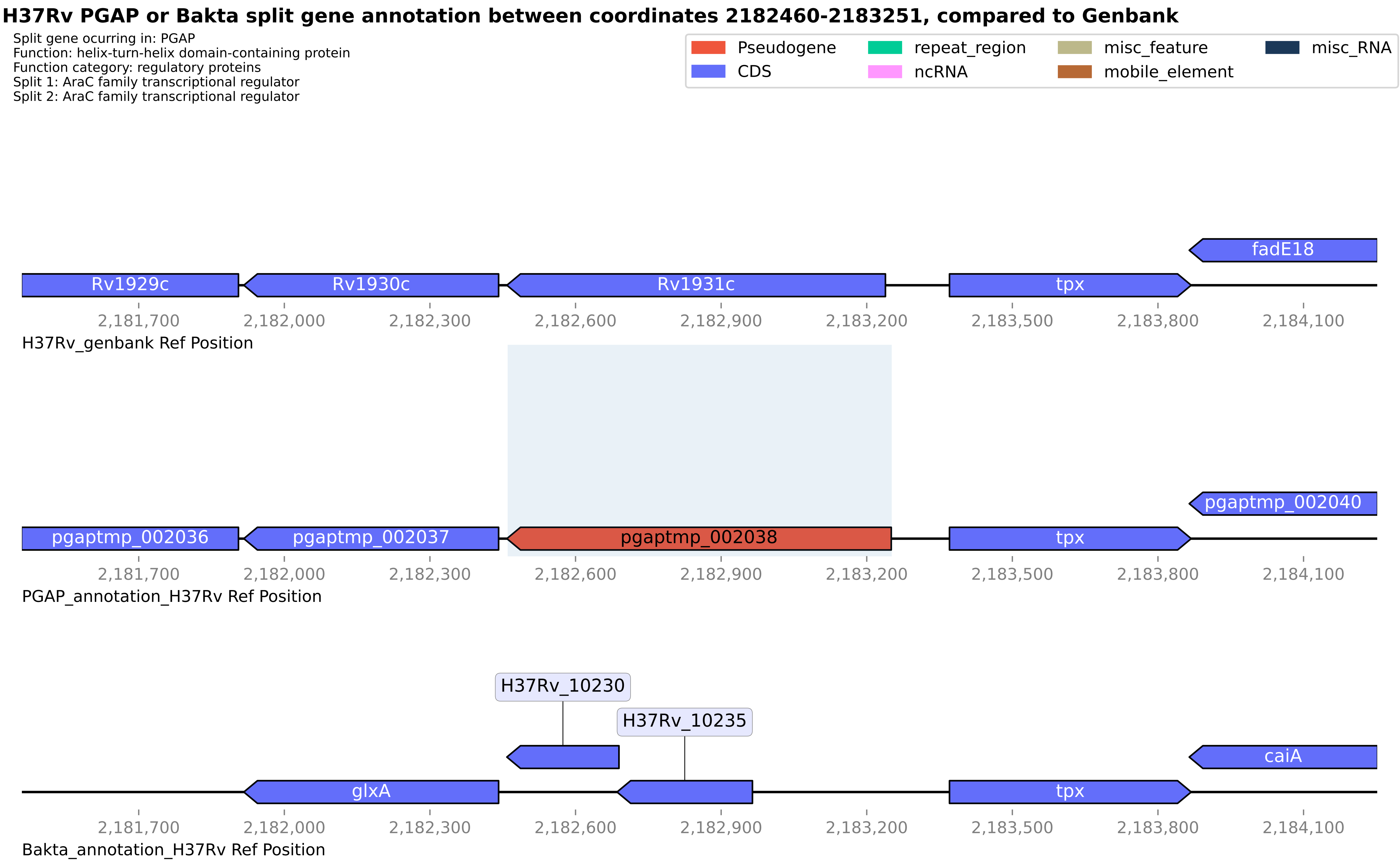

H37Rv PGAP or Bakta split gene annotation between coordinates 2182460-2183251, compared to Genbank
Split gene ocurring in: PGAP
Function: helix-turn-helix domain-containing protein
Function category: regulatory proteins
Split 1: AraC family transcriptional regulator
Split 2: AraC family transcriptional regulator
Pseudogene
CDS
repeat_region
ncRNA
misc_feature
mobile_element
misc_RNA
fadE18
Rv1929c
Rv1930c
Rv1931c
tpx
2,181,700
2,182,000
2,182,300
2,182,600
2,182,900
2,183,200
2,183,500
2,183,800
2,184,100
H37Rv_genbank Ref Position
pgaptmp_002040
pgaptmp_002036
pgaptmp_002037
pgaptmp_002038
tpx
PGAP_annotation_H37Rv Ref Position
H37Rv_10230
H37Rv_10235
caiA
glxA
tpx
Bakta_annotation_H37Rv Ref Position

**H37Rv PGAP or Bakta split gene annotation between coordinates 1173945-1174700, compared to Genbank**

Split gene ocurring in: Bakta
Function: HTH-17 domain-containing protein
Function category: conserved hypotheticals
Split 1: helix-turn-helix domain-containing protein
Split 2: nucleotidyl transferase AbiEii/AbiGii toxin family protein

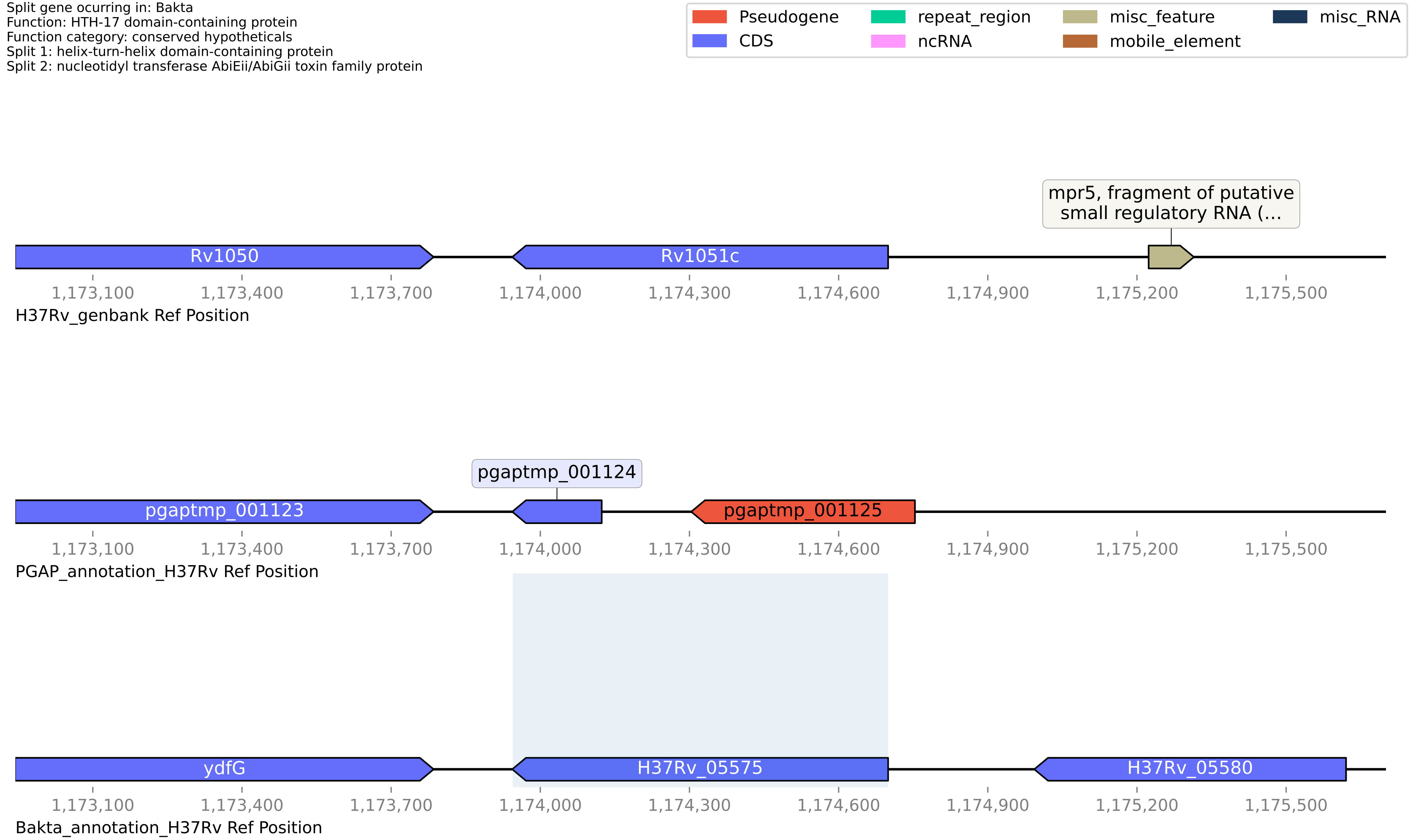

#### H37Rv PGAP or Bakta split gene annotation between coordinates 3609781-3611189, compared to Genbank

Split gene ocurring in: PGAP
Function: wax ester/triacylglycerol synthase family O-acyltransferase
Function category: lipid metabolism
Split 1: Diacylglycerol O-acyltransferase
Split 2: putative diacyglycerol O-acyltransferase tgs3

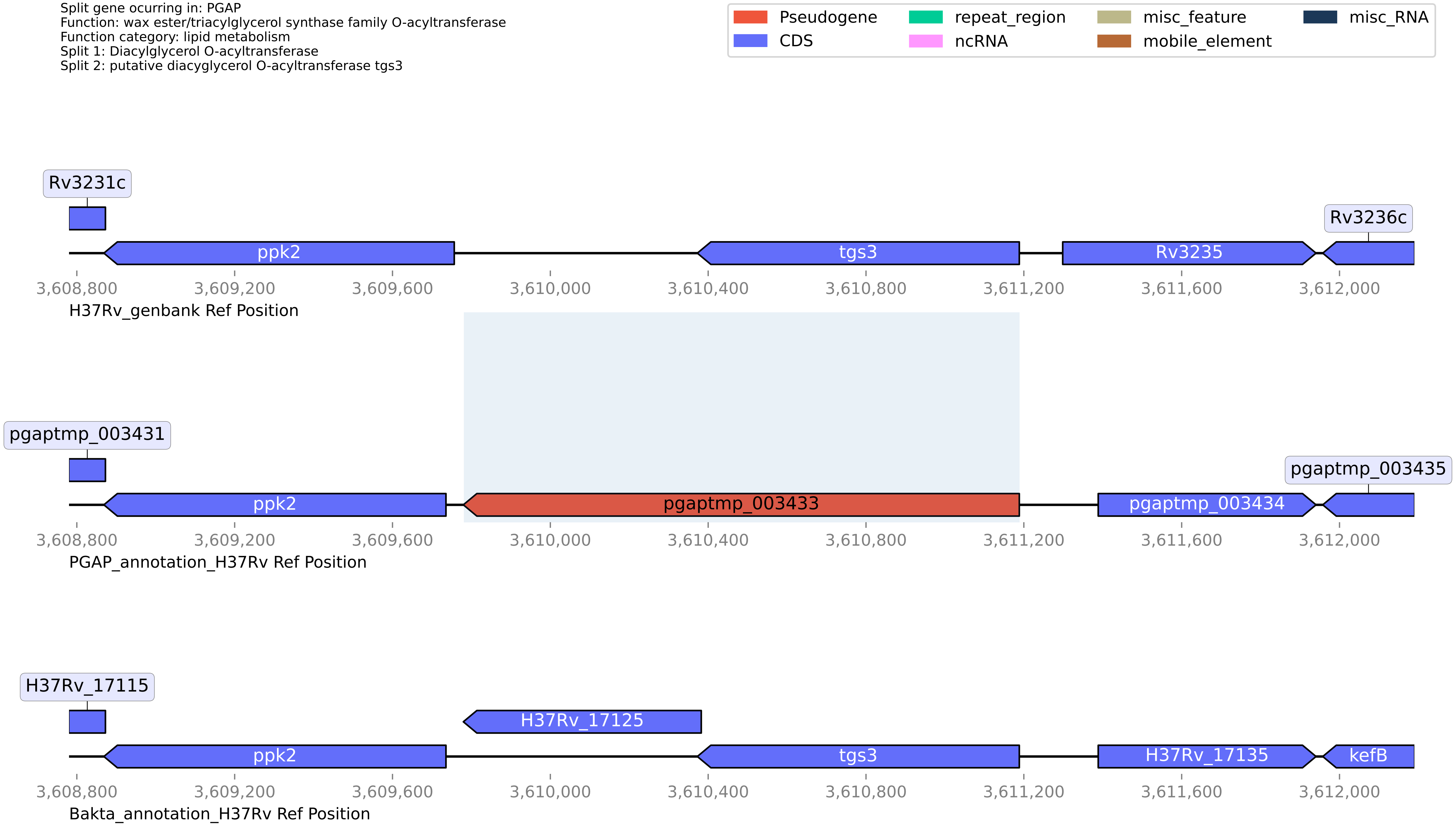

**H37Rv PGAP or Bakta split gene annotation between coordinates 3874404-3876090, compared to Genbank**

Split gene ocurring in: PGAP
Function: hypothetical protein
Function category: cell wall and cell processes
Split 1: Transmembrane protein
Split 2: Transmembrane protein

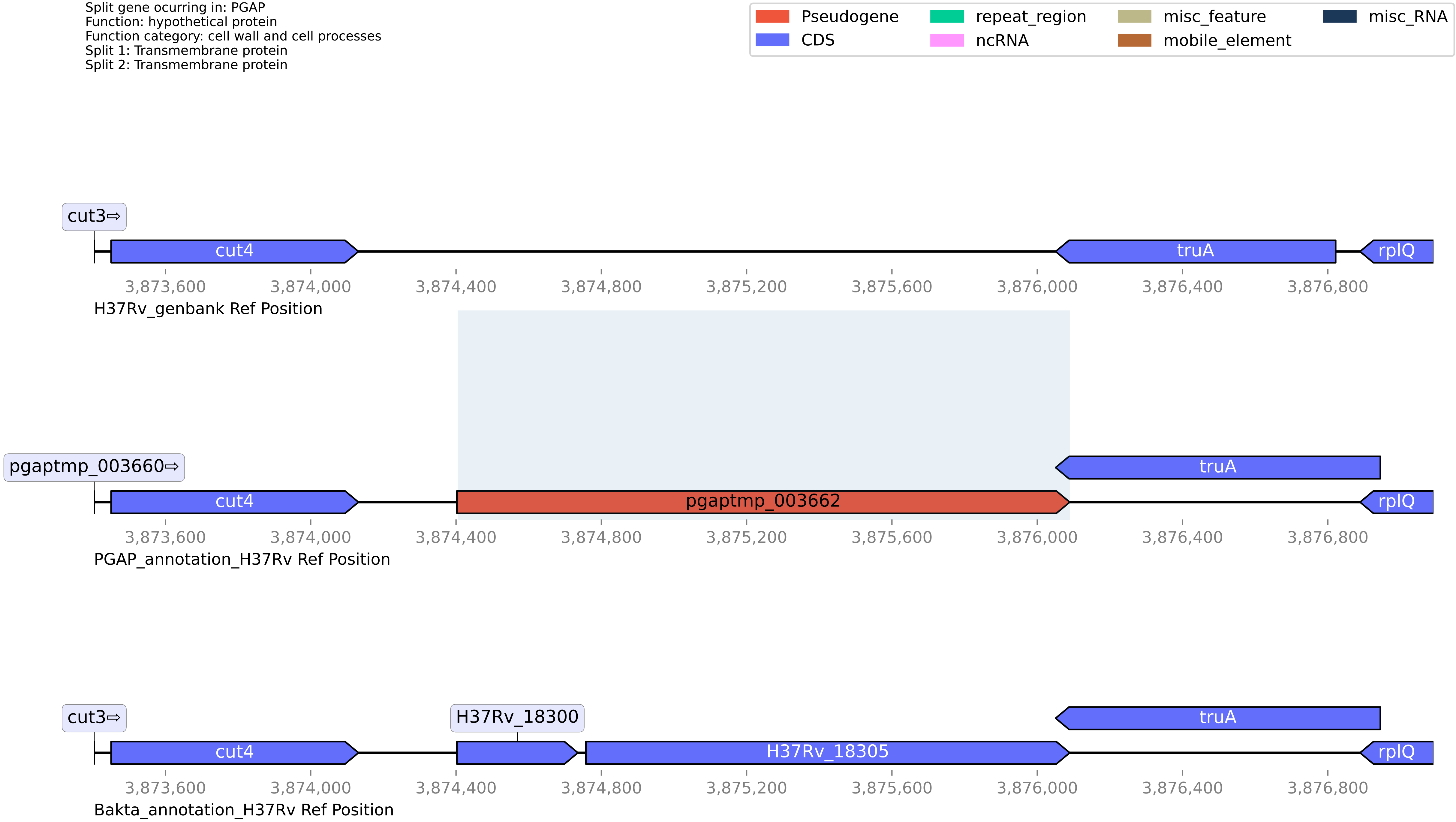

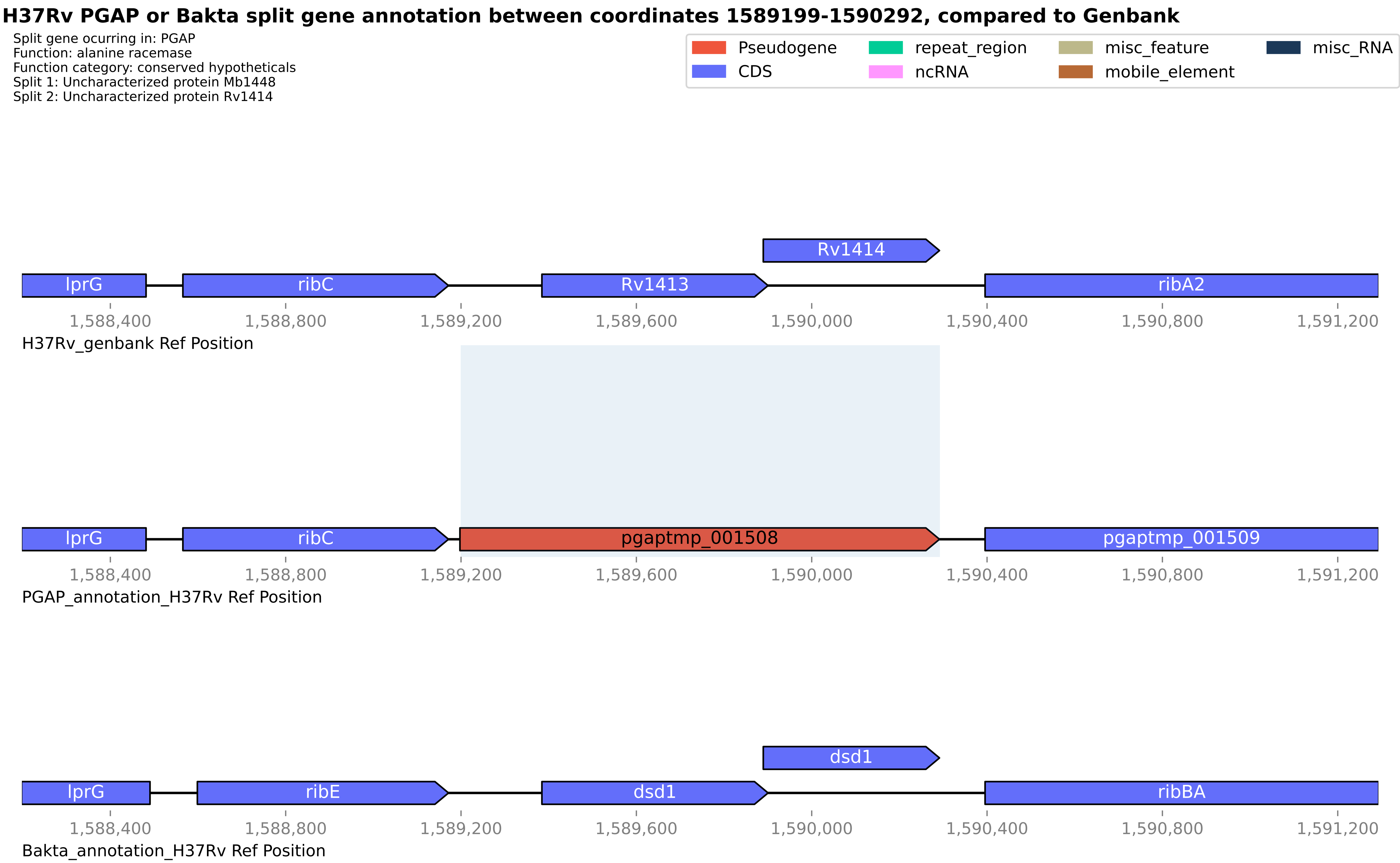
H37Rv PGAP or Bakta split gene annotation between coordinates 1589199-1590292, compared to Genbank
Split gene ocurring in: PGAP
Function: alanine racemase
Function category: conserved hypotheticals
Split 1: Uncharacterized protein Mb1448
Split 2: Uncharacterized protein Rv1414
Pseudogene
CDS
repeat_region
ncRNA
misc_feature
mobile_element
misc_RNA
Rv1414
lprG
ribC
Rv1413
ribA2
1,588,400
1,588,800
1,589,200
1,589,600
1,590,000
1,590,400
1,590,800
1,591,200
H37Rv_genbank Ref Position
lprG
ribC
pgaptmp_001508
pgaptmp_001509
PGAP_annotation_H37Rv Ref Position
dsd1
lprG
ribE
dsd1
ribBA
Bakta_annotation_H37Rv Ref Position

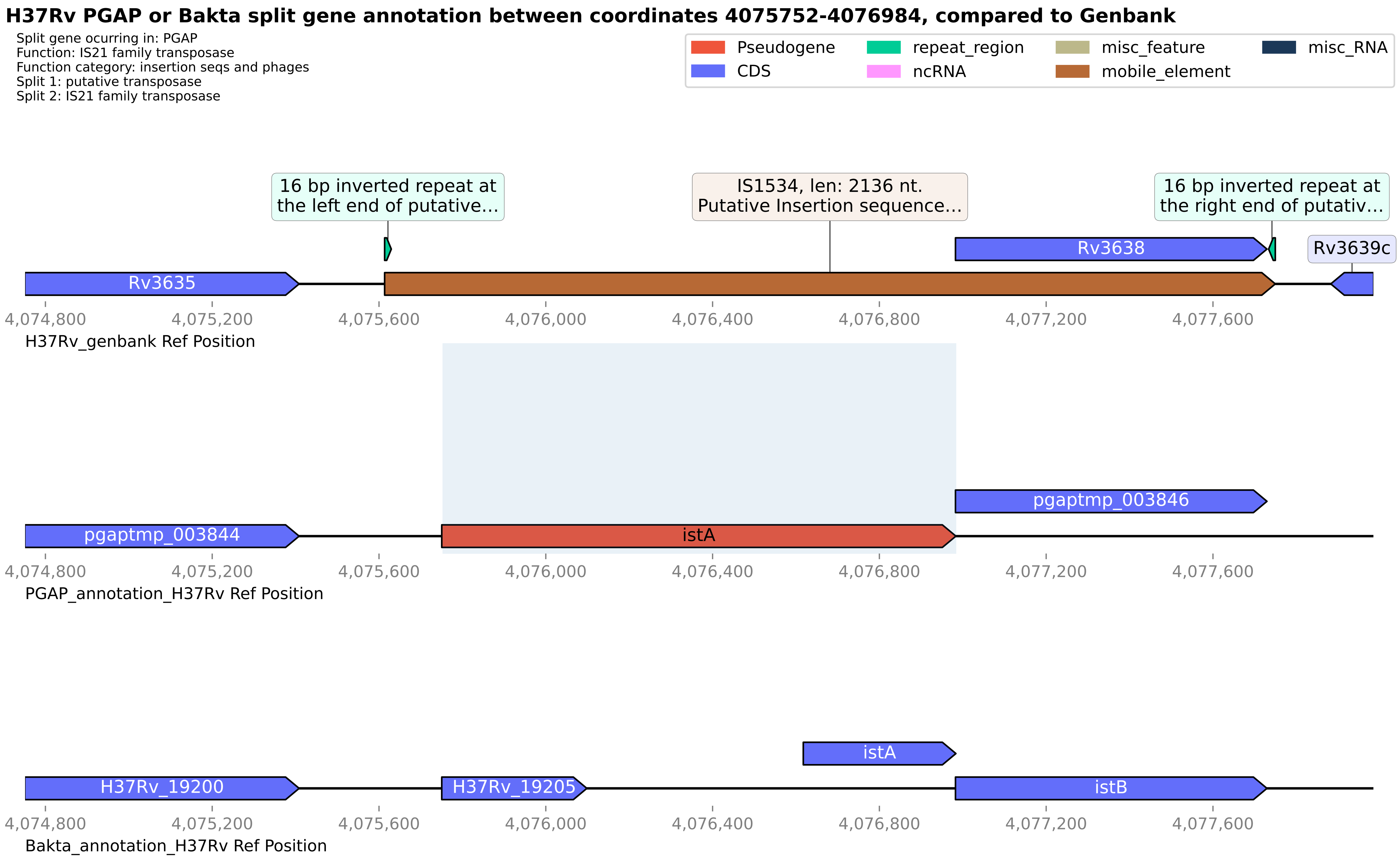
H37Rv PGAP or Bakta split gene annotation between coordinates 4075752-4076984, compared to Genbank
Split gene ocurring in: PGAP
Function: IS21 family transposase
Function category: insertion seqs and phages
Split 1: putative transposase
Split 2: IS21 family transposase
Pseudogene
CDS
repeat_region
ncRNA
misc_feature
mobile_element
misc_RNA
16 bp inverted repeat at the left end of putative...
IS1534, len: 2136 nt. Putative Insertion sequence...
16 bp inverted repeat at the right end of putativ...
Rv3638
Rv3639c
Rv3635
4,074,800
4,075,200
4,075,600
4,076,000
4,076,400
4,076,800
4,077,200
4,077,600
H37Rv_genbank Ref Position
pgaptmp_003846
pgaptmp_003844
istA
PGAP_annotation_H37Rv Ref Position
istA
H37Rv_19200
H37Rv_19205
istB
Bakta_annotation_H37Rv Ref Position

**H37Rv PGAP or Bakta split gene annotation between coordinates 2030347-2030643, compared to Genbank**

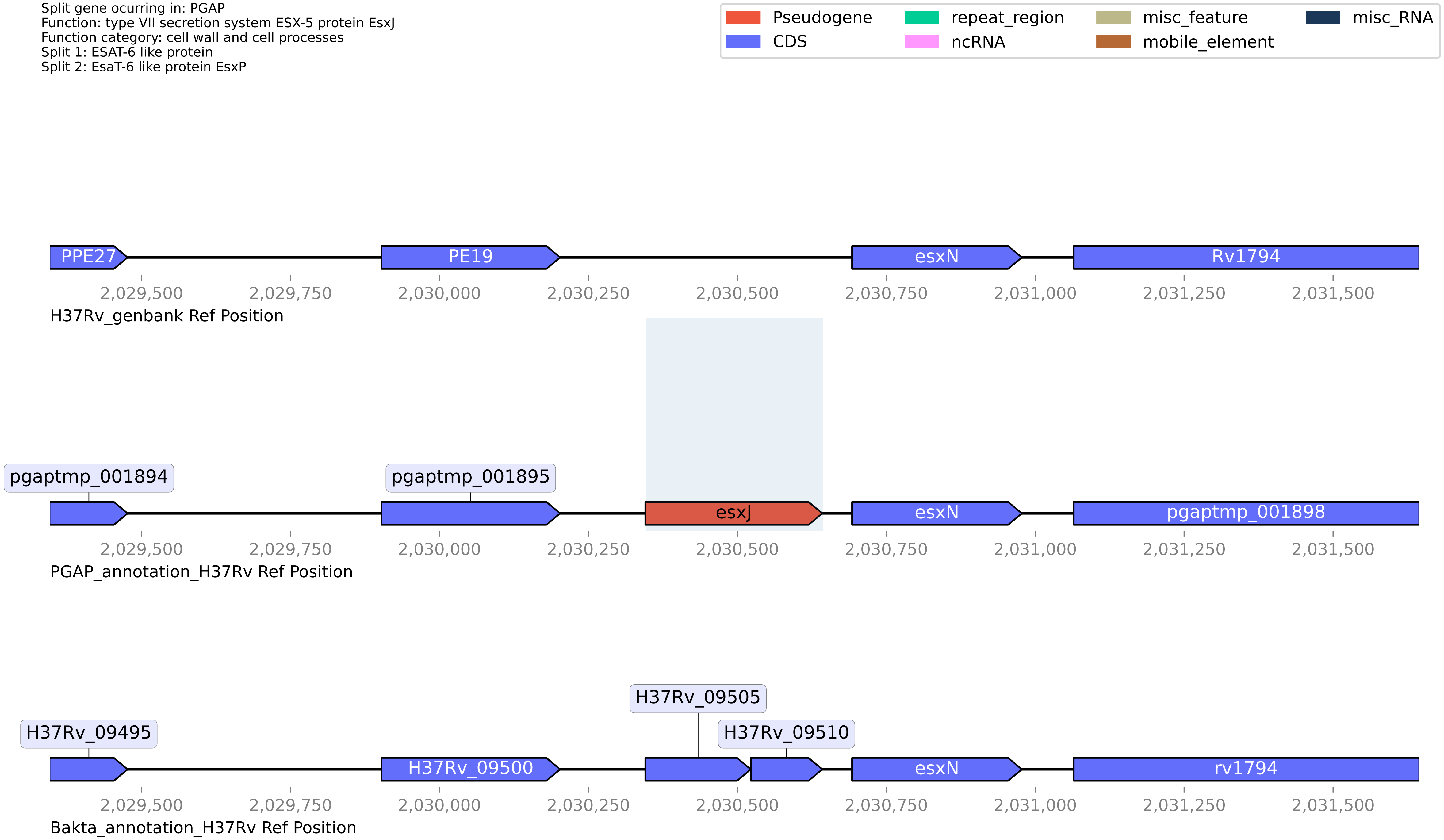

**H37Rv PGAP or Bakta split gene annotation between coordinates 366150-372764, compared to Genbank**

Split gene ocurring in: Bakta
Function: PPE family
Function category: PE/PPE
Split 1: hypothetical protein
Split 2: pseudogene

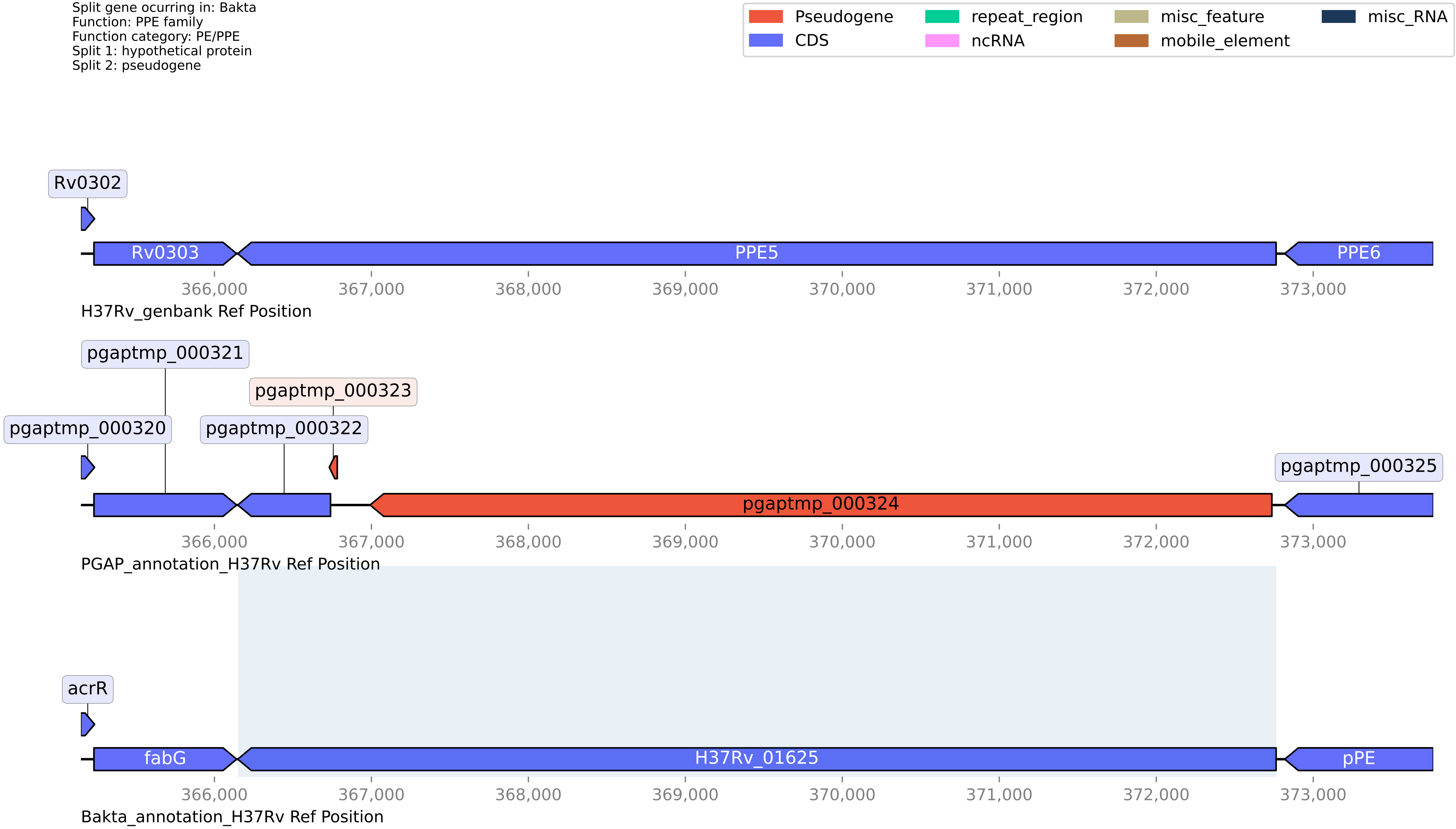

#### H37Rv PGAP or Bakta split gene annotation between coordinates 688032-689062, compared to Genbank

Split gene ocurring in: PGAP
Function: pseudogene
Function category: virulence
Split 1: Virulence factor mce family protein
Split 2: MCE-family protein

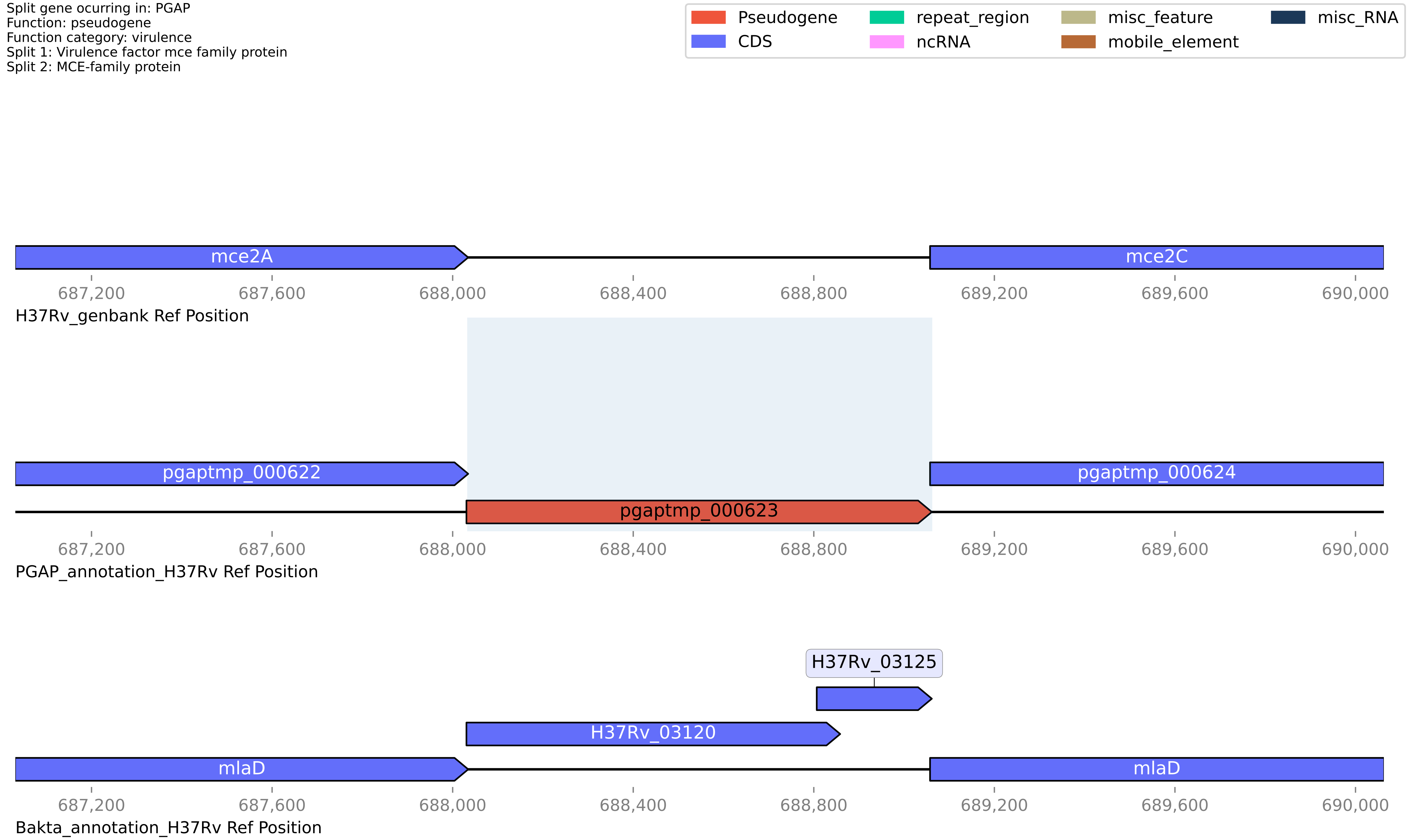

**H37Rv PGAP or Bakta split gene annotation between coordinates 4215881-4216295, compared to Genbank**

Split gene ocurring in: PGAP
Function: helix-turn-helix domain-containing protein
Function category: insertion seqs and phages
Split 1: hypothetical protein
Split 2: Transposase

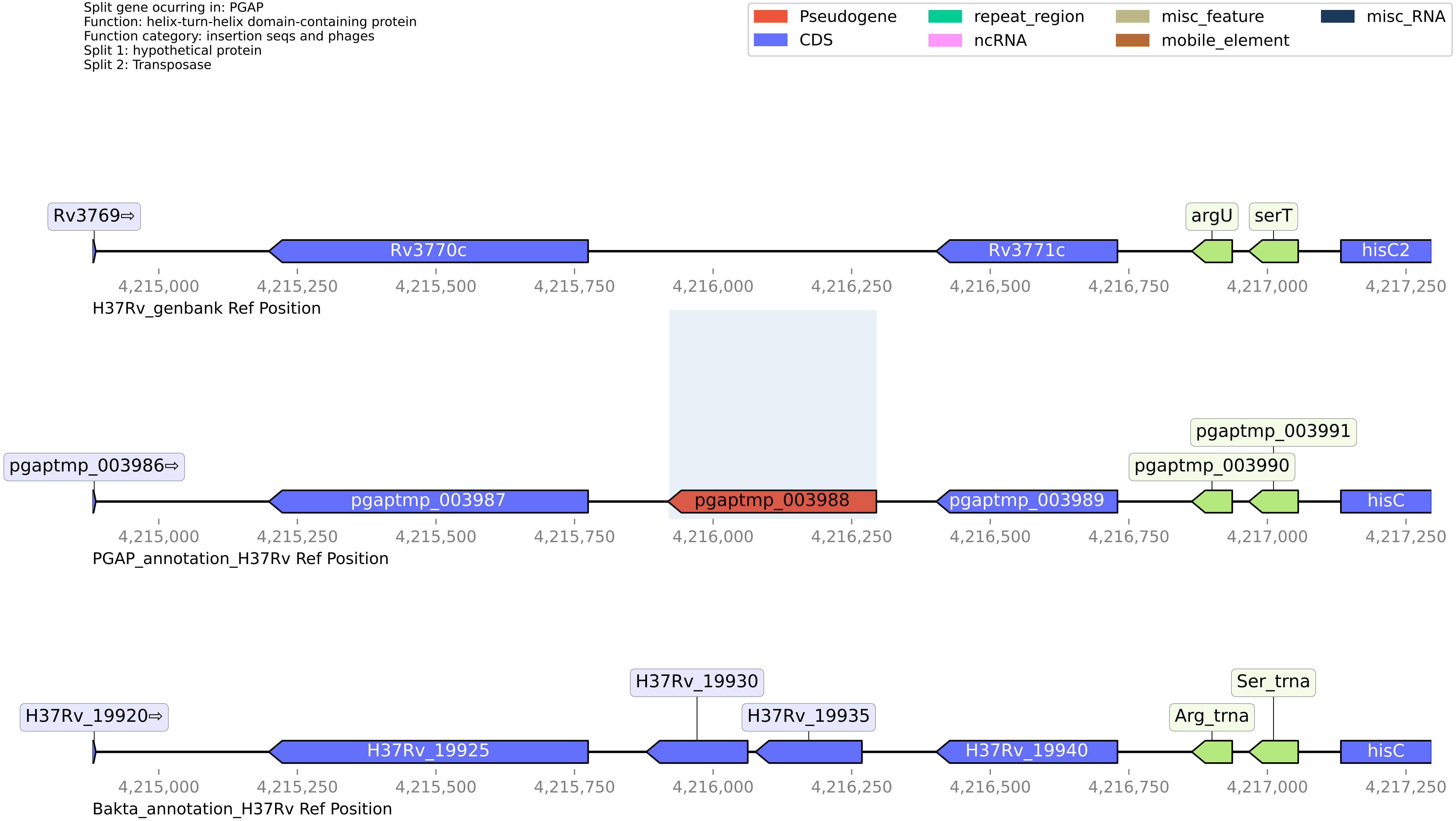

#### H37Rv PGAP or Bakta split gene annotation between coordinates 1753606-1755431, compared to Genbank

Split gene ocurring in: PGAP
Function: fatty acid--CoA ligase FadD11
Function category: lipid metabolism
Split 1: Uncharacterized protein Rv1549
Split 2: Putative fatty-acid--CoA ligase fadD11

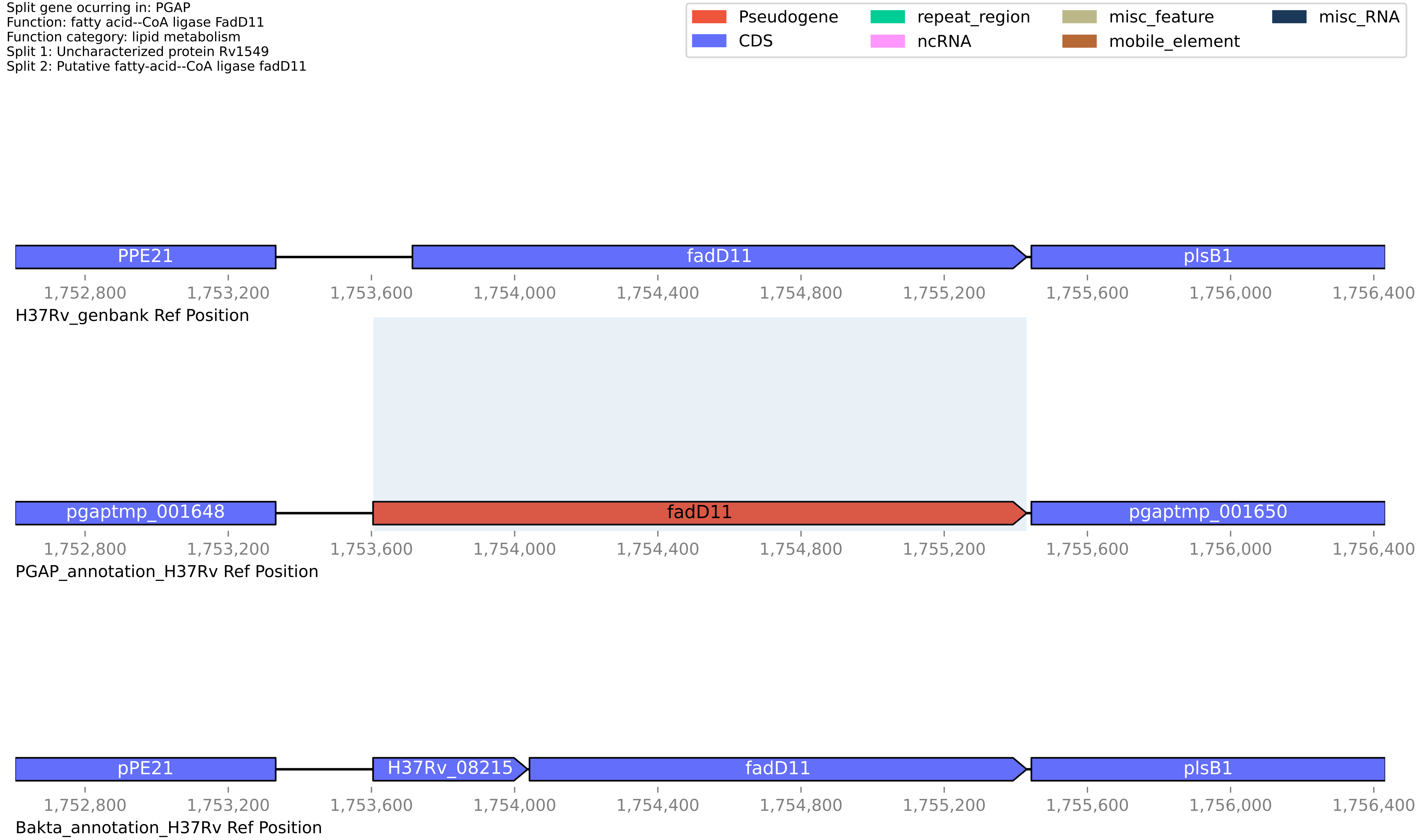

**H37Rv PGAP or Bakta split gene annotation between coordinates 2356729-2358206, compared to Genbank**

Split gene ocurring in: PGAP
Function: PE family protein
Function category: PE/PPE
Split 1: PE domain-containing protein
Split 2: PE-PGRS family protein

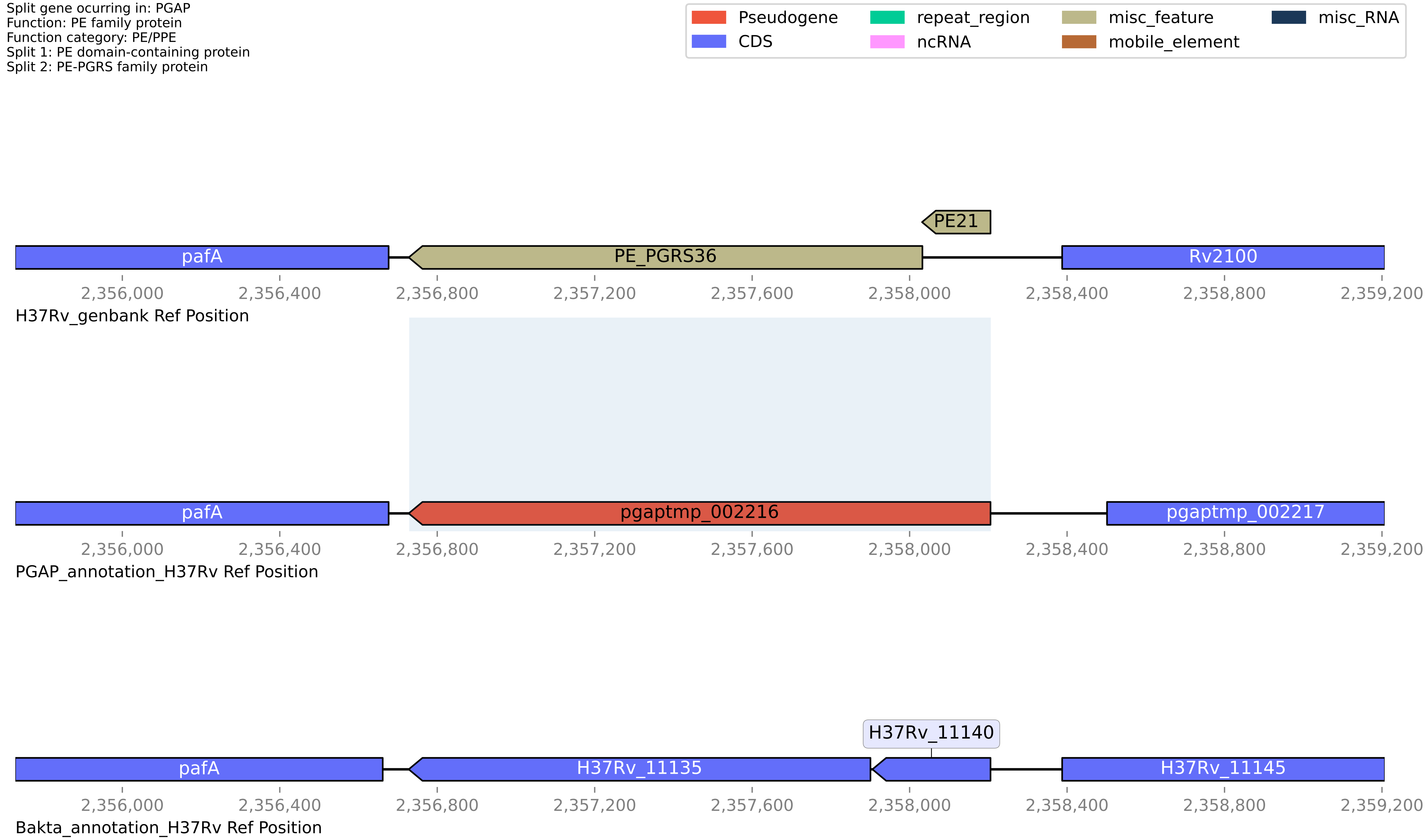

#### H37Rv PGAP or Bakta split gene annotation between coordinates 3329949-3331612, compared to Genbank

Split gene ocurring in: PGAP
Function: DAK2 domain-containing protein
Function category: conserved hypotheticals
Split 1: dihydroxyacetone kinase yloV
Split 2: DhaL domain-containing protein

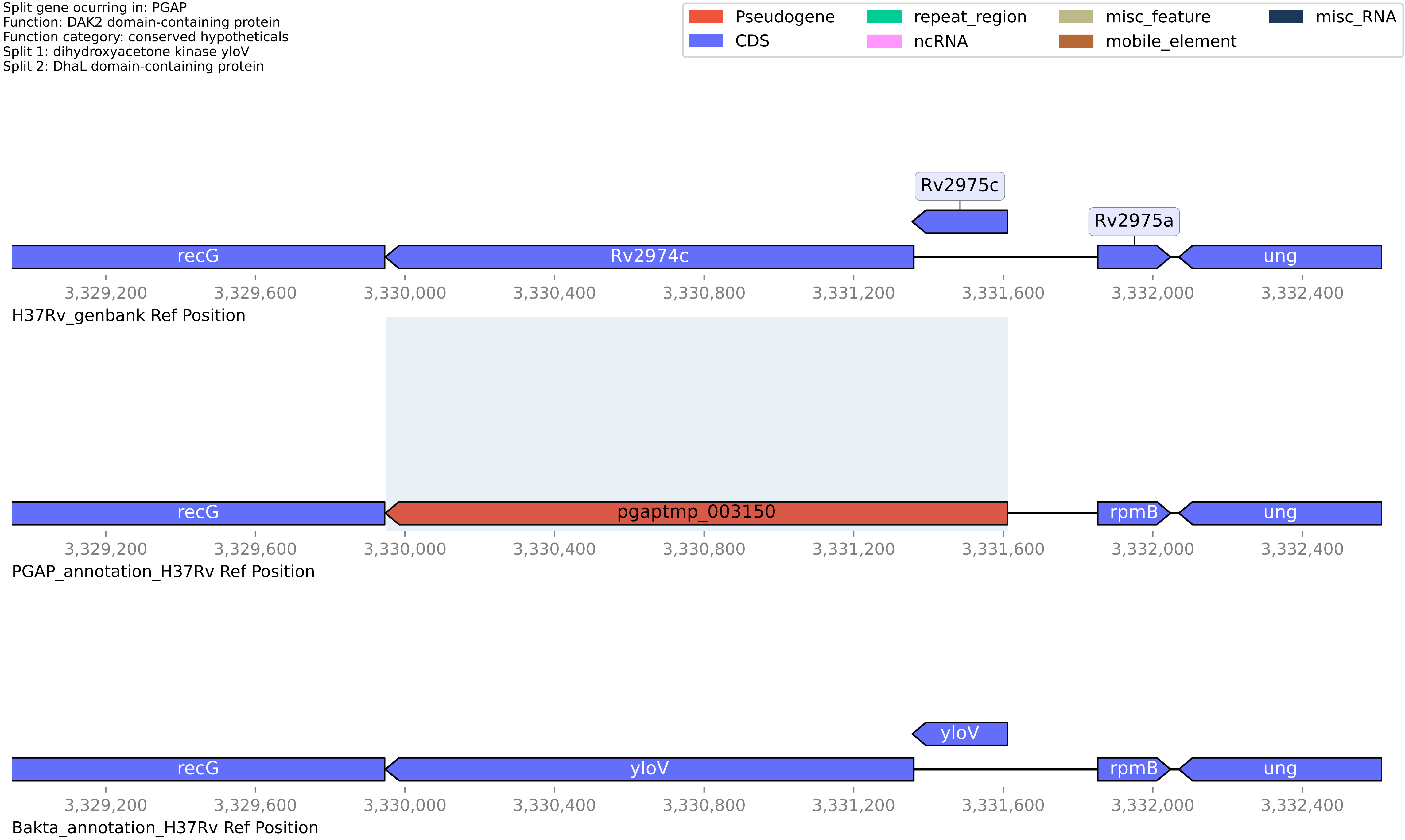

### H37Rv PGAP or Bakta split gene annotation between coordinates 103710-105101, compared to Genbank

Split gene ocurring in: PGAP
Function: pseudogene
Function category: insertion seqs and phages
Split 1: HNHc domain-containing protein
Split 2: Putative uncharacterized protein Rv0095c

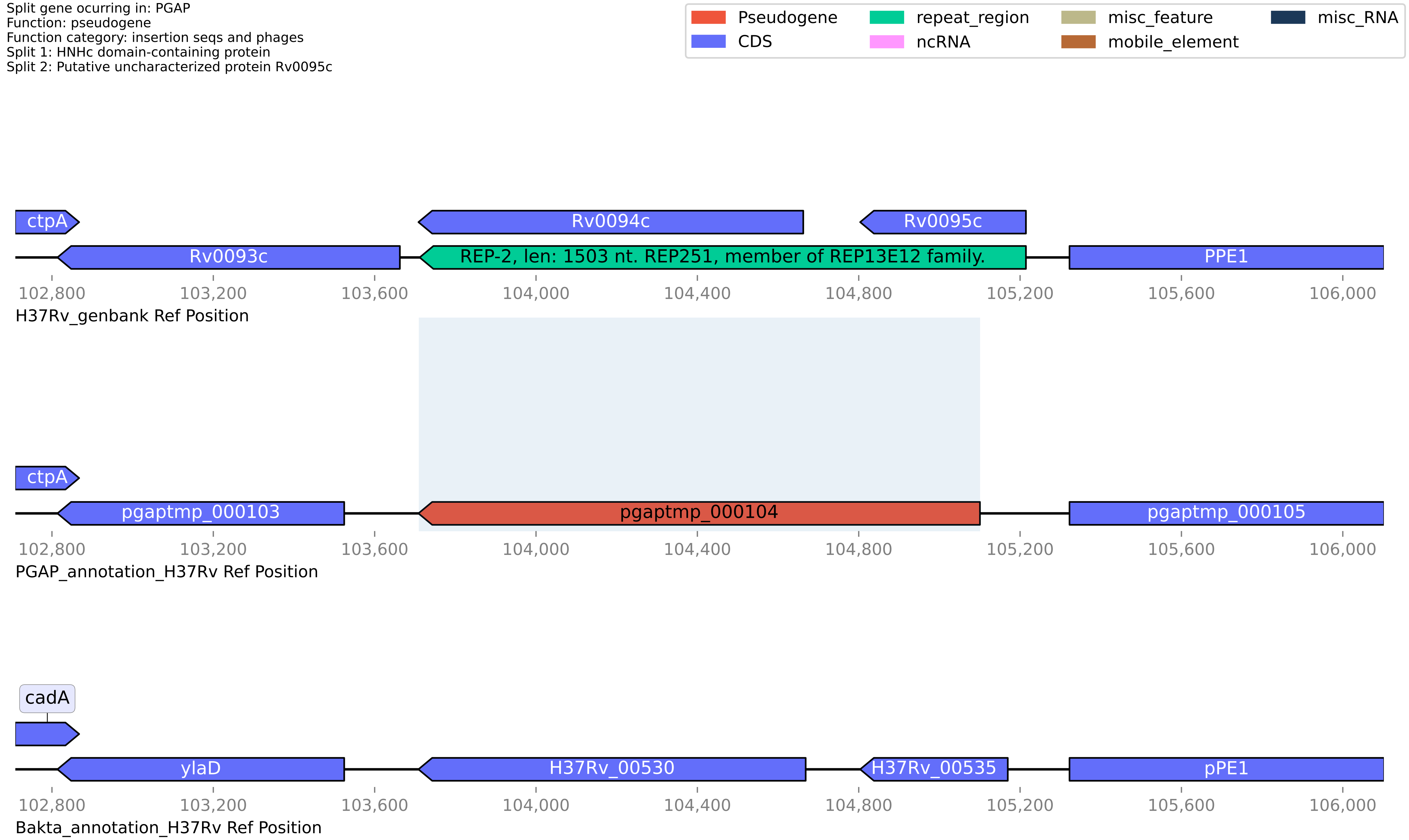

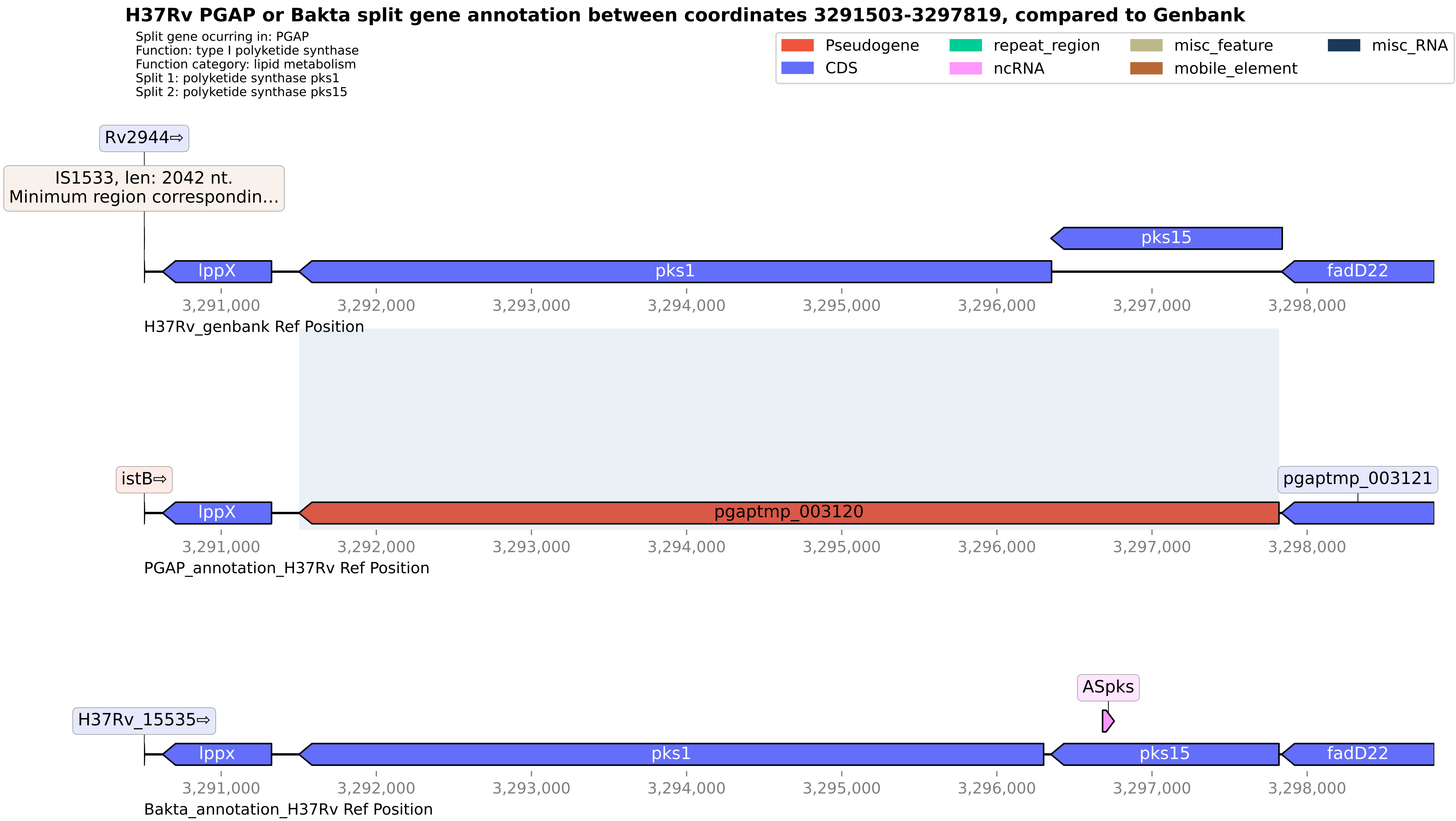
H37Rv PGAP or Bakta split gene annotation between coordinates 3291503-3297819, compared to Genbank
Split gene ocurring in: PGAP
Function: type I polyketide synthase
Function category: lipid metabolism
Split 1: polyketide synthase pks1
Split 2: polyketide synthase pks15
Pseudogene
CDS
repeat_region
ncRNA
misc_feature
mobile_element
misc_RNA
Rv2944⇨
IS1533, len: 2042 nt.
Minimum region correspondin...
pks15
lppX
pks1
fadD22
3,291,000
3,292,000
3,293,000
3,294,000
3,295,000
3,296,000
3,297,000
3,298,000
H37Rv_genbank Ref Position
istB⇨
pgaptmp_003121
lppX
pgaptmp_003120
PGAP_annotation_H37Rv Ref Position
ASpks
H37Rv_15535⇨
lppx
pks1
pks15
fadD22
Bakta_annotation_H37Rv Ref Position

#### H37Rv PGAP or Bakta split gene annotation between coordinates 1277893-1278820, compared to Genbank

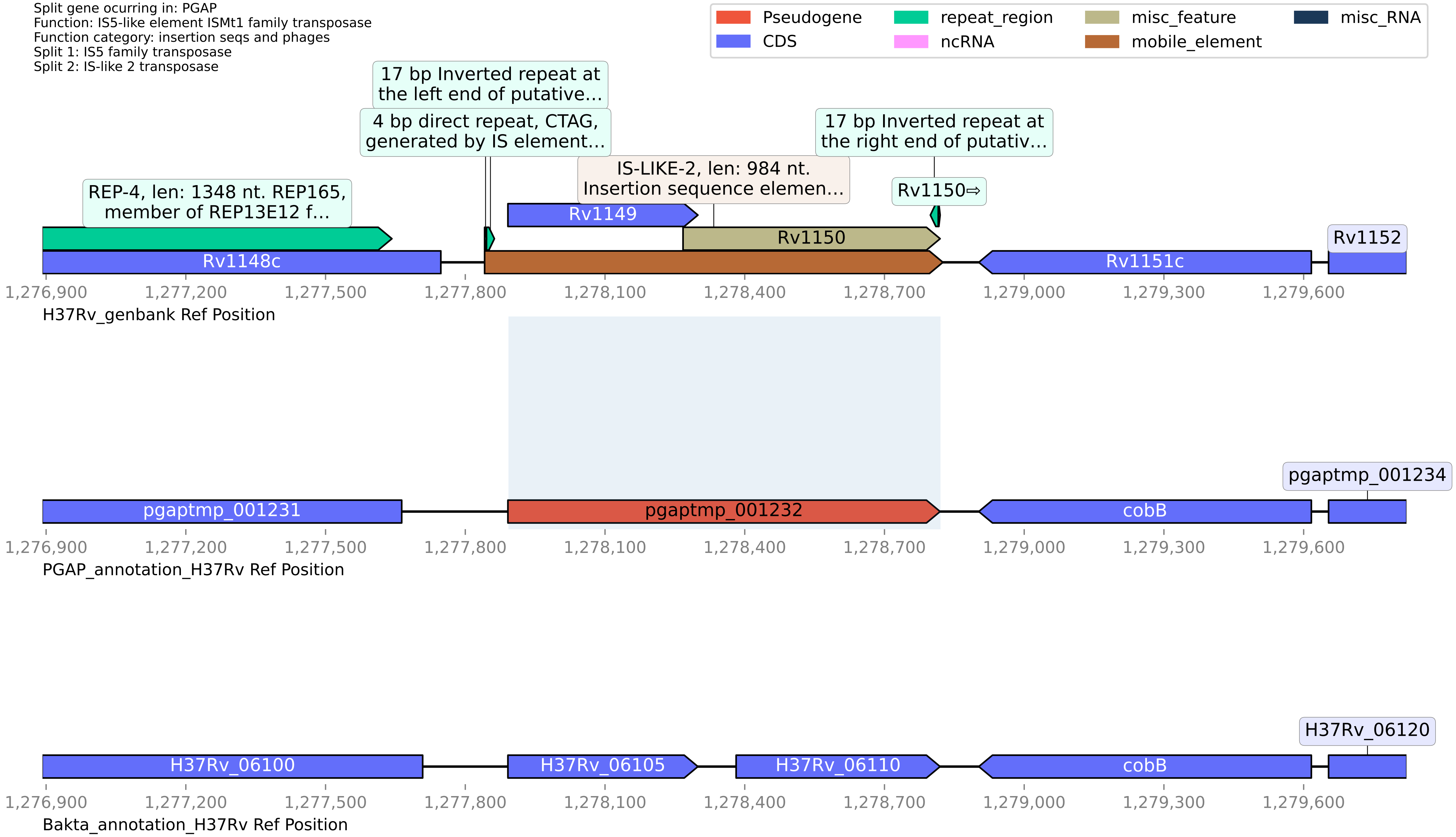

**H37Rv PGAP or Bakta split gene annotation between coordinates 1272423-1274767, compared to Genbank**

Split gene ocurring in: PGAP
Function: MMPL family transporter
Function category: cell wall and cell processes
Split 1: transporter
Split 2: MMPL family

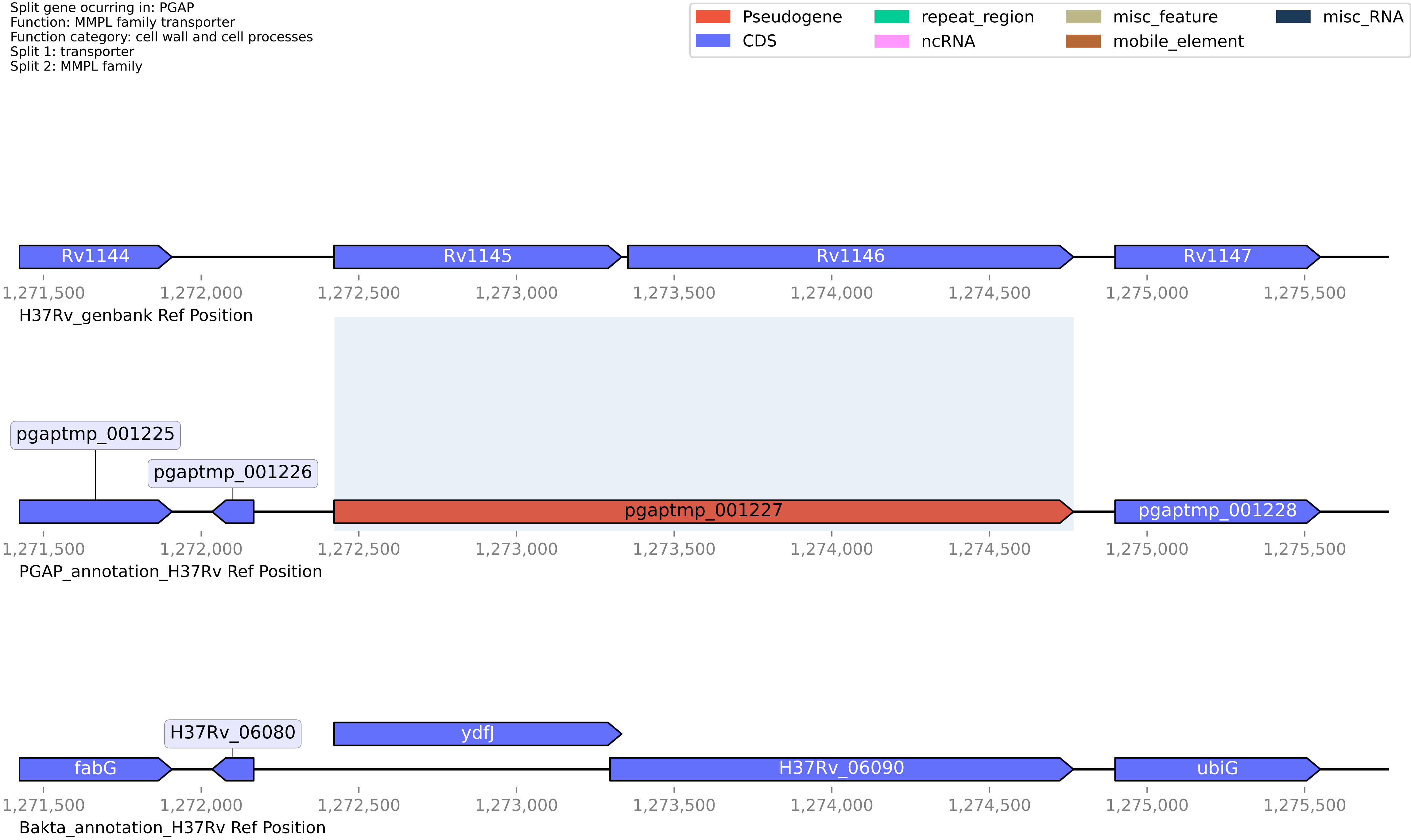

**H37Rv PGAP or Bakta split gene annotation between coordinates 4189285-4190517, compared to Genbank**

Split gene ocurring in: PGAP
Function: PPE family protein
Function category: PE/PPE
Split 1: Uncharacterized PPE family protein PPE66
Split 2: Uncharacterized PPE family protein PPE66

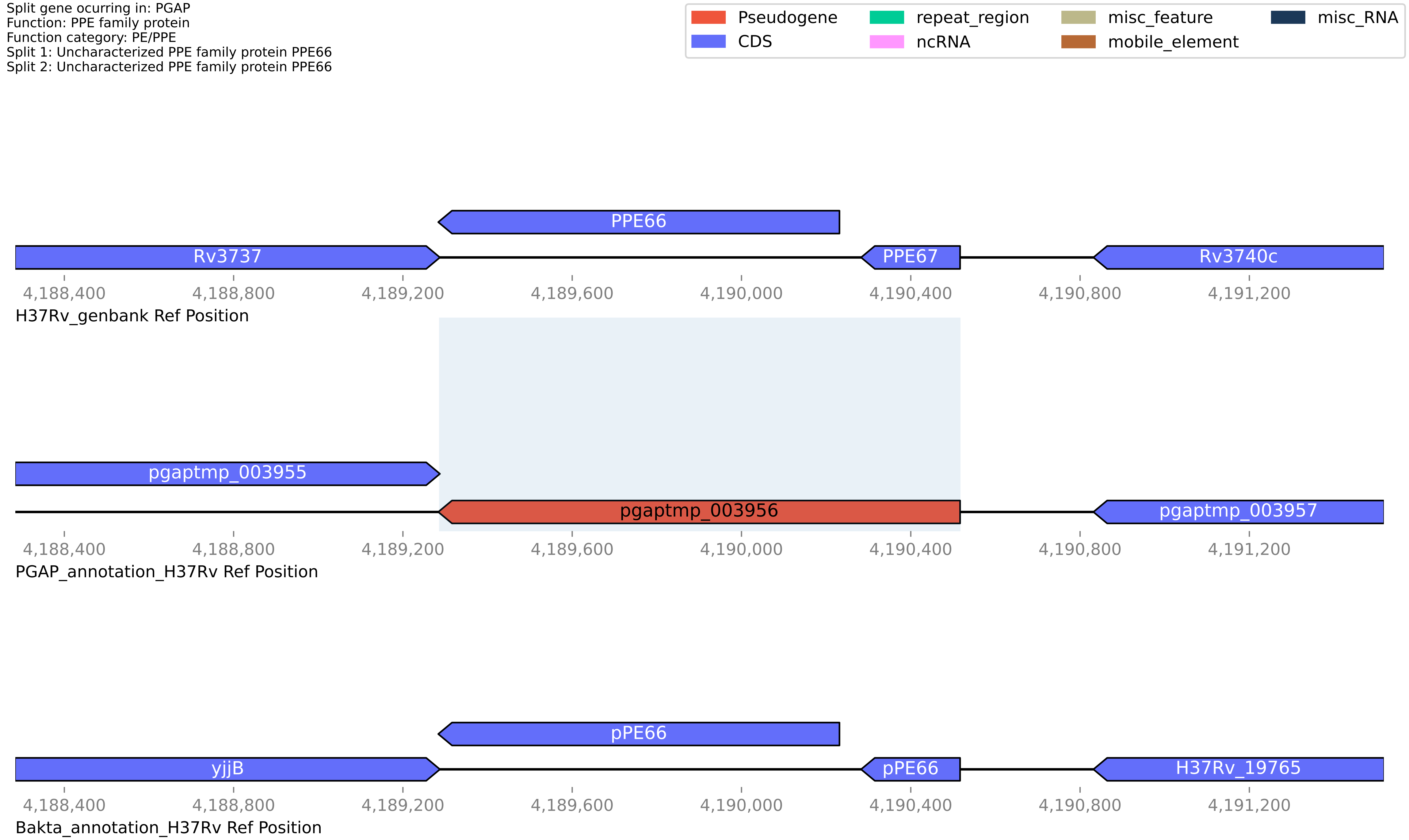

**H37Rv PGAP or Bakta split gene annotation between coordinates 1158918-1160358, compared to Genbank**

Split gene ocurring in: PGAP
Function: ISNCY family transposase
Function category: insertion seqs and phages
Split 1: Transposase
Split 2: Putative transposase

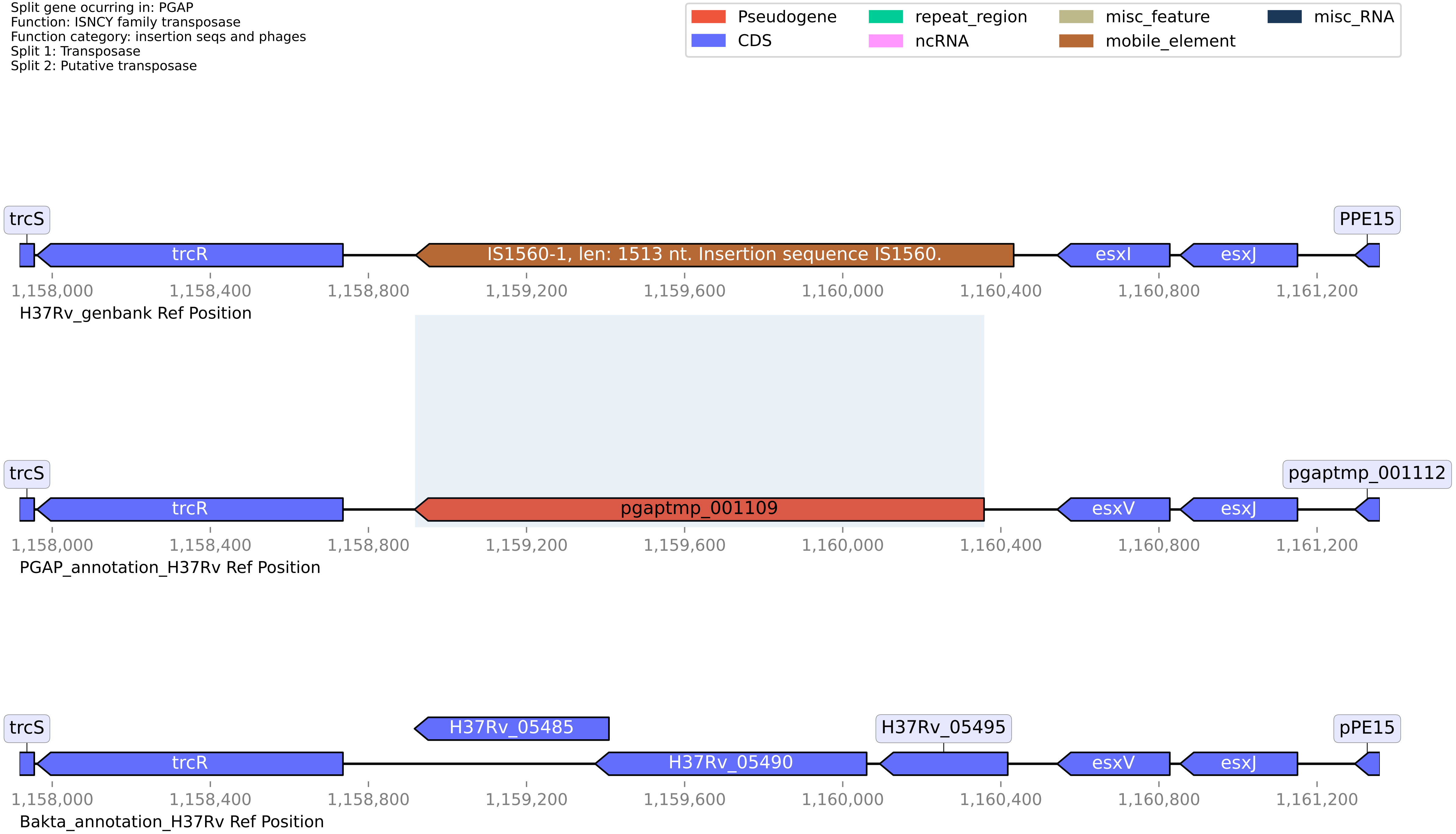

**H37Rv PGAP or Bakta split gene annotation between coordinates 1231301-1232837, compared to Genbank**

Split gene ocurring in: PGAP
Function: carboxylesterase/lipase family protein
Function category: intermediary metabolism and respiration
Split 1: Para-nitrobenzyl esterase
Split 2: Para-nitrobenzyl esterase

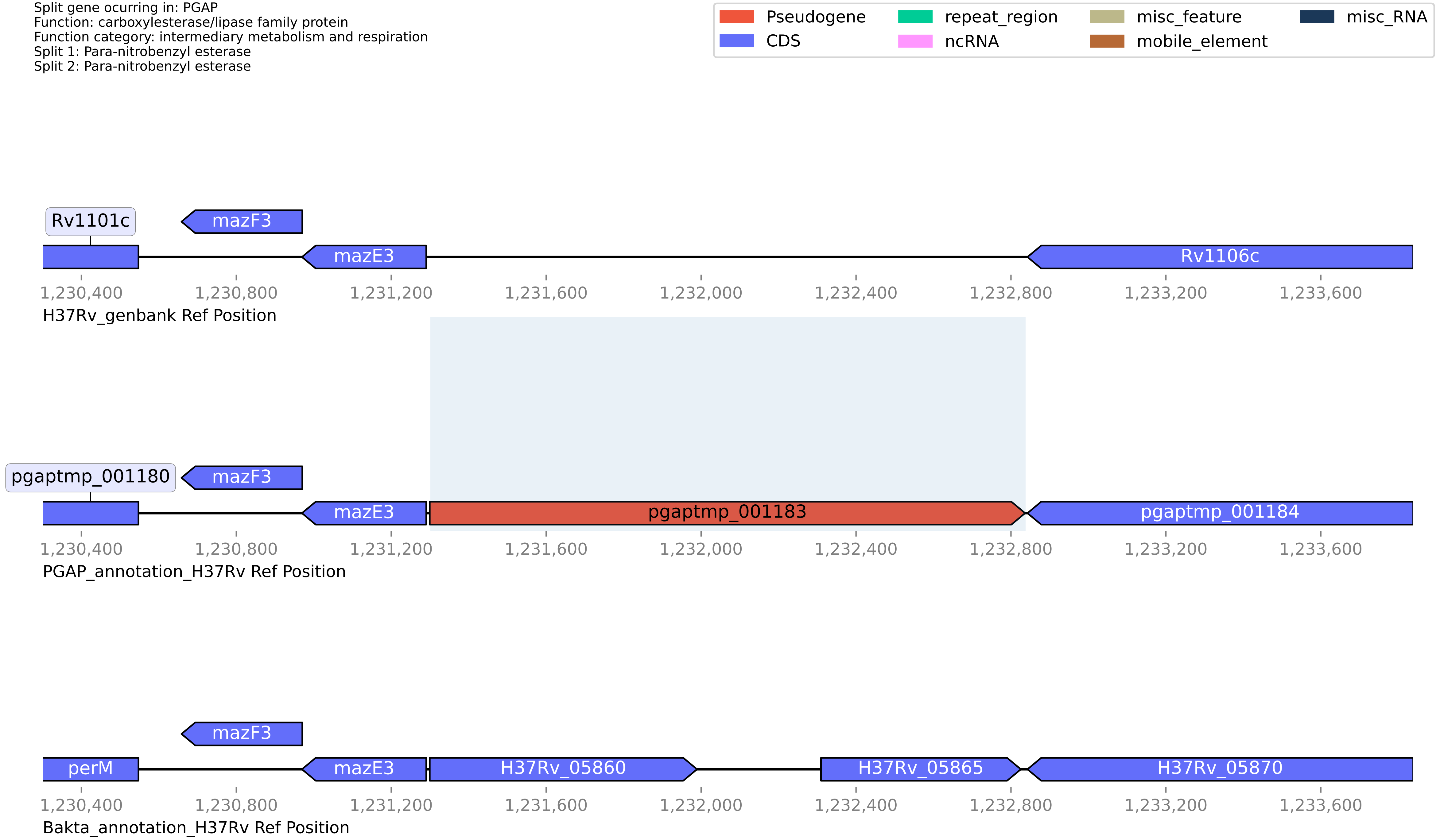

#### H37Rv PGAP or Bakta split gene annotation between coordinates 1164572-1165499, compared to Genbank

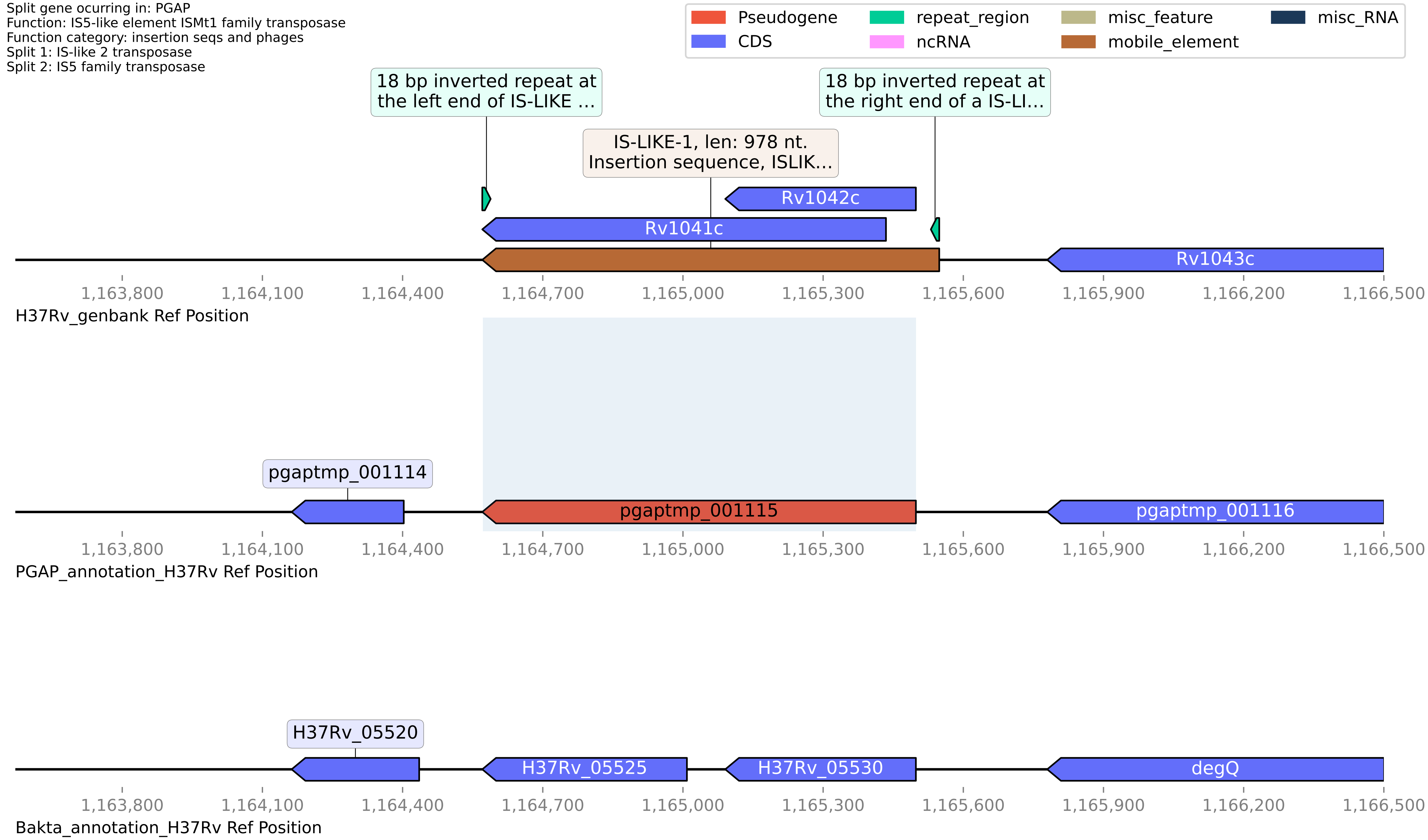

**H37Rv PGAP or Bakta split gene annotation between coordinates 1313725-1319982, compared to Genbank**

Split gene ocurring in: PGAP
Function: sulfolipid-1 biosynthesis phthioceranic/hydroxyphthioceranic acid synthase pks2 gene
Function category: lipid metabolism
Split 1: Mycolipanoate synthase
Split 2: polyketide synthase

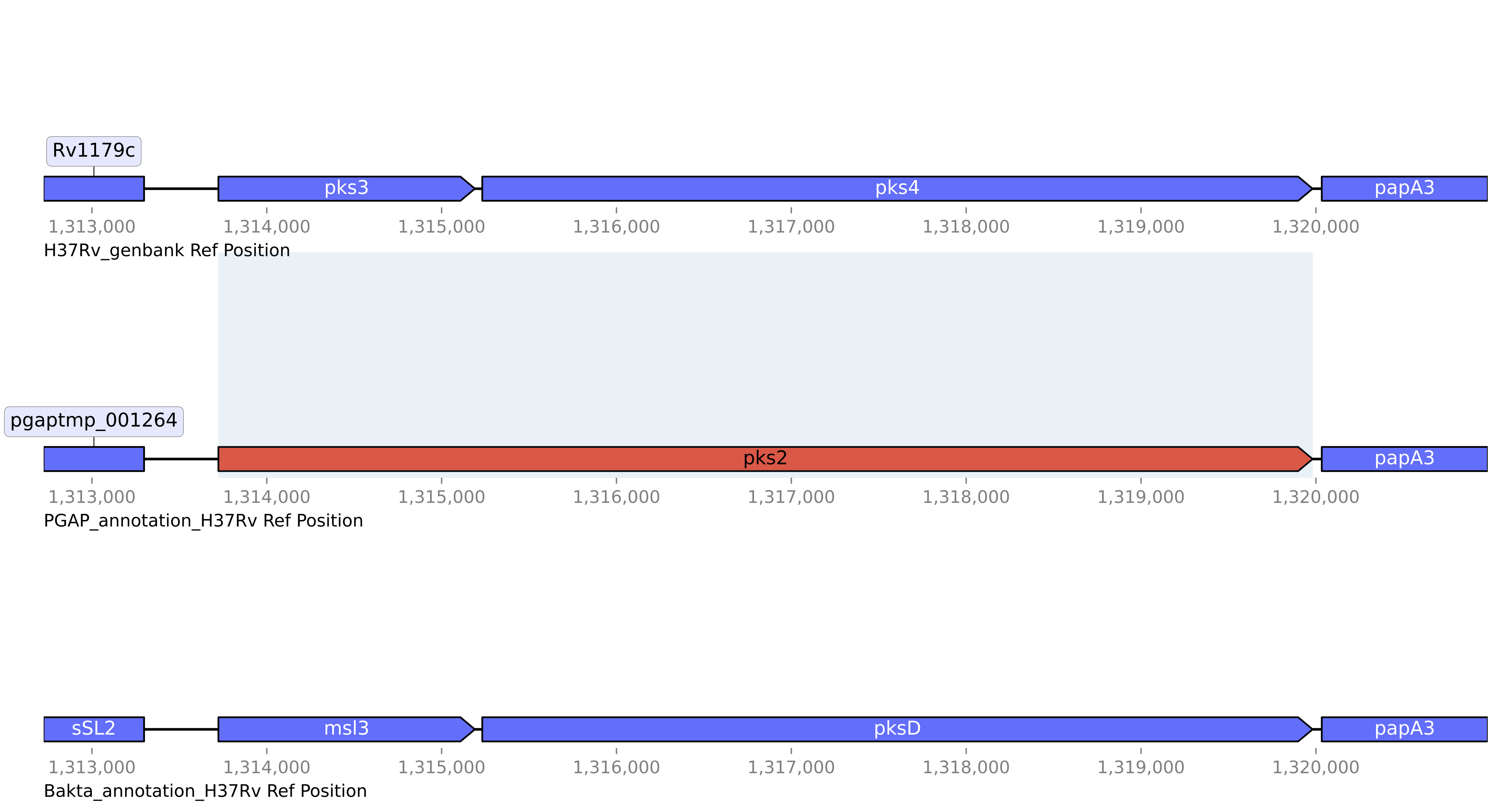

#### H37Rv PGAP or Bakta split gene annotation between coordinates 2534042-2535552, compared to Genbank

Split gene ocurring in: PGAP
Function: apolipoprotein N-acyltransferase Int
Function category: lipid metabolism
Split 1: CN hydrolase domain-containing protein
Split 2: apolipoprotein N-acyltransferase Int

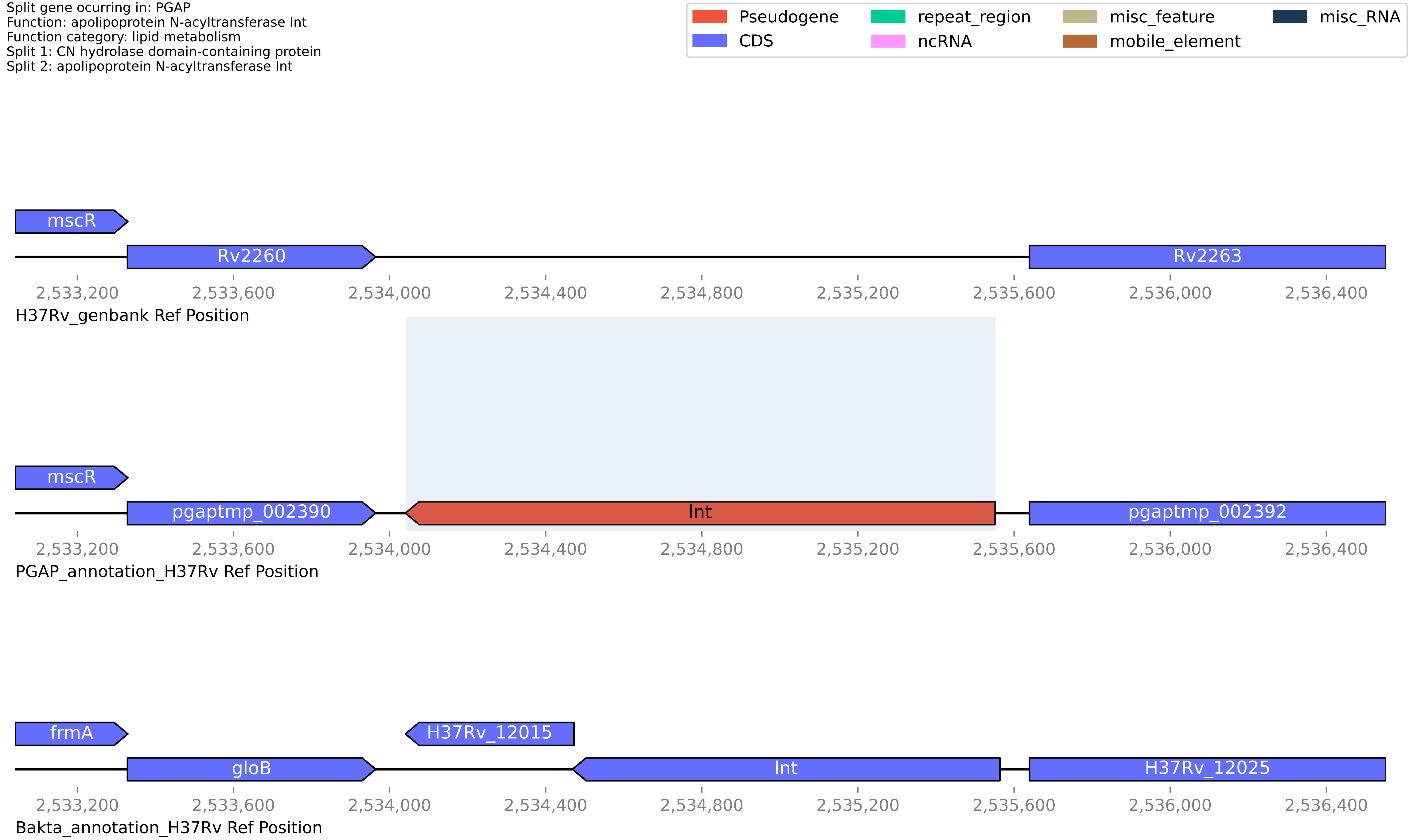

**H37Rv PGAP or Bakta split gene annotation between coordinates 1242864-1243634, compared to Genbank**

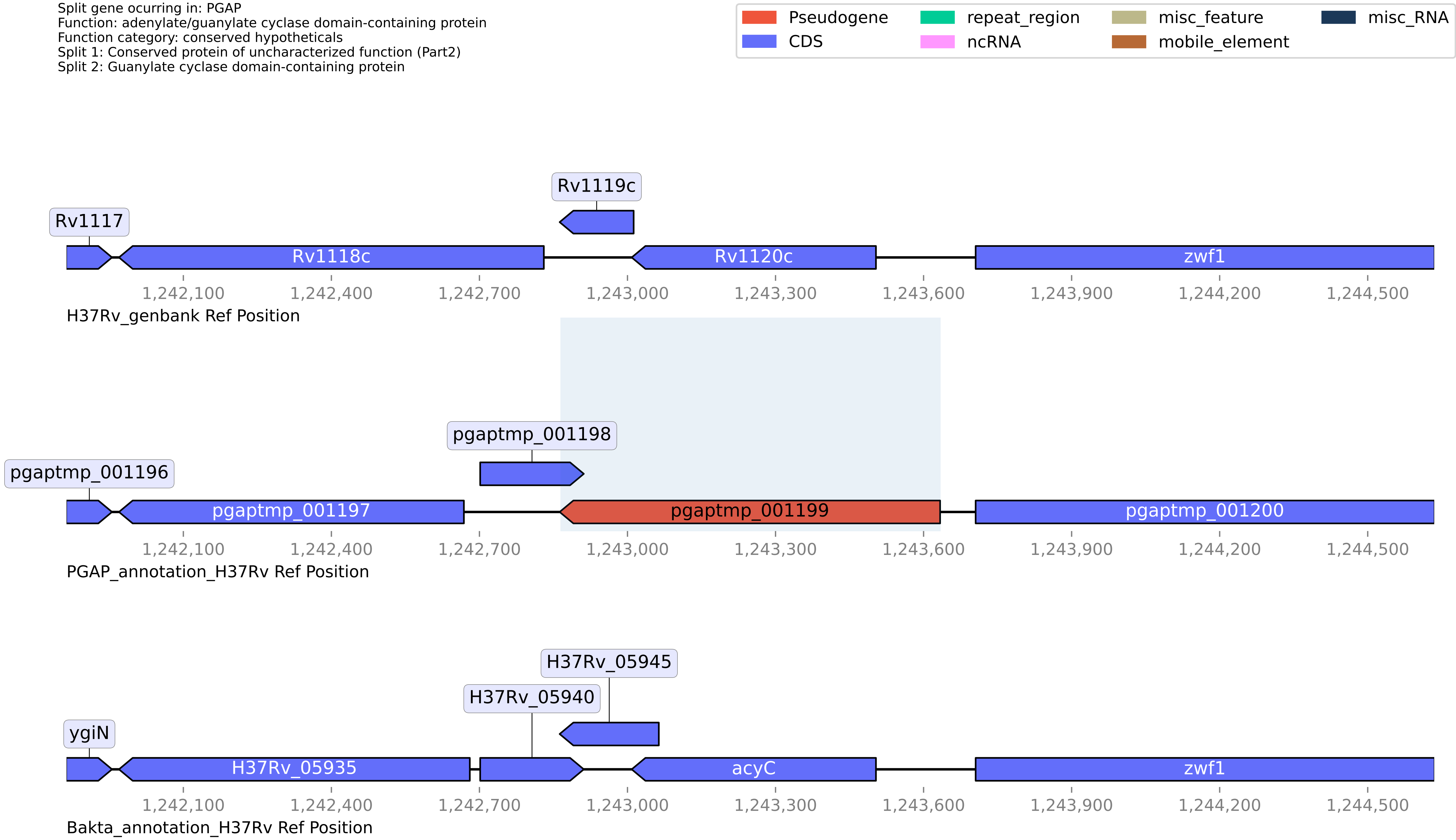

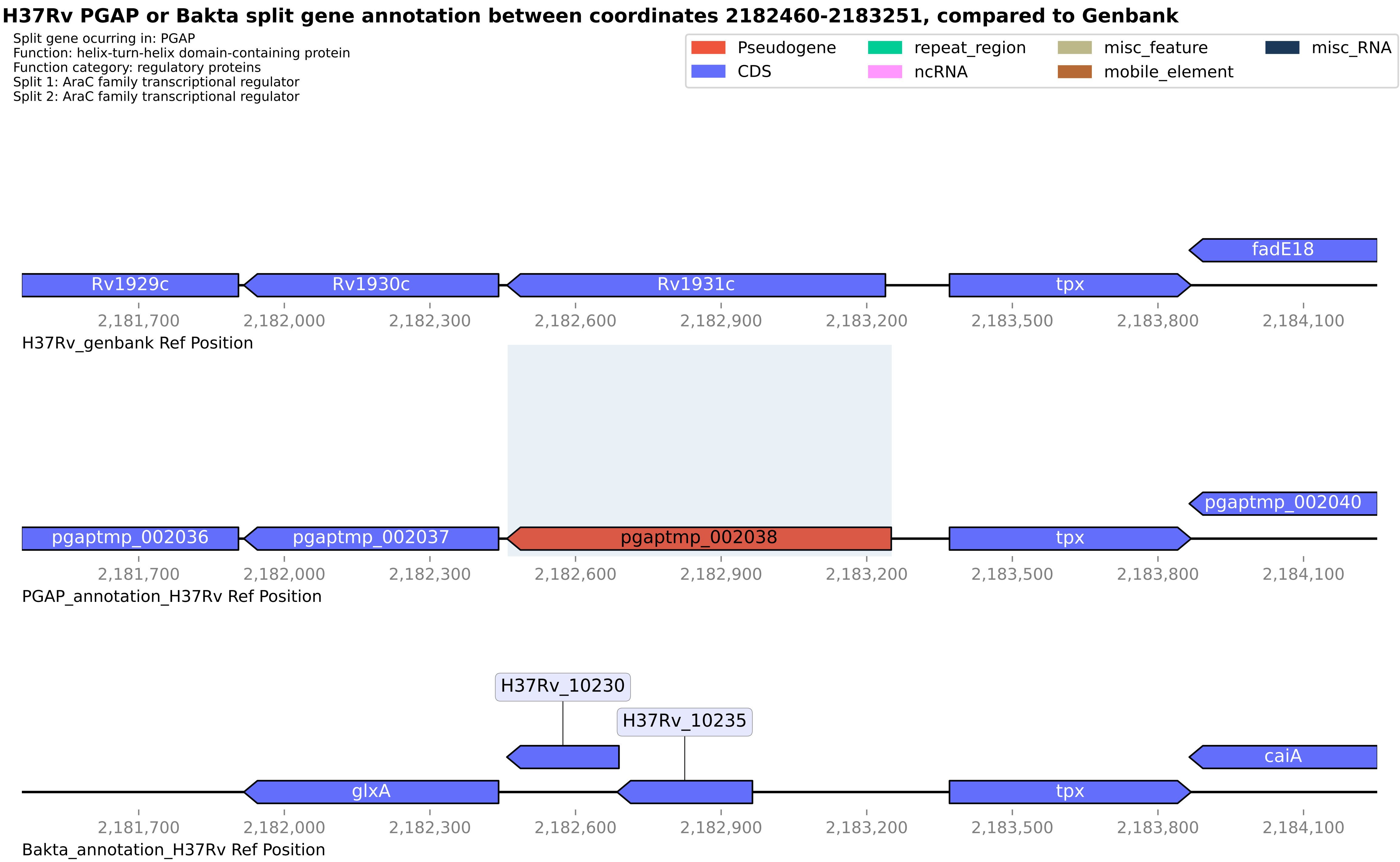

H37Rv PGAP or Bakta split gene annotation between coordinates 2182460-2183251, compared to Genbank
Split gene ocurring in: PGAP
Function: helix-turn-helix domain-containing protein
Function category: regulatory proteins
Split 1: AraC family transcriptional regulator
Split 2: AraC family transcriptional regulator
Pseudogene
CDS
repeat_region
ncRNA
misc_feature
mobile_element
misc_RNA
fadE18
Rv1929c
Rv1930c
Rv1931c
tpx
2,181,700
2,182,000
2,182,300
2,182,600
2,182,900
2,183,200
2,183,500
2,183,800
2,184,100
H37Rv_genbank Ref Position
pgaptmp_002040
pgaptmp_002036
pgaptmp_002037
pgaptmp_002038
tpx
PGAP_annotation_H37Rv Ref Position
H37Rv_10230
H37Rv_10235
caiA
glxA
tpx
Bakta_annotation_H37Rv Ref Position

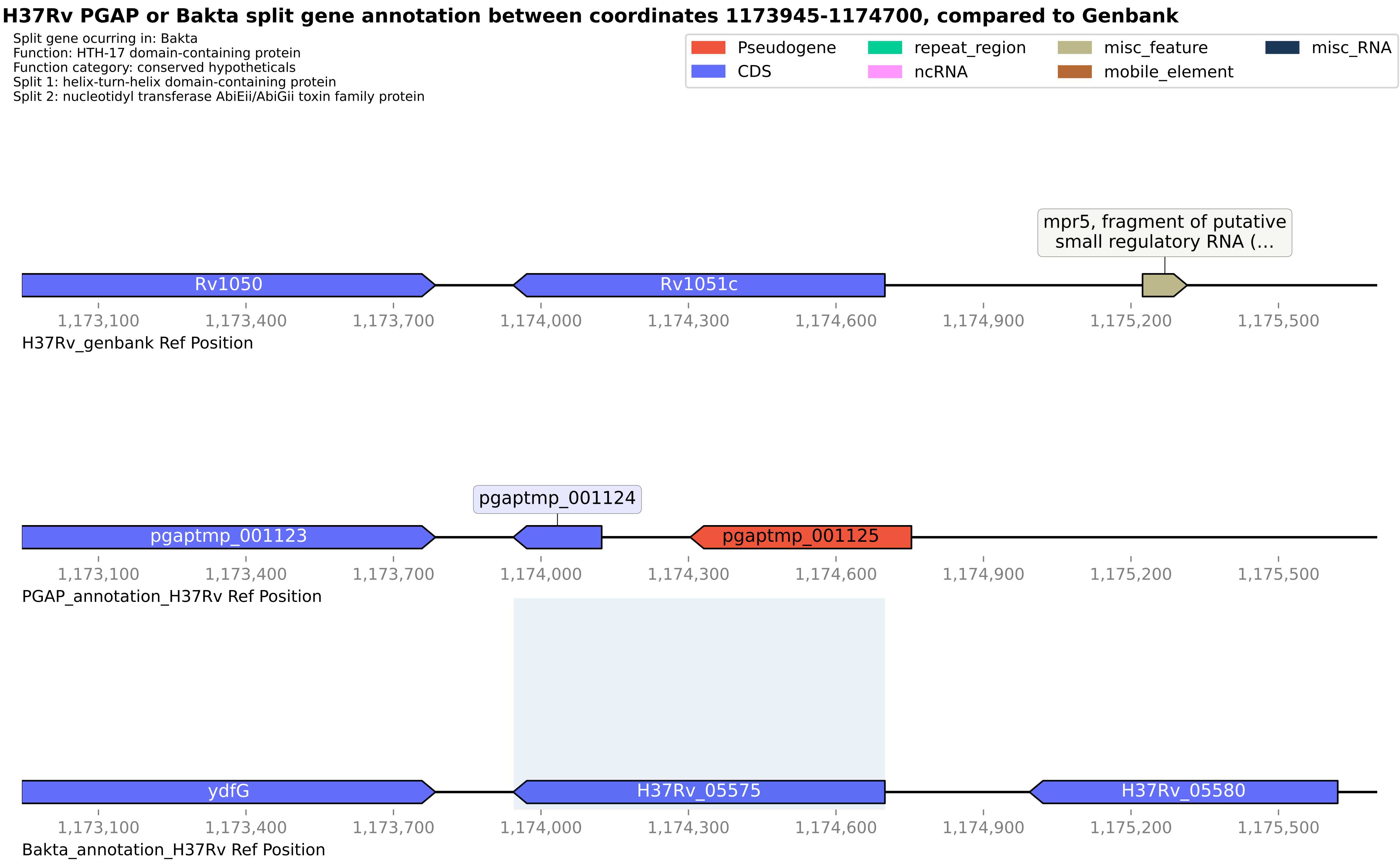
H37Rv PGAP or Bakta split gene annotation between coordinates 1173945-1174700, compared to Genbank
Split gene ocurring in: Bakta
Function: HTH-17 domain-containing protein
Function category: conserved hypotheticals
Split 1: helix-turn-helix domain-containing protein
Split 2: nucleotidyl transferase AbiEii/AbiGii toxin family protein
Pseudogene
CDS
repeat_region
ncRNA
misc_feature
mobile_element
misc_RNA
mpr5, fragment of putative small regulatory RNA (...
Rv1050
Rv1051c
1,173,100
1,173,400
1,173,700
1,174,000
1,174,300
1,174,600
1,174,900
1,175,200
1,175,500
H37Rv_genbank Ref Position
pgaptmp_001124
pgaptmp_001123
pgaptmp_001125
PGAP_annotation_H37Rv Ref Position
ydfG
H37Rv_05575
H37Rv_05580
Bakta_annotation_H37Rv Ref Position

#### H37Rv PGAP or Bakta split gene annotation between coordinates 3609781-3611189, compared to Genbank

Split gene ocurring in: PGAP
Function: wax ester/triacylglycerol synthase family O-acyltransferase
Function category: lipid metabolism
Split 1: Diacylglycerol O-acyltransferase
Split 2: putative diacyglycerol O-acyltransferase tgs3

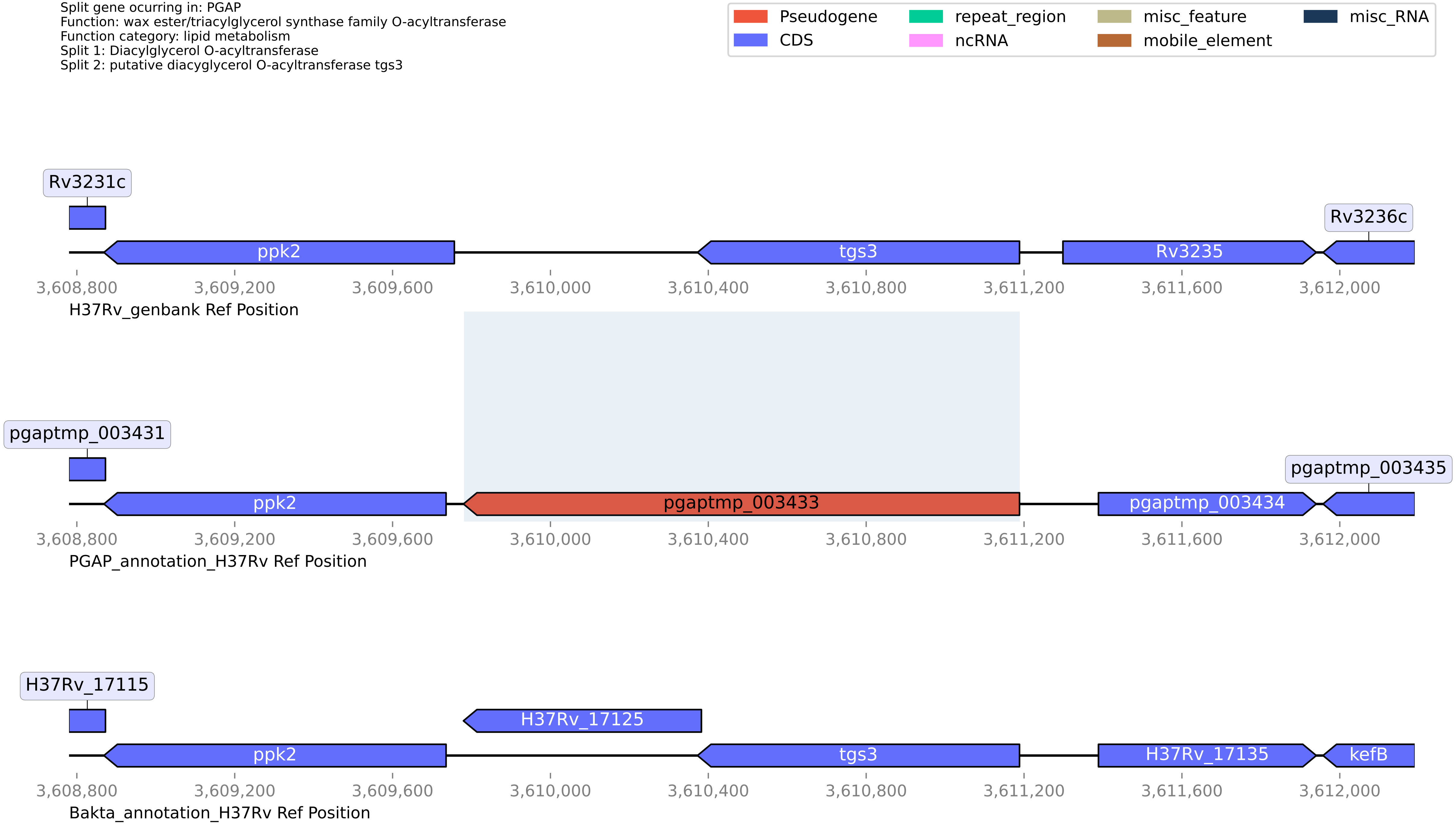

**H37Rv PGAP or Bakta split gene annotation between coordinates 3874404-3876090, compared to Genbank**

Split gene ocurring in: PGAP
Function: hypothetical protein
Function category: cell wall and cell processes
Split 1: Transmembrane protein
Split 2: Transmembrane protein

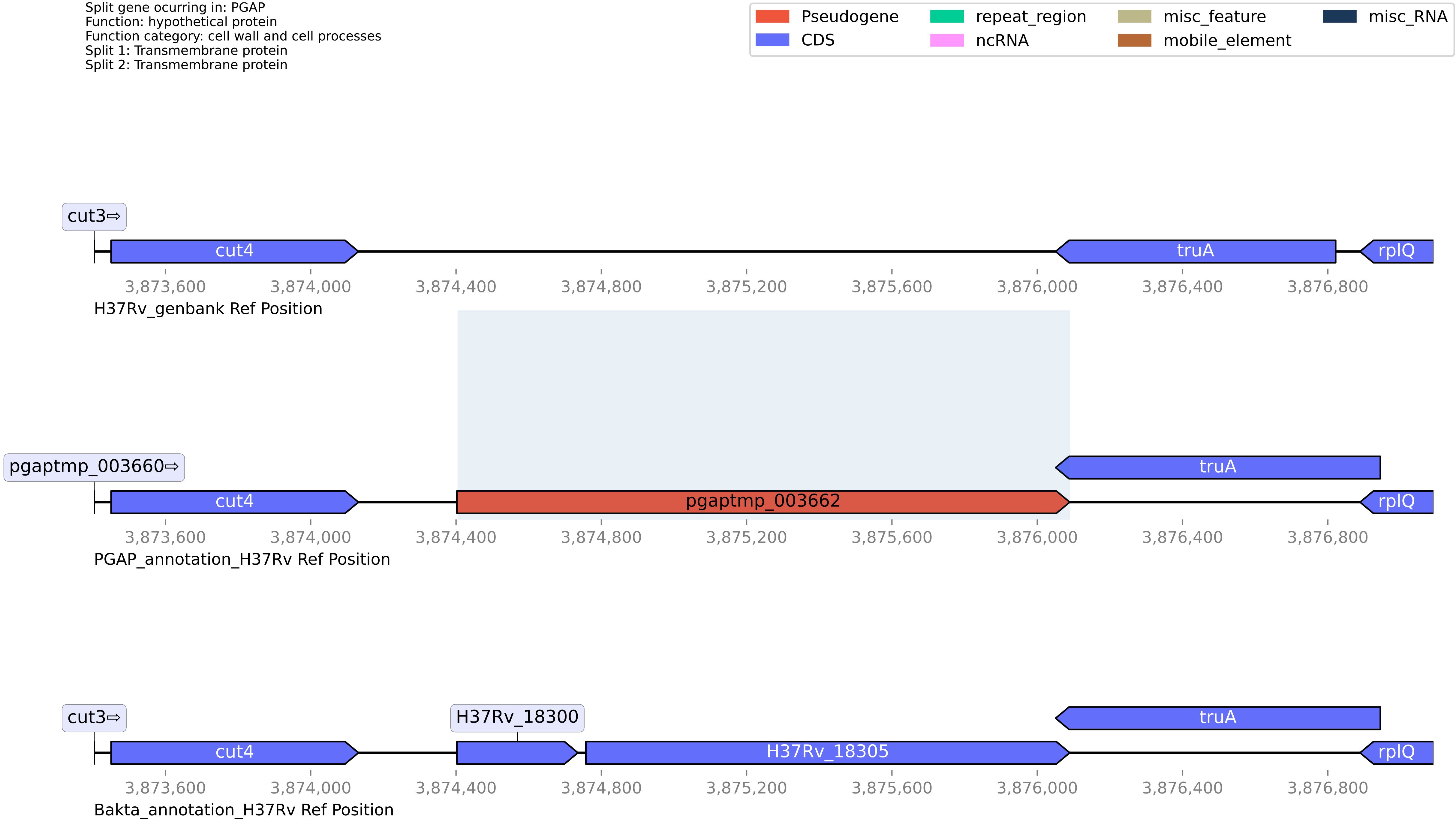

#### H37Rv PGAP or Bakta split gene annotation between coordinates 1589199-1590292, compared to Genbank

Split gene ocurring in: PGAP
Function: alanine racemase
Function category: conserved hypotheticals
Split 1: Uncharacterized protein Mb1448
Split 2: Uncharacterized protein Rv1414

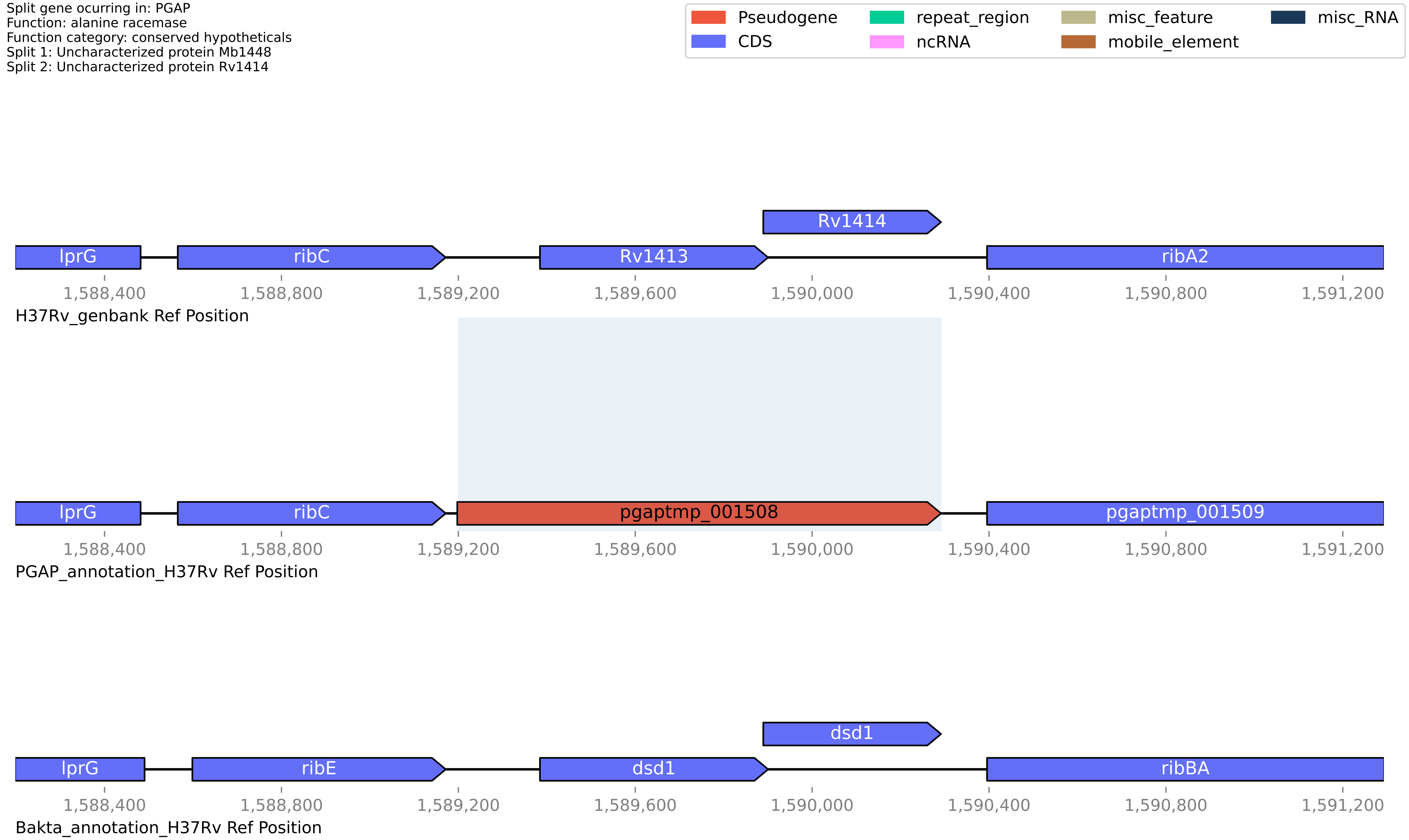

**H37Rv PGAP or Bakta split gene annotation between coordinates 4075752-4076984, compared to Genbank**

Split gene ocurring in: PGAP
Function: IS21 family transposase
Function category: insertion seqs and phages
Split 1: putative transposase
Split 2: IS21 family transposase

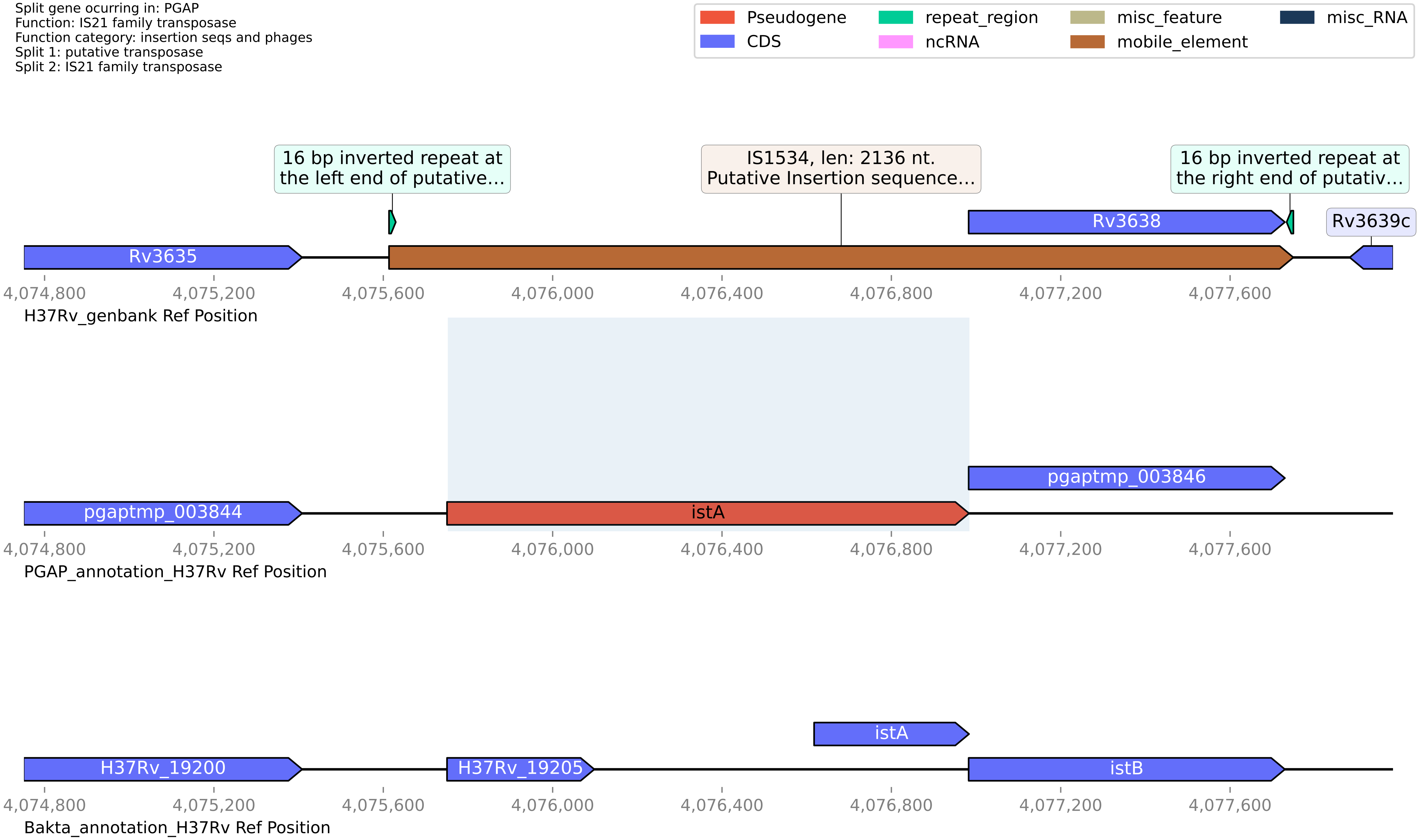

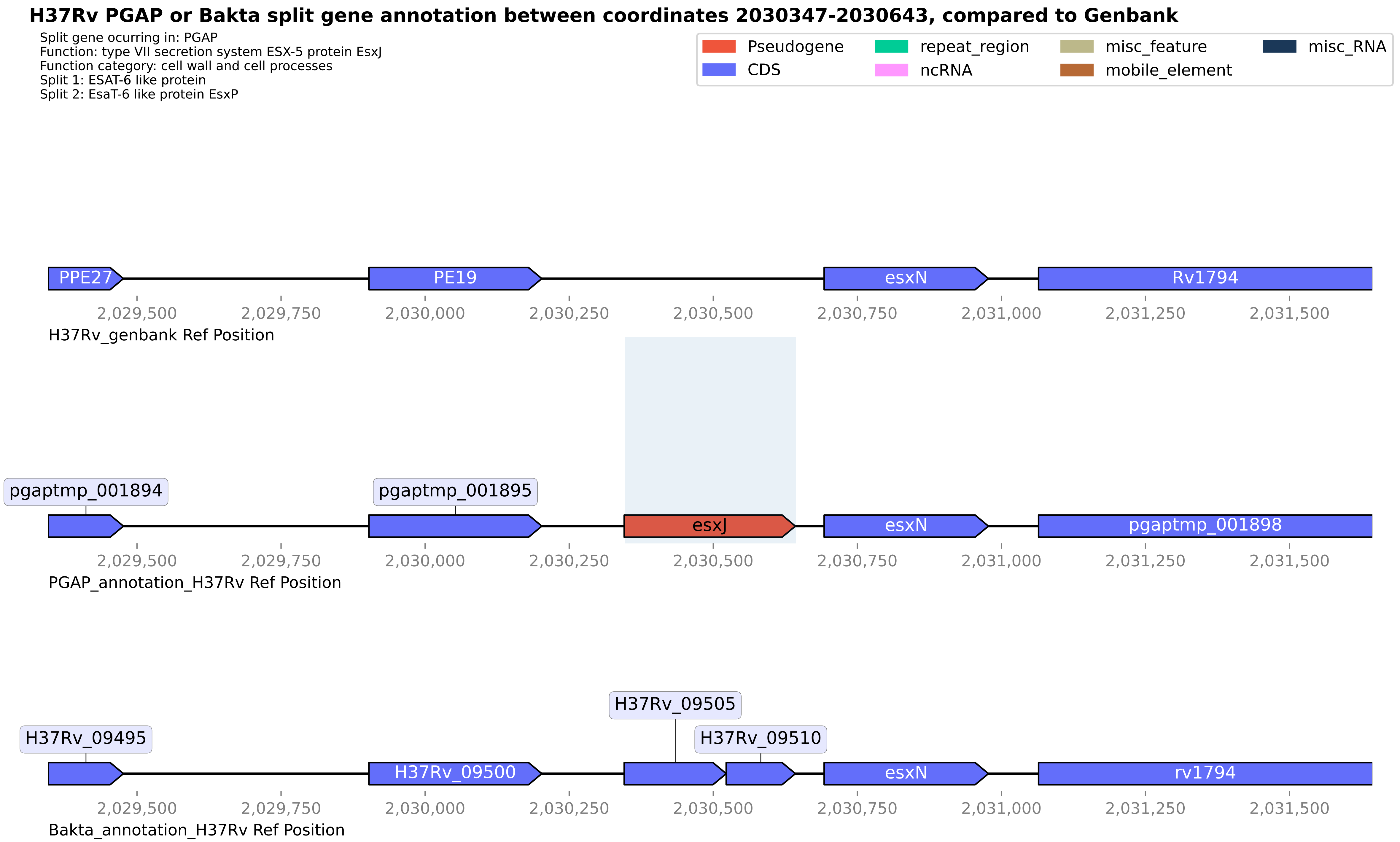

H37Rv PGAP or Bakta split gene annotation between coordinates 2030347-2030643, compared to Genbank
Split gene ocurring in: PGAP
Function: type VII secretion system ESX-5 protein EsxJ
Function category: cell wall and cell processes
Split 1: ESAT-6 like protein
Split 2: EsaT-6 like protein EsxP
Pseudogene
CDS
repeat_region
ncRNA
misc_feature
mobile_element
misc_RNA
PPE27
PE19
esxN
Rv1794
2,029,500
2,029,750
2,030,000
2,030,250
2,030,500
2,030,750
2,031,000
2,031,250
2,031,500
H37Rv_genbank Ref Position
pgaptmp_001894
pgaptmp_001895
esxJ
esxN
pgaptmp_001898
PGAP_annotation_H37Rv Ref Position
H37Rv_09495
H37Rv_09500
H37Rv_09505
H37Rv_09510
esxN
rv1794
Bakta_annotation_H37Rv Ref Position

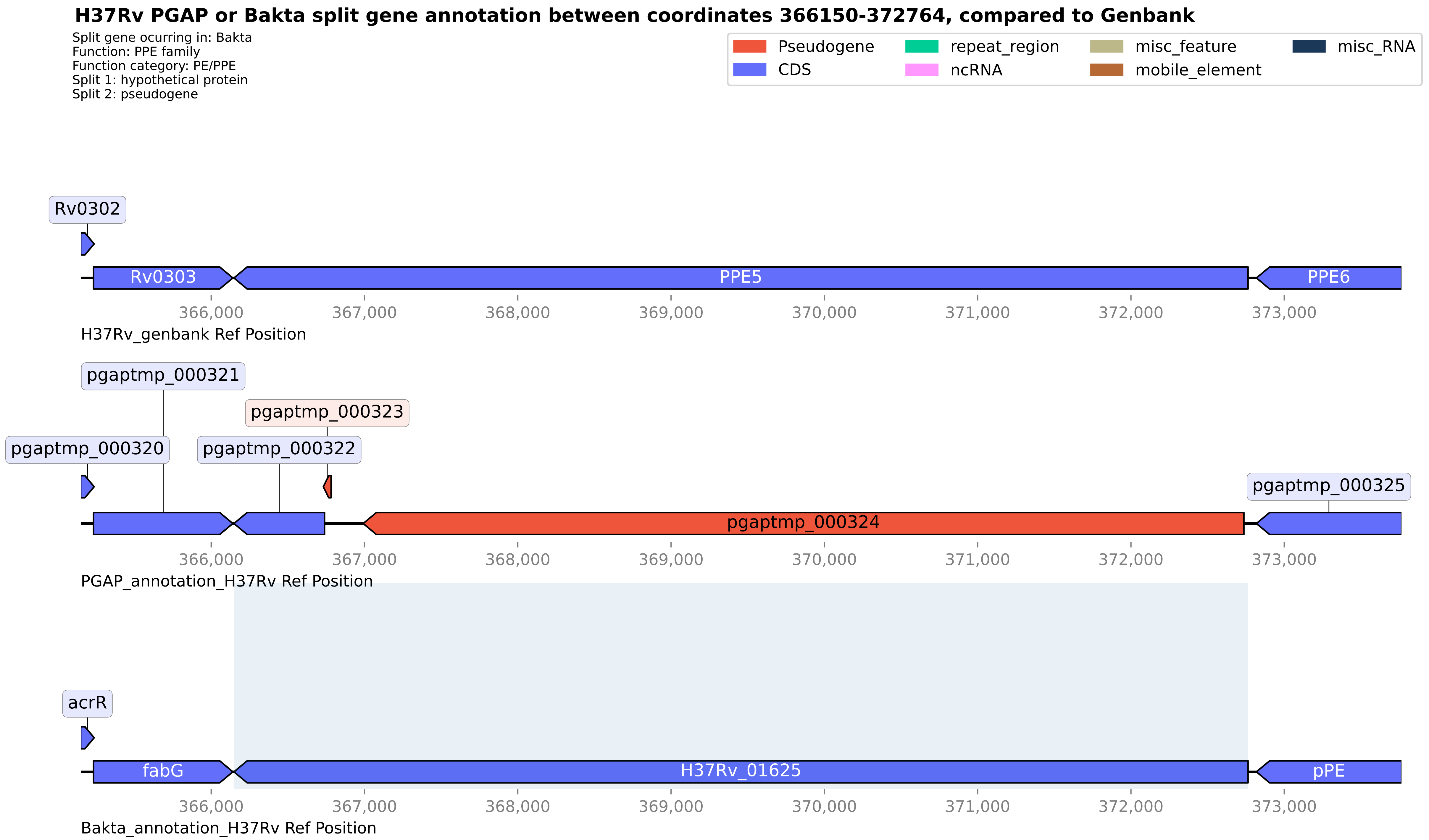
H37Rv PGAP or Bakta split gene annotation between coordinates 366150-372764, compared to Genbank
Split gene ocurring in: Bakta
Function: PPE family
Function category: PE/PPE
Split 1: hypothetical protein
Split 2: pseudogene
Pseudogene
CDS
repeat_region
ncRNA
misc_feature
mobile_element
misc_RNA
Rv0302
Rv0303
PPE5
PPE6
366,000
367,000
368,000
369,000
370,000
371,000
372,000
373,000
H37Rv_genbank Ref Position
pgaptmp_000321
pgaptmp_000323
pgaptmp_000320
pgaptmp_000322
pgaptmp_000325
pgaptmp_000324
PGAP_annotation_H37Rv Ref Position
acrR
fabG
H37Rv_01625
pPE
Bakta_annotation_H37Rv Ref Position

**H37Rv PGAP or Bakta split gene annotation between coordinates 688032-689062, compared to Genbank**

Split gene ocurring in: PGAP
Function: pseudogene
Function category: virulence
Split 1: Virulence factor mce family protein
Split 2: MCE-family protein

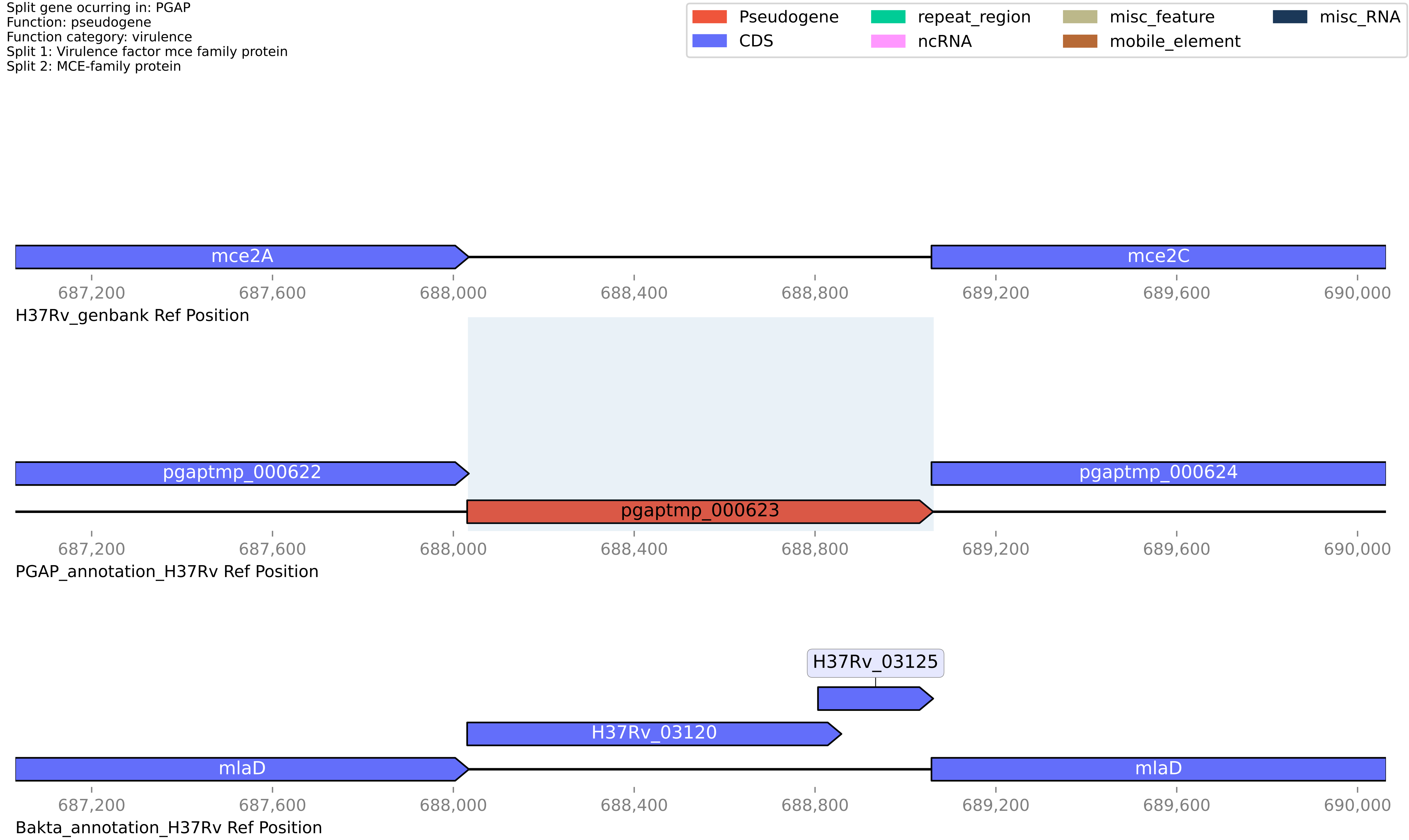

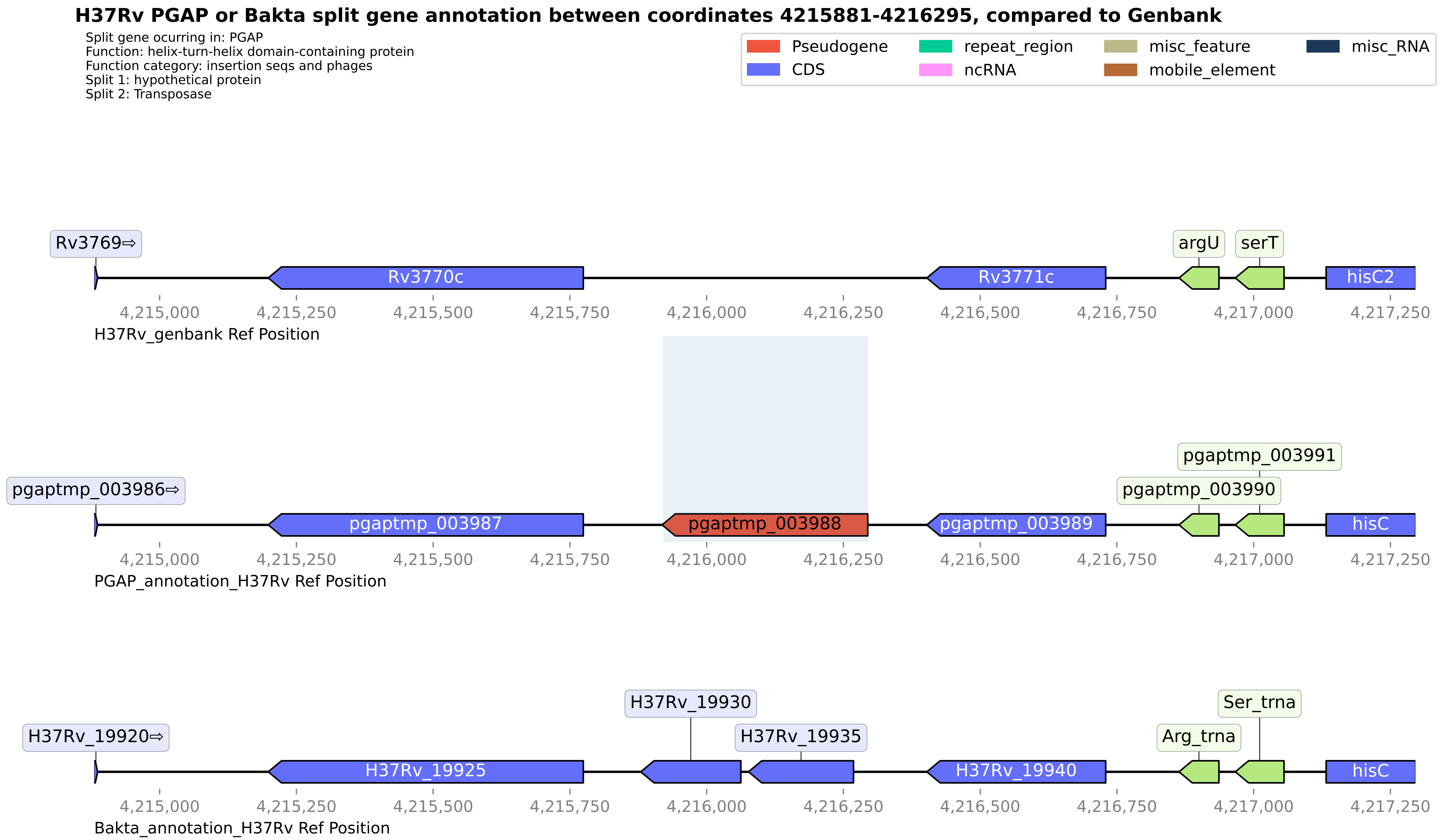

H37Rv PGAP or Bakta split gene annotation between coordinates 4215881-4216295, compared to Genbank
Split gene ocurring in: PGAP
Function: helix-turn-helix domain-containing protein
Function category: insertion seqs and phages
Split 1: hypothetical protein
Split 2: Transposase
Pseudogene
CDS
repeat_region
ncRNA
misc_feature
mobile_element
misc_RNA
Rv3769⇨
Rv3770c
Rv3771c
argU
serT
hisC2
4,215,000
4,215,250
4,215,500
4,215,750
4,216,000
4,216,250
4,216,500
4,216,750
4,217,000
4,217,250
H37Rv_genbank Ref Position
pgaptmp_003986⇨
pgaptmp_003987
pgaptmp_003988
pgaptmp_003989
pgaptmp_003990
pgaptmp_003991
hisC
PGAP_annotation_H37Rv Ref Position
H37Rv_19920⇨
H37Rv_19925
H37Rv_19930
H37Rv_19935
H37Rv_19940
Arg_trna
Ser_trna
hisC
Bakta_annotation_H37Rv Ref Position

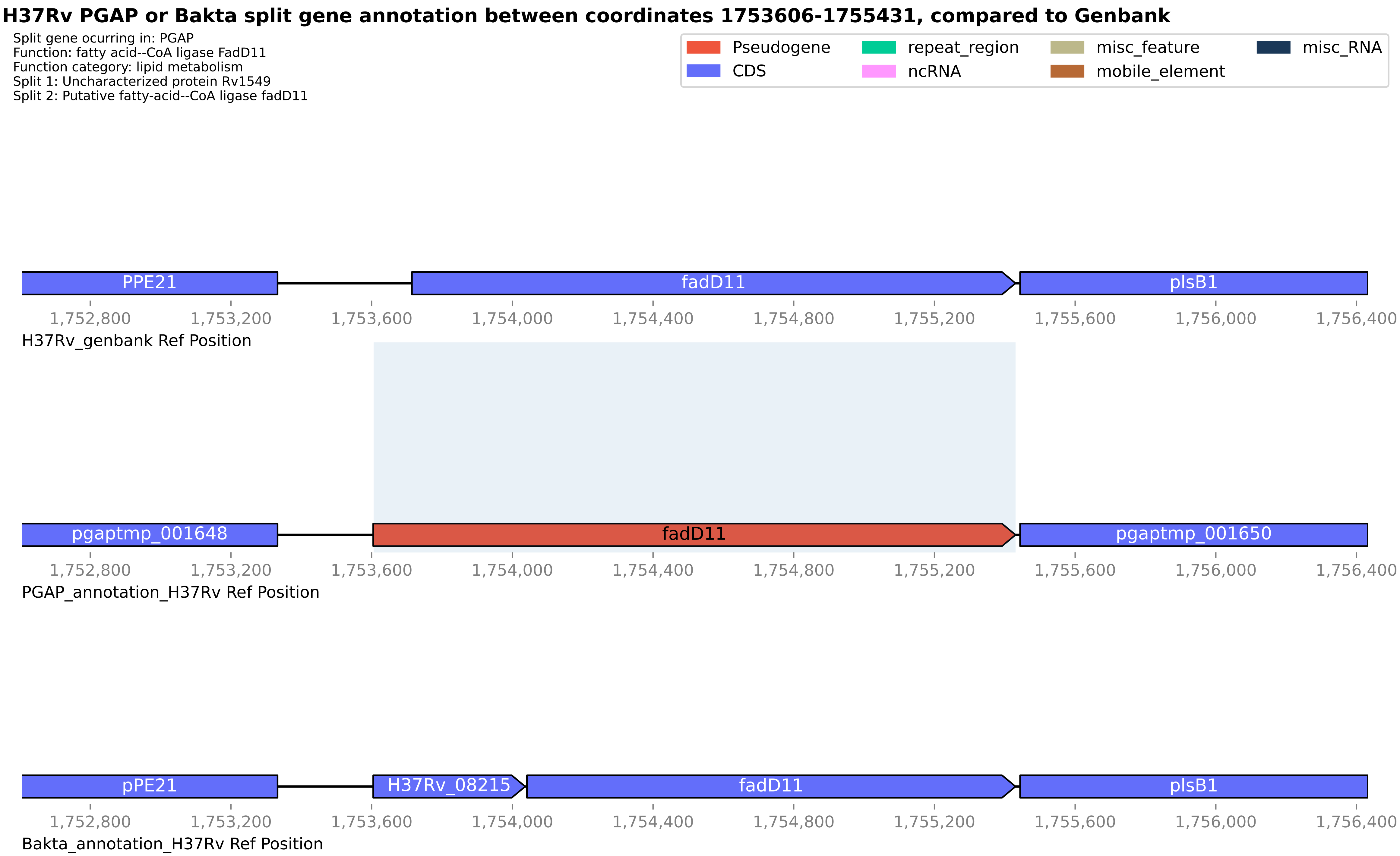
H37Rv PGAP or Bakta split gene annotation between coordinates 1753606-1755431, compared to Genbank
Split gene ocurring in: PGAP
Function: fatty acid--CoA ligase FadD11
Function category: lipid metabolism
Split 1: Uncharacterized protein Rv1549
Split 2: Putative fatty-acid--CoA ligase fadD11
Pseudogene
CDS
repeat_region
ncRNA
misc_feature
mobile_element
misc_RNA
PPE21
fadD11
plsB1
1,752,800
1,753,200
1,753,600
1,754,000
1,754,400
1,754,800
1,755,200
1,755,600
1,756,000
1,756,400
H37Rv_genbank Ref Position
pgaptmp_001648
fadD11
pgaptmp_001650
PGAP_annotation_H37Rv Ref Position
pPE21
H37Rv_08215
fadD11
plsB1
Bakta_annotation_H37Rv Ref Position

**H37Rv PGAP or Bakta split gene annotation between coordinates 2356729-2358206, compared to Genbank**

Split gene ocurring in: PGAP
Function: PE family protein
Function category: PE/PPE
Split 1: PE domain-containing protein
Split 2: PE-PGRS family protein

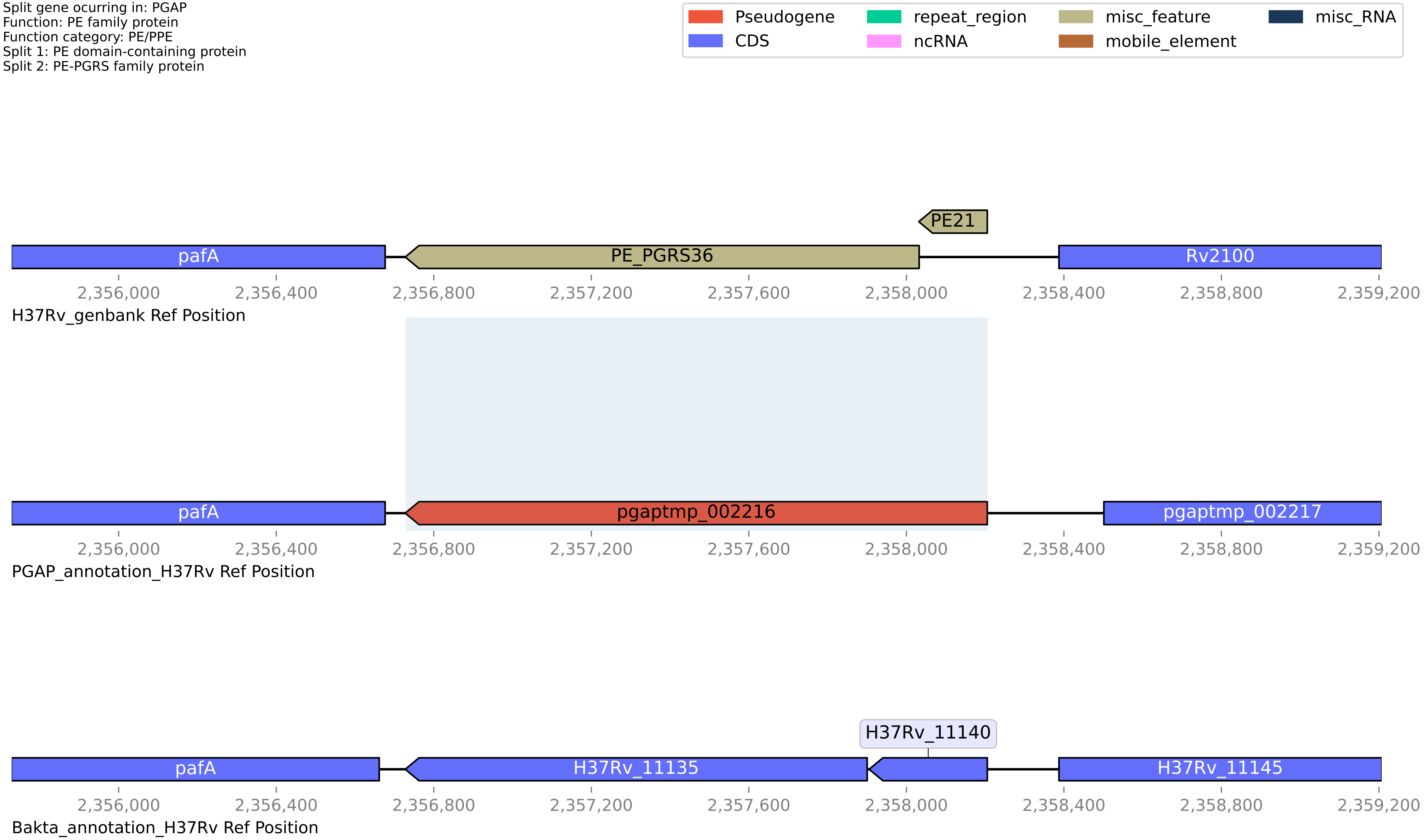

**H37Rv PGAP or Bakta split gene annotation between coordinates 3329949-3331612, compared to Genbank**

Split gene ocurring in: PGAP
Function: DAK2 domain-containing protein
Function category: conserved hypotheticals
Split 1: dihydroxyacetone kinase yloV
Split 2: DhaL domain-containing protein

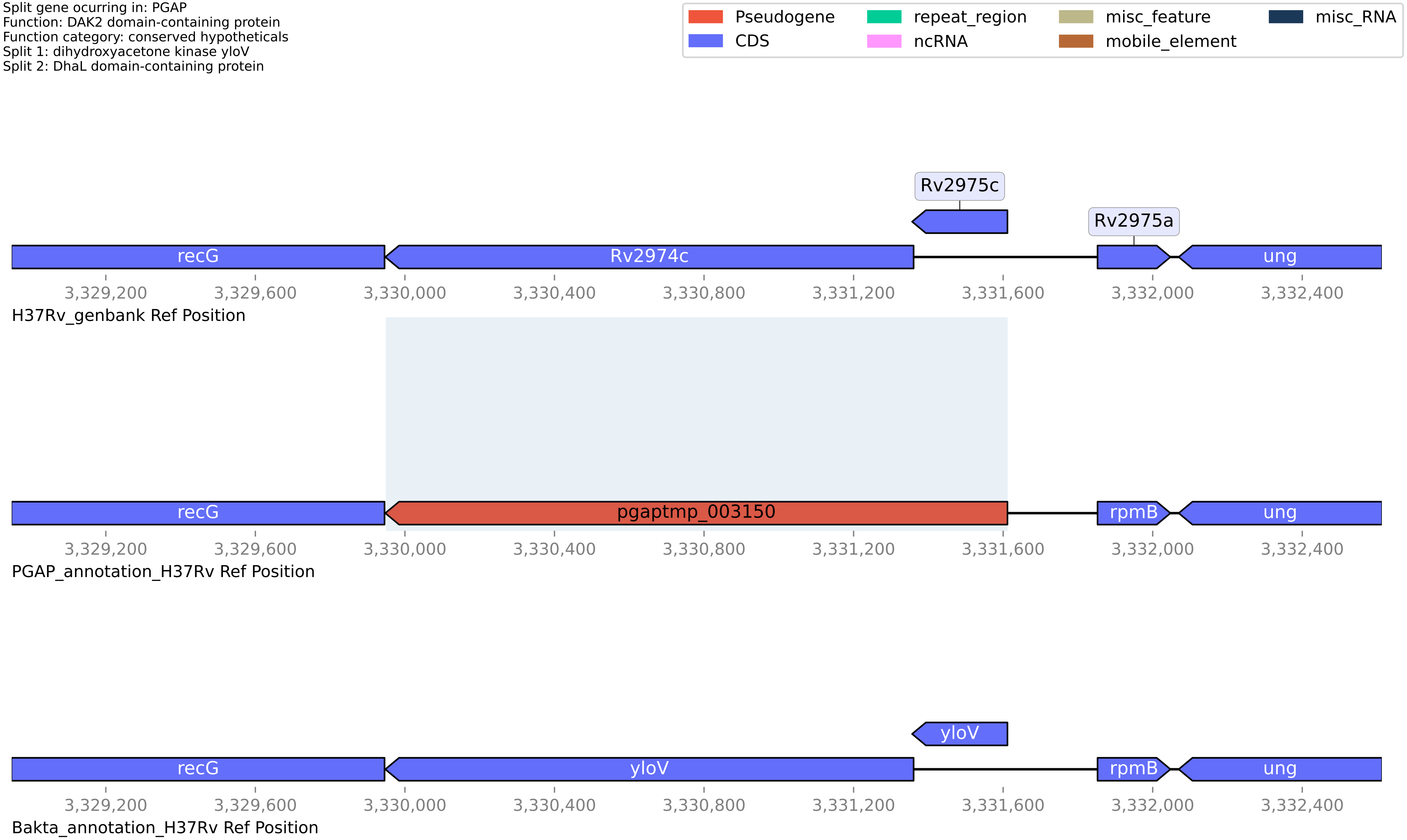

#### H37Rv PGAP or Bakta split gene annotation between coordinates 103710-105101, compared to Genbank

Split gene ocurring in: PGAP
Function: pseudogene
Function category: insertion seqs and phages
Split 1: HNHc domain-containing protein
Split 2: Putative uncharacterized protein Rv0095c

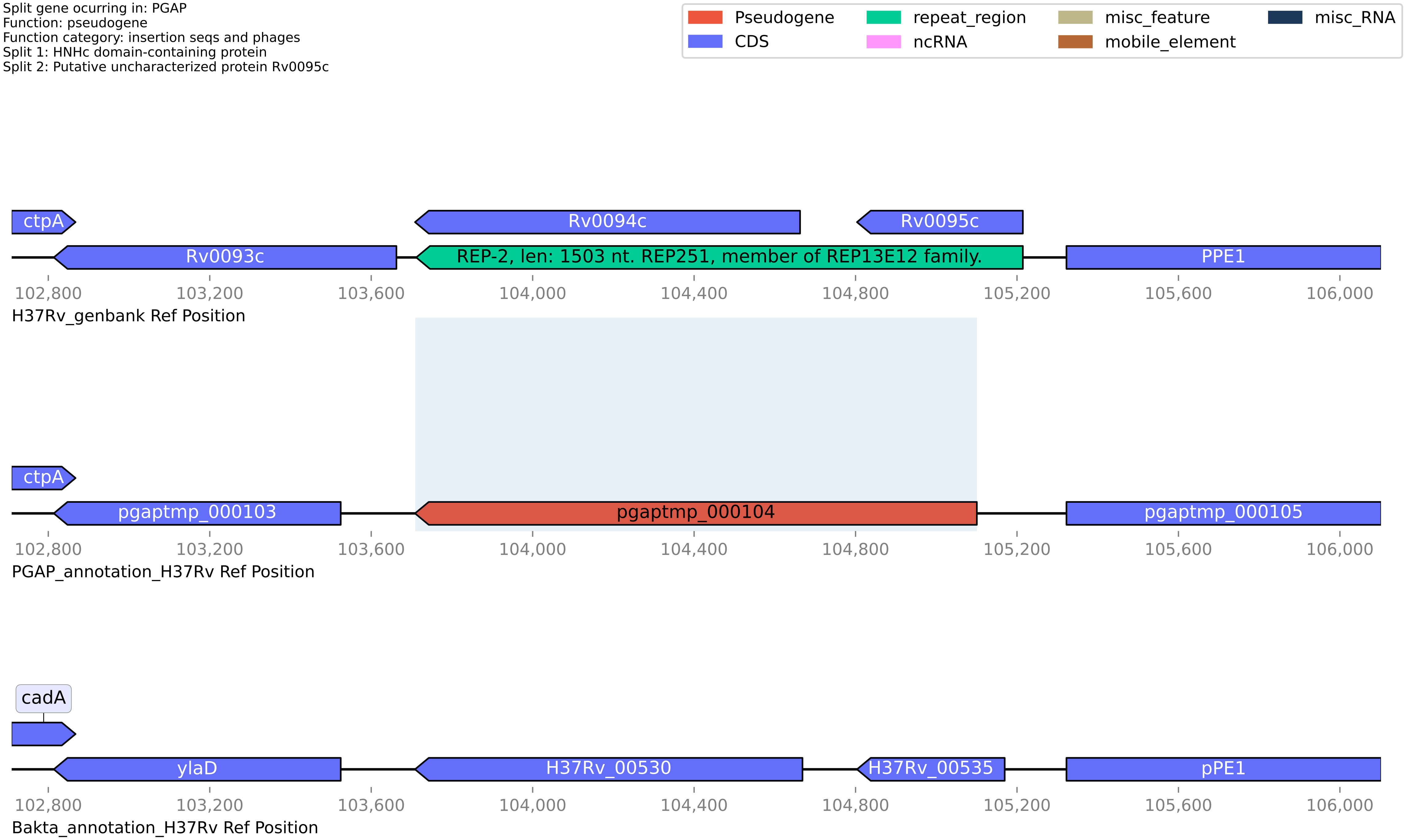

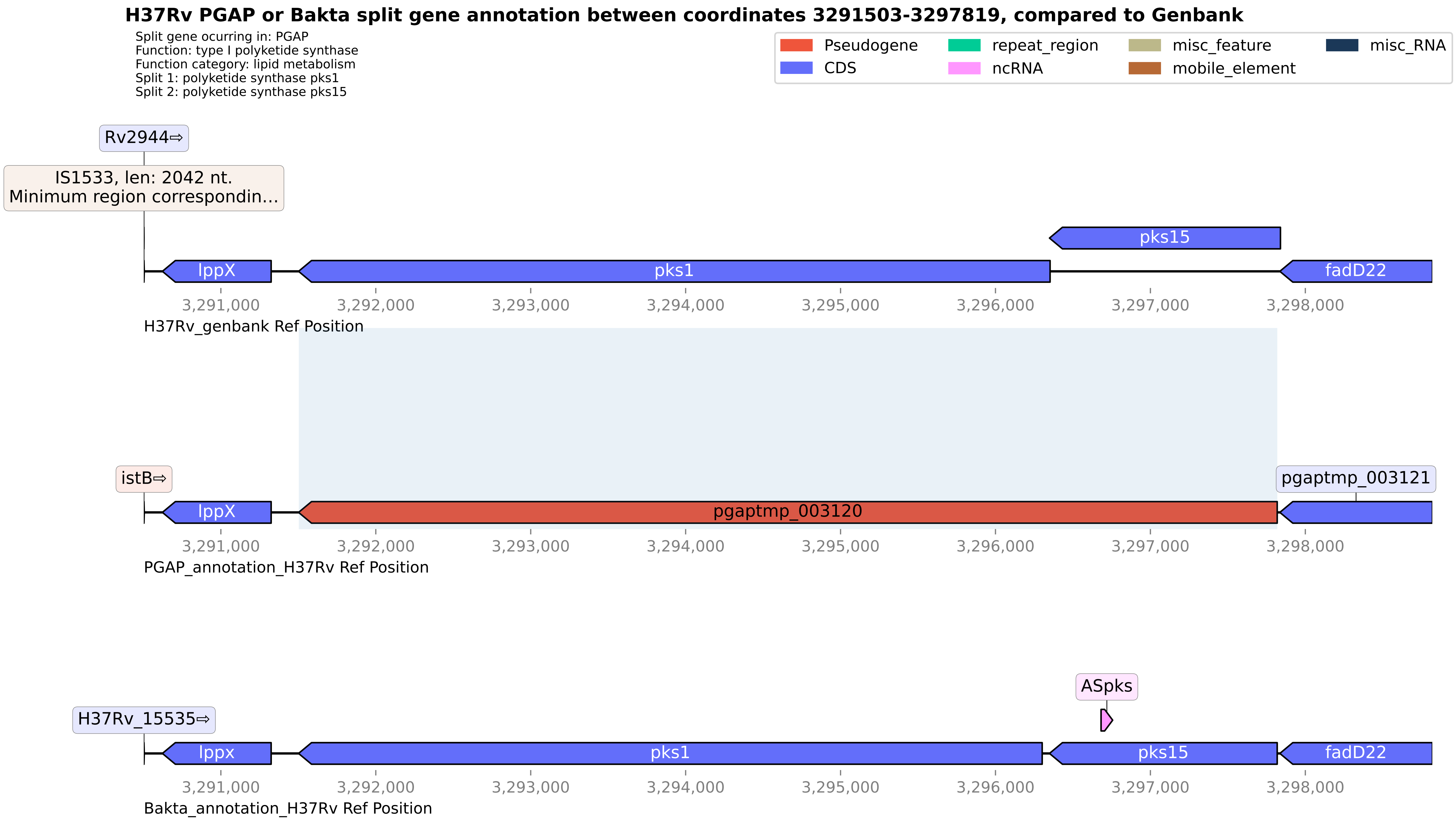
H37Rv PGAP or Bakta split gene annotation between coordinates 3291503-3297819, compared to Genbank
Split gene ocurring in: PGAP
Function: type I polyketide synthase
Function category: lipid metabolism
Split 1: polyketide synthase pks1
Split 2: polyketide synthase pks15
Pseudogene
CDS
repeat_region
ncRNA
misc_feature
mobile_element
misc_RNA
Rv2944⇨
IS1533, len: 2042 nt.
Minimum region correspondin...
pks15
lppX
pks1
fadD22
3,291,000
3,292,000
3,293,000
3,294,000
3,295,000
3,296,000
3,297,000
3,298,000
H37Rv_genbank Ref Position
istB⇨
pgaptmp_003121
lppX
pgaptmp_003120
PGAP_annotation_H37Rv Ref Position
ASpks
H37Rv_15535⇨
lppx
pks1
pks15
fadD22
Bakta_annotation_H37Rv Ref Position

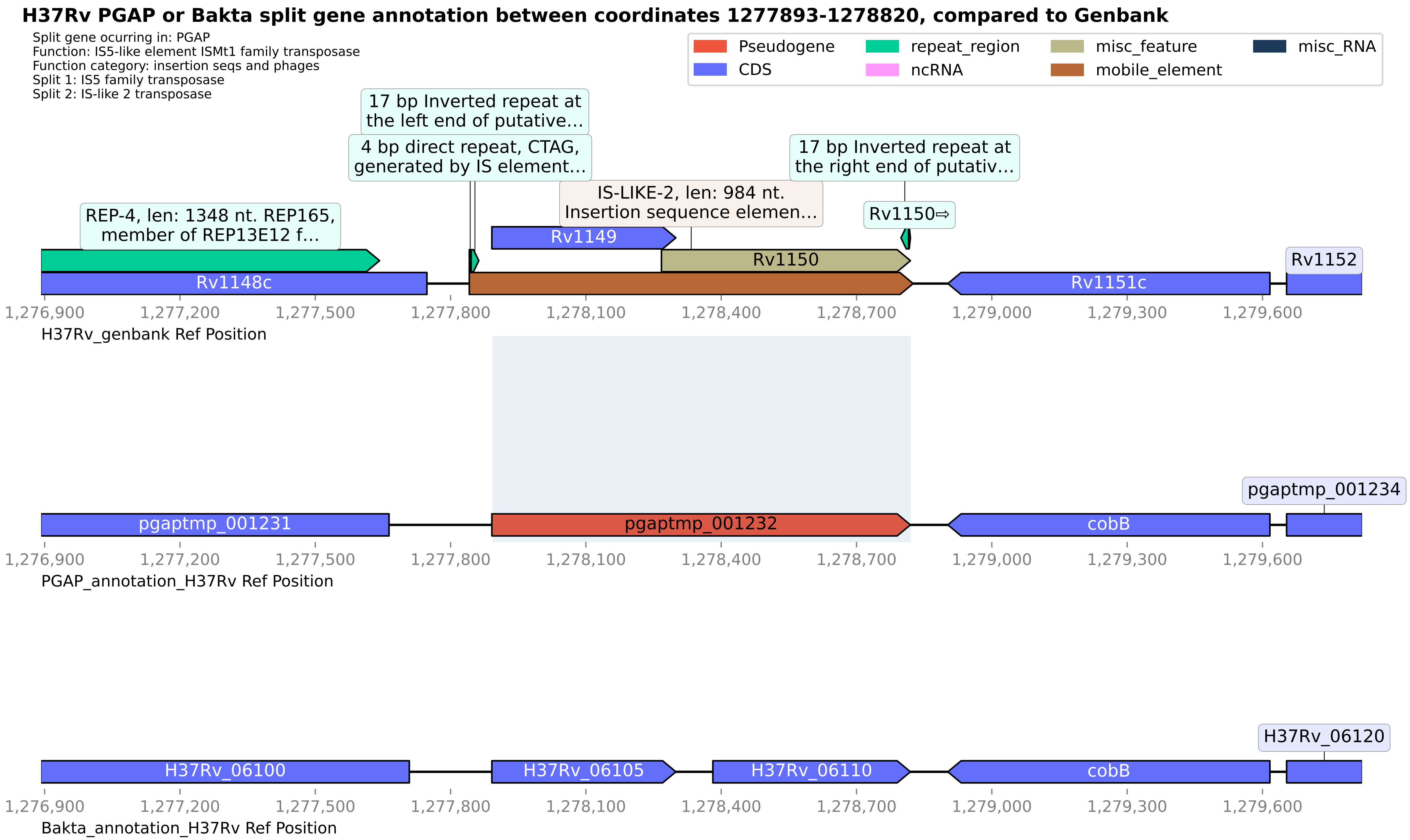
H37Rv PGAP or Bakta split gene annotation between coordinates 1277893-1278820, compared to Genbank
Split gene ocurring in: PGAP
Function: IS5-like element ISMt1 family transposase
Function category: insertion seqs and phages
Split 1: IS5 family transposase
Split 2: IS-like 2 transposase
Pseudogene
CDS
repeat_region
ncRNA
misc_feature
mobile_element
misc_RNA
17 bp Inverted repeat at the left end of putative...
4 bp direct repeat, CTAG, generated by IS element...
17 bp Inverted repeat at the right end of putativ...
IS-LIKE-2, len: 984 nt. Insertion sequence elemen...
REP-4, len: 1348 nt. REP165, member of REP13E12 f...
Rv1150⇨
Rv1149
Rv1150
Rv1148c
Rv1151c
Rv1152
1,276,900
1,277,200
1,277,500
1,277,800
1,278,100
1,278,400
1,278,700
1,279,000
1,279,300
1,279,600
H37Rv_genbank Ref Position
pgaptmp_001234
pgaptmp_001231
pgaptmp_001232
cobB
PGAP_annotation_H37Rv Ref Position
H37Rv_06120
H37Rv_06100
H37Rv_06105
H37Rv_06110
cobB
Bakta_annotation_H37Rv Ref Position

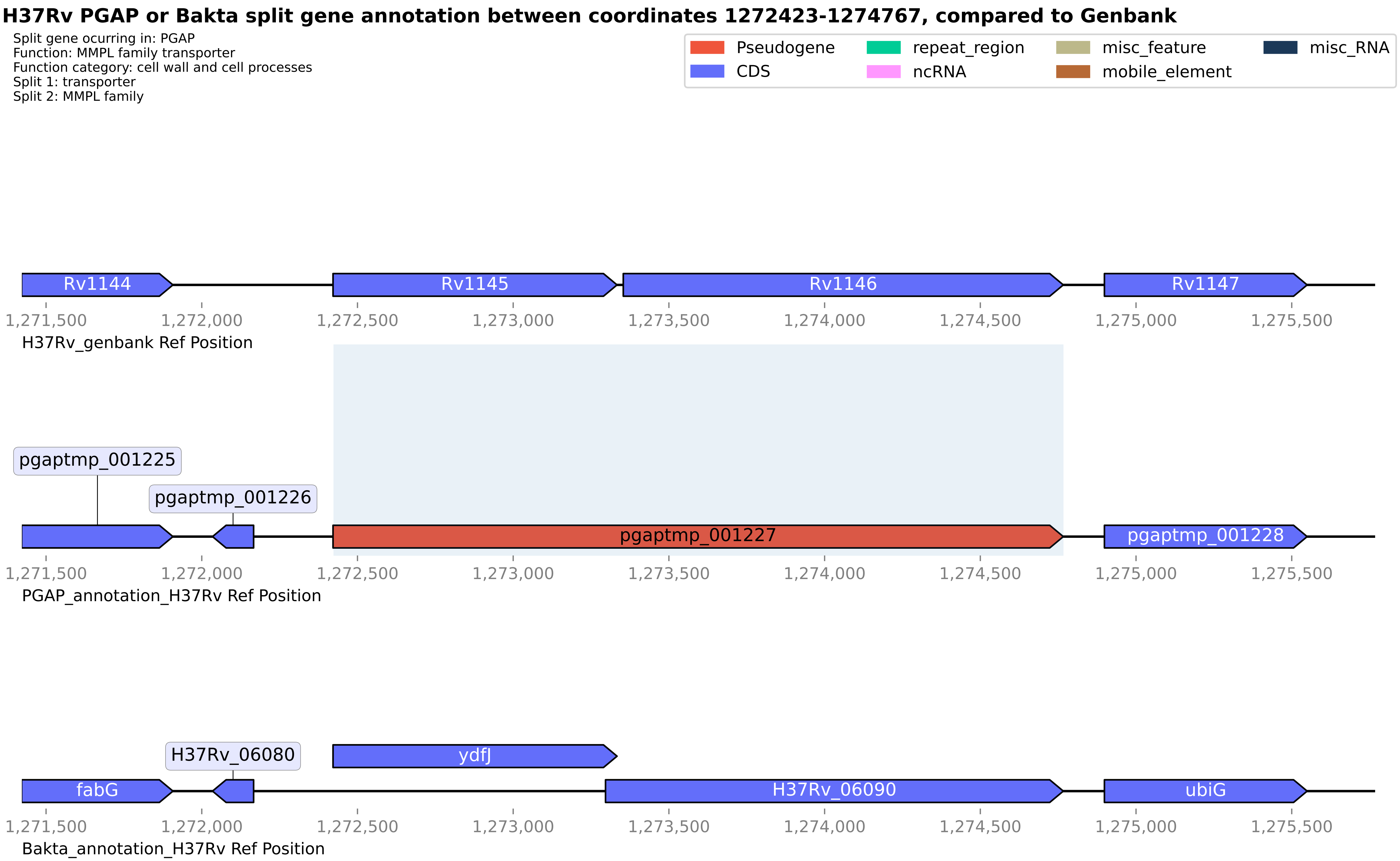
H37Rv PGAP or Bakta split gene annotation between coordinates 1272423-1274767, compared to Genbank
Split gene ocurring in: PGAP
Function: MMPL family transporter
Function category: cell wall and cell processes
Split 1: transporter
Split 2: MMPL family
Pseudogene
CDS
repeat_region
ncRNA
misc_feature
mobile_element
misc_RNA
Rv1144
Rv1145
Rv1146
Rv1147
1,271,500
1,272,000
1,272,500
1,273,000
1,273,500
1,274,000
1,274,500
1,275,000
1,275,500
H37Rv_genbank Ref Position
pgaptmp_001225
pgaptmp_001226
pgaptmp_001227
pgaptmp_001228
PGAP_annotation_H37Rv Ref Position
H37Rv_06080
ydfJ
fabG
H37Rv_06090
ubiG
Bakta_annotation_H37Rv Ref Position

**H37Rv PGAP or Bakta split gene annotation between coordinates 4189285-4190517, compared to Genbank**

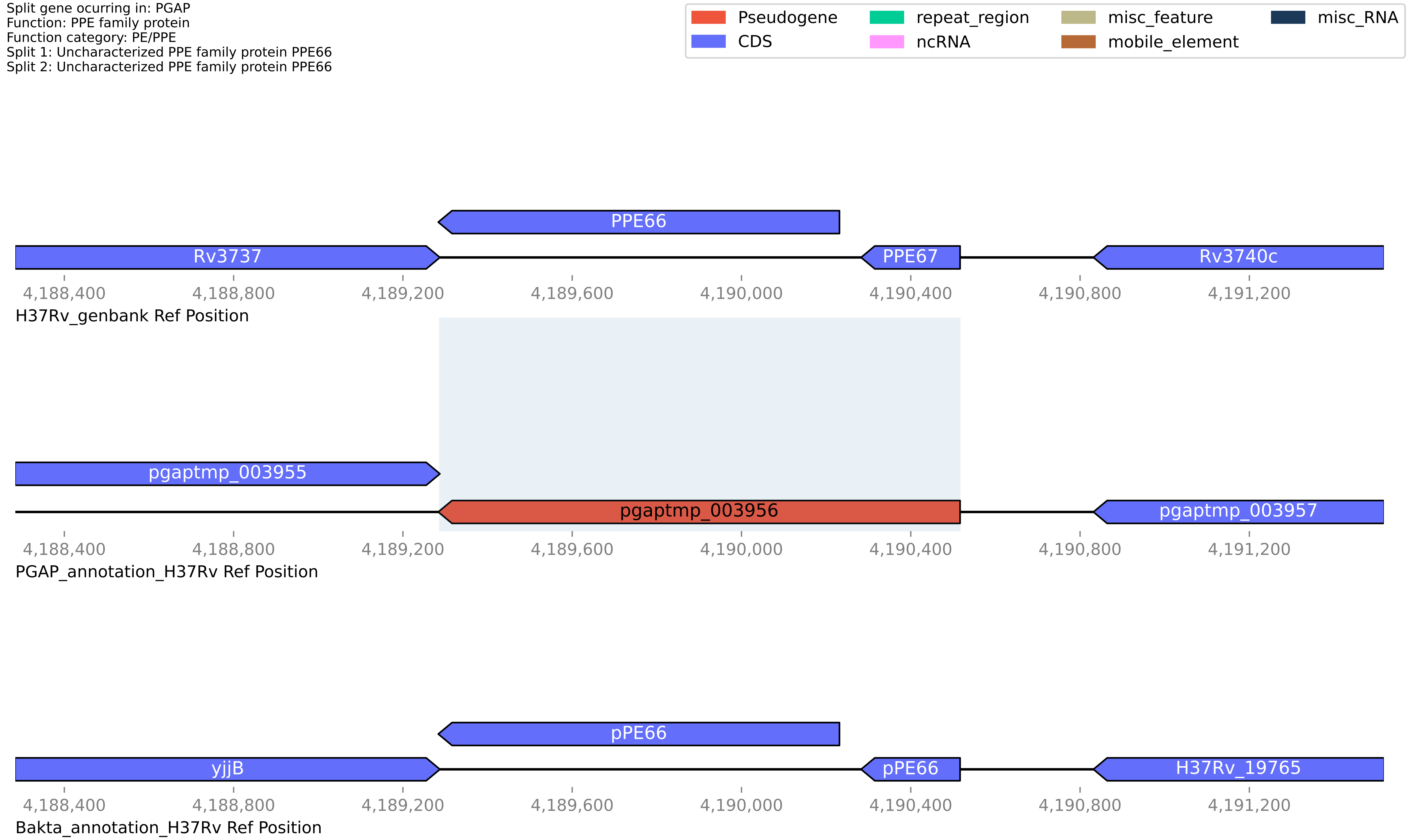

**H37Rv PGAP or Bakta split gene annotation between coordinates 1158918-1160358, compared to Genbank**

Split gene ocurring in: PGAP
Function: ISNCY family transposase
Function category: insertion seqs and phages
Split 1: Transposase
Split 2: Putative transposase

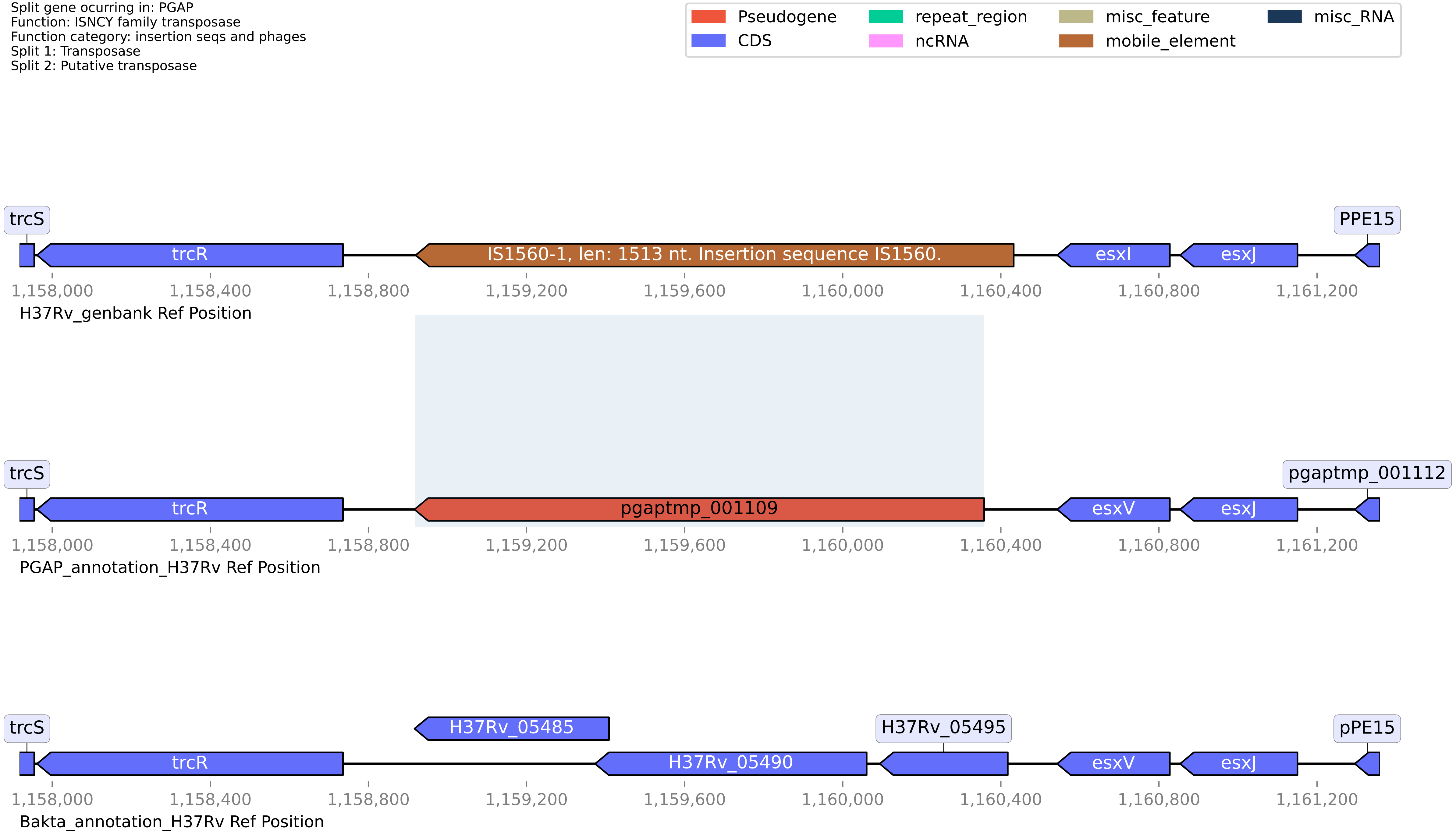

#### H37Rv PGAP or Bakta split gene annotation between coordinates 1231301-1232837, compared to Genbank

Split gene ocurring in: PGAP
Function: carboxylesterase/lipase family protein
Function category: intermediary metabolism and respiration
Split 1: Para-nitrobenzyl esterase
Split 2: Para-nitrobenzyl esterase

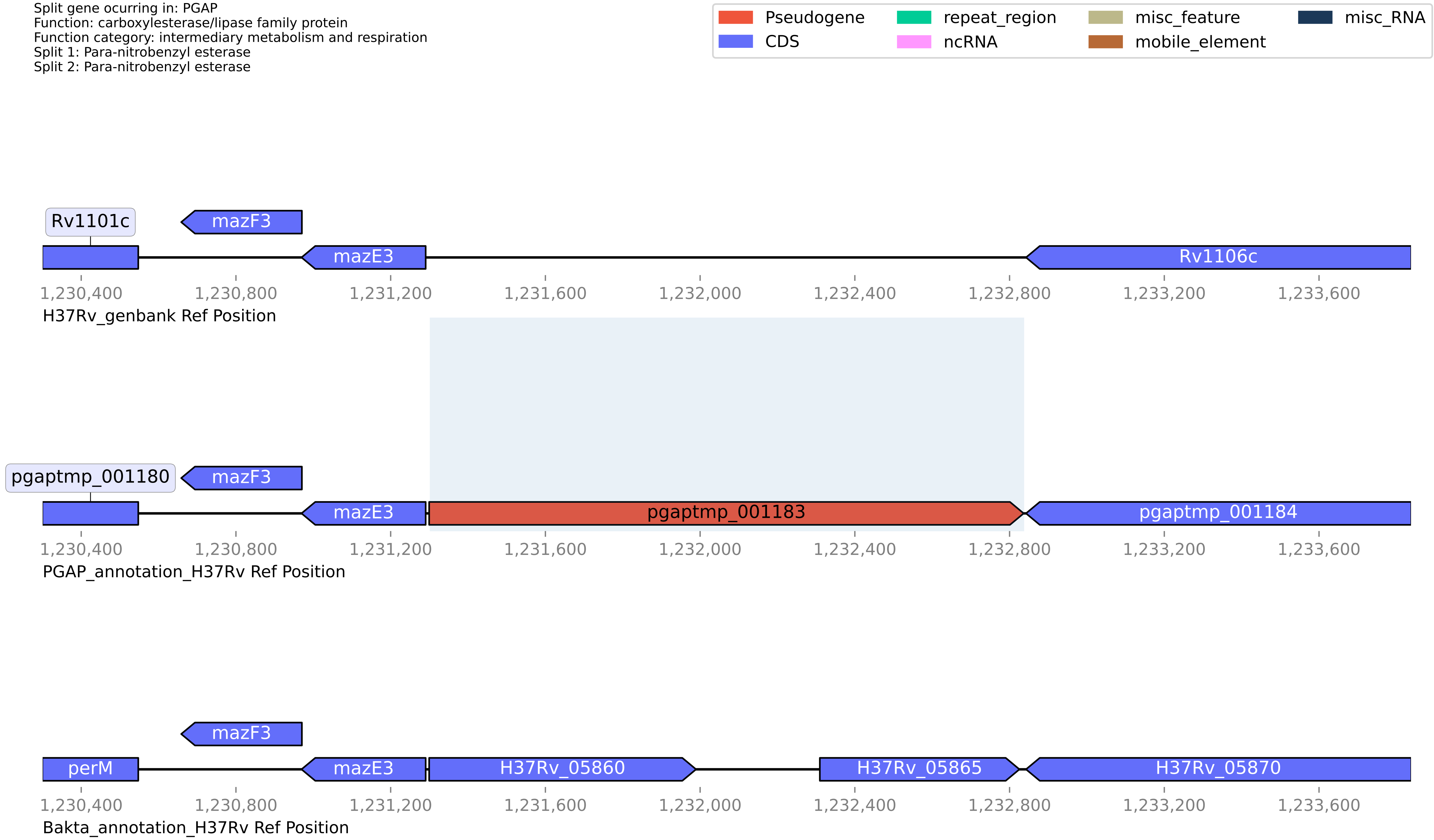

**H37Rv PGAP or Bakta split gene annotation between coordinates 1164572-1165499, compared to Genbank**

Split gene ocurring in: PGAP
Function: IS5-like element ISMt1 family transposase
Function category: insertion seqs and phages
Split 1: IS-like 2 transposase
Split 2: IS5 family transposase

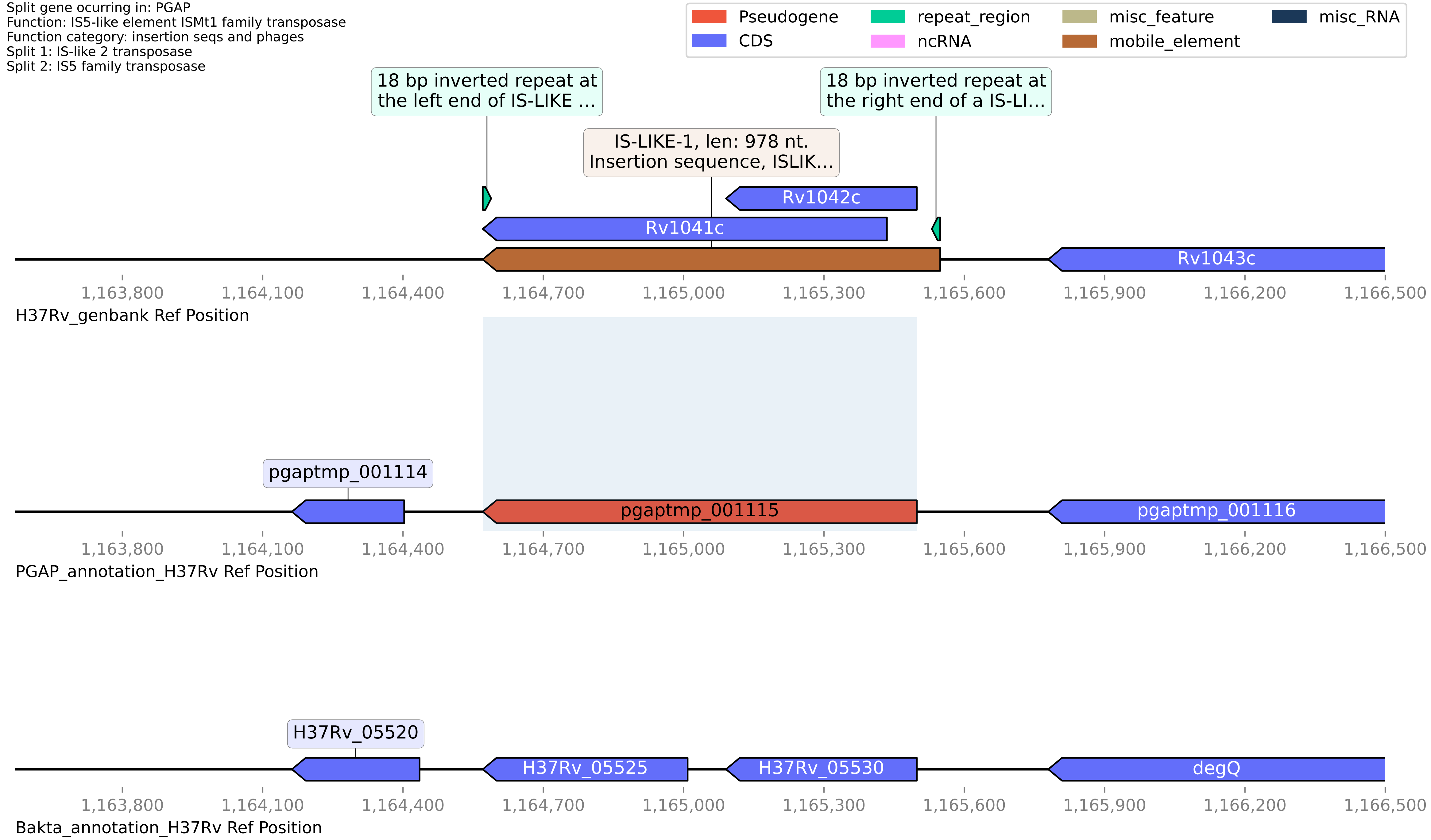

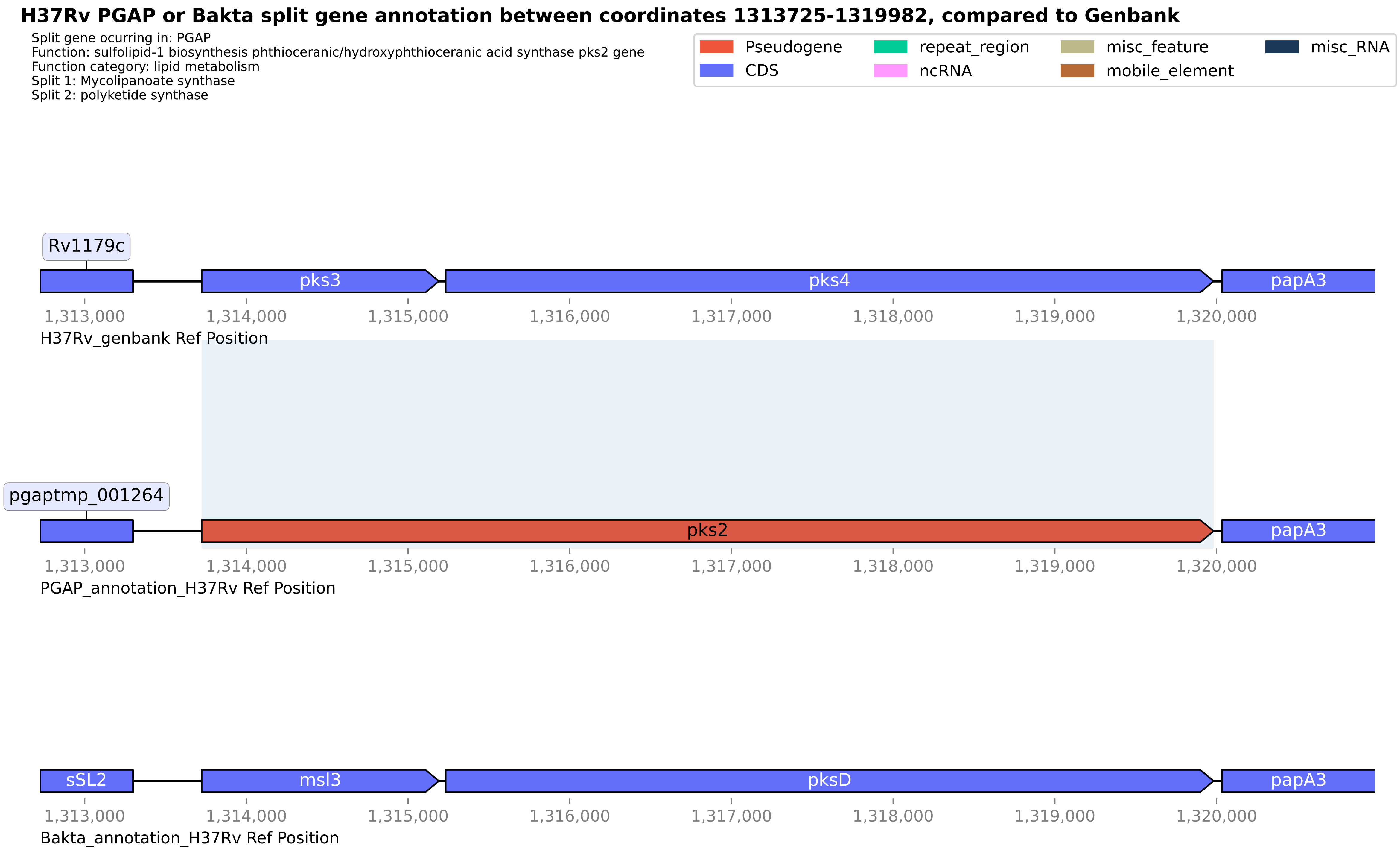

H37Rv PGAP or Bakta split gene annotation between coordinates 1313725-1319982, compared to Genbank
Split gene ocurring in: PGAP
Function: sulfolipid-1 biosynthesis phthioceranic/hydroxyphthioceranic acid synthase pks2 gene
Function category: lipid metabolism
Split 1: Mycolipanoate synthase
Split 2: polyketide synthase
Pseudogene
CDS
repeat_region
ncRNA
misc_feature
mobile_element
misc_RNA
Rv1179c
pks3
pks4
papA3
1,313,000
1,314,000
1,315,000
1,316,000
1,317,000
1,318,000
1,319,000
1,320,000
H37Rv_genbank Ref Position
pgaptmp_001264
pks2
papA3
PGAP_annotation_H37Rv Ref Position
sSL2
msl3
pksD
papA3
Bakta_annotation_H37Rv Ref Position

**H37Rv PGAP or Bakta split gene annotation between coordinates 2534042-2535552, compared to Genbank**

Split gene ocurring in: PGAP
Function: apolipoprotein N-acyltransferase Int
Function category: lipid metabolism
Split 1: CN hydrolase domain-containing protein
Split 2: apolipoprotein N-acyltransferase Int

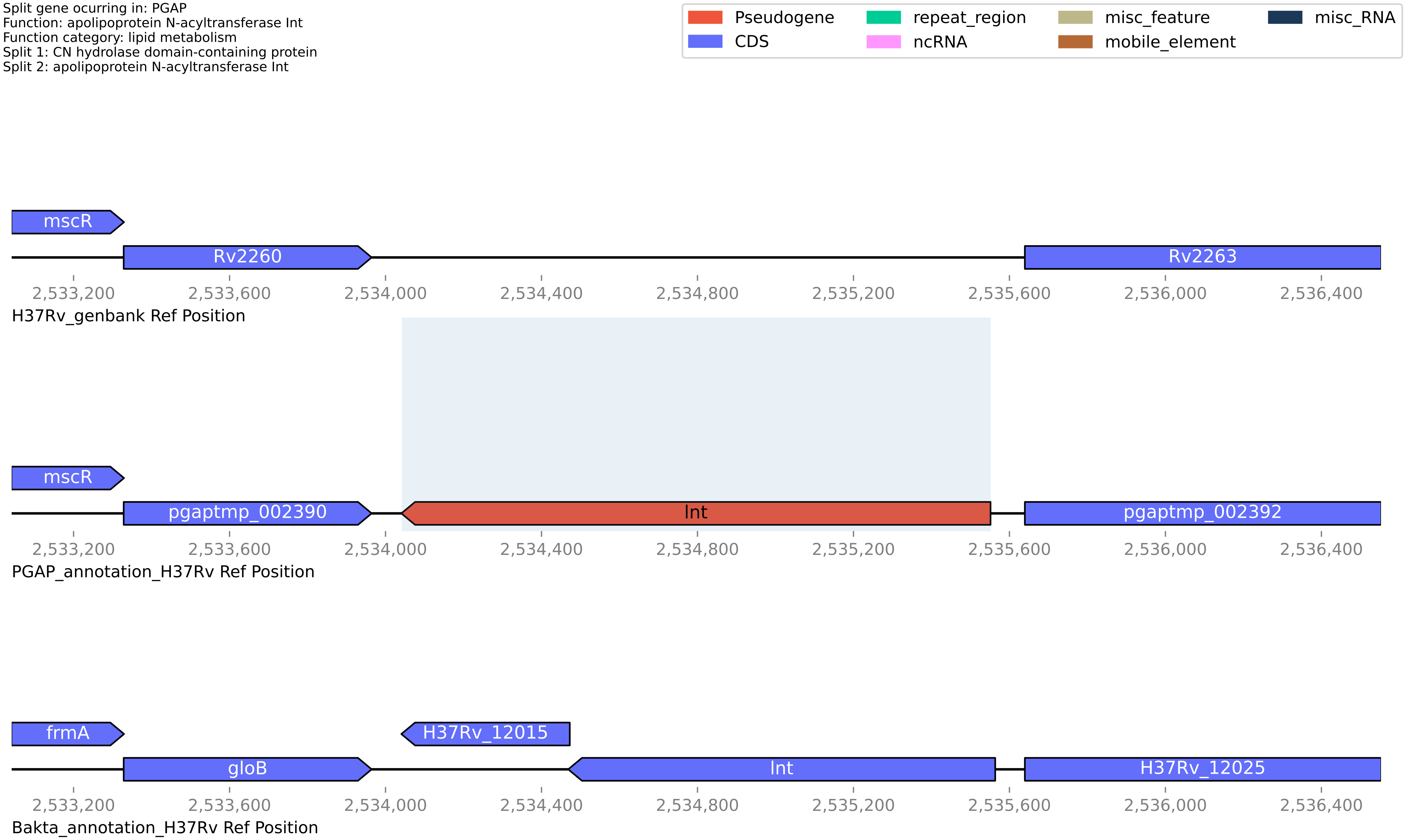

#### H37Rv PGAP or Bakta split gene annotation between coordinates 1242864-1243634, compared to Genbank

Split gene ocurring in: PGAP
Function: adenylate/guanylate cyclase domain-containing protein
Function category: conserved hypotheticals
Split 1: Conserved protein of uncharacterized function (Part2)
Split 2: Guanylate cyclase domain-containing protein

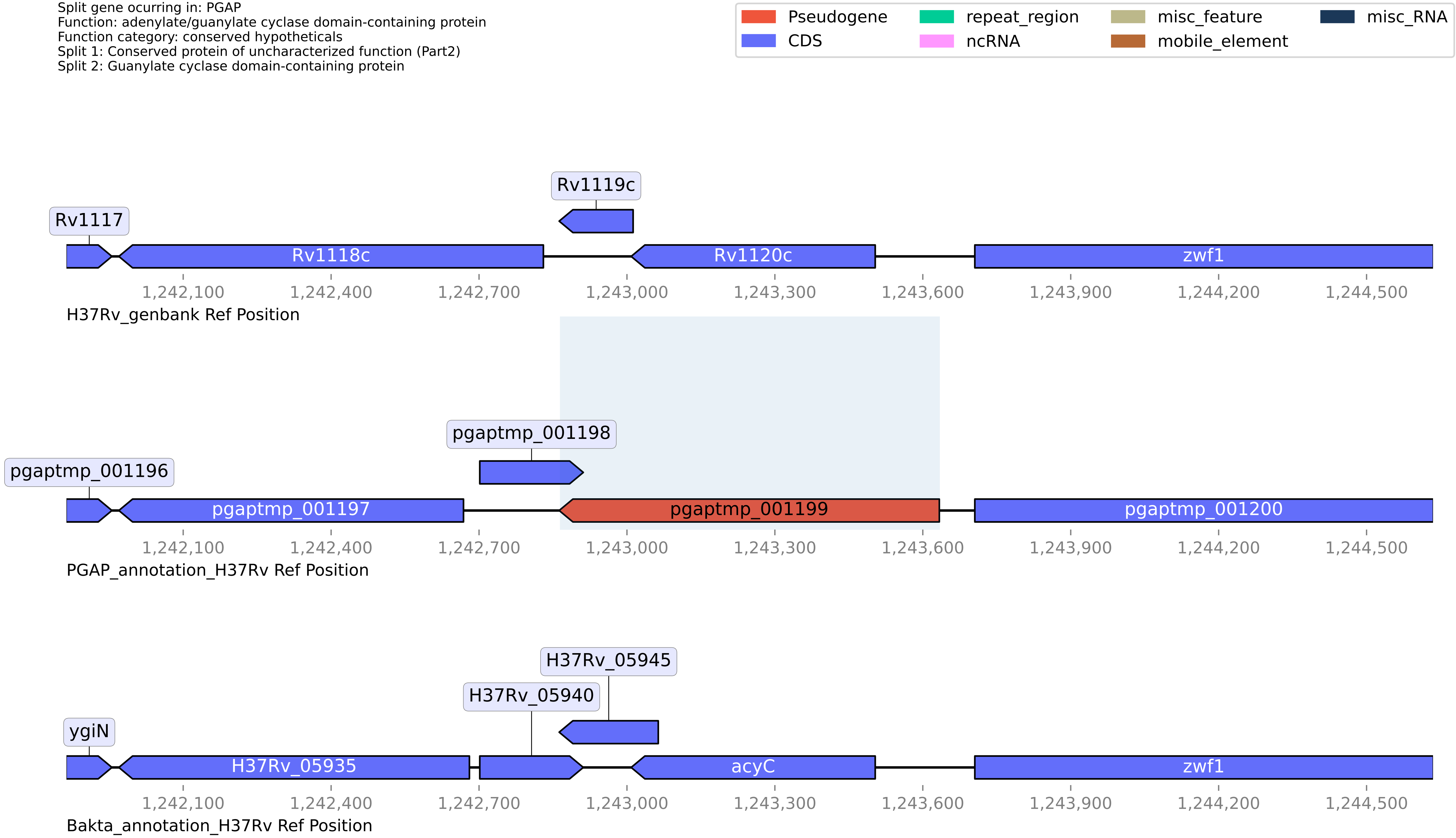

#### H37Rv PGAP or Bakta split gene annotation between coordinates 103710-105101, compared to Genbank

Split gene ocurring in: PGAP
Function: pseudogene
Function category: insertion seqs and phages
Split 1: HNHc domain-containing protein
Split 2: Putative uncharacterized protein Rv0095c

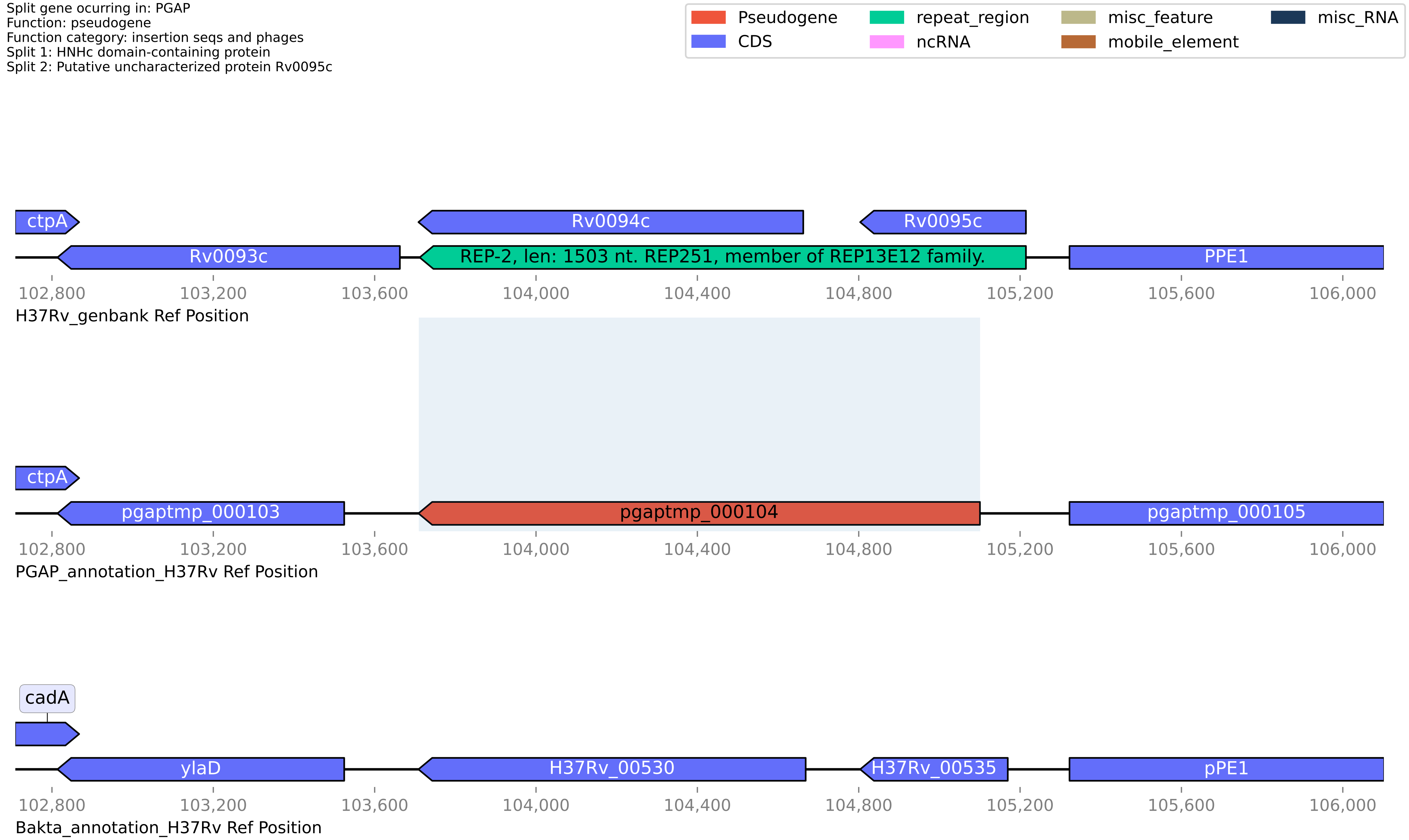

#### H37Rv PGAP or Bakta split gene annotation between coordinates 1158918-1160358, compared to Genbank

Split gene ocurring in: PGAP
Function: ISNCY family transposase
Function category: insertion seqs and phages
Split 1: Transposase
Split 2: Putative transposase

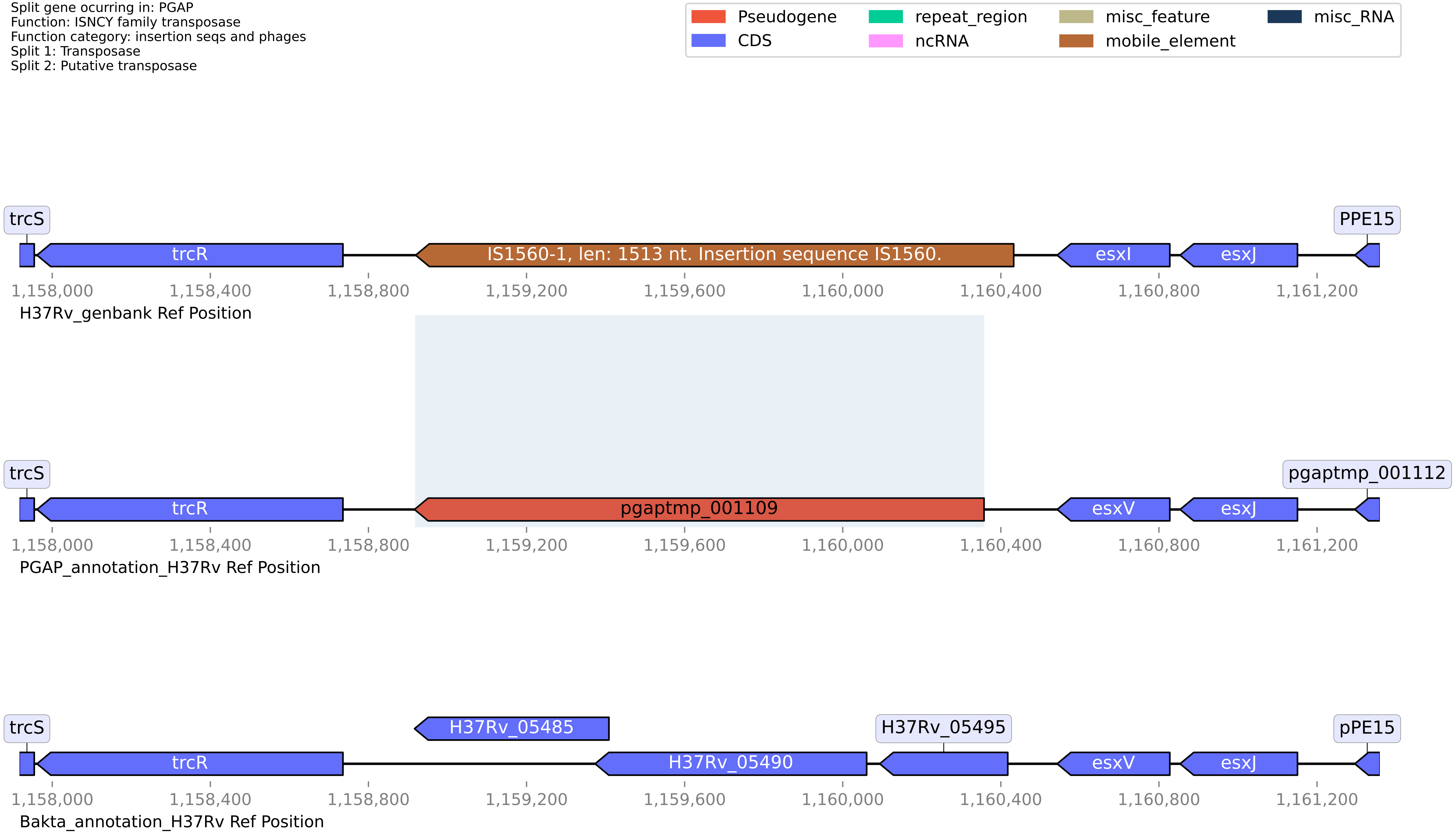

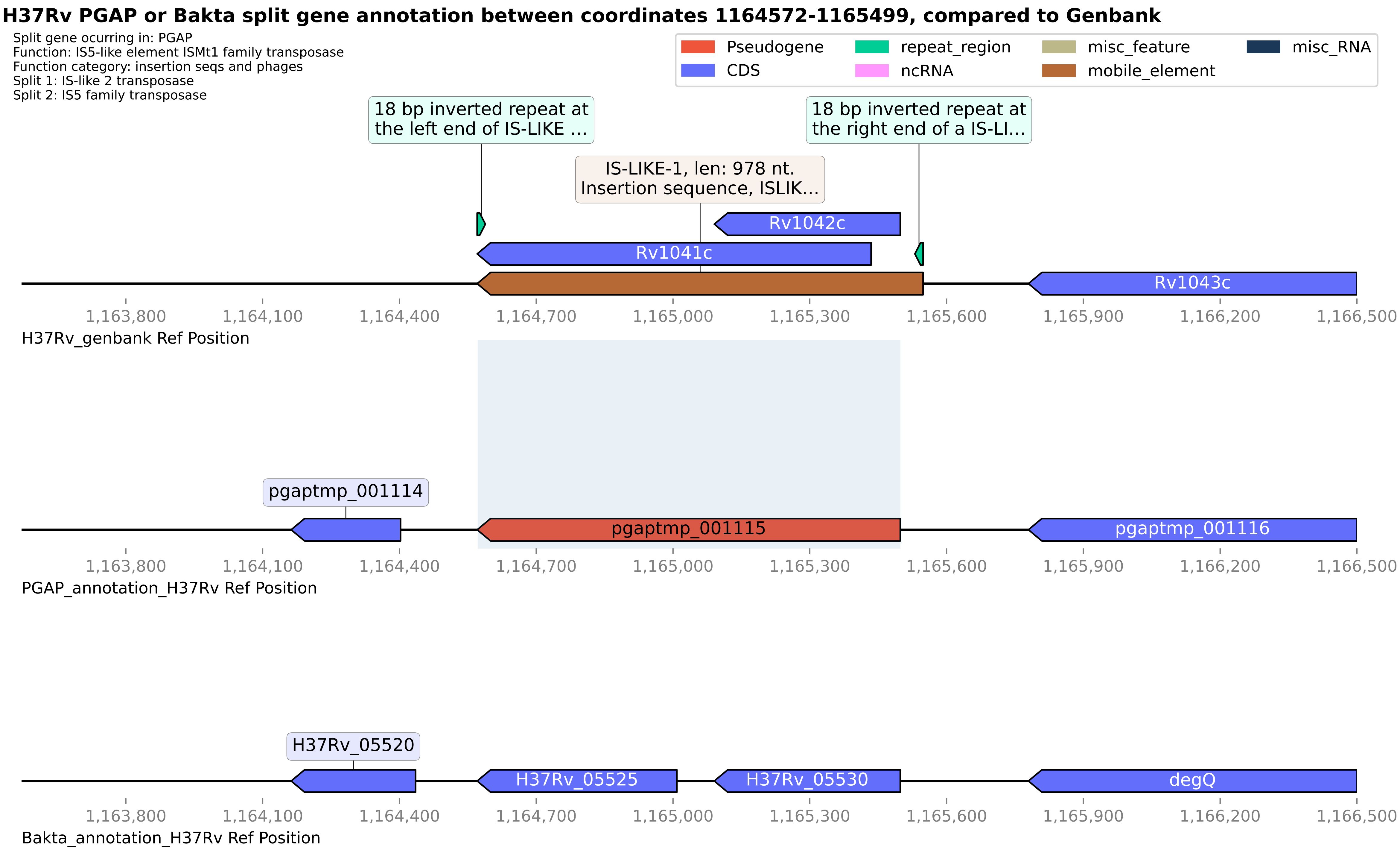
H37Rv PGAP or Bakta split gene annotation between coordinates 1164572-1165499, compared to Genbank
Split gene ocurring in: PGAP
Function: IS5-like element ISMt1 family transposase
Function category: insertion seqs and phages
Split 1: IS-like 2 transposase
Split 2: IS5 family transposase
Pseudogene
CDS
repeat_region
ncRNA
misc_feature
mobile_element
misc_RNA
18 bp inverted repeat at the left end of IS-LIKE ...
18 bp inverted repeat at the right end of a IS-LI...
IS-LIKE-1, len: 978 nt. Insertion sequence, ISLIK...
Rv1042c
Rv1041c
Rv1043c
1,163,800
1,164,100
1,164,400
1,164,700
1,165,000
1,165,300
1,165,600
1,165,900
1,166,200
1,166,500
H37Rv_genbank Ref Position
pgaptmp_001114
pgaptmp_001115
pgaptmp_001116
PGAP_annotation_H37Rv Ref Position
H37Rv_05520
H37Rv_05525
H37Rv_05530
degQ
Bakta_annotation_H37Rv Ref Position

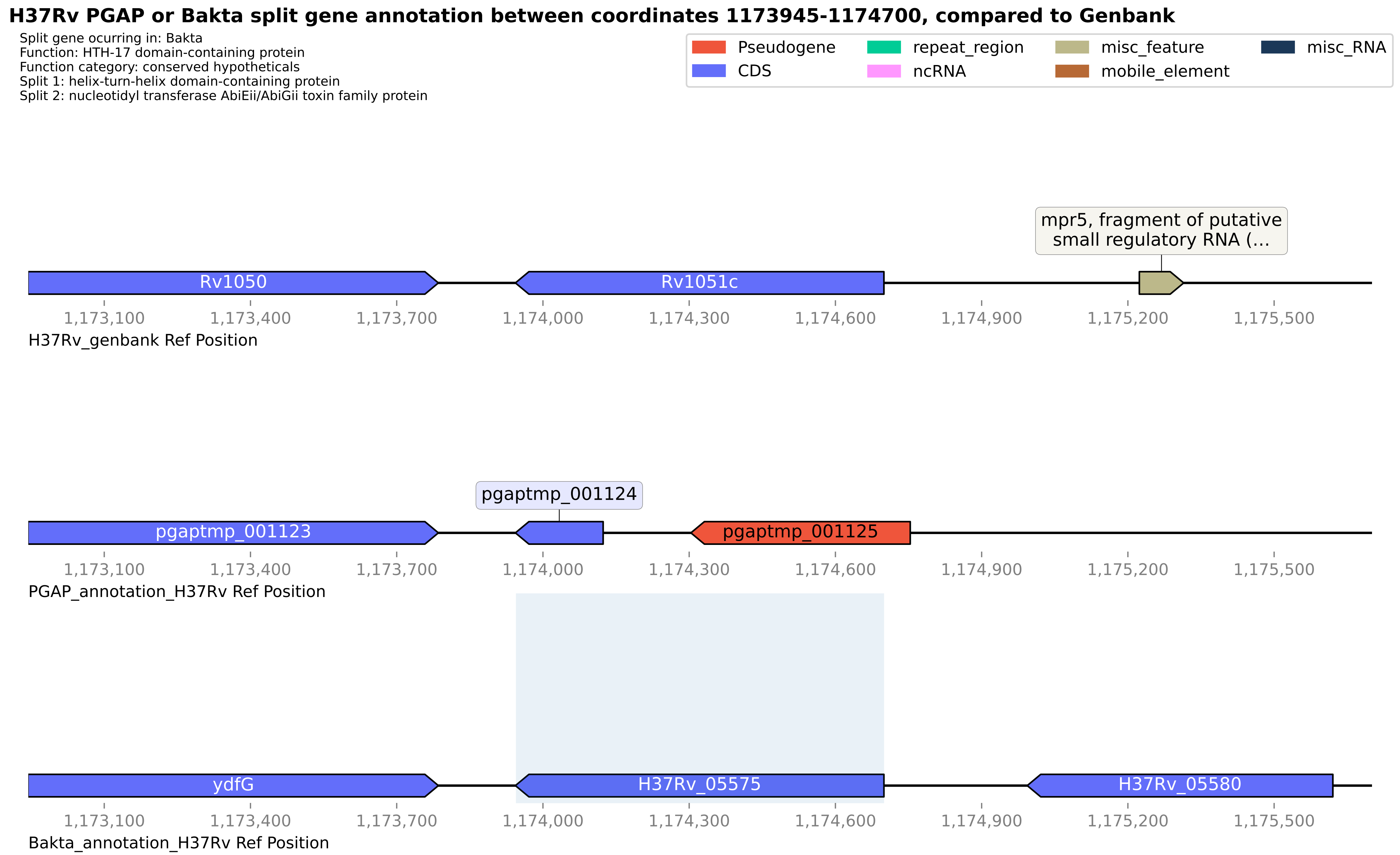
H37Rv PGAP or Bakta split gene annotation between coordinates 1173945-1174700, compared to Genbank
Split gene ocurring in: Bakta
Function: HTH-17 domain-containing protein
Function category: conserved hypotheticals
Split 1: helix-turn-helix domain-containing protein
Split 2: nucleotidyl transferase AbiEii/AbiGii toxin family protein
Pseudogene
CDS
repeat_region
ncRNA
misc_feature
mobile_element
misc_RNA
mpr5, fragment of putative small regulatory RNA (...
Rv1050
Rv1051c
1,173,100
1,173,400
1,173,700
1,174,000
1,174,300
1,174,600
1,174,900
1,175,200
1,175,500
H37Rv_genbank Ref Position
pgaptmp_001124
pgaptmp_001123
pgaptmp_001125
PGAP_annotation_H37Rv Ref Position
ydfG
H37Rv_05575
H37Rv_05580
Bakta_annotation_H37Rv Ref Position

#### H37Rv PGAP or Bakta split gene annotation between coordinates 1231301-1232837, compared to Genbank

Split gene ocurring in: PGAP
Function: carboxylesterase/lipase family protein
Function category: intermediary metabolism and respiration
Split 1: Para-nitrobenzyl esterase
Split 2: Para-nitrobenzyl esterase

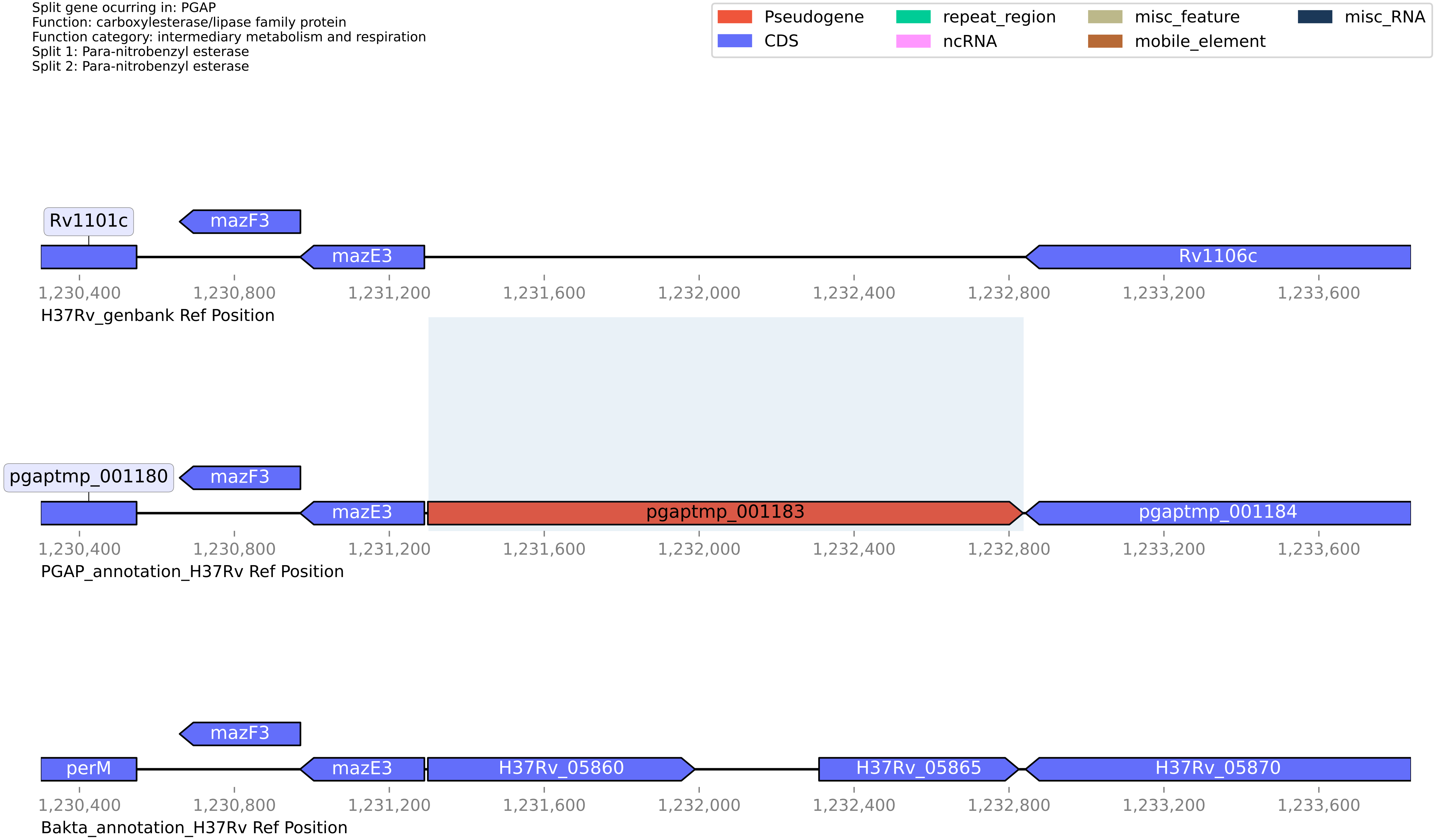

**H37Rv PGAP or Bakta split gene annotation between coordinates 1242864-1243634, compared to Genbank**

Split gene ocurring in: PGAP
Function: adenylate/guanylate cyclase domain-containing protein
Function category: conserved hypotheticals
Split 1: Conserved protein of uncharacterized function (Part2)
Split 2: Guanylate cyclase domain-containing protein

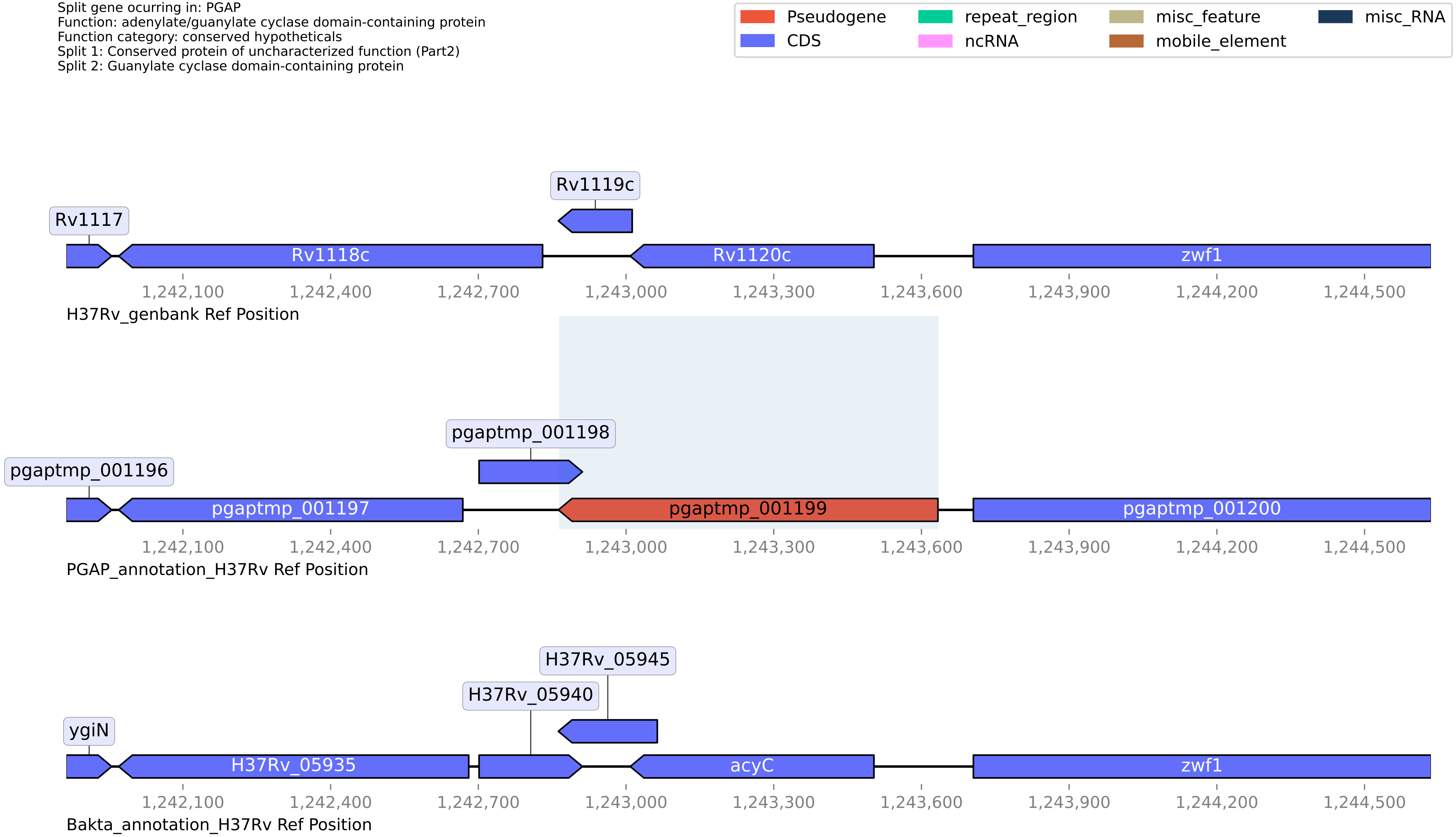

#### H37Rv PGAP or Bakta split gene annotation between coordinates 1272423-1274767, compared to Genbank

Split gene ocurring in: PGAP
Function: MMPL family transporter
Function category: cell wall and cell processes
Split 1: transporter
Split 2: MMPL family

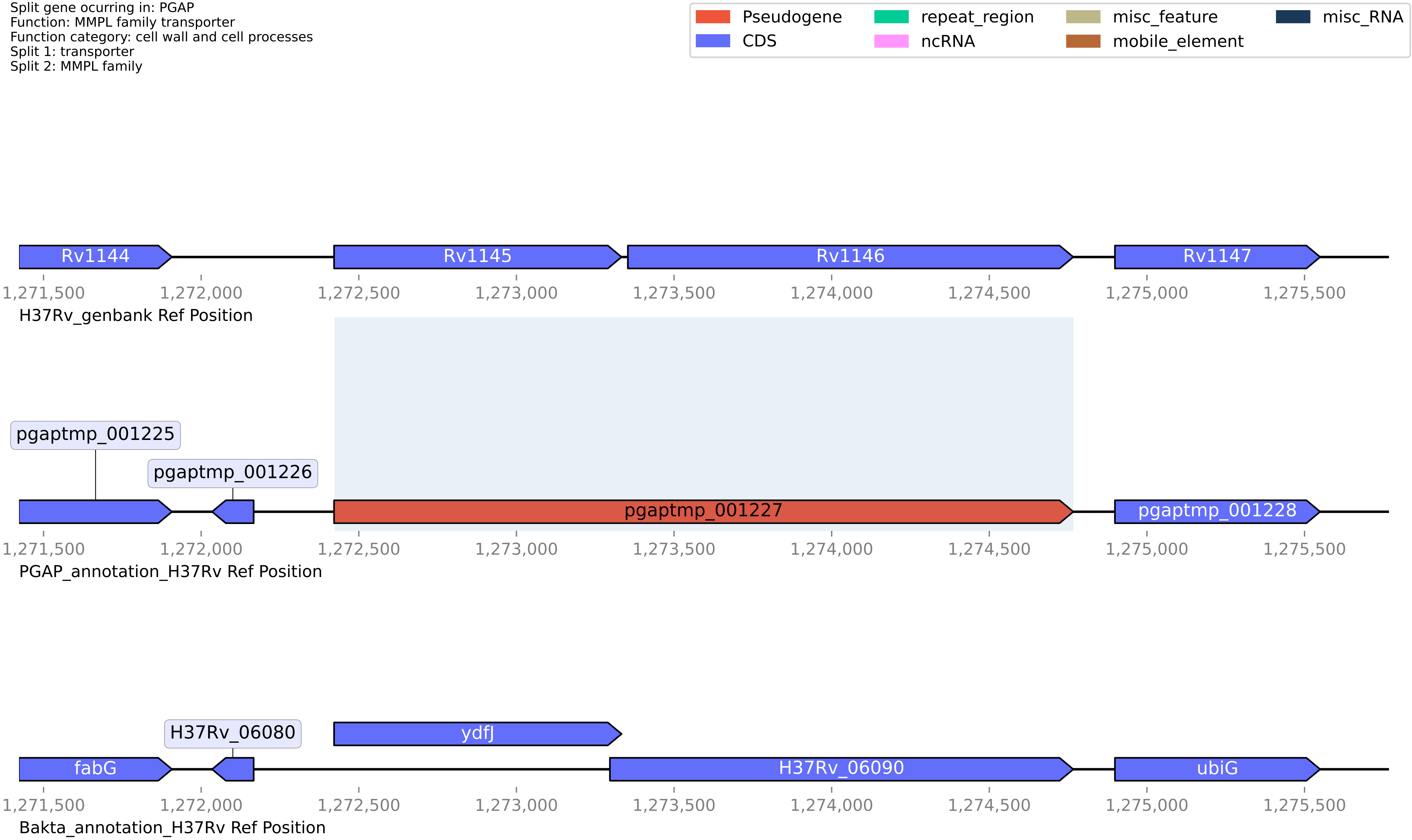

#### H37Rv PGAP or Bakta split gene annotation between coordinates 1277893-1278820, compared to Genbank

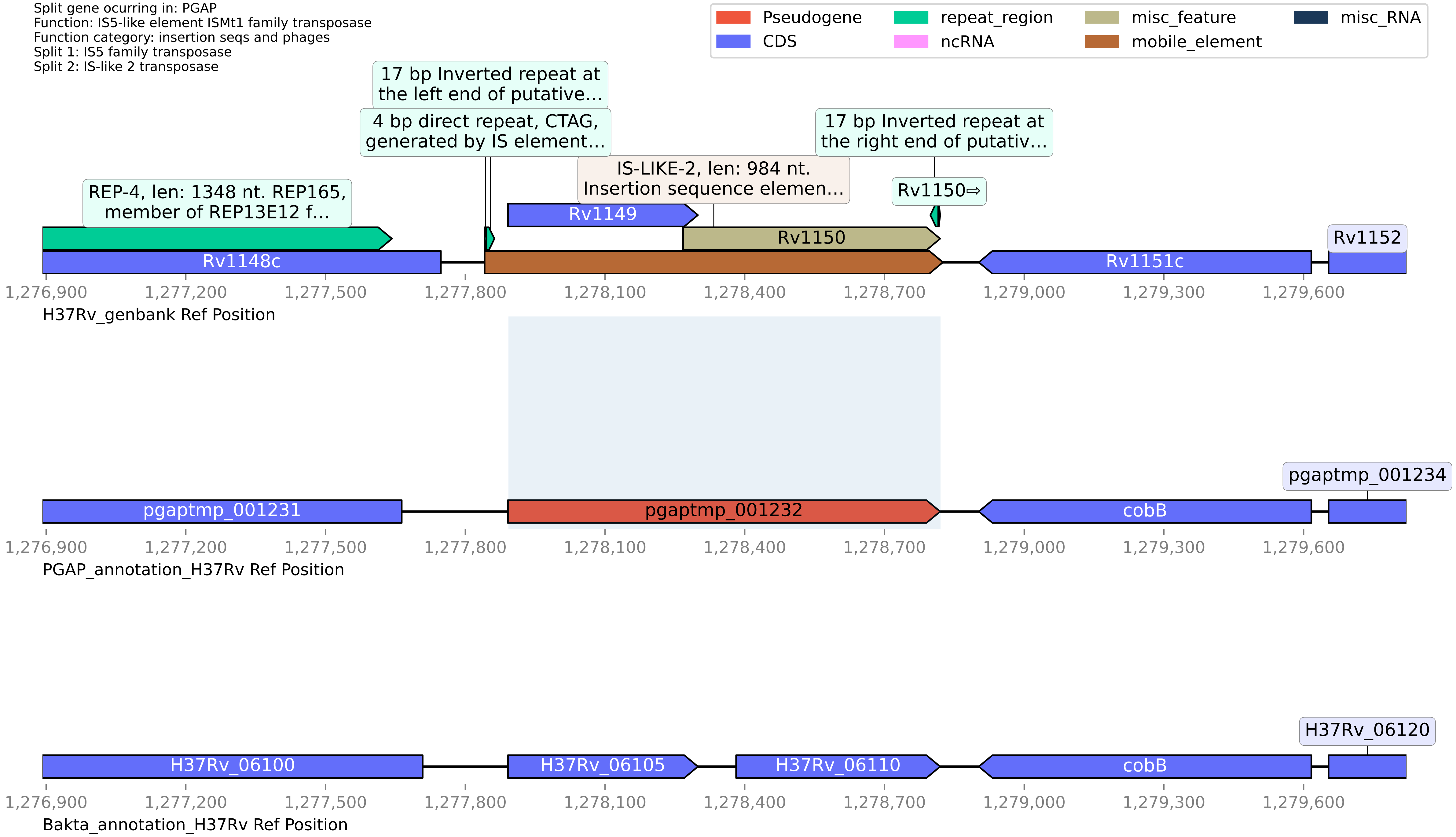

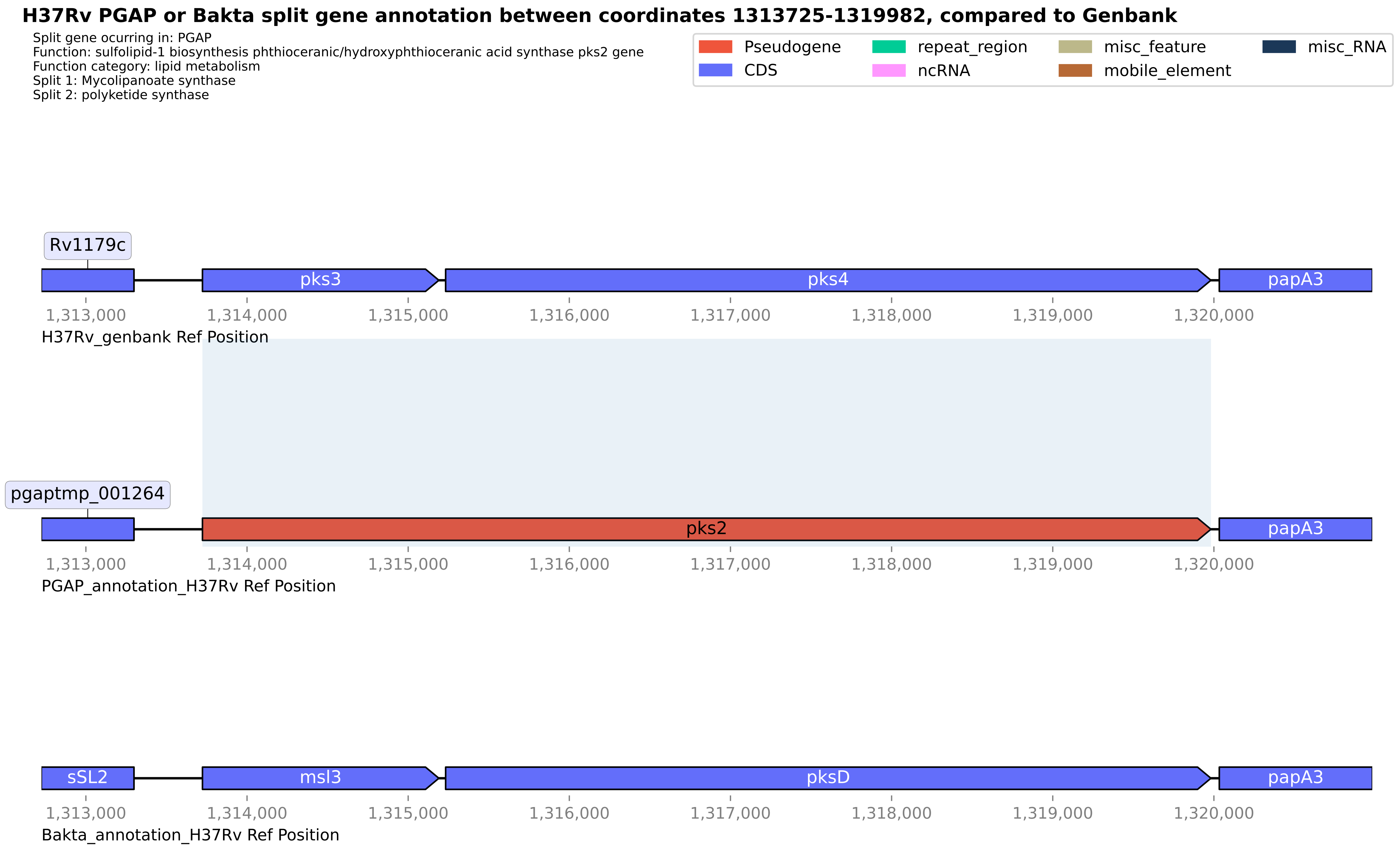
H37Rv PGAP or Bakta split gene annotation between coordinates 1313725-1319982, compared to Genbank
Split gene ocurring in: PGAP
Function: sulfolipid-1 biosynthesis phthioceranic/hydroxyphthioceranic acid synthase pks2 gene
Function category: lipid metabolism
Split 1: Mycolipanoate synthase
Split 2: polyketide synthase
Pseudogene
CDS
repeat_region
ncRNA
misc_feature
mobile_element
misc_RNA
Rv1179c
pks3
pks4
papA3
1,313,000
1,314,000
1,315,000
1,316,000
1,317,000
1,318,000
1,319,000
1,320,000
H37Rv_genbank Ref Position
pgaptmp_001264
pks2
papA3
PGAP_annotation_H37Rv Ref Position
sSL2
msl3
pksD
papA3
Bakta_annotation_H37Rv Ref Position

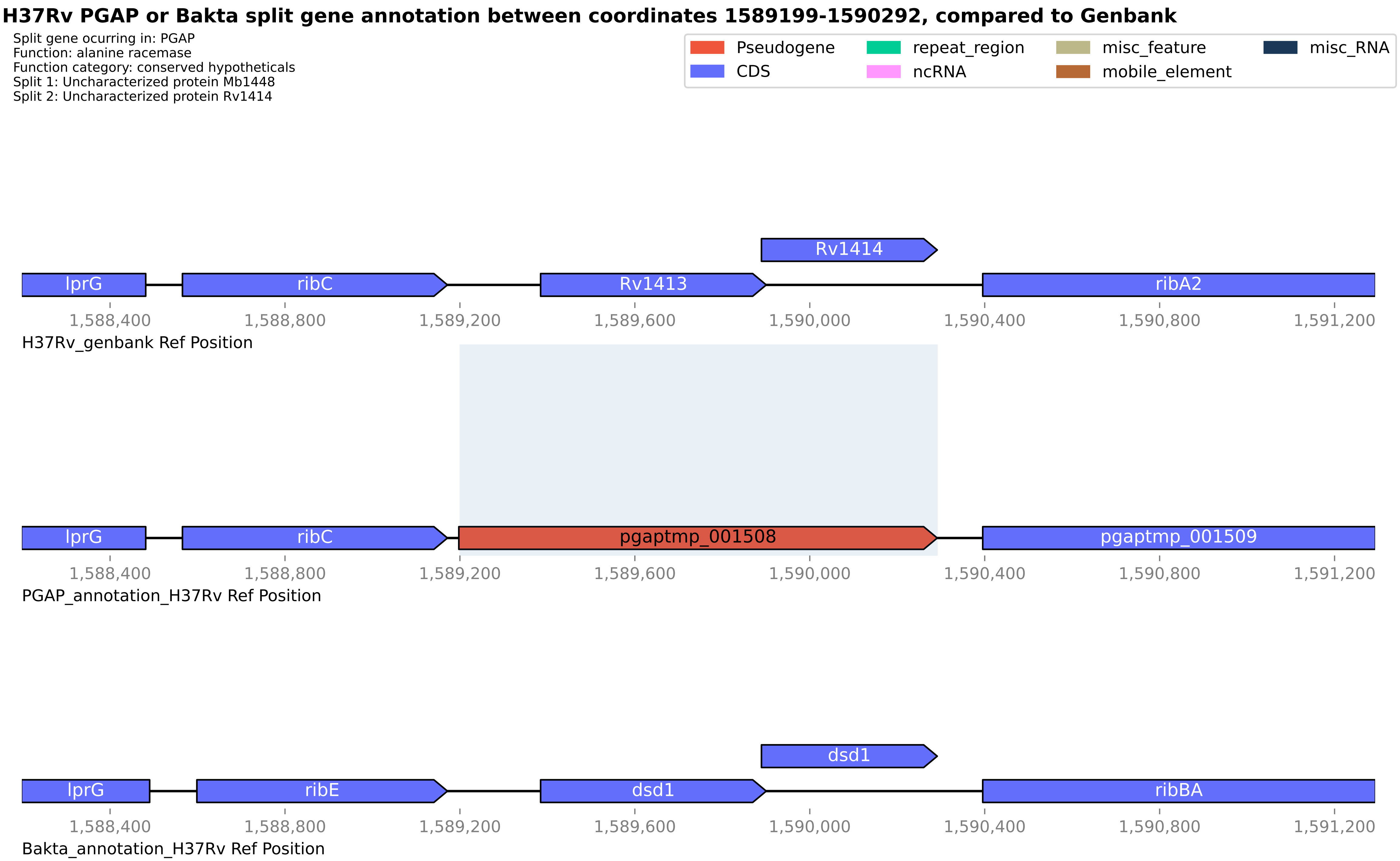
H37Rv PGAP or Bakta split gene annotation between coordinates 1589199-1590292, compared to Genbank
Split gene ocurring in: PGAP
Function: alanine racemase
Function category: conserved hypotheticals
Split 1: Uncharacterized protein Mb1448
Split 2: Uncharacterized protein Rv1414
Pseudogene
CDS
repeat_region
ncRNA
misc_feature
mobile_element
misc_RNA
Rv1414
lprG
ribC
Rv1413
ribA2
1,588,400
1,588,800
1,589,200
1,589,600
1,590,000
1,590,400
1,590,800
1,591,200
H37Rv_genbank Ref Position
lprG
ribC
pgaptmp_001508
pgaptmp_001509
PGAP_annotation_H37Rv Ref Position
dsd1
lprG
ribE
dsd1
ribBA
Bakta_annotation_H37Rv Ref Position

#### H37Rv PGAP or Bakta split gene annotation between coordinates 1693996-1695108, compared to Genbank

Split gene ocurring in: PGAP
Function: dTDP-4-amino-4,6-dideoxygalactose transaminase rffa
Function category: conserved hypotheticals
Split 1: TDP-4-oxo-6-deoxy-D-glucose aminotransferase
Split 2: dTDP-4-amino-4,6-dideoxygalactose transaminase rffa

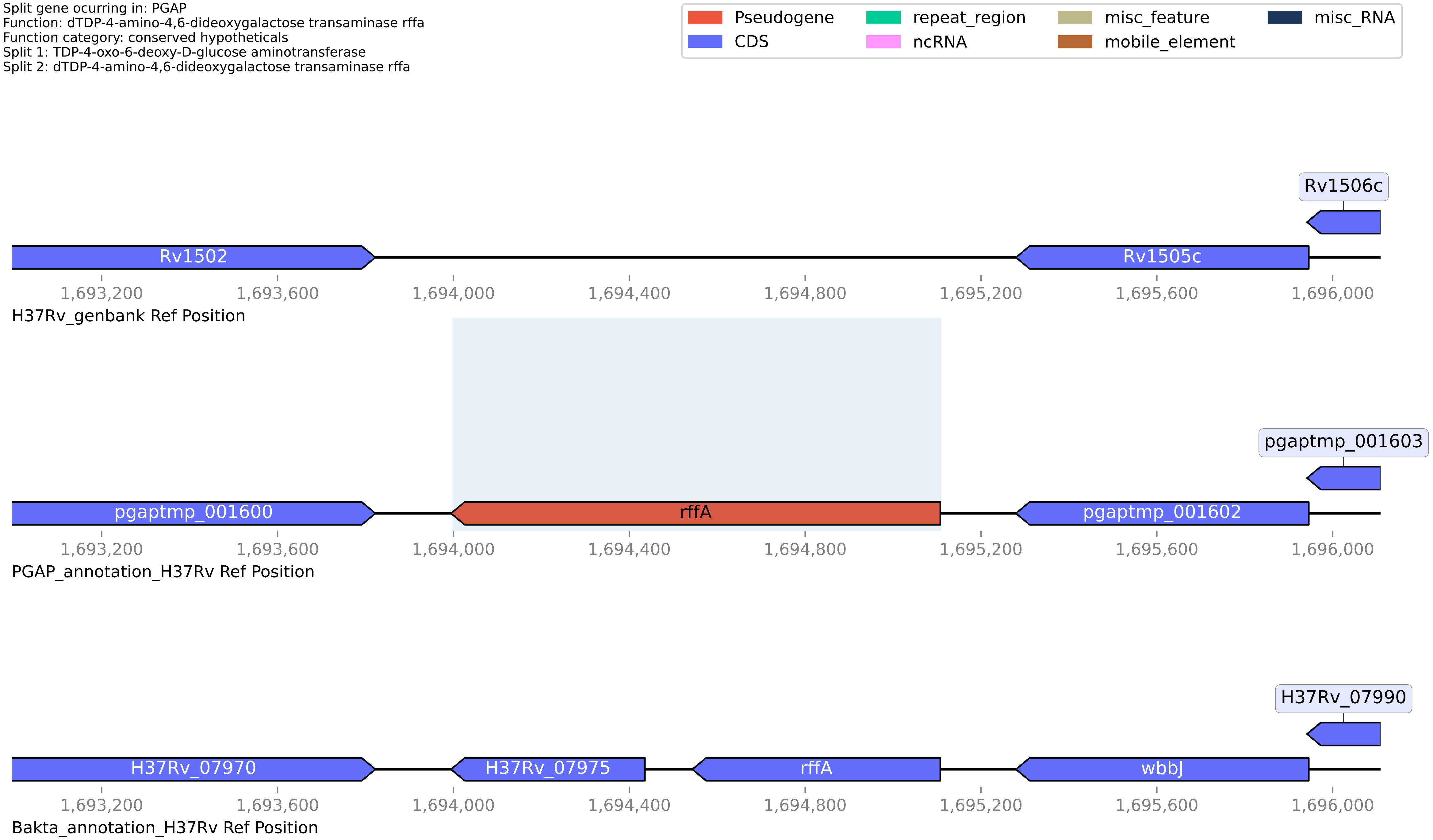

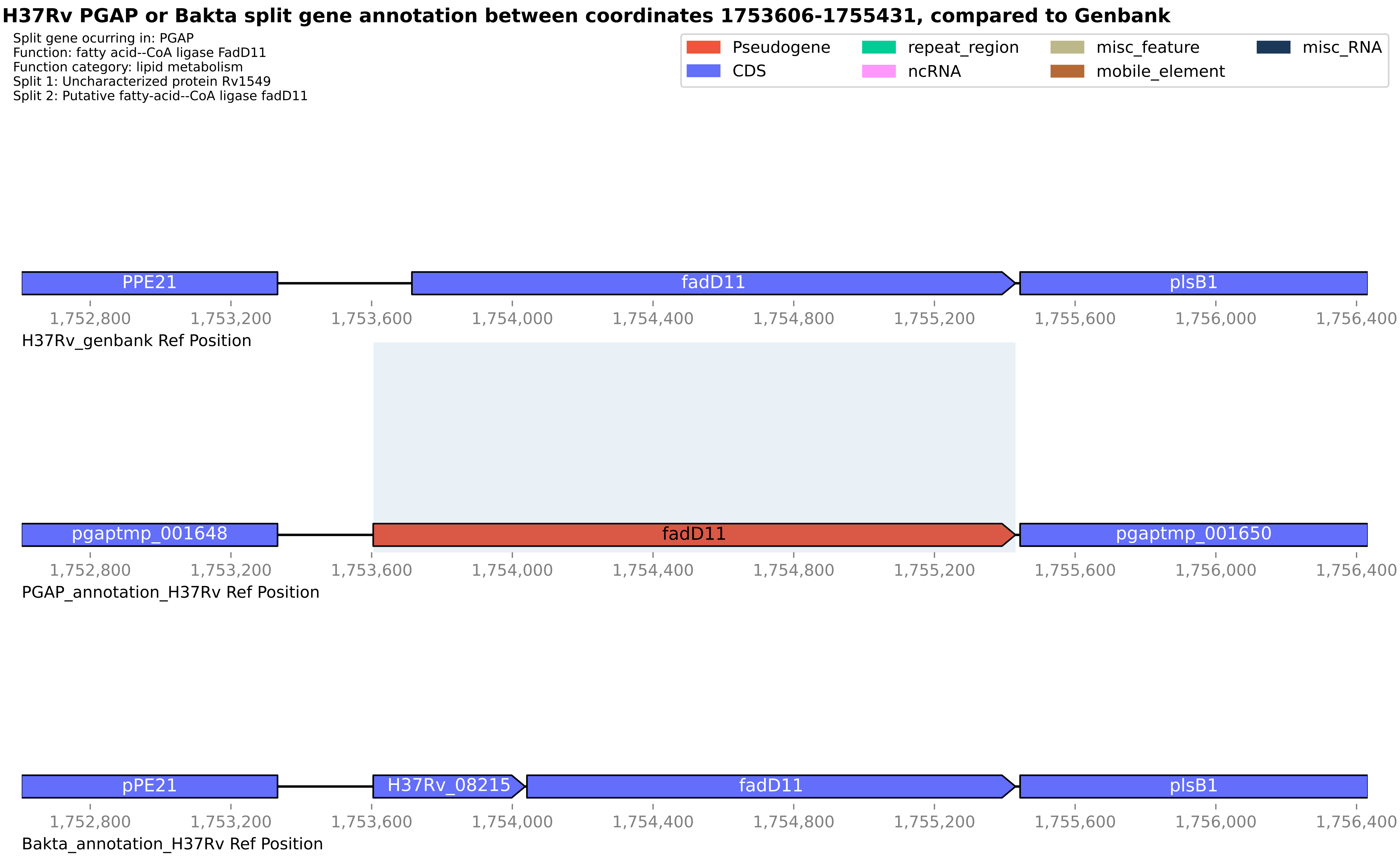
H37Rv PGAP or Bakta split gene annotation between coordinates 1753606-1755431, compared to Genbank
Split gene ocurring in: PGAP
Function: fatty acid--CoA ligase FadD11
Function category: lipid metabolism
Split 1: Uncharacterized protein Rv1549
Split 2: Putative fatty-acid--CoA ligase fadD11
Pseudogene
CDS
repeat_region
ncRNA
misc_feature
mobile_element
misc_RNA
PPE21
fadD11
plsB1
1,752,800
1,753,200
1,753,600
1,754,000
1,754,400
1,754,800
1,755,200
1,755,600
1,756,000
1,756,400
H37Rv_genbank Ref Position
pgaptmp_001648
fadD11
pgaptmp_001650
PGAP_annotation_H37Rv Ref Position
pPE21
H37Rv_08215
fadD11
plsB1
Bakta_annotation_H37Rv Ref Position

#### H37Rv PGAP or Bakta split gene annotation between coordinates 179319-181029, compared to Genbank

Split gene ocurring in: PGAP
Function: PE-PPE domain-containing protein
Function category: PE/PPE
Split 1: PE family protein
Split 2: PE-PGRS family protein

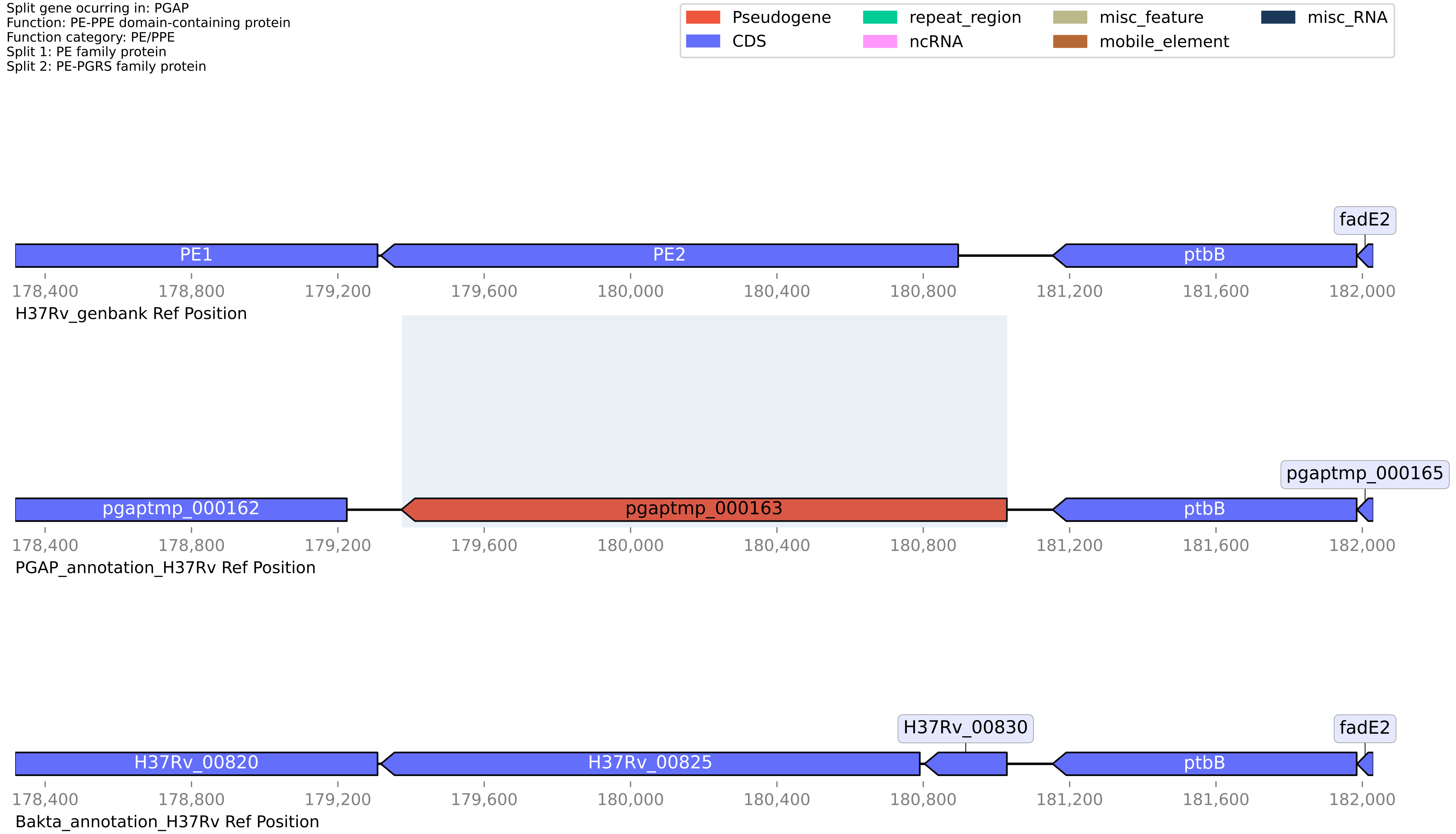

**H37Rv PGAP or Bakta split gene annotation between coordinates 1893577-1895342, compared to Genbank**

Split gene ocurring in: PGAP
Function: ABC-F family ATP-binding cassette domain-containing protein
Function category: cell wall and cell processes
Split 1: Macrolide-transport ATP-binding protein ABC transporter first part
Split 2: Macrolide ABC transporter ATP-binding protein second part

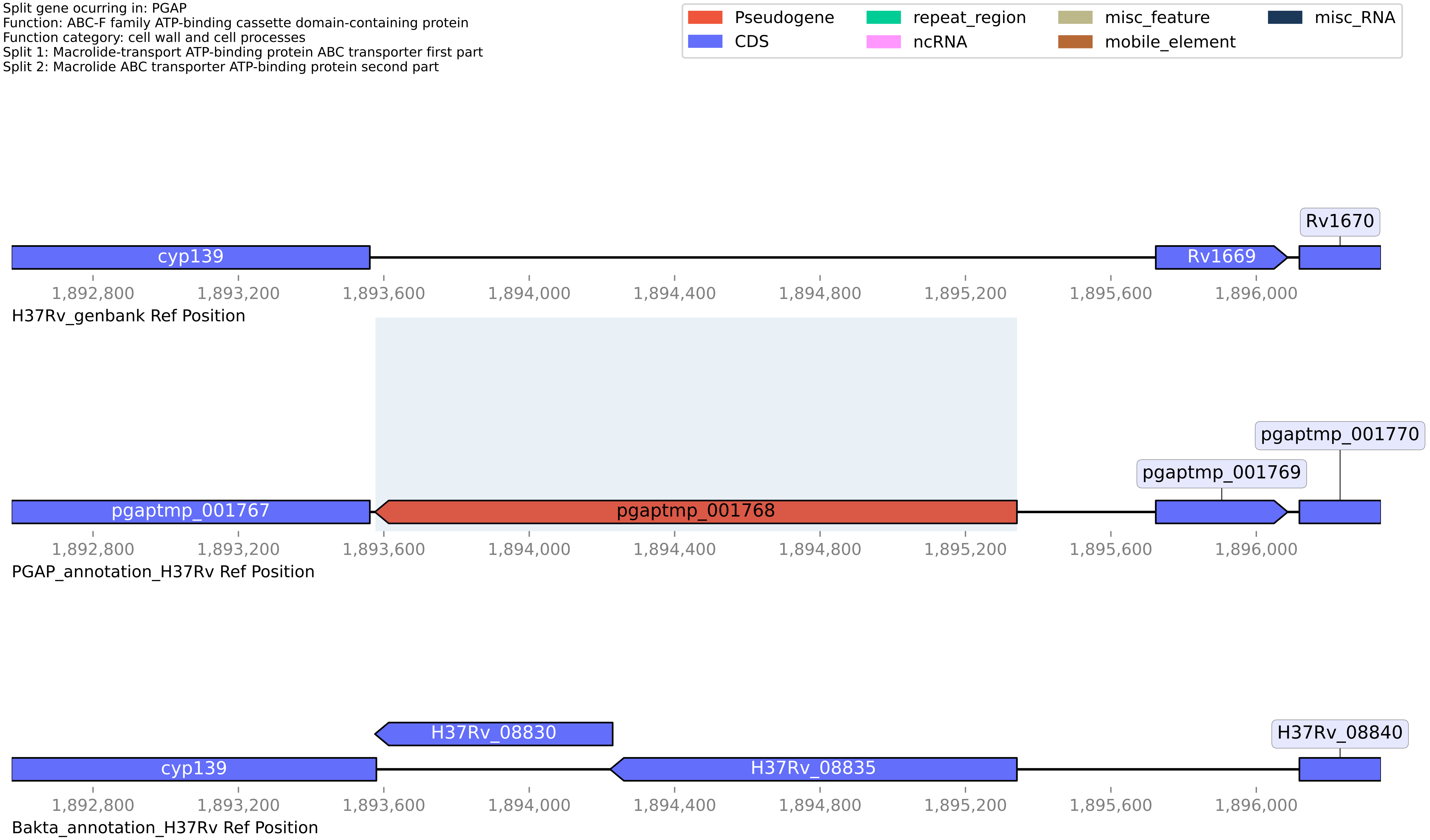

#### H37Rv PGAP or Bakta split gene annotation between coordinates 2030347-2030643, compared to Genbank

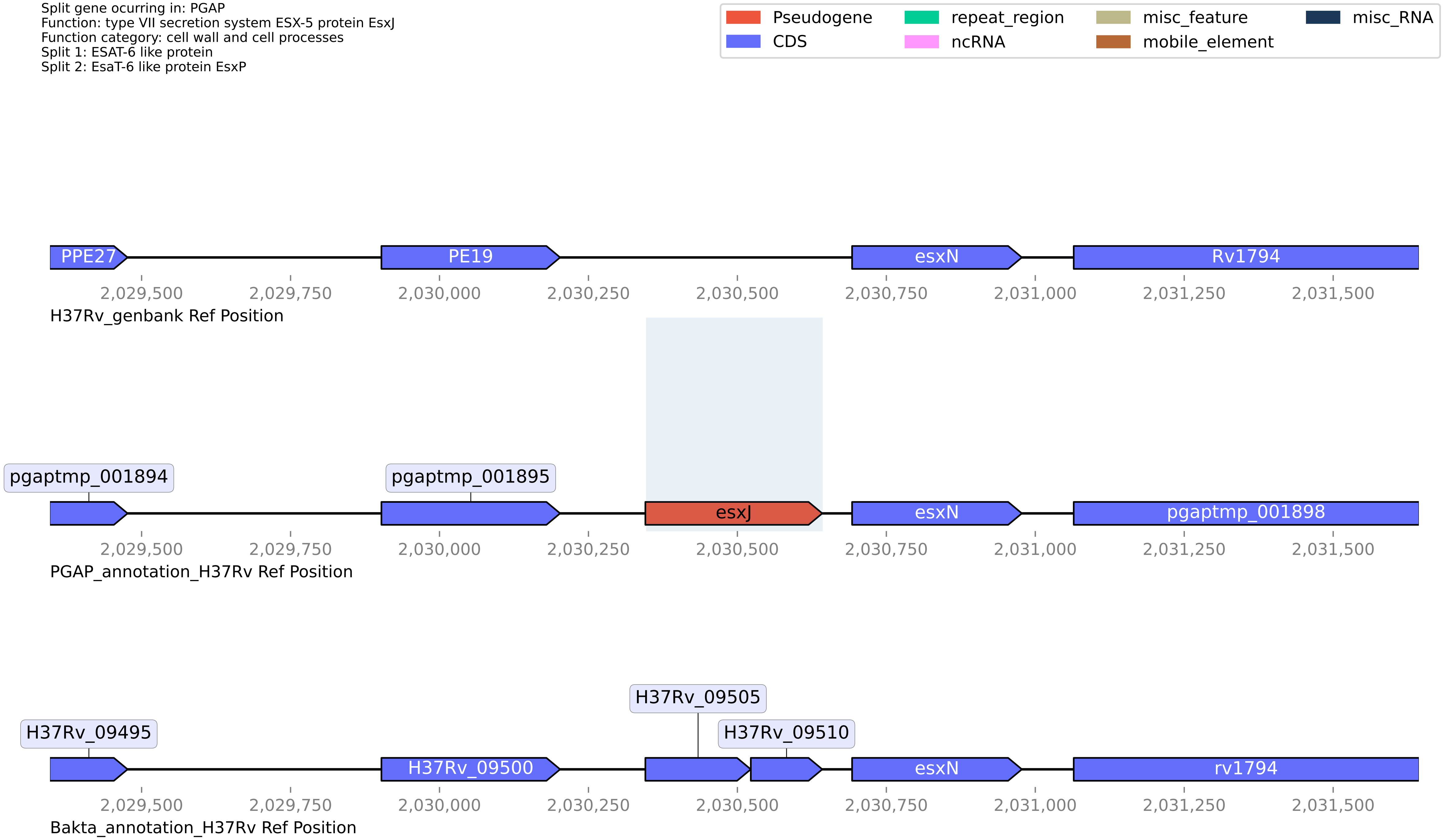

**H37Rv PGAP or Bakta split gene annotation between coordinates 2138174-2139017, compared to Genbank**

Split gene ocurring in: PGAP
Function: class I SAM-dependent methyltransferase
Function category: conserved hypotheticals
Split 1: O-methyltransferase
Split 2: S-adenosyl-L-methionine-dependent methyltransferase (Part1)

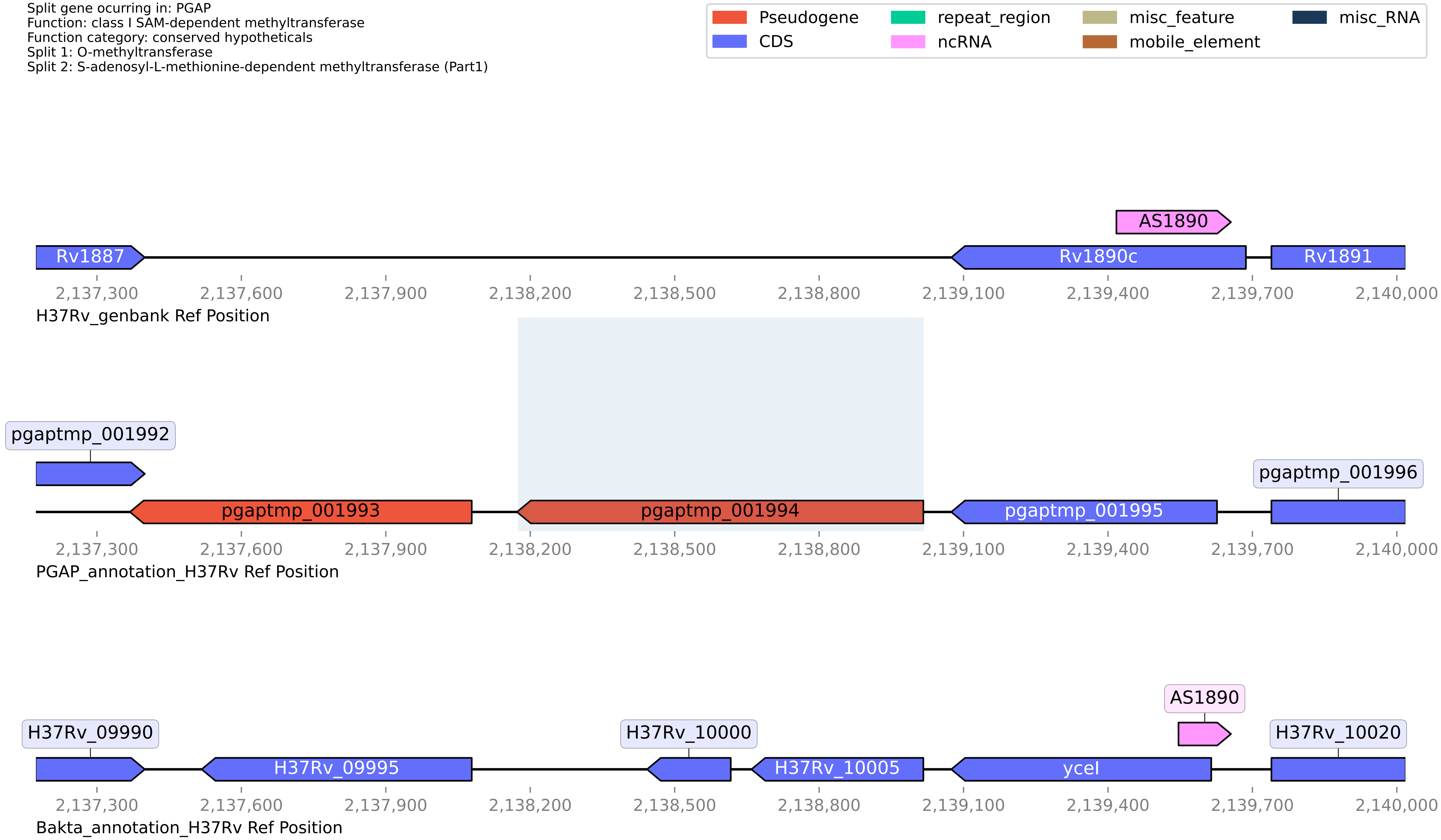

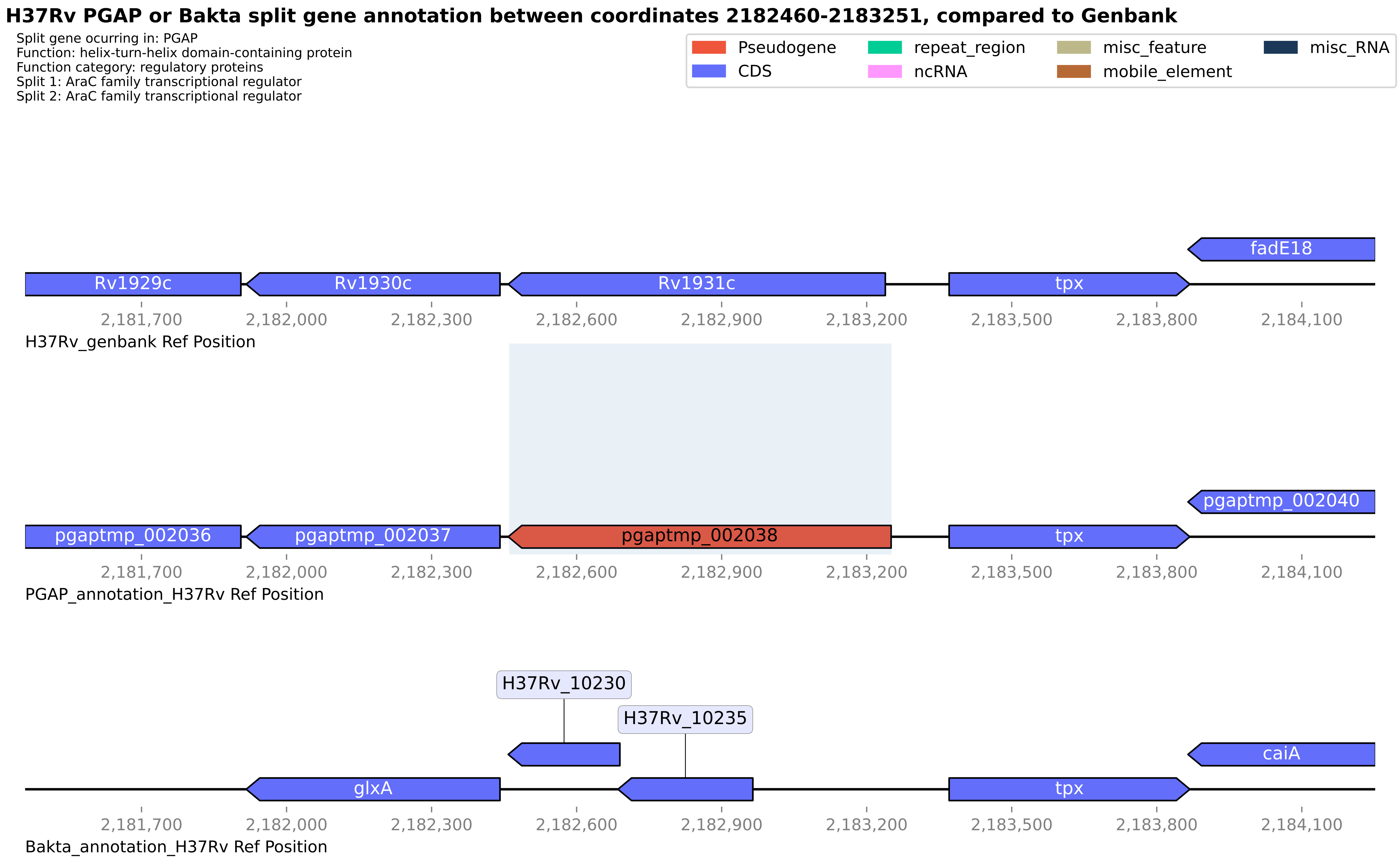
H37Rv PGAP or Bakta split gene annotation between coordinates 2182460-2183251, compared to Genbank
Split gene ocurring in: PGAP
Function: helix-turn-helix domain-containing protein
Function category: regulatory proteins
Split 1: AraC family transcriptional regulator
Split 2: AraC family transcriptional regulator
Pseudogene
CDS
repeat_region
ncRNA
misc_feature
mobile_element
misc_RNA
fadE18
Rv1929c
Rv1930c
Rv1931c
tpx
2,181,700
2,182,000
2,182,300
2,182,600
2,182,900
2,183,200
2,183,500
2,183,800
2,184,100
H37Rv_genbank Ref Position
pgaptmp_002040
pgaptmp_002036
pgaptmp_002037
pgaptmp_002038
tpx
PGAP_annotation_H37Rv Ref Position
H37Rv_10230
H37Rv_10235
caiA
glxA
tpx
Bakta_annotation_H37Rv Ref Position

#### H37Rv PGAP or Bakta split gene annotation between coordinates 2356729-2358206, compared to Genbank

Split gene ocurring in: PGAP
Function: PE family protein
Function category: PE/PPE
Split 1: PE domain-containing protein
Split 2: PE-PGRS family protein

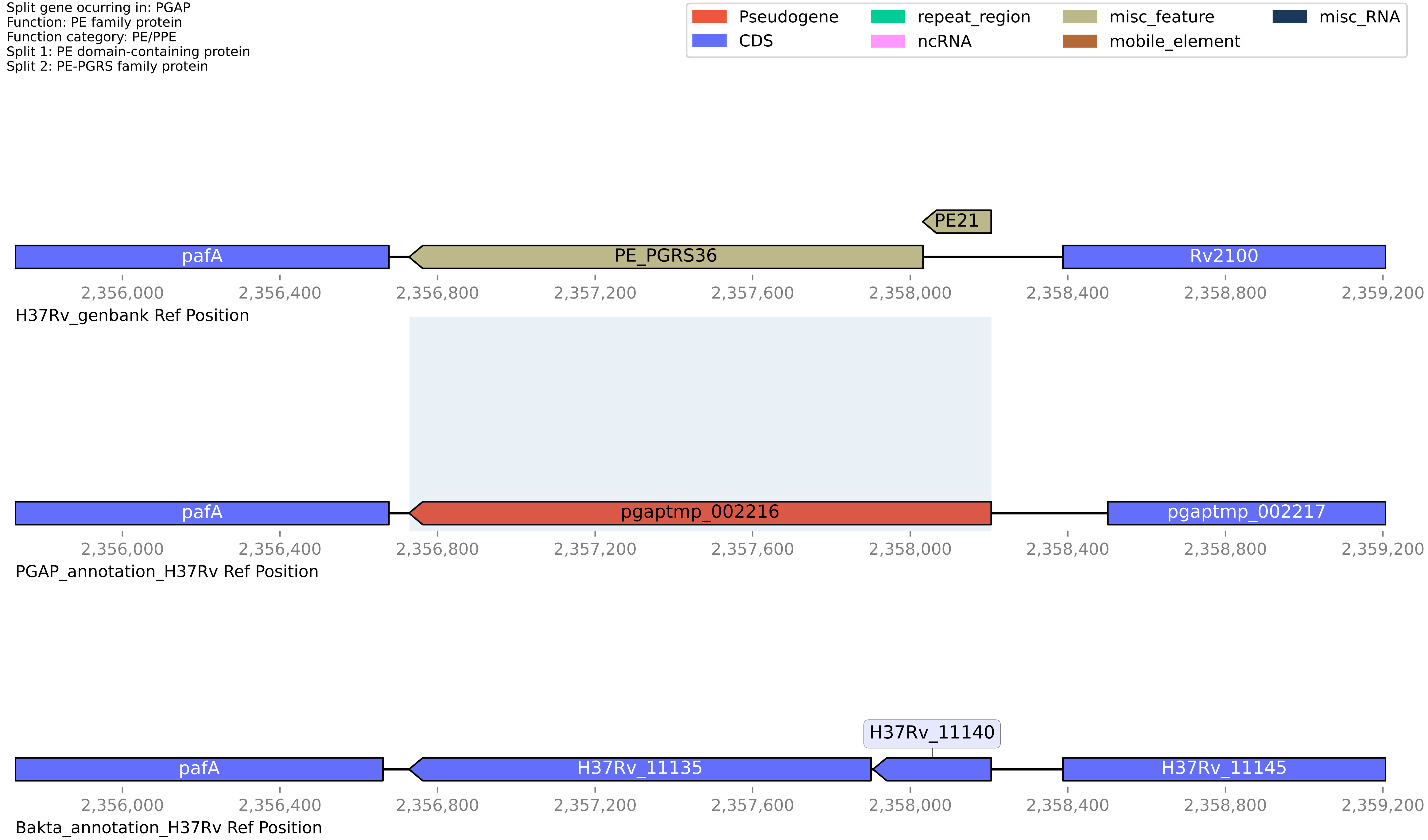

**H37Rv PGAP or Bakta split gene annotation between coordinates 2500923-2501632, compared to Genbank**

Split gene ocurring in: PGAP
Function: 2OG-Fe(II) oxygenase
Function category: conserved hypotheticals
Split 1: proline hydroxylase
Split 2: DUF2086 domain-containing protein

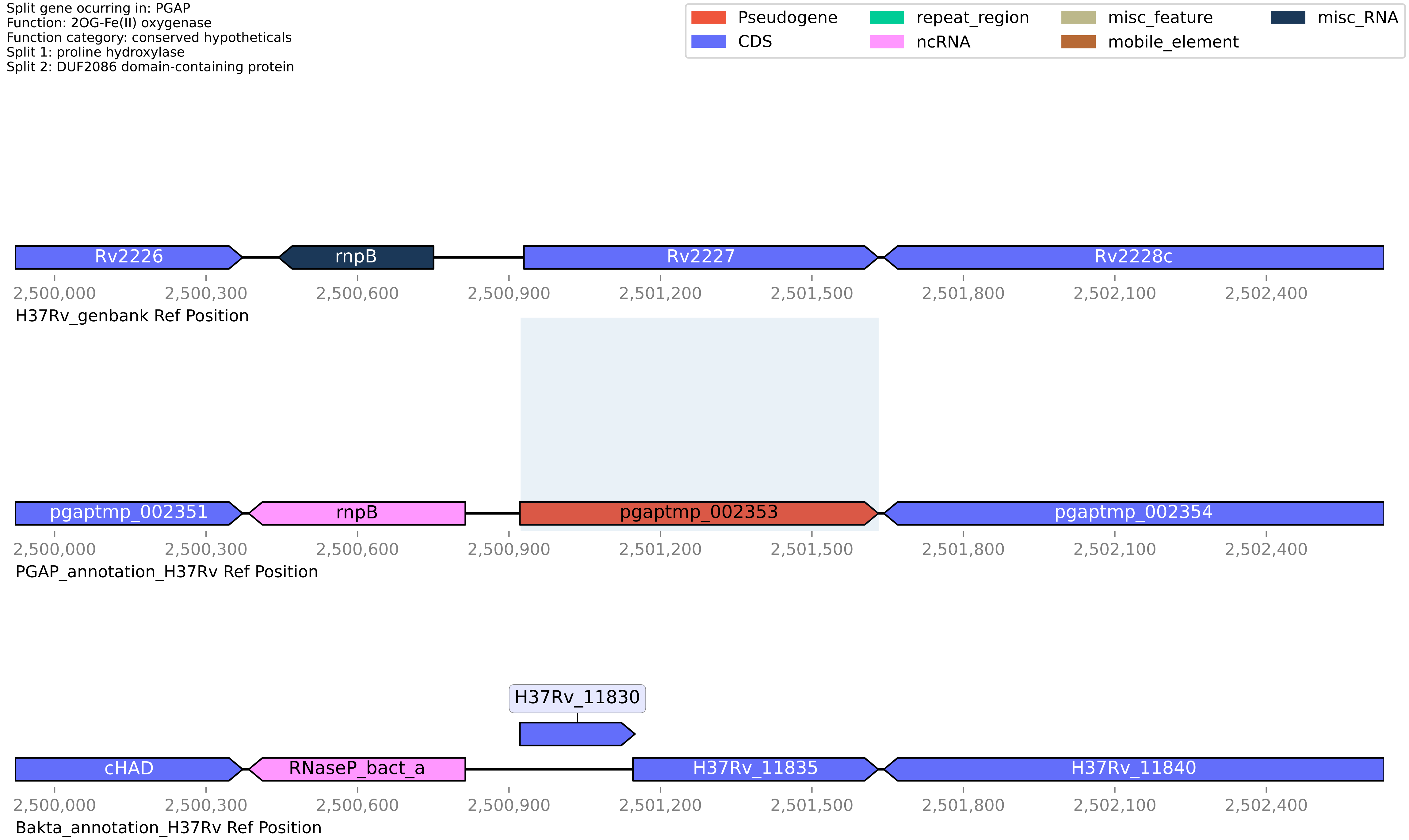

#### H37Rv PGAP or Bakta split gene annotation between coordinates 2525402-2526992, compared to Genbank

Split gene ocurring in: PGAP
Function: FAD-binding oxidoreductase
Function category: intermediary metabolism and respiration
Split 1: putative flavoprotein
Split 2: FAD/FMN-containing lactate dehydrogenase/glycolate oxidase (glcD)

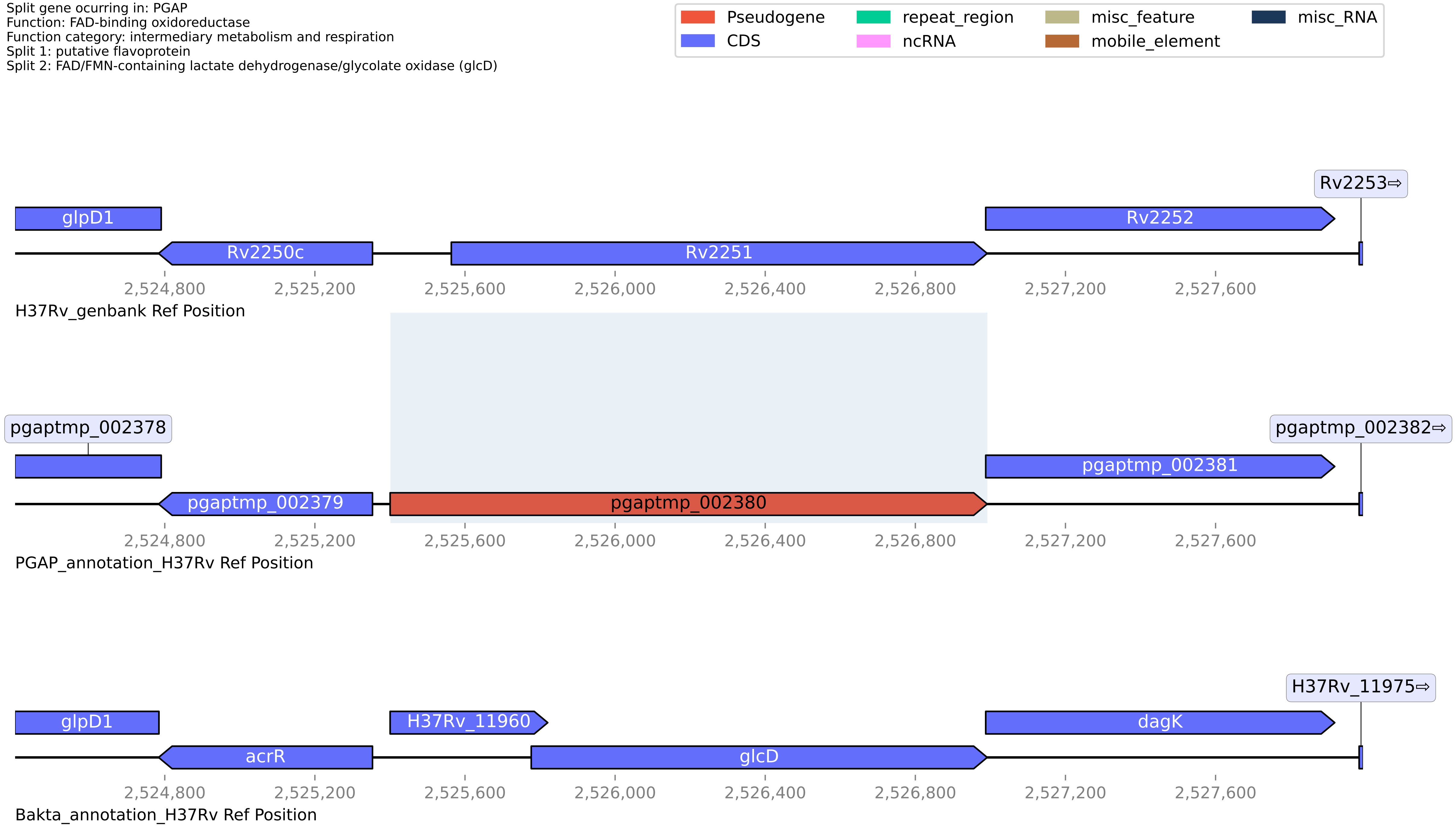

**H37Rv PGAP or Bakta split gene annotation between coordinates 2534042-2535552, compared to Genbank**

Split gene ocurring in: PGAP
Function: apolipoprotein N-acyltransferase Int
Function category: lipid metabolism
Split 1: CN hydrolase domain-containing protein
Split 2: apolipoprotein N-acyltransferase Int

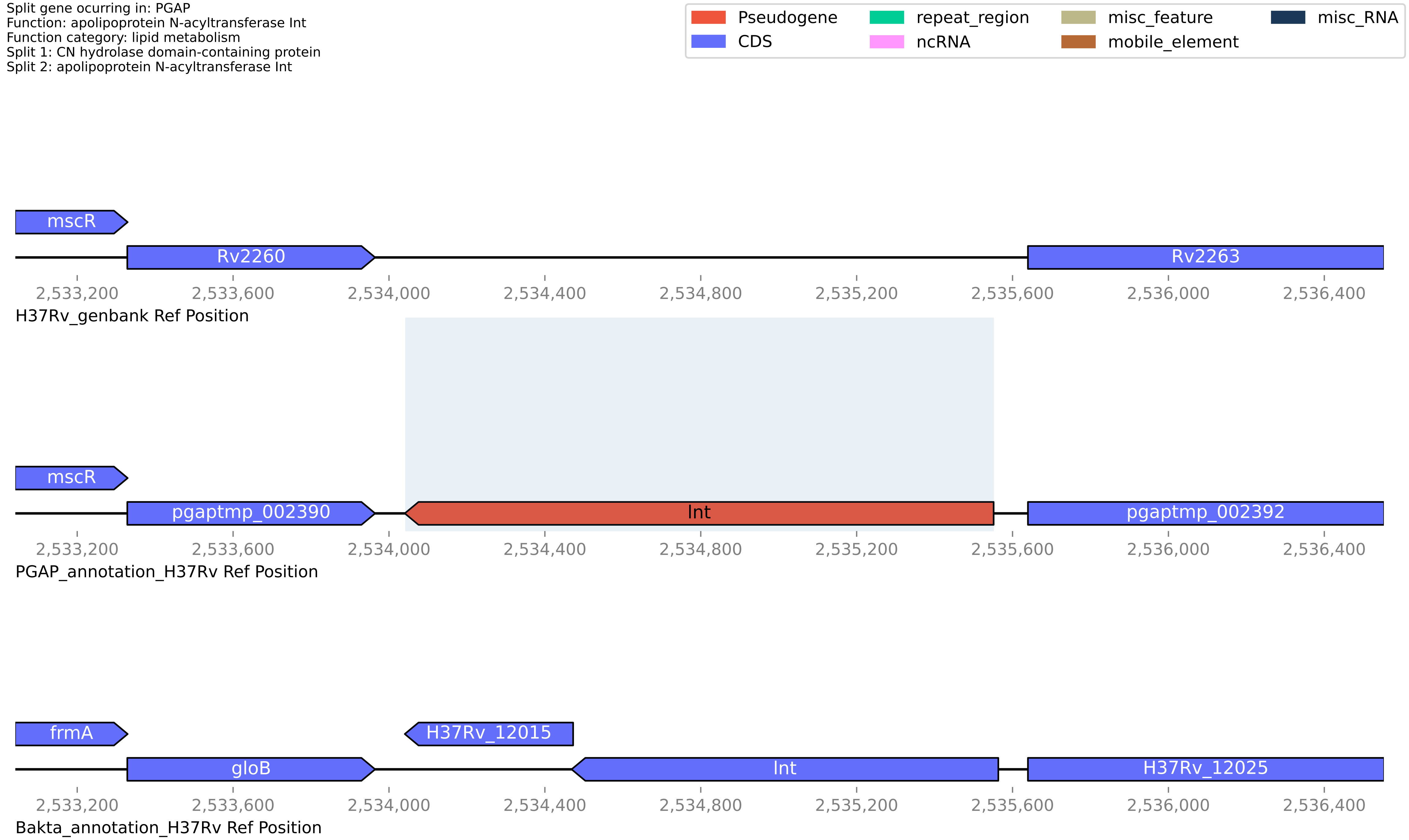

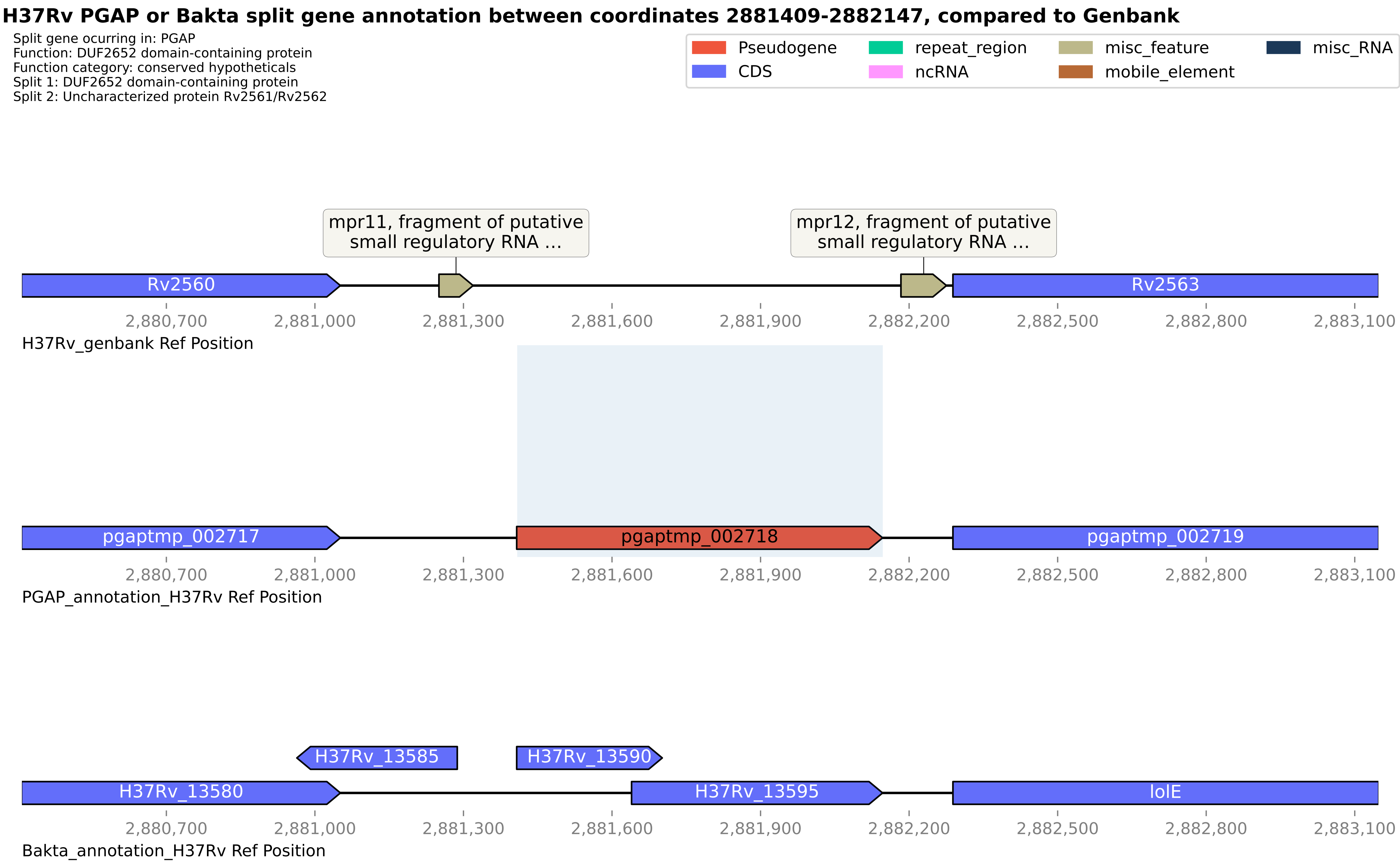
H37Rv PGAP or Bakta split gene annotation between coordinates 2881409-2882147, compared to Genbank
Split gene ocurring in: PGAP
Function: DUF2652 domain-containing protein
Function category: conserved hypotheticals
Split 1: DUF2652 domain-containing protein
Split 2: Uncharacterized protein Rv2561/Rv2562
Pseudogene
CDS
repeat_region
ncRNA
misc_feature
mobile_element
misc_RNA
mpr11, fragment of putative small regulatory RNA ...
mpr12, fragment of putative small regulatory RNA ...
Rv2560
Rv2563
2,880,700
2,881,000
2,881,300
2,881,600
2,881,900
2,882,200
2,882,500
2,882,800
2,883,100
H37Rv_genbank Ref Position
pgaptmp_002717
pgaptmp_002718
pgaptmp_002719
PGAP_annotation_H37Rv Ref Position
H37Rv_13585
H37Rv_13590
H37Rv_13580
H37Rv_13595
lolE
Bakta_annotation_H37Rv Ref Position

**H37Rv PGAP or Bakta split gene annotation between coordinates 3291503-3297819, compared to Genbank**

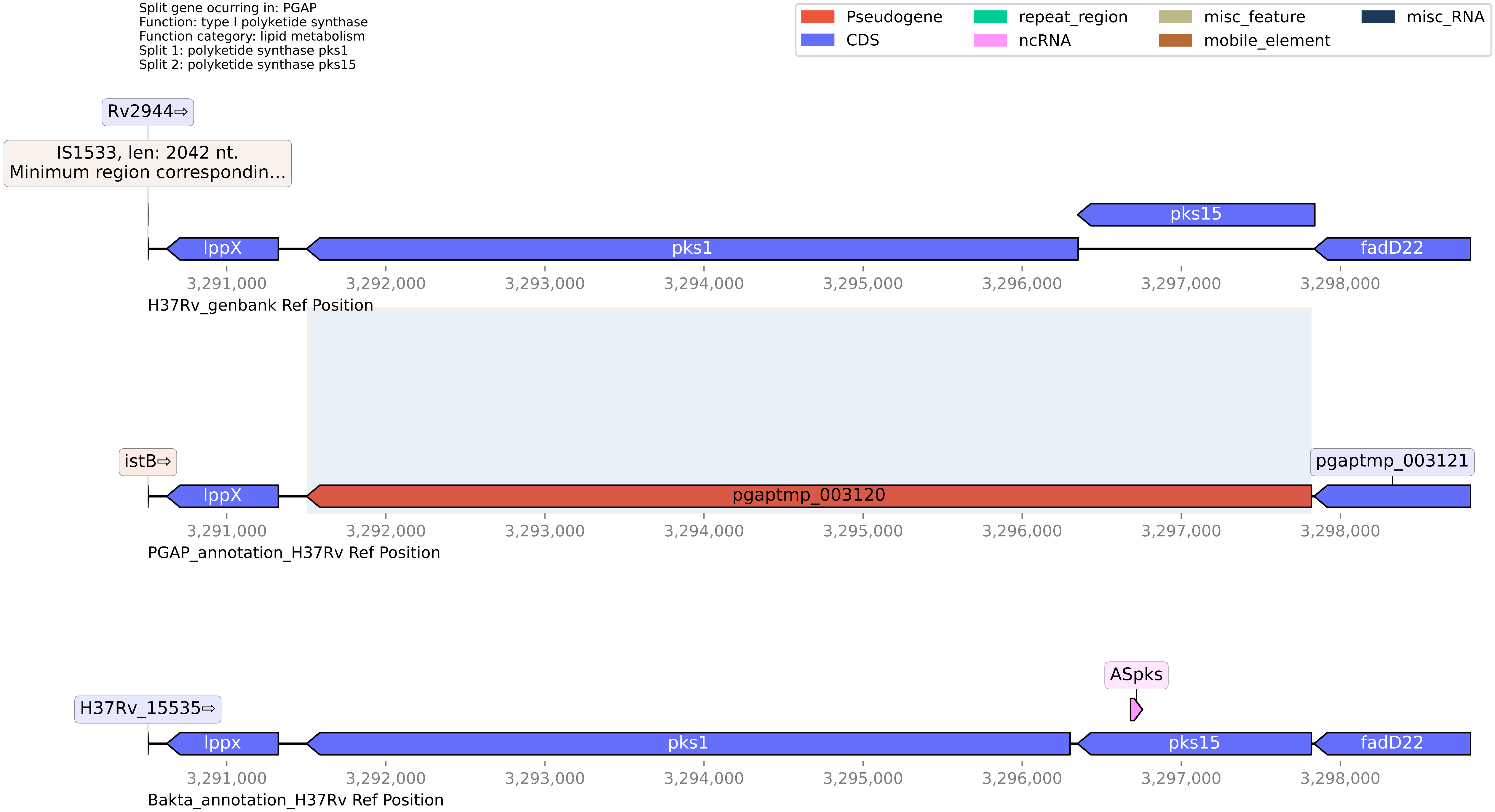

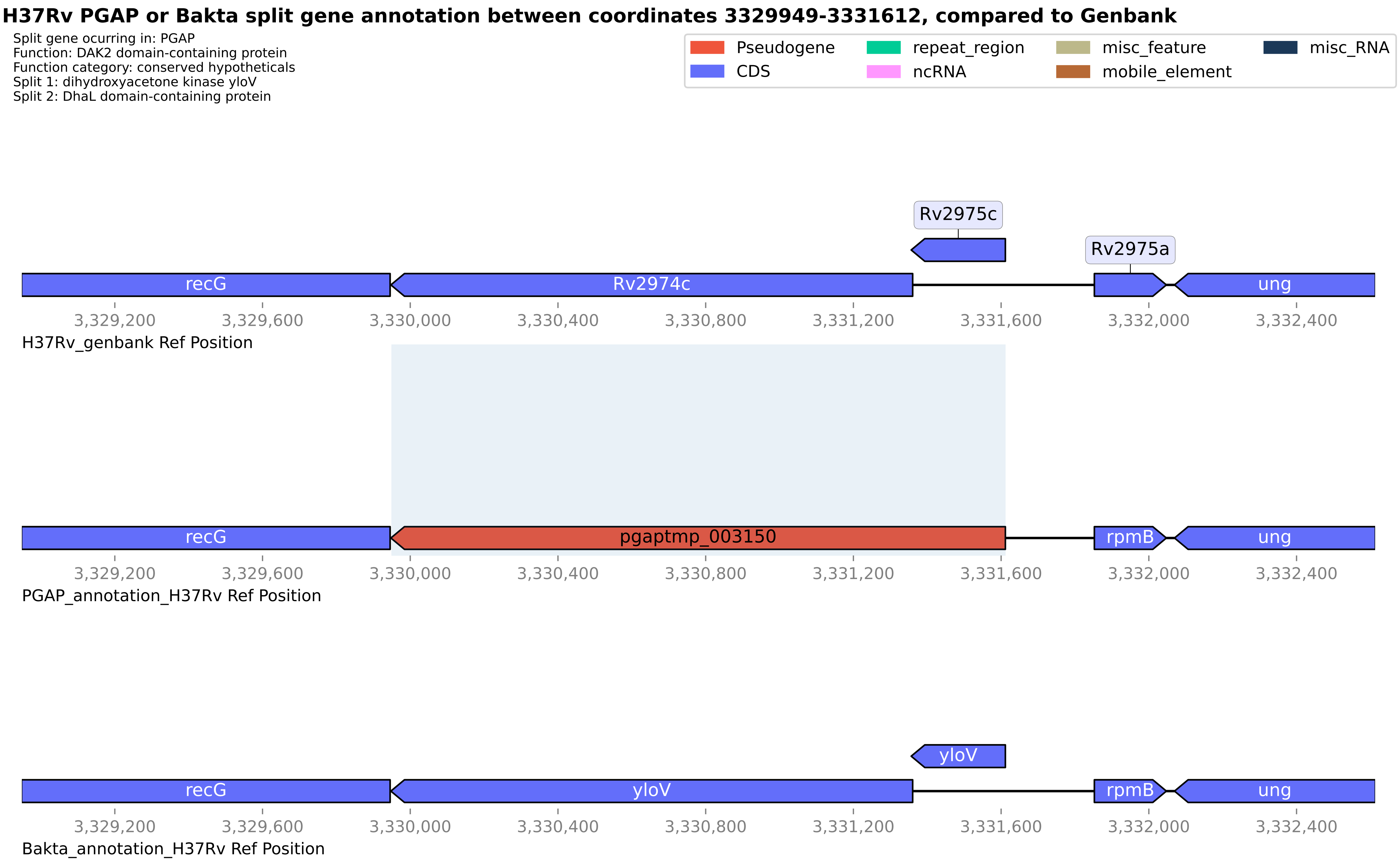
H37Rv PGAP or Bakta split gene annotation between coordinates 3329949-3331612, compared to Genbank
Split gene ocurring in: PGAP
Function: DAK2 domain-containing protein
Function category: conserved hypotheticals
Split 1: dihydroxyacetone kinase yloV
Split 2: DhaL domain-containing protein
Pseudogene
CDS
repeat_region
ncRNA
misc_feature
mobile_element
misc_RNA
Rv2975c
Rv2975a
recG
Rv2974c
ung
3,329,200
3,329,600
3,330,000
3,330,400
3,330,800
3,331,200
3,331,600
3,332,000
3,332,400
H37Rv_genbank Ref Position
recG
pgaptmp_003150
rpmB
ung
PGAP_annotation_H37Rv Ref Position
yloV
recG
yloV
rpmB
ung
Bakta_annotation_H37Rv Ref Position

**H37Rv PGAP or Bakta split gene annotation between coordinates 3435718-3436295, compared to Genbank**

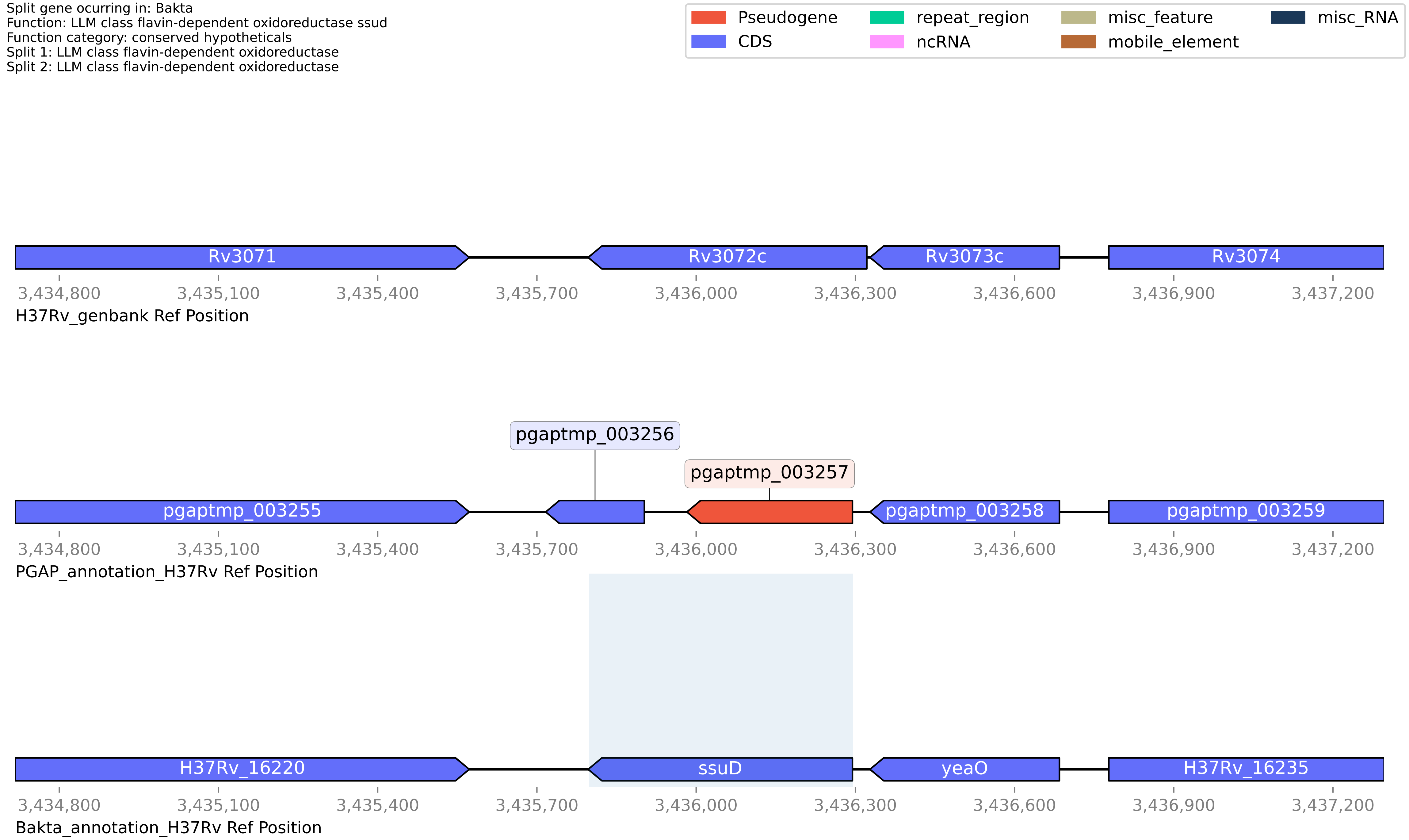

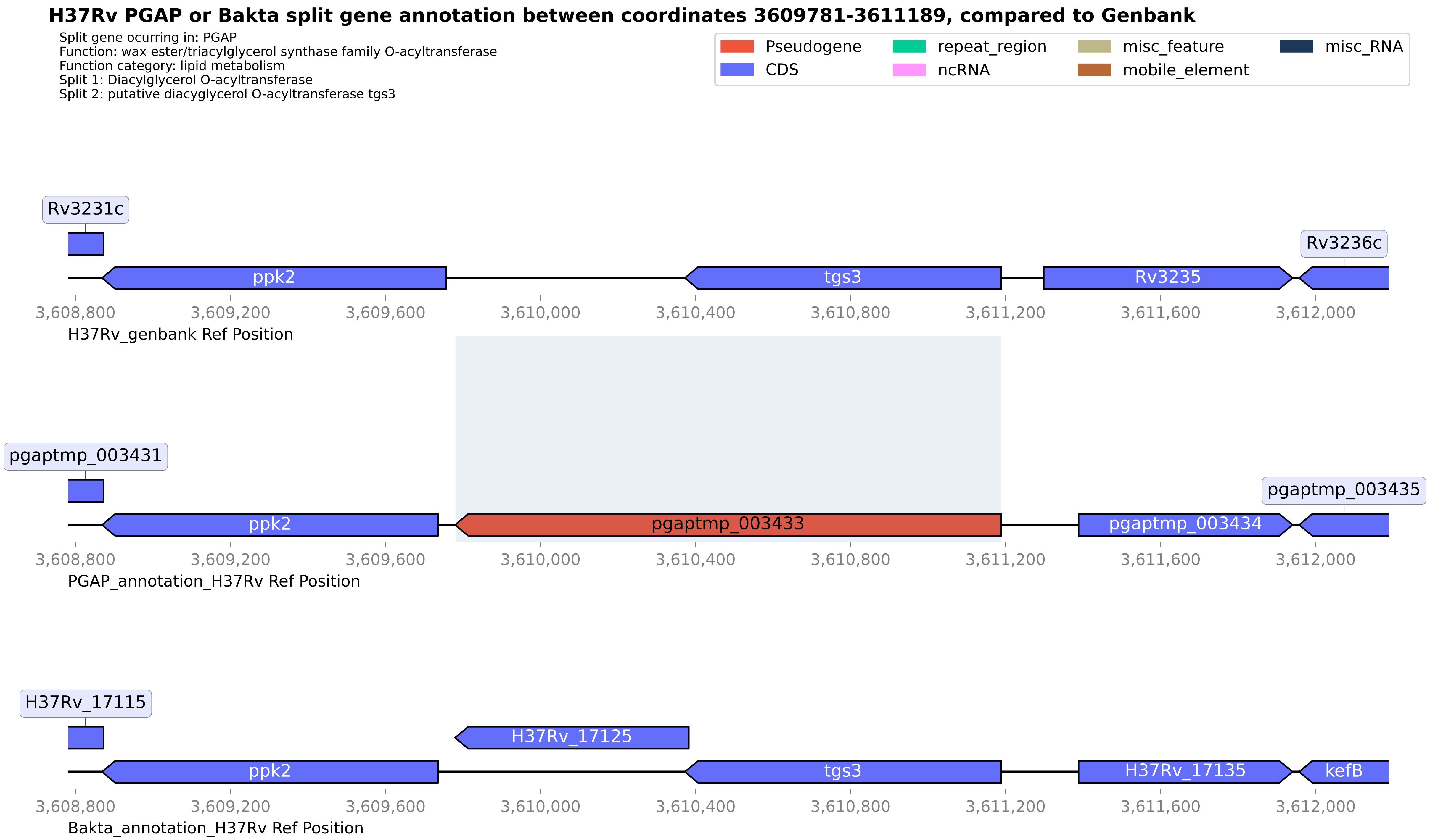
H37Rv PGAP or Bakta split gene annotation between coordinates 3609781-3611189, compared to Genbank
Split gene ocurring in: PGAP
Function: wax ester/triacylglycerol synthase family O-acyltransferase
Function category: lipid metabolism
Split 1: Diacylglycerol O-acyltransferase
Split 2: putative diacyglycerol O-acyltransferase tgs3
Pseudogene
CDS
repeat_region
ncRNA
misc_feature
mobile_element
misc_RNA
Rv3231c
ppk2
tgs3
Rv3235
Rv3236c
3,608,800
3,609,200
3,609,600
3,610,000
3,610,400
3,610,800
3,611,200
3,611,600
3,612,000
H37Rv_genbank Ref Position
pgaptmp_003431
ppk2
pgaptmp_003433
pgaptmp_003434
pgaptmp_003435
PGAP_annotation_H37Rv Ref Position
H37Rv_17115
H37Rv_17125
ppk2
tgs3
H37Rv_17135
kefB
Bakta_annotation_H37Rv Ref Position

**H37Rv PGAP or Bakta split gene annotation between coordinates 366150-372764, compared to Genbank**

Split gene ocurring in: Bakta
Function: PPE family
Function category: PE/PPE
Split 1: hypothetical protein
Split 2: pseudogene

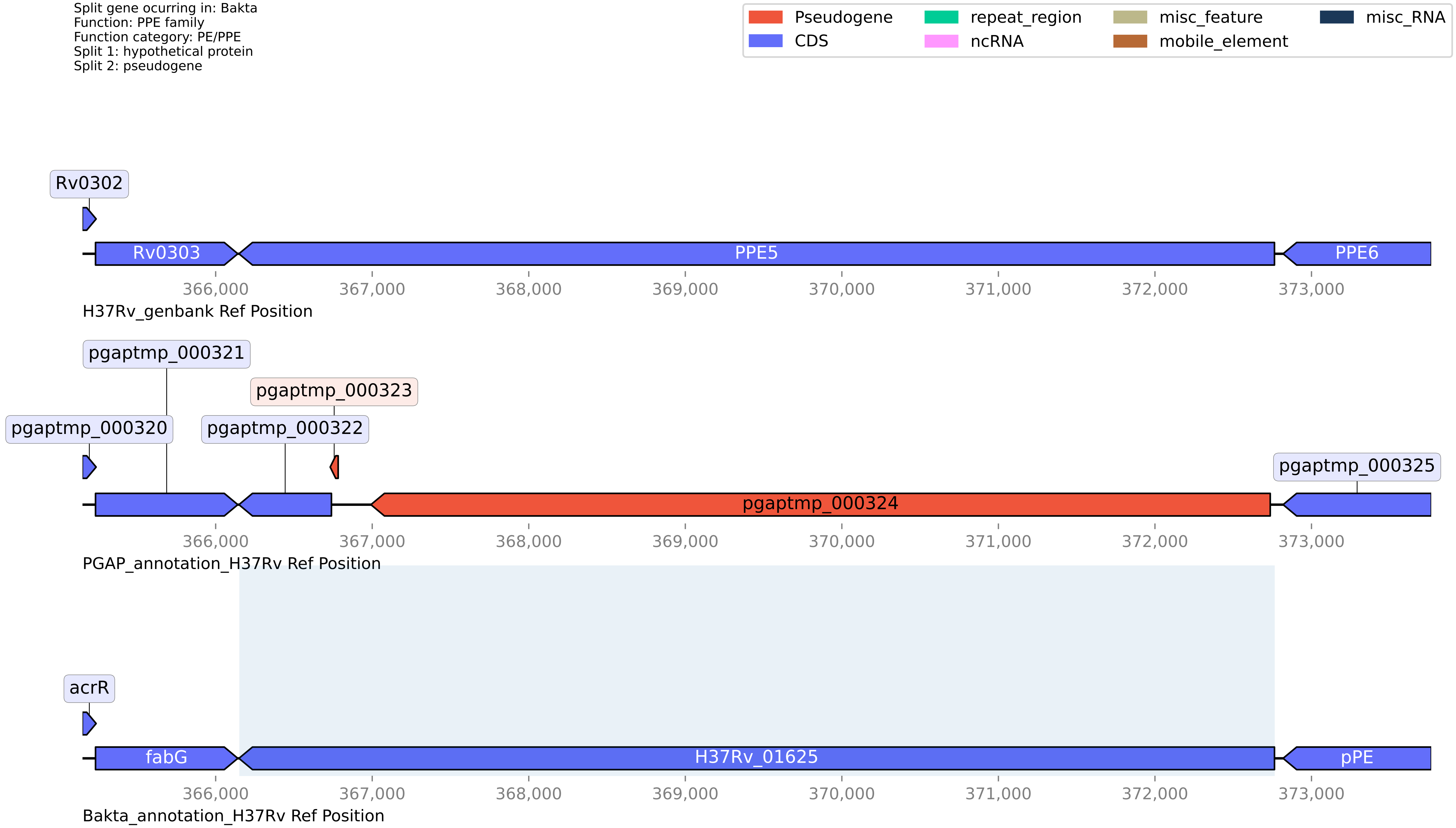

**H37Rv PGAP or Bakta split gene annotation between coordinates 3800017-3801463, compared to Genbank**

Split gene ocurring in: PGAP
Function: ISNCY family transposase
Function category: insertion seqs and phages
Split 1: Transposase and inactivated derivatives, IS5 family
Split 2: Transposase

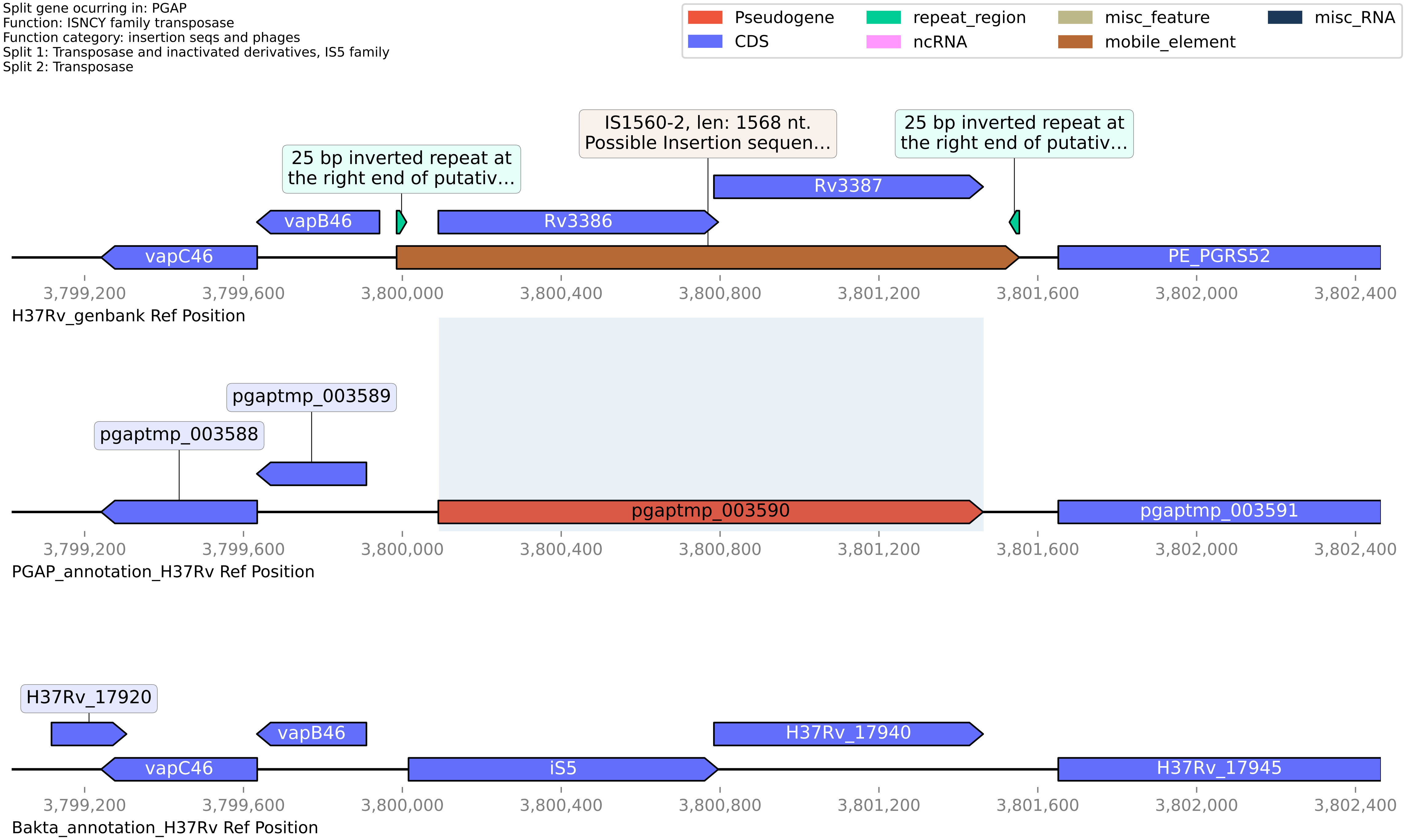

**H37Rv PGAP or Bakta split gene annotation between coordinates 3874404-3876090, compared to Genbank**

Split gene ocurring in: PGAP
Function: hypothetical protein
Function category: cell wall and cell processes
Split 1: Transmembrane protein
Split 2: Transmembrane protein

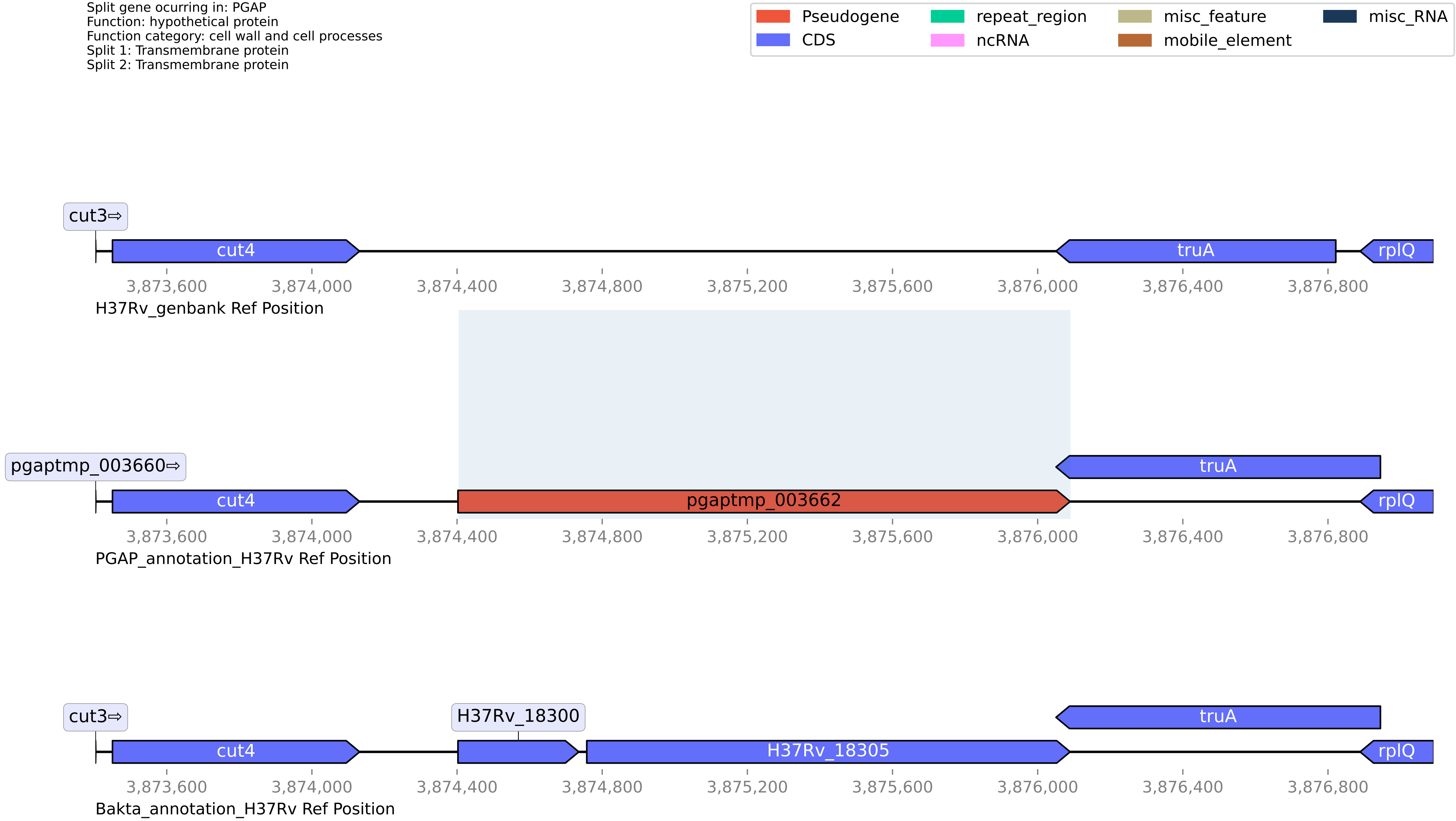

**H37Rv PGAP or Bakta split gene annotation between coordinates 4075752-4076984, compared to Genbank**

Split gene ocurring in: PGAP
Function: IS21 family transposase
Function category: insertion seqs and phages
Split 1: putative transposase
Split 2: IS21 family transposase

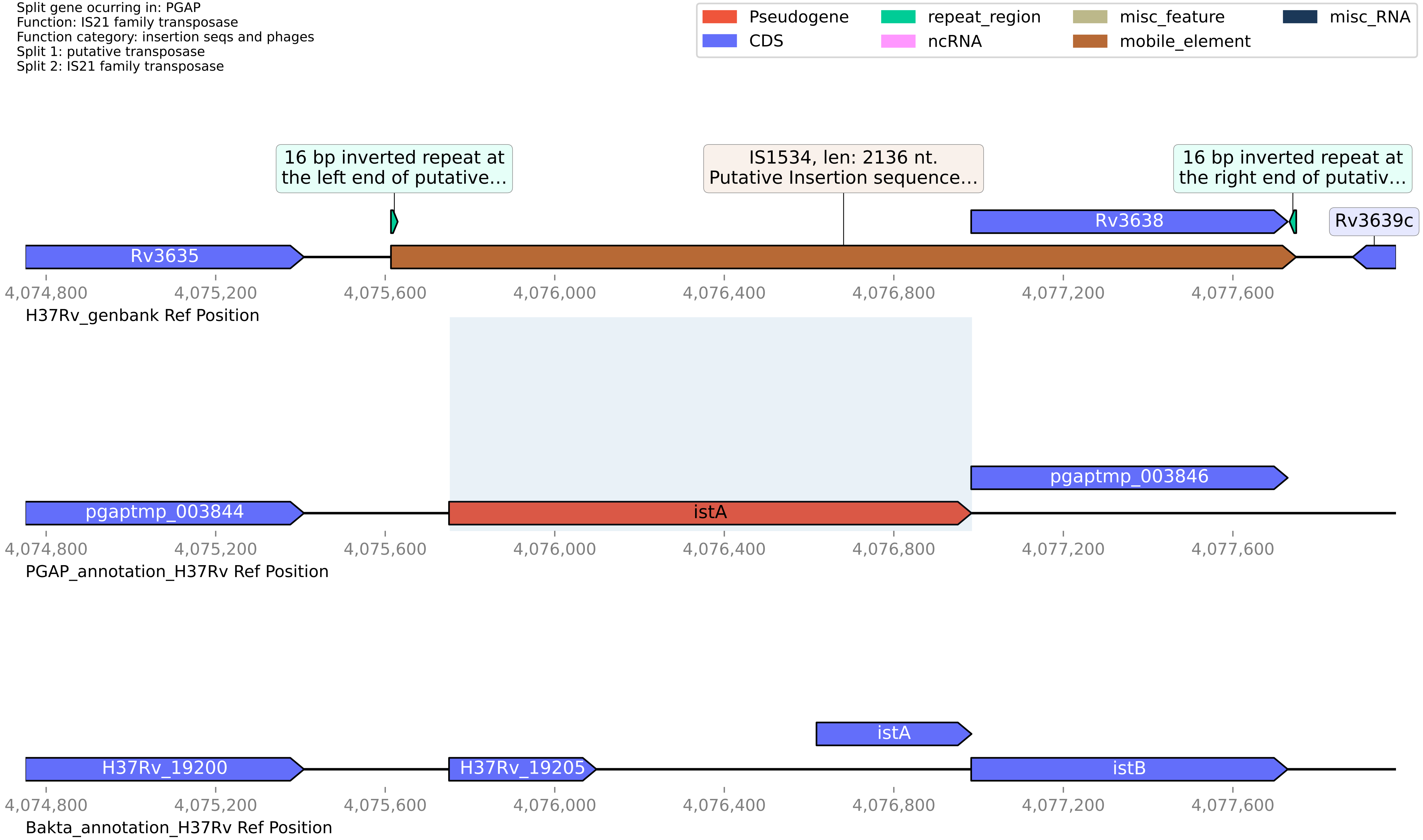

**H37Rv PGAP or Bakta split gene annotation between coordinates 4189285-4190517, compared to Genbank**

Split gene ocurring in: PGAP
Function: PPE family protein
Function category: PE/PPE
Split 1: Uncharacterized PPE family protein PPE66
Split 2: Uncharacterized PPE family protein PPE66

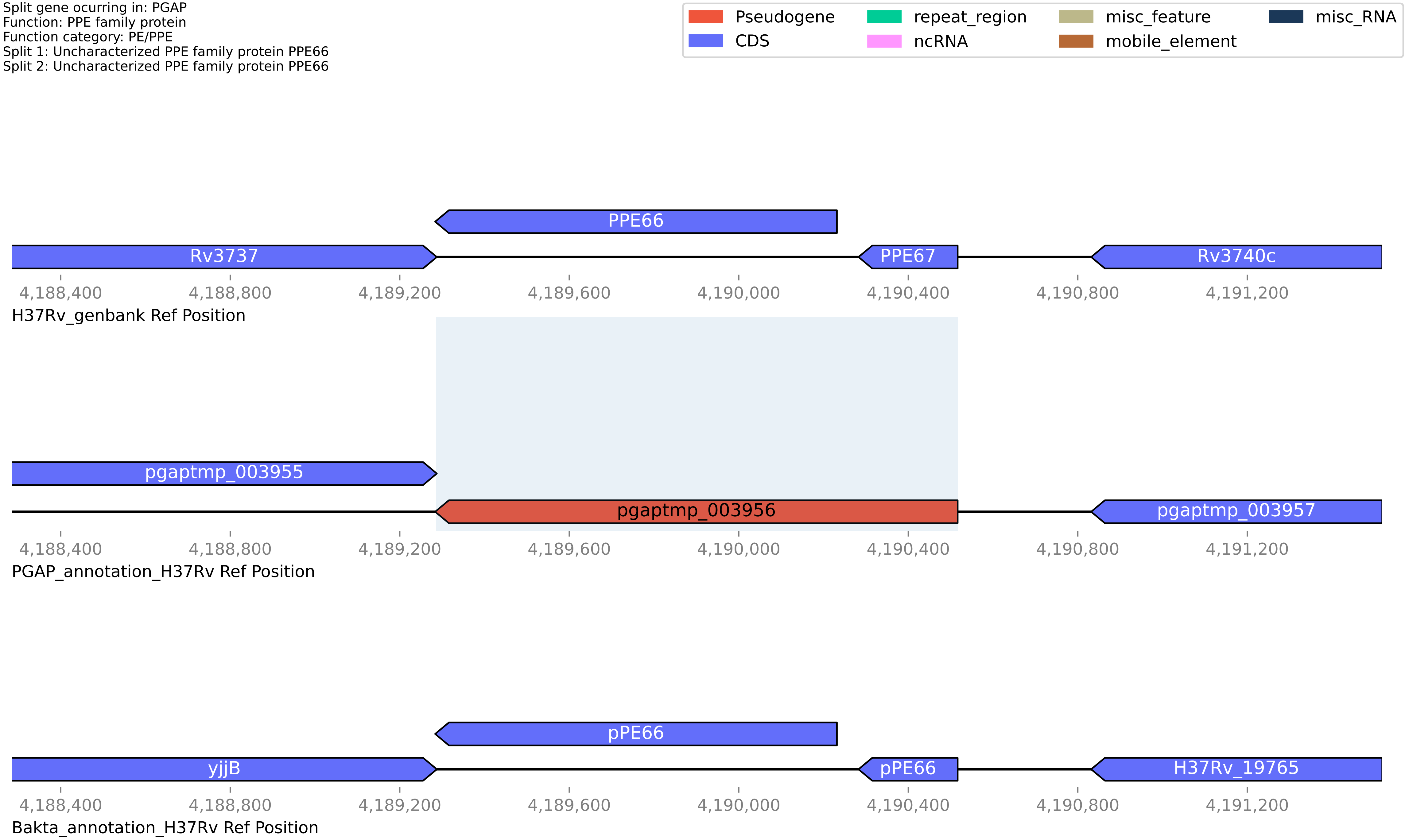

### H37Rv PGAP or Bakta split gene annotation between coordinates 4192179-4193245, compared to Genbank

Split gene ocurring in: PGAP
Function: NAD(P)/FAD-dependent oxidoreductase
Function category: intermediary metabolism and respiration
Split 1: NAD(P)/FAD-dependent oxidoreductase
Split 2: Oxidoreductase

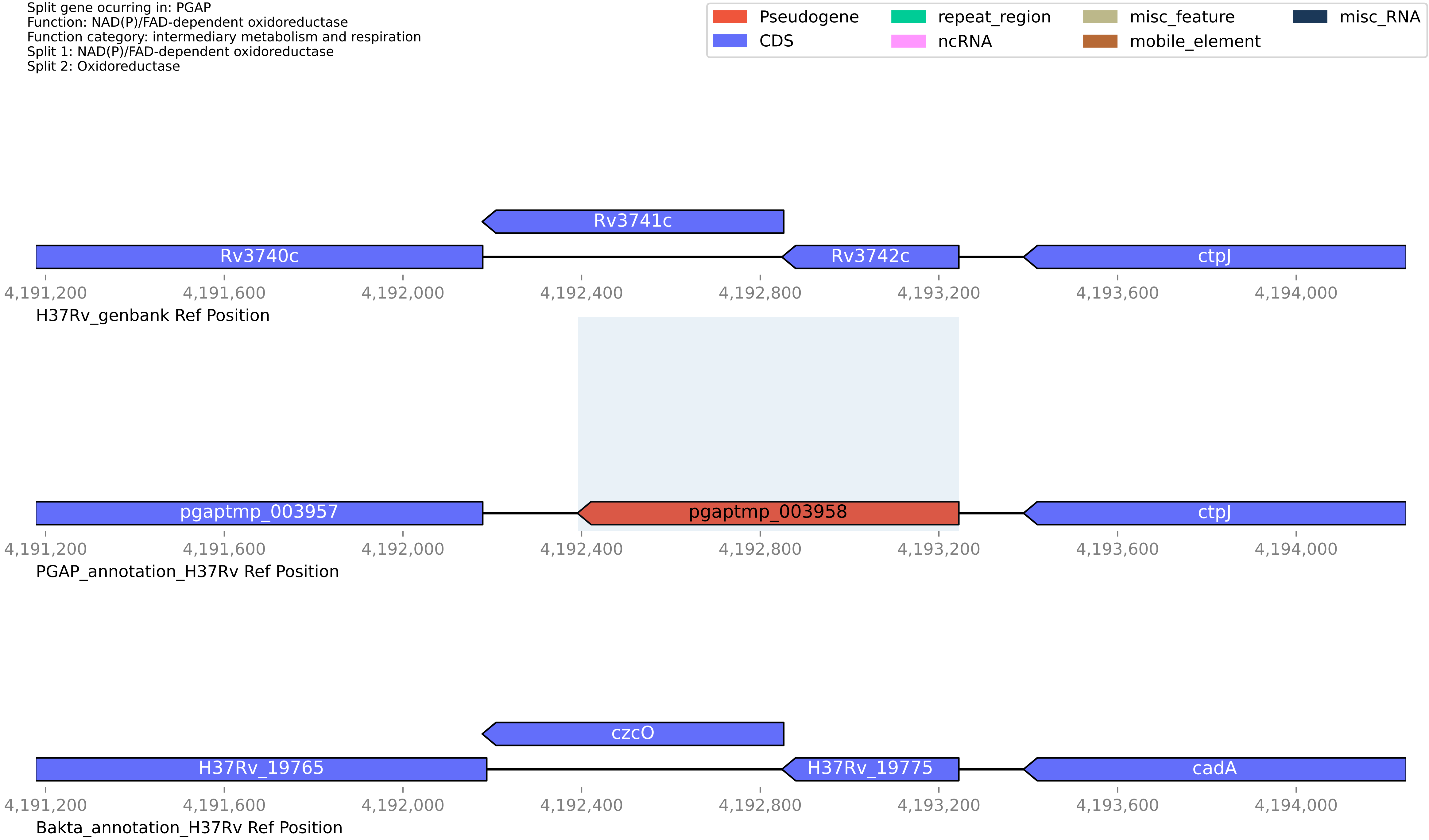

#### H37Rv PGAP or Bakta split gene annotation between coordinates 4215881-4216295, compared to Genbank

Split gene ocurring in: PGAP
Function: helix-turn-helix domain-containing protein
Function category: insertion seqs and phages
Split 1: hypothetical protein
Split 2: Transposase

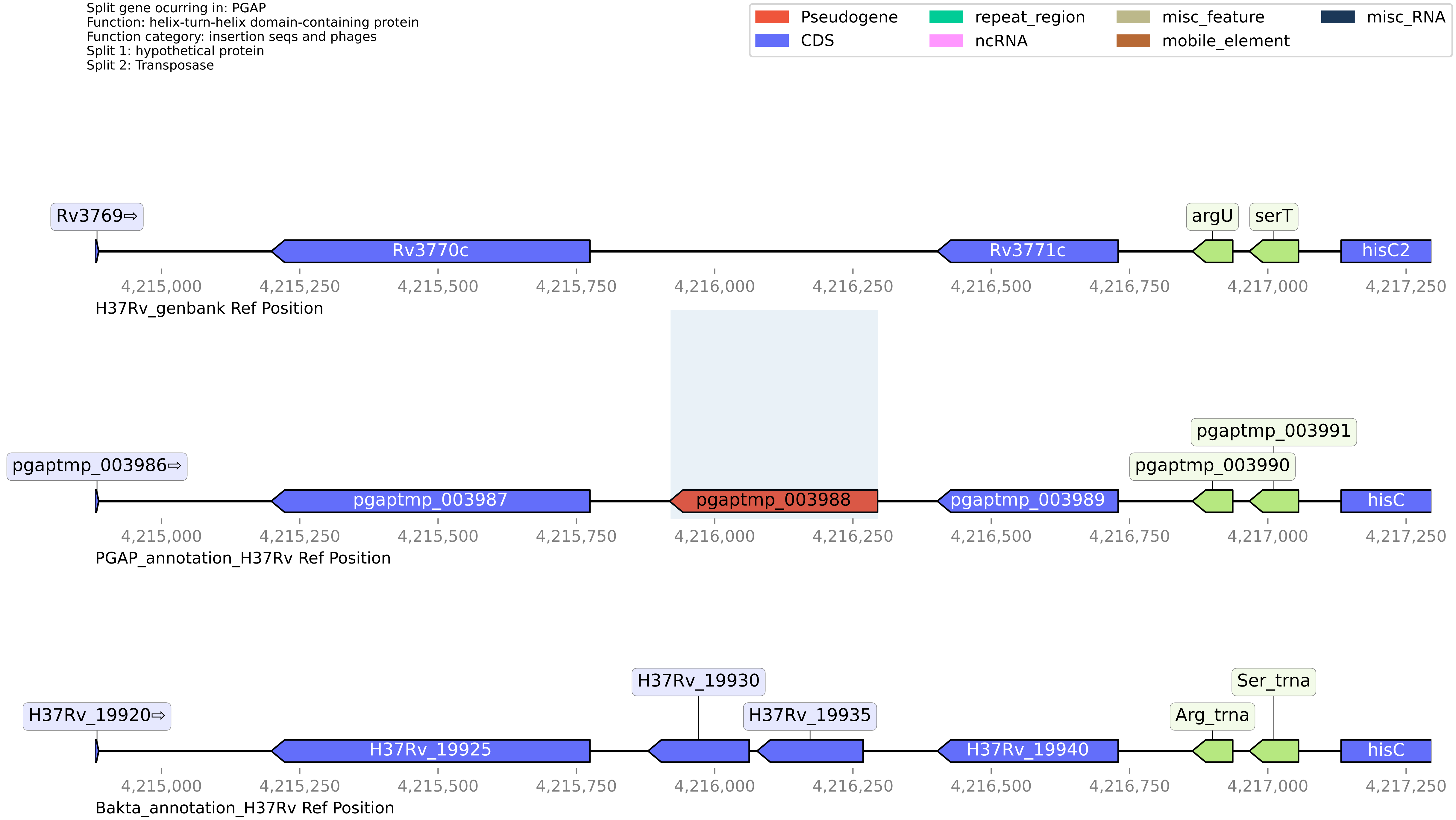

#### H37Rv PGAP or Bakta split gene annotation between coordinates 472890-474106, compared to Genbank

Split gene ocurring in: PGAP
Function: pseudogene
Function category: insertion seqs and phages
Split 1: 13E12 repeat family protein
Split 2: 13E12 repeat family protein

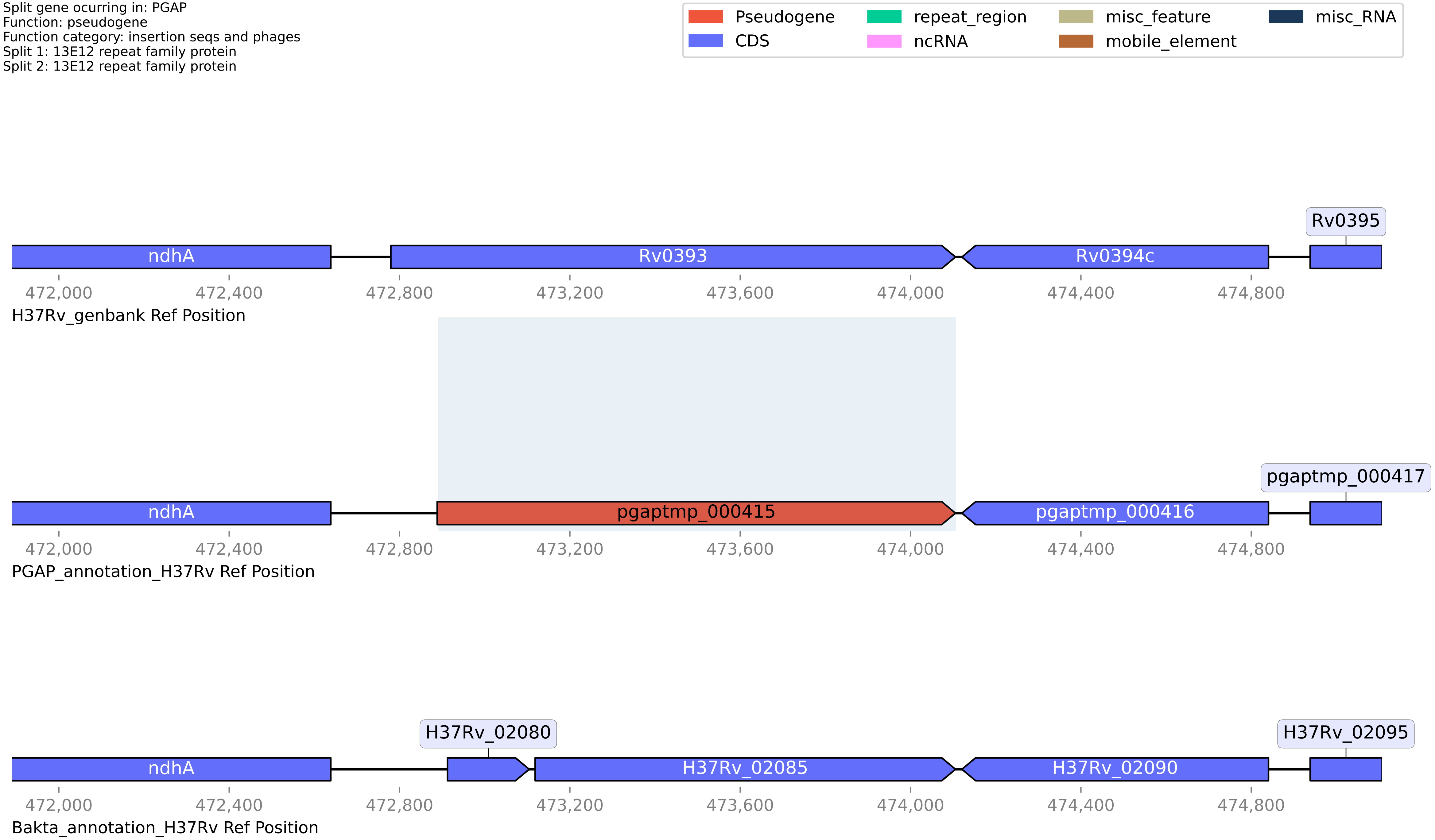

#### H37Rv PGAP or Bakta split gene annotation between coordinates 688032-689062, compared to Genbank

Split gene ocurring in: PGAP
Function: pseudogene
Function category: virulence
Split 1: Virulence factor mce family protein
Split 2: MCE-family protein

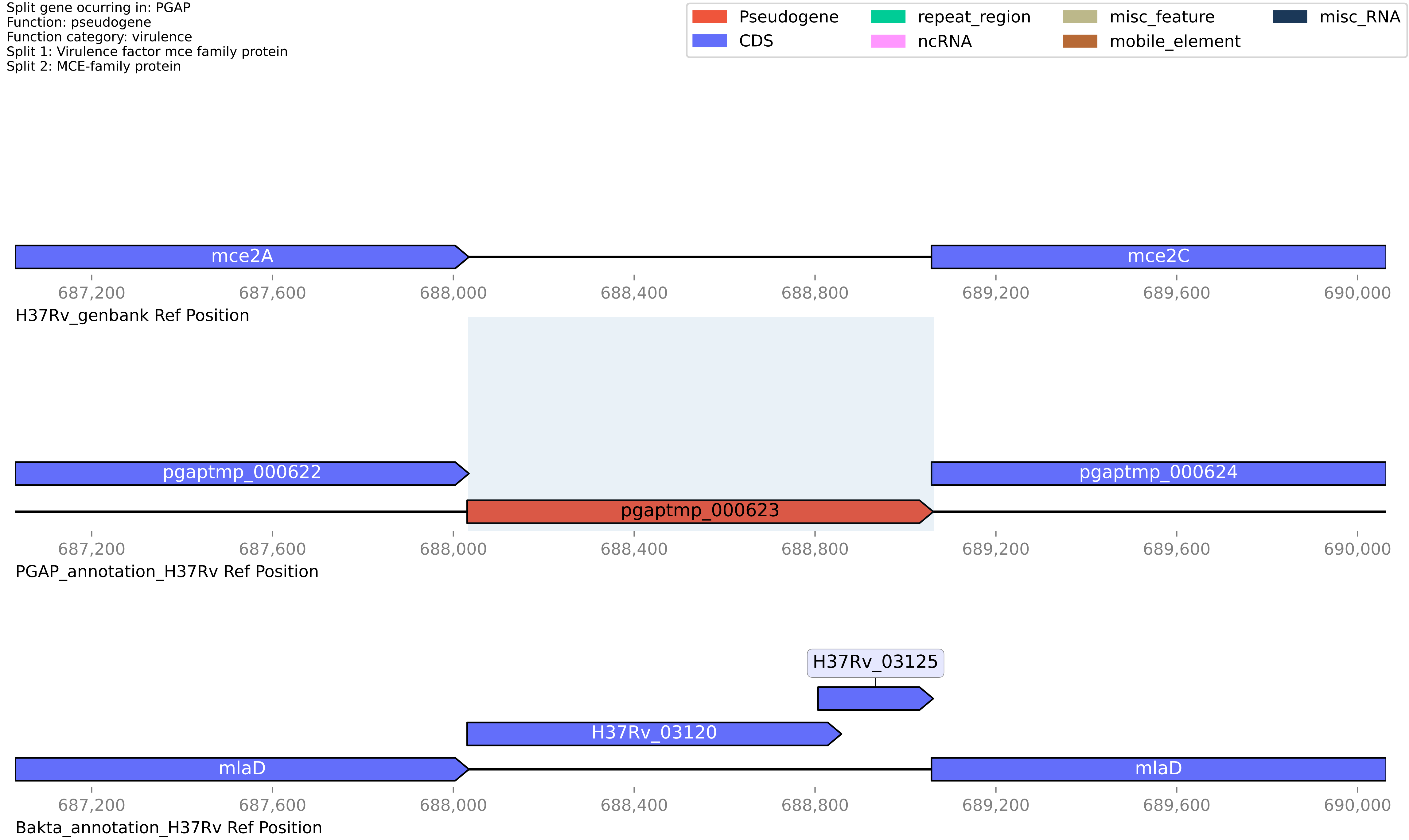

**H37Rv PGAP or Bakta split gene annotation between coordinates 711536-712719, compared to Genbank**

Split gene ocurring in: PGAP
Function: galT
Function category: intermediary metabolism and respiration
Split 1: galactose-1-phosphate uridylyltransferase
Split 2: Galactose-1-phosphate uridylyltransferase

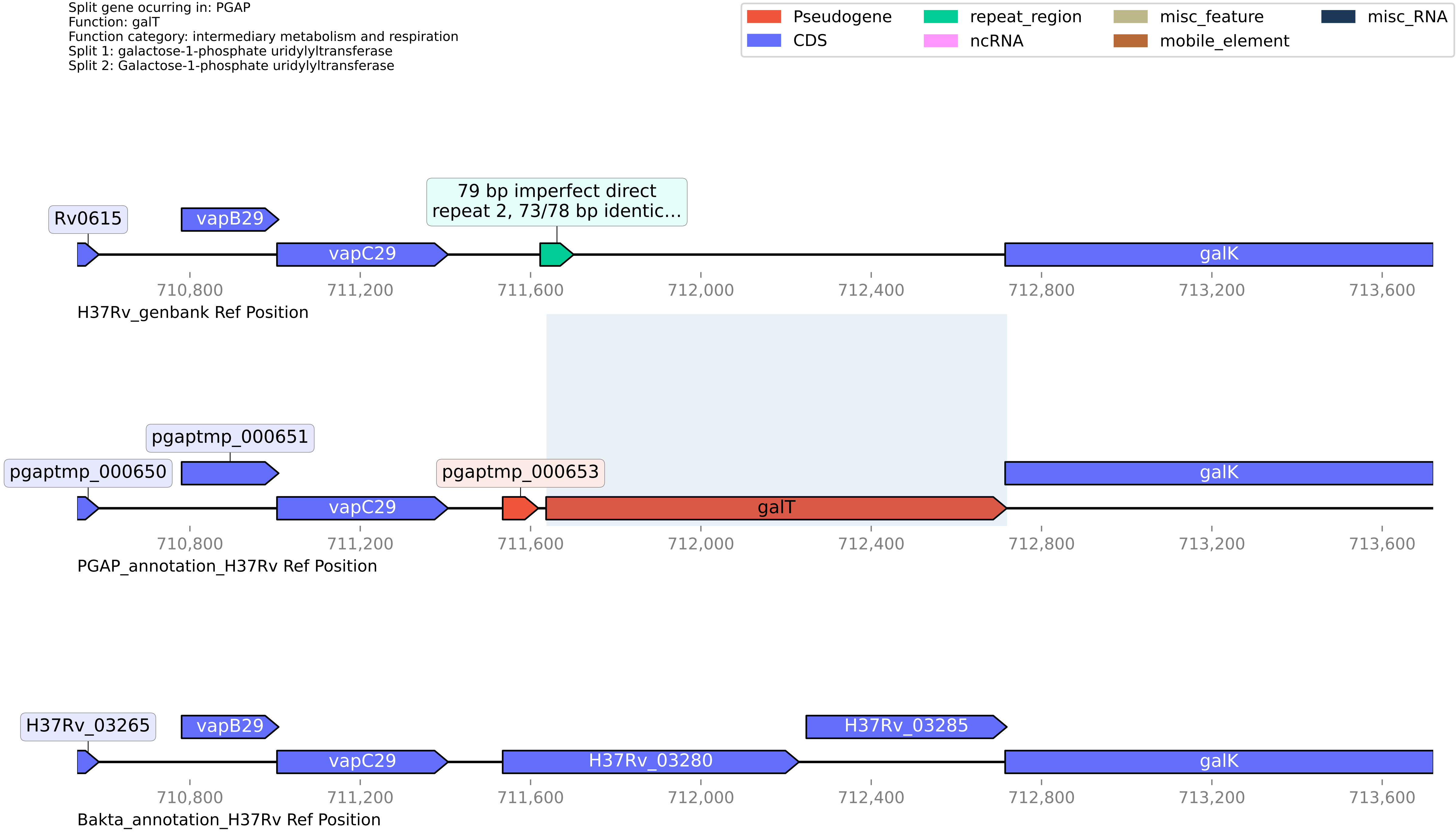

#### H37Rv PGAP or Bakta split gene annotation between coordinates 874233-876390, compared to Genbank

Split gene ocurring in: PGAP
Function: S9 family peptidase
Function category: intermediary metabolism and respiration
Split 1: putative protease II PtrBa [first part] (Oligopeptidase B)
Split 2: putative protease II PtrBb [second part] (Oligopeptidase B)

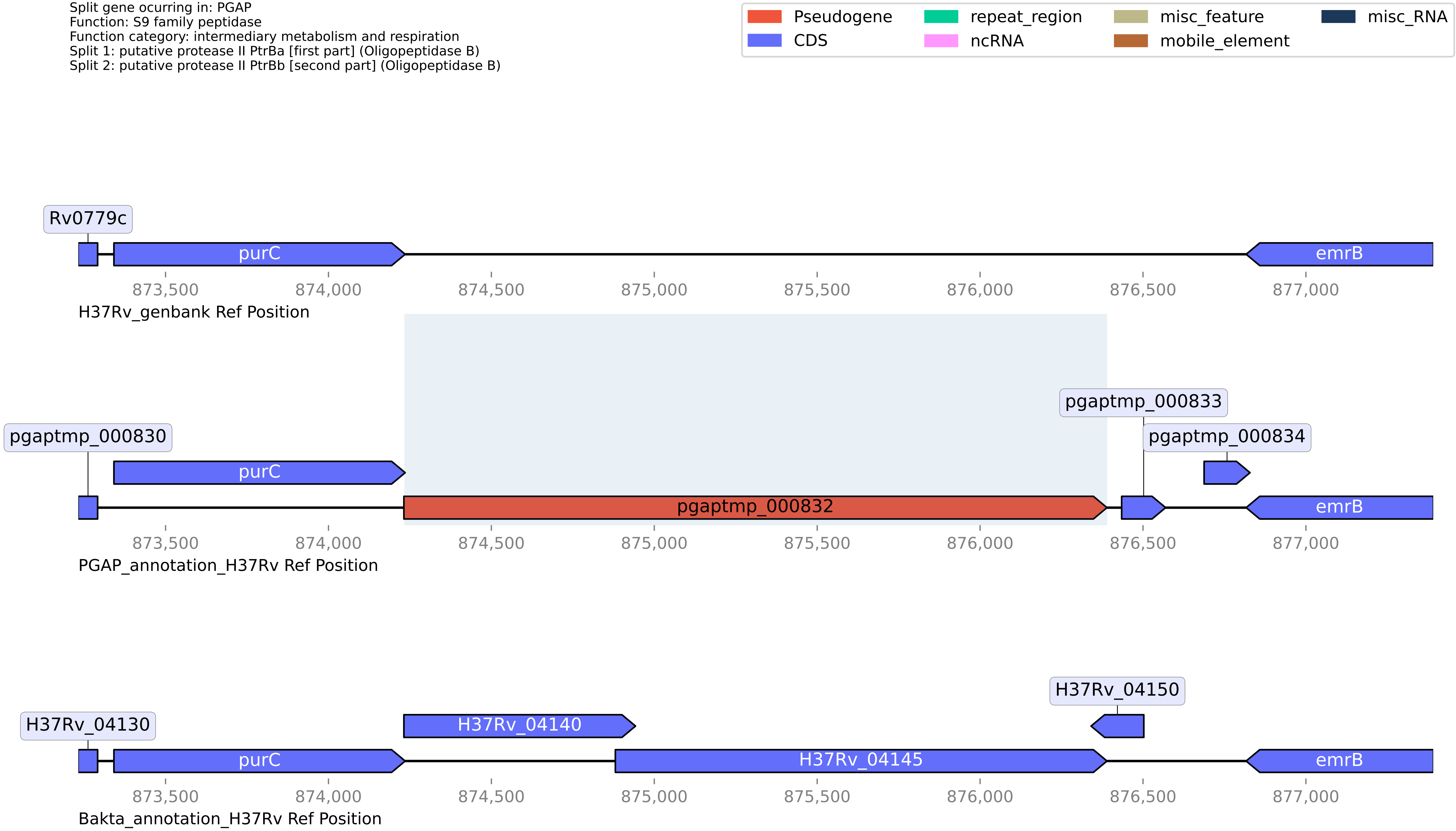

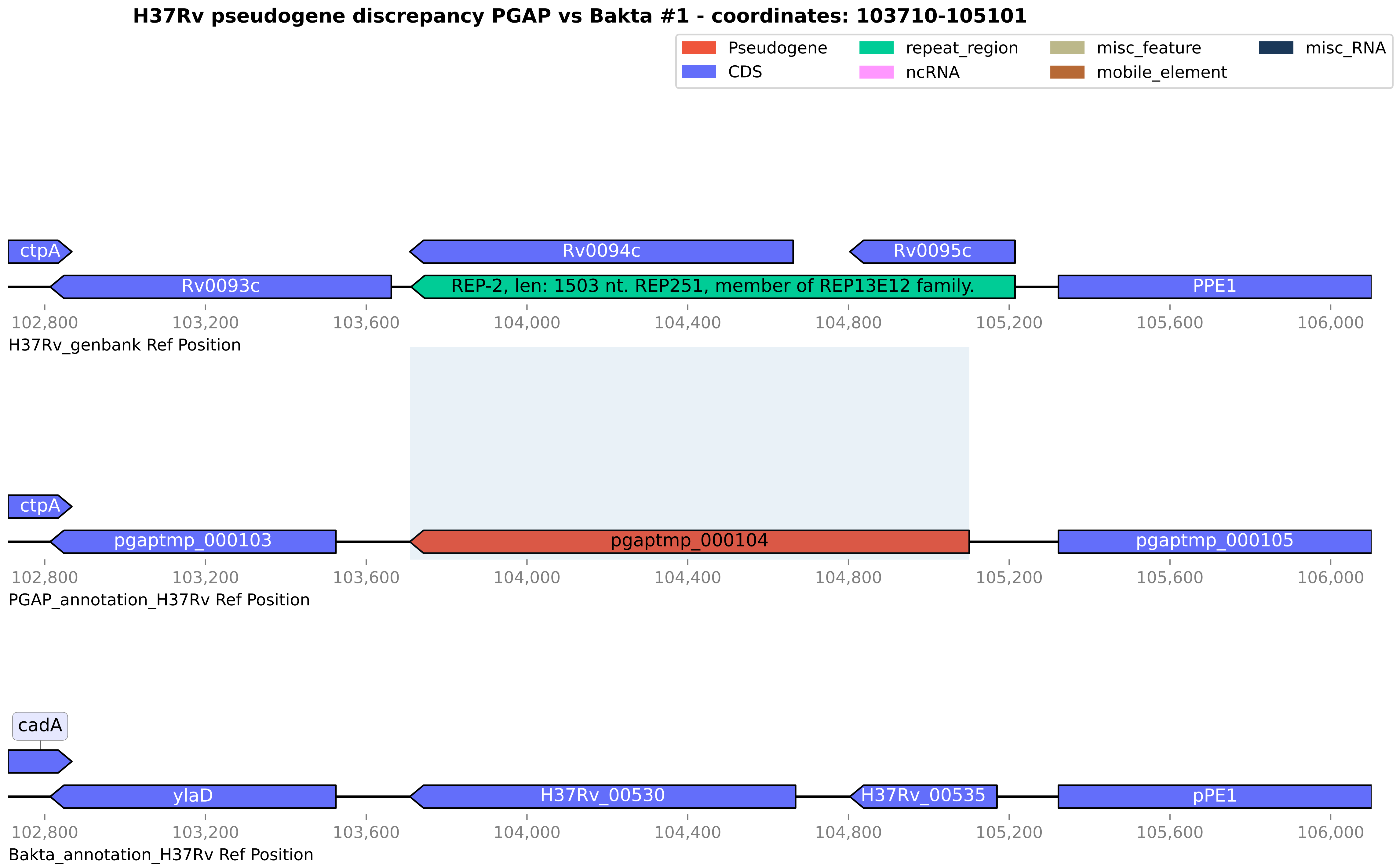

H37Rv pseudogene discrepancy PGAP vs Bakta #1 - coordinates: 103710-105101
Pseudogene
CDS
repeat_region
ncRNA
misc_feature
mobile_element
misc_RNA
ctpA
Rv0094c
Rv0095c
Rv0093c
REP-2, len: 1503 nt. REP251, member of REP13E12 family.
PPE1
102,800
103,200
103,600
104,000
104,400
104,800
105,200
105,600
106,000
H37Rv_genbank Ref Position
ctpA
pgaptmp_000103
pgaptmp_000104
pgaptmp_000105
PGAP_annotation_H37Rv Ref Position
cadA
ylaD
H37Rv_00530
H37Rv_00535
pPE1
Bakta_annotation_H37Rv Ref Position

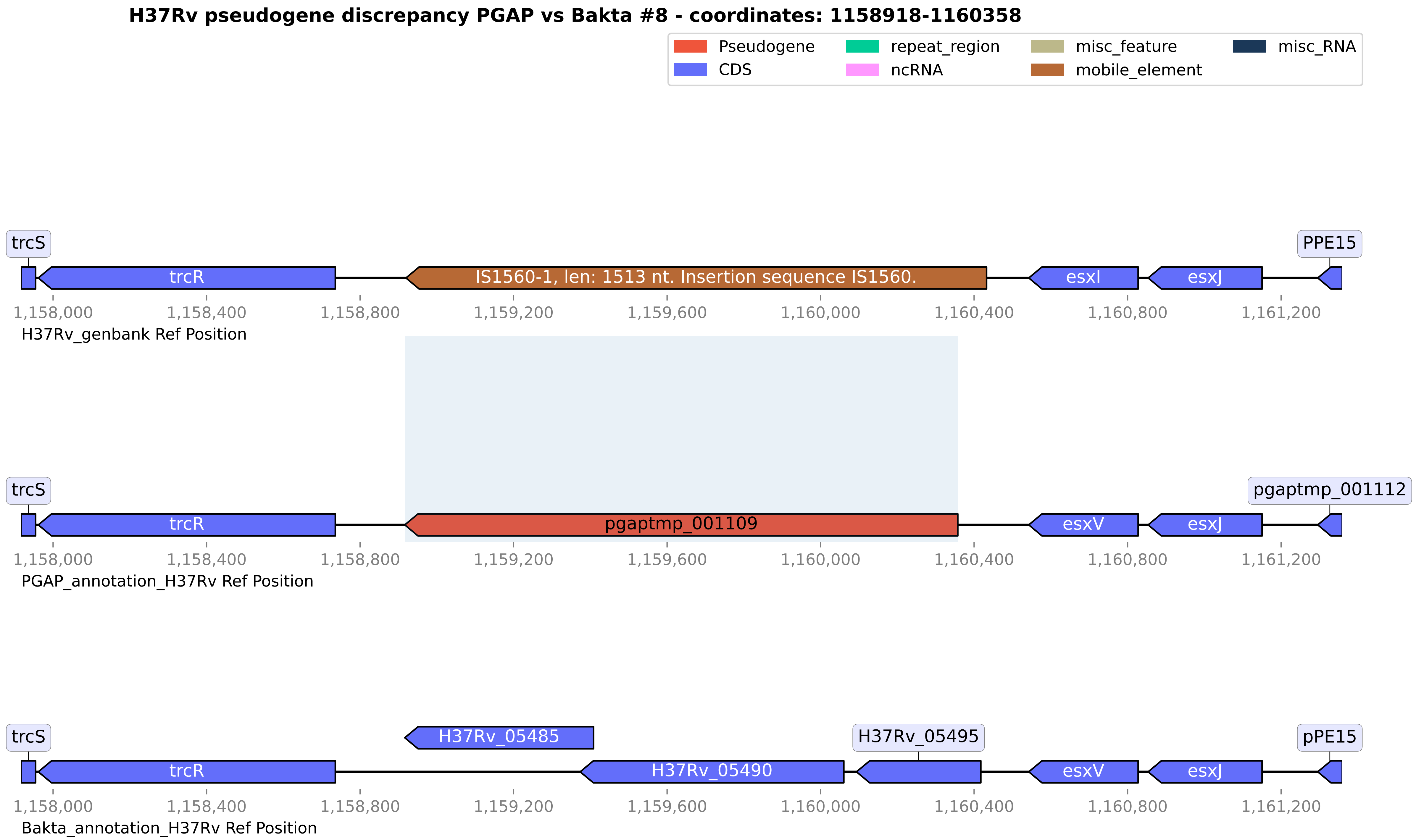
H37Rv pseudogene discrepancy PGAP vs Bakta #8 - coordinates: 1158918-1160358
Pseudogene
CDS
repeat_region
ncRNA
misc_feature
mobile_element
misc_RNA
trcS
trcR
IS1560-1, len: 1513 nt. Insertion sequence IS1560.
esxI
esxJ
PPE15
1,158,000
1,158,400
1,158,800
1,159,200
1,159,600
1,160,000
1,160,400
1,160,800
1,161,200
H37Rv_genbank Ref Position
trcS
trcR
pgaptmp_001109
esxV
esxJ
pgaptmp_001112
PGAP_annotation_H37Rv Ref Position
trcS
trcR
H37Rv_05485
H37Rv_05490
H37Rv_05495
esxV
esxJ
pPE15
Bakta_annotation_H37Rv Ref Position

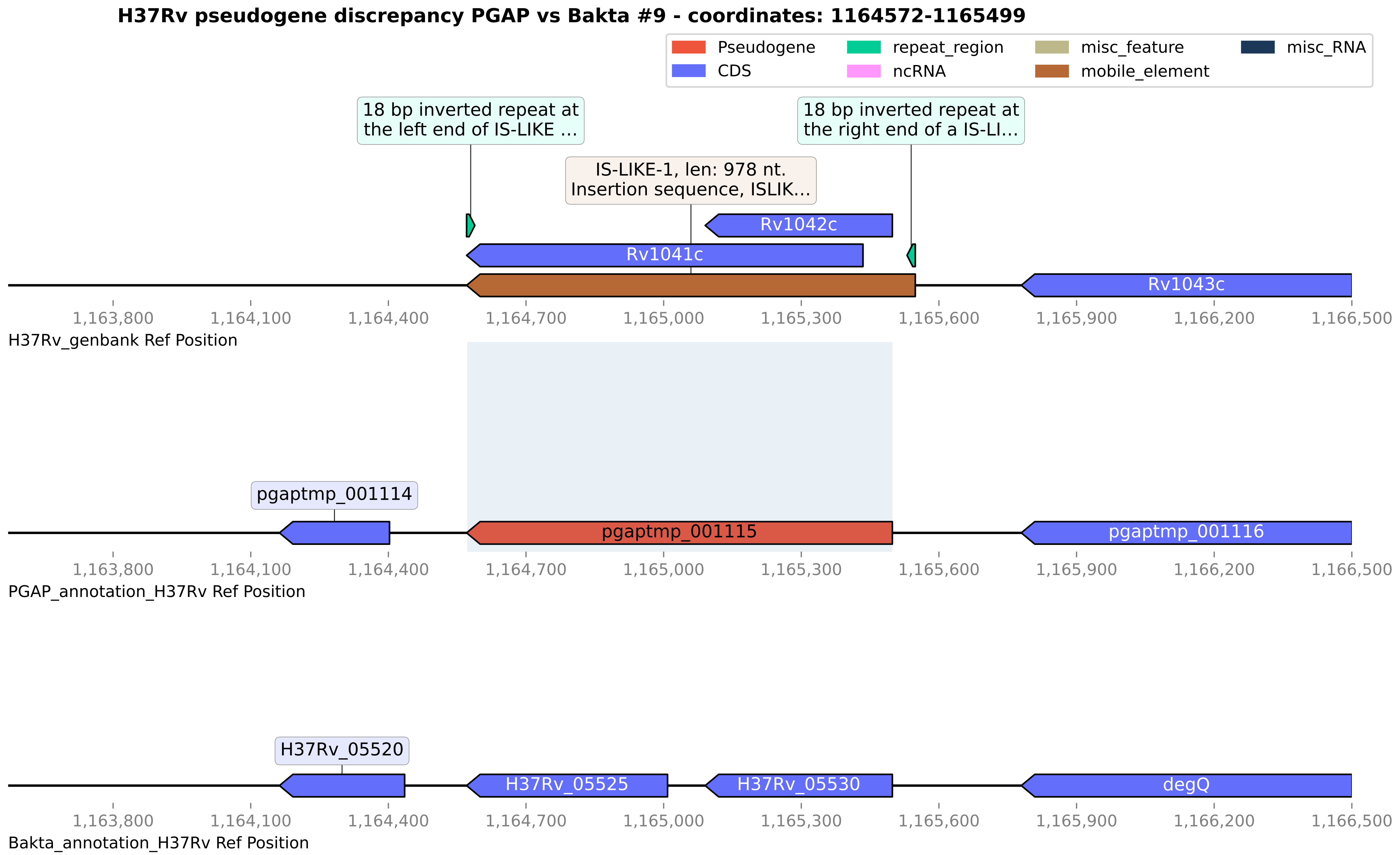

H37Rv pseudogene discrepancy PGAP vs Bakta #9 - coordinates: 1164572-1165499
Pseudogene
CDS
repeat_region
ncRNA
misc_feature
mobile_element
misc_RNA
18 bp inverted repeat at the left end of IS-LIKE ...
18 bp inverted repeat at the right end of a IS-LI...
IS-LIKE-1, len: 978 nt. Insertion sequence, ISLIK...
Rv1042c
Rv1041c
Rv1043c
1,163,800
1,164,100
1,164,400
1,164,700
1,165,000
1,165,300
1,165,600
1,165,900
1,166,200
1,166,500
H37Rv_genbank Ref Position
pgaptmp_001114
pgaptmp_001115
pgaptmp_001116
PGAP_annotation_H37Rv Ref Position
H37Rv_05520
H37Rv_05525
H37Rv_05530
degQ
Bakta_annotation_H37Rv Ref Position

**H37Rv pseudogene discrepancy PGAP vs Bakta #10 - coordinates: 1173945-1174700**

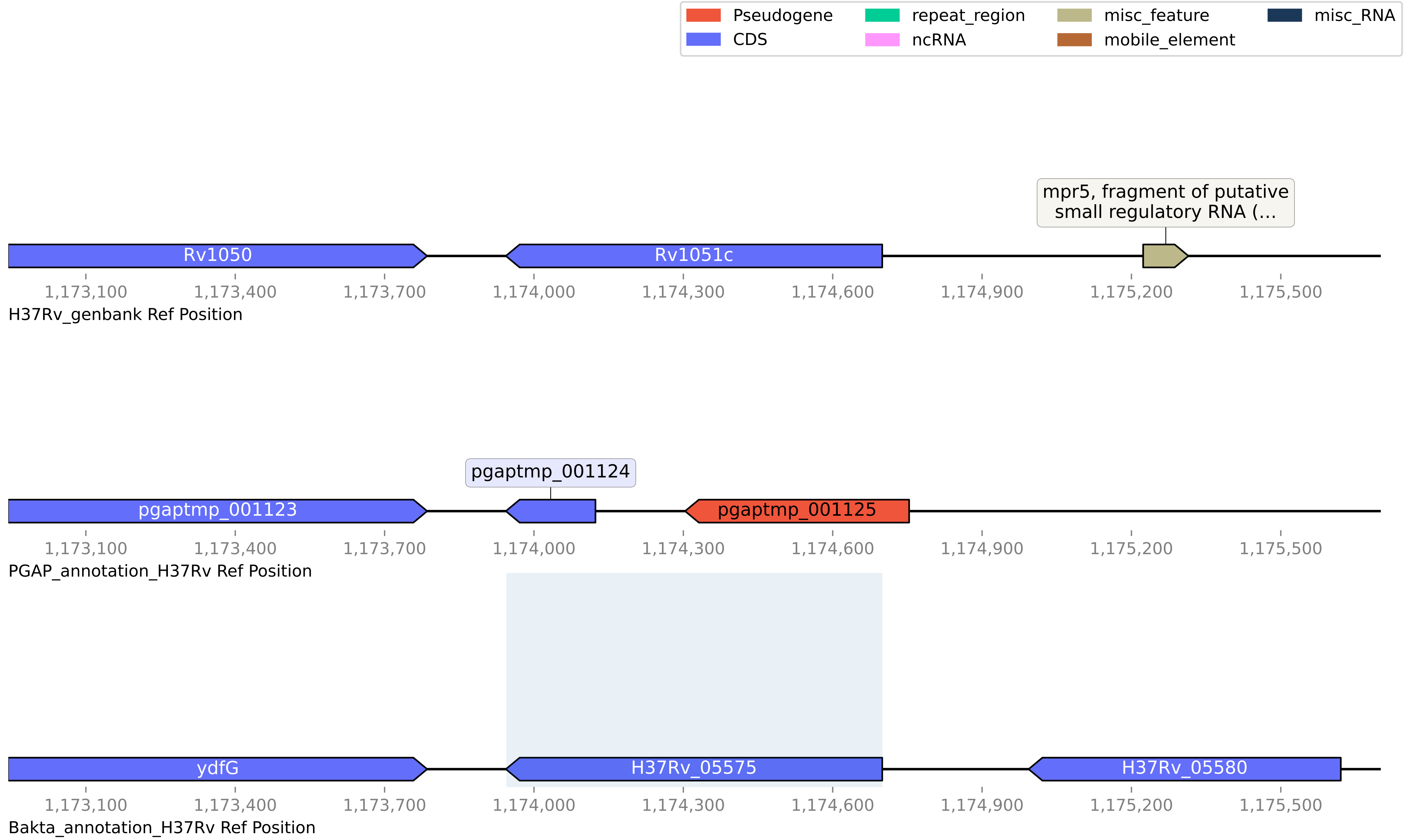

#### H37Rv pseudogene discrepancy PGAP vs Bakta #11 - coordinates: 1231301-1232837

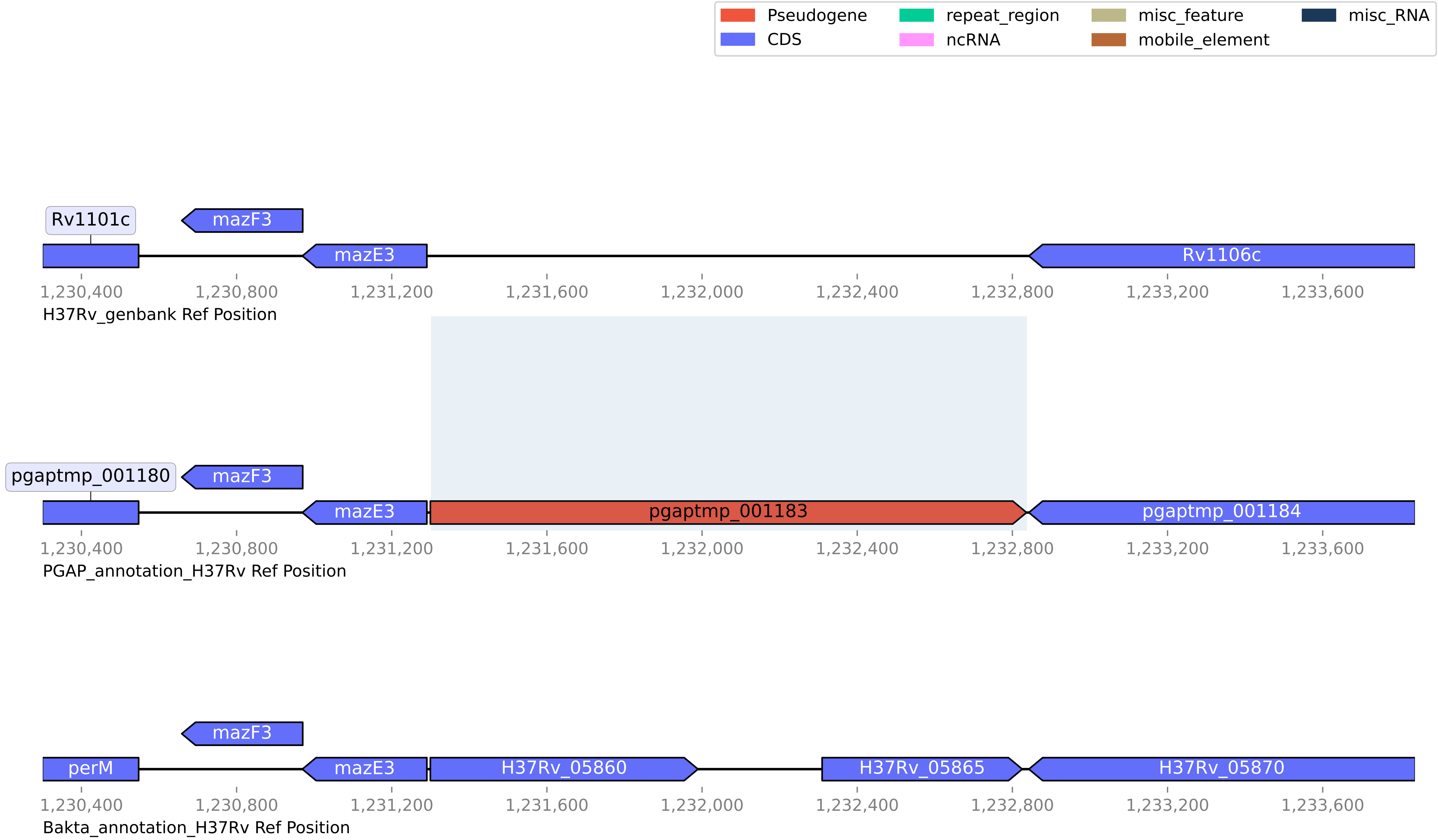

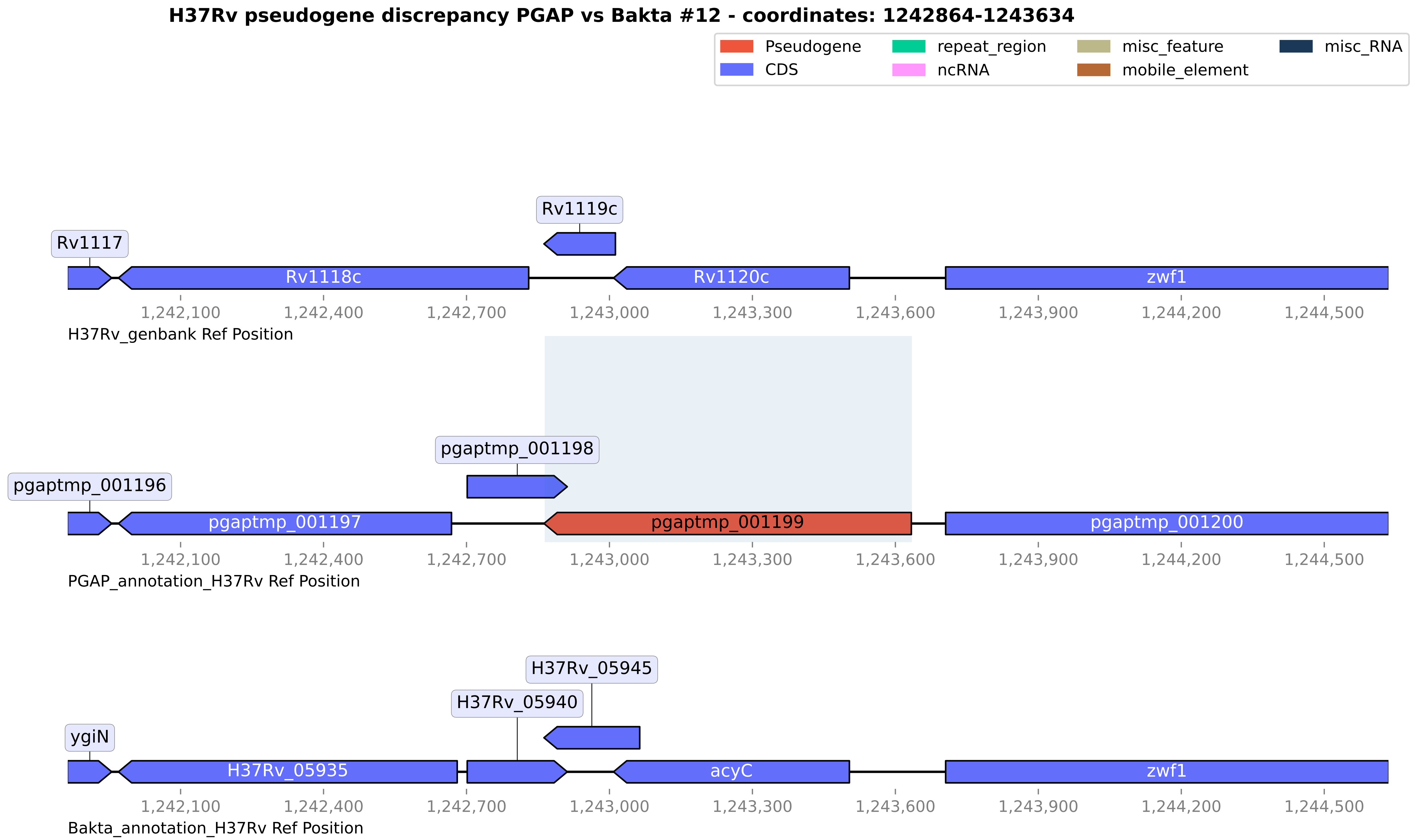
H37Rv pseudogene discrepancy PGAP vs Bakta #12 - coordinates: 1242864-1243634
Pseudogene
CDS
repeat_region
ncRNA
misc_feature
mobile_element
misc_RNA
Rv1119c
Rv1117
Rv1118c
Rv1120c
zwf1
1,242,100
1,242,400
1,242,700
1,243,000
1,243,300
1,243,600
1,243,900
1,244,200
1,244,500
H37Rv_genbank Ref Position
pgaptmp_001198
pgaptmp_001196
pgaptmp_001197
pgaptmp_001199
pgaptmp_001200
PGAP_annotation_H37Rv Ref Position
H37Rv_05945
H37Rv_05940
ygiN
H37Rv_05935
acyC
zwf1
Bakta_annotation_H37Rv Ref Position

#### H37Rv pseudogene discrepancy PGAP vs Bakta #13 - coordinates: 1272423-1274767

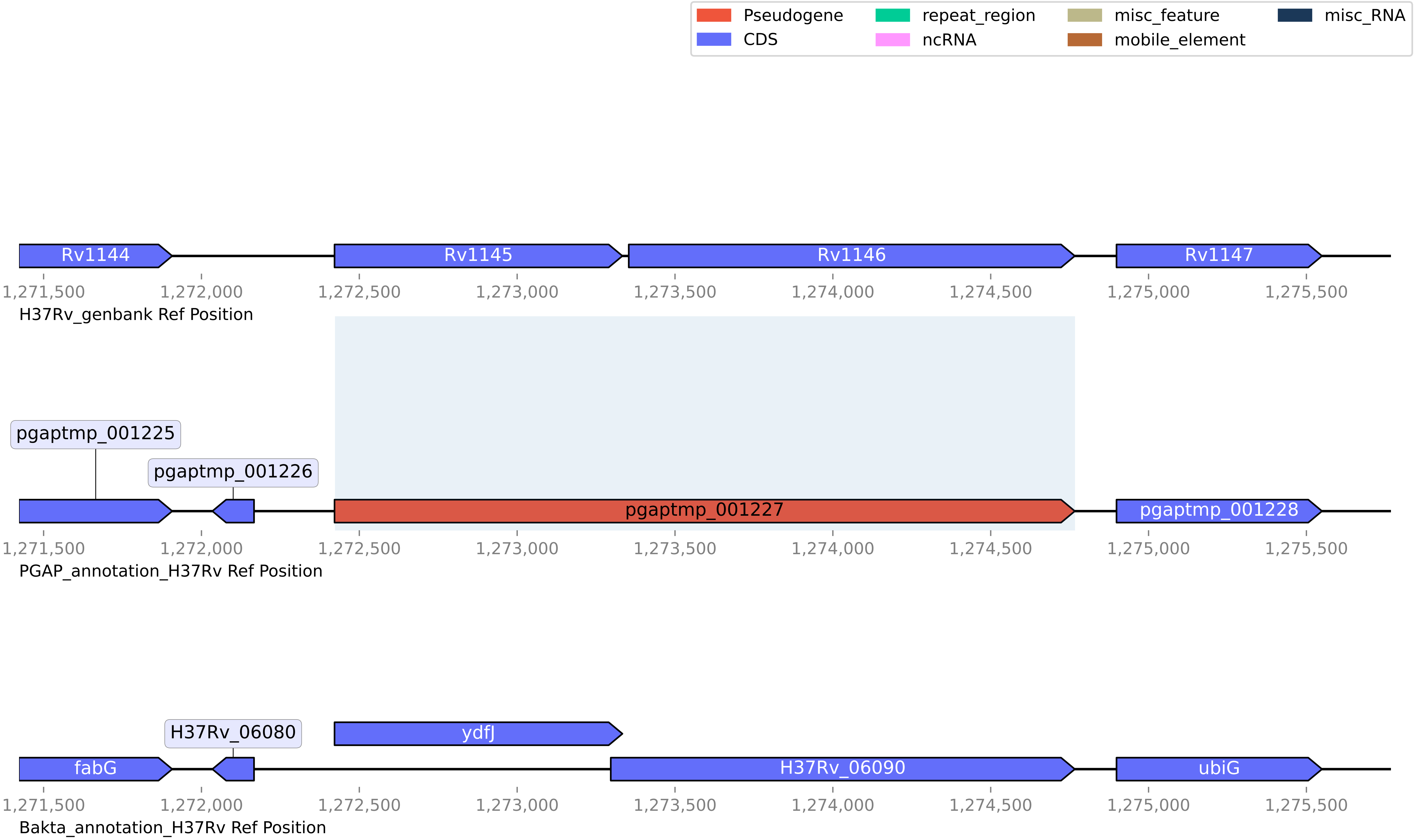

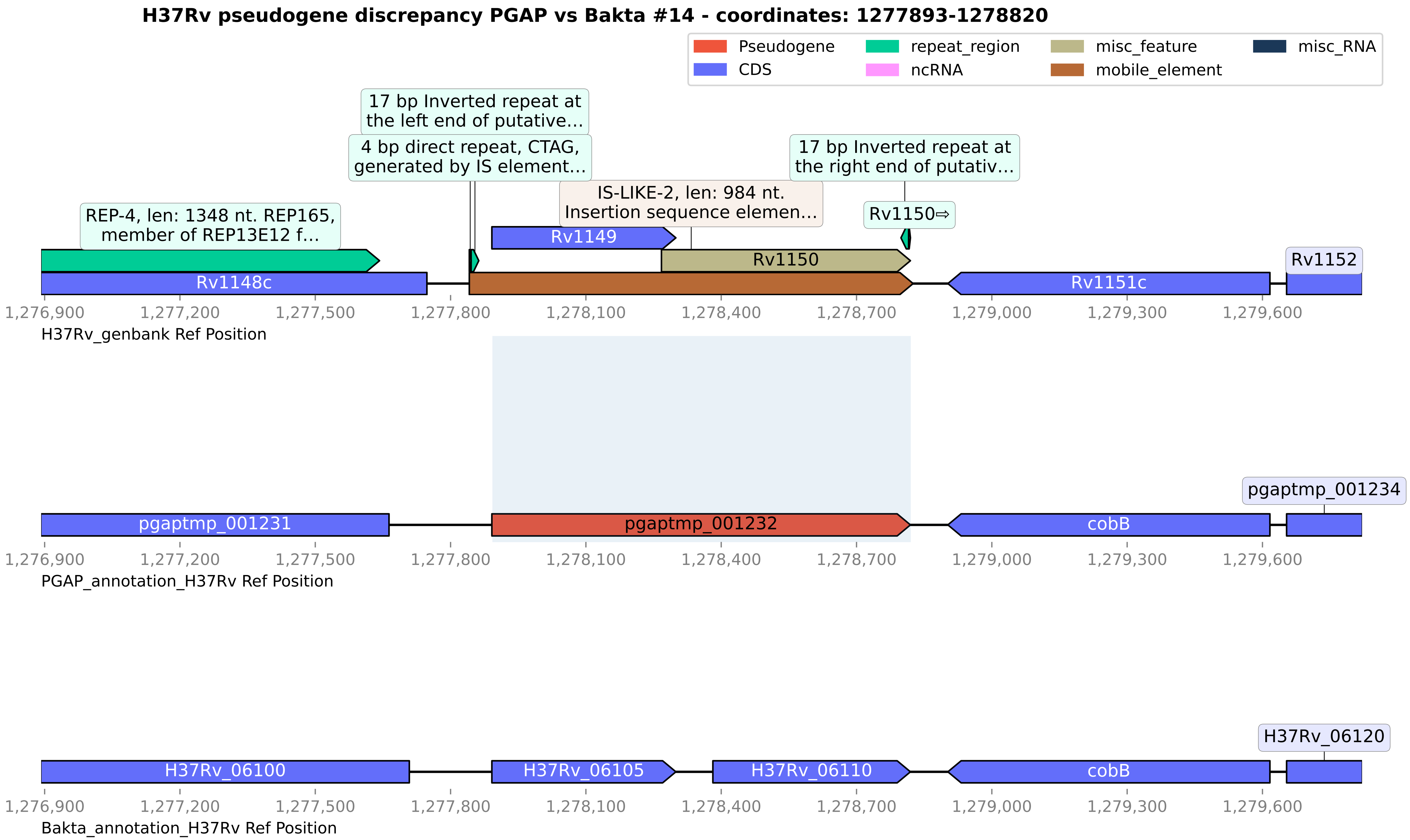
H37Rv pseudogene discrepancy PGAP vs Bakta #14 - coordinates: 1277893-1278820
Pseudogene
CDS
repeat_region
ncRNA
misc_feature
mobile_element
misc_RNA
17 bp Inverted repeat at the left end of putative...
4 bp direct repeat, CTAG, generated by IS element...
17 bp Inverted repeat at the right end of putativ...
IS-LIKE-2, len: 984 nt. Insertion sequence elemen...
REP-4, len: 1348 nt. REP165, member of REP13E12 f...
Rv1150⇨
Rv1149
Rv1150
Rv1148c
Rv1151c
Rv1152
1,276,900
1,277,200
1,277,500
1,277,800
1,278,100
1,278,400
1,278,700
1,279,000
1,279,300
1,279,600
H37Rv_genbank Ref Position
pgaptmp_001231
pgaptmp_001232
cobB
pgaptmp_001234
PGAP_annotation_H37Rv Ref Position
H37Rv_06100
H37Rv_06105
H37Rv_06110
cobB
H37Rv_06120
Bakta_annotation_H37Rv Ref Position

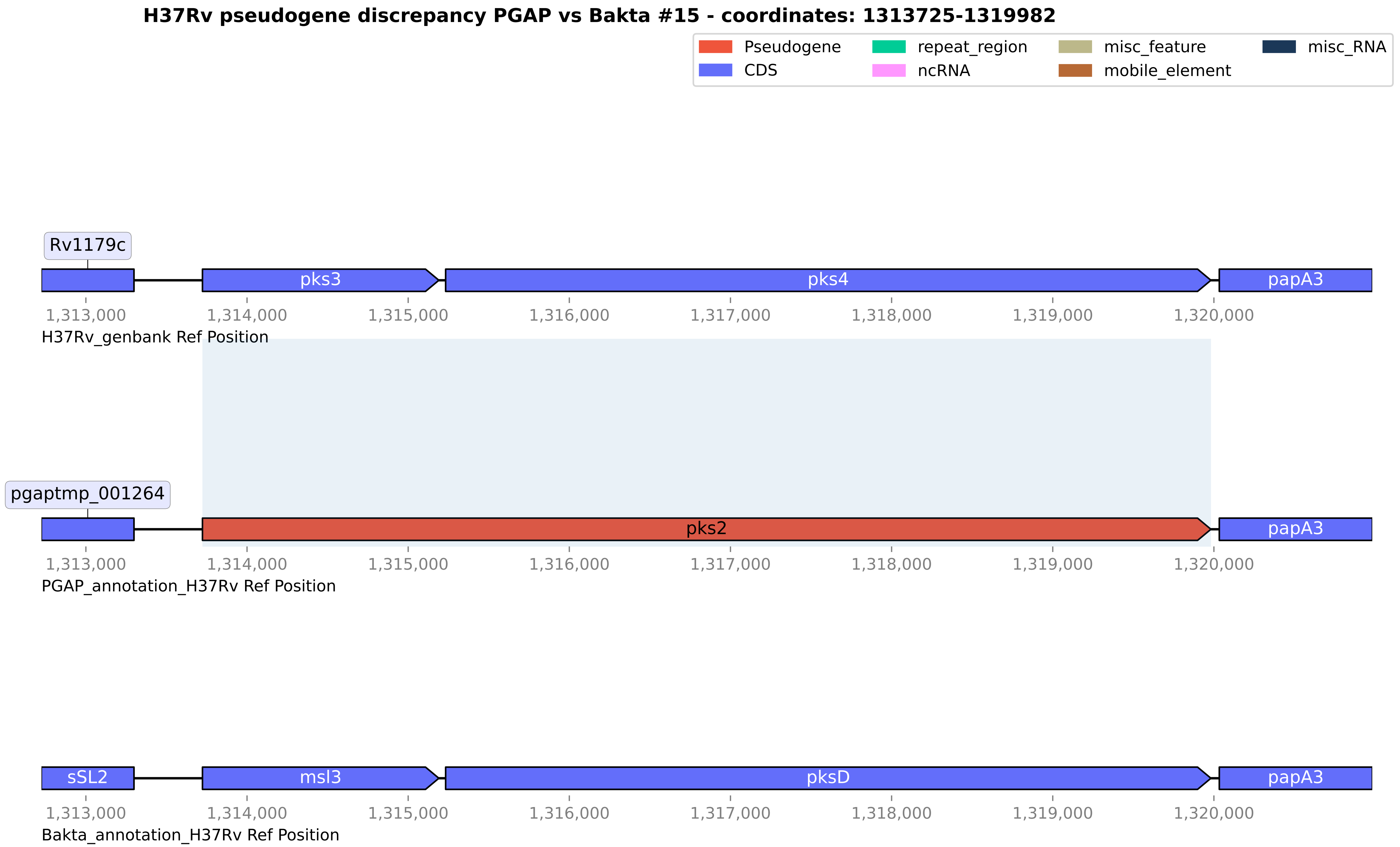

H37Rv pseudogene discrepancy PGAP vs Bakta #15 - coordinates: 1313725-1319982
Pseudogene
CDS
repeat_region
ncRNA
misc_feature
mobile_element
misc_RNA
Rv1179c
pks3
pks4
papA3
1,313,000
1,314,000
1,315,000
1,316,000
1,317,000
1,318,000
1,319,000
1,320,000
H37Rv_genbank Ref Position
pgaptmp_001264
pks2
papA3
1,313,000
1,314,000
1,315,000
1,316,000
1,317,000
1,318,000
1,319,000
1,320,000
PGAP_annotation_H37Rv Ref Position
sSL2
msl3
pksD
papA3
1,313,000
1,314,000
1,315,000
1,316,000
1,317,000
1,318,000
1,319,000
1,320,000
Bakta_annotation_H37Rv Ref Position

H37Rv pseudogene discrepancy PGAP vs Bakta #16 - coordinates: 1589199-1590292

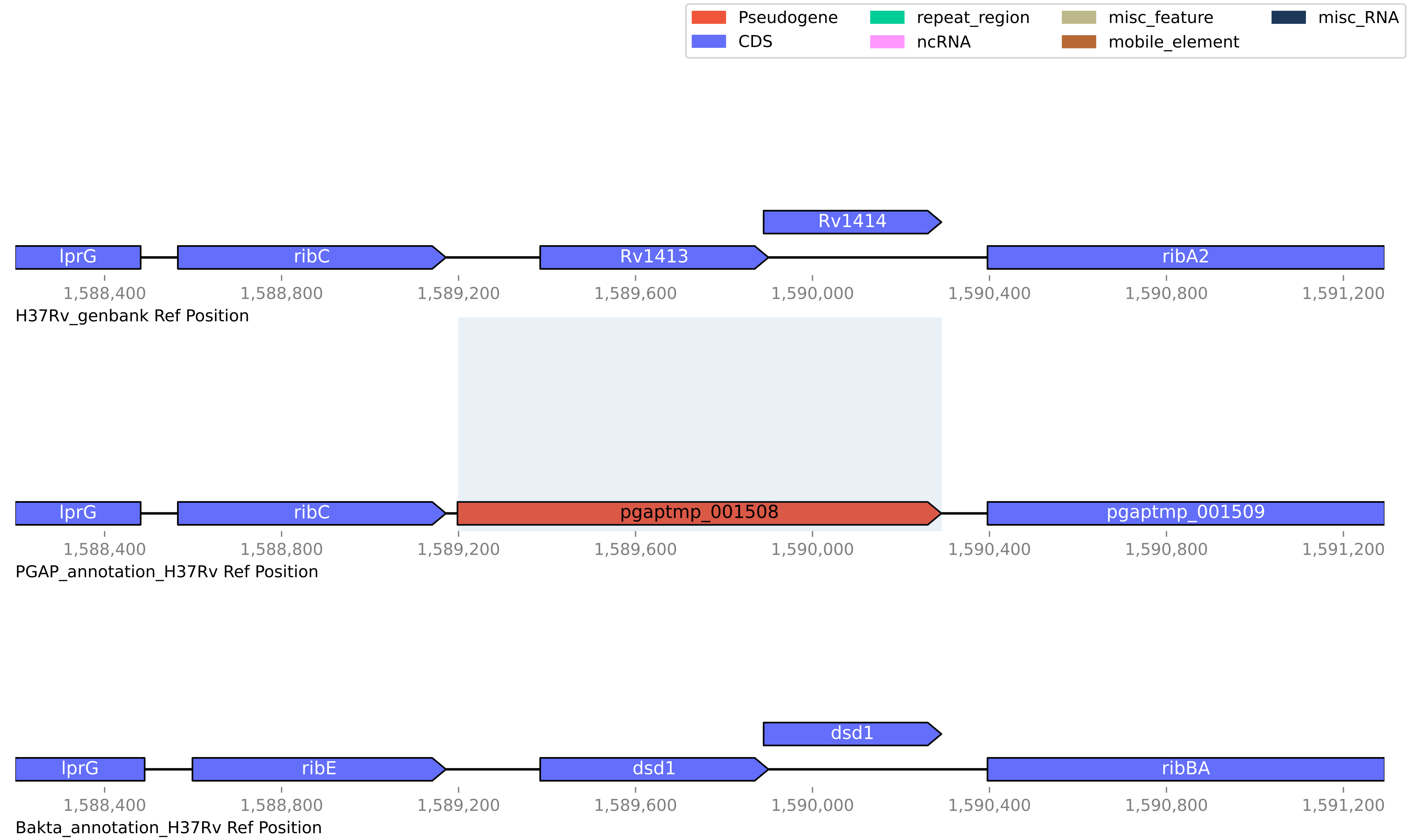

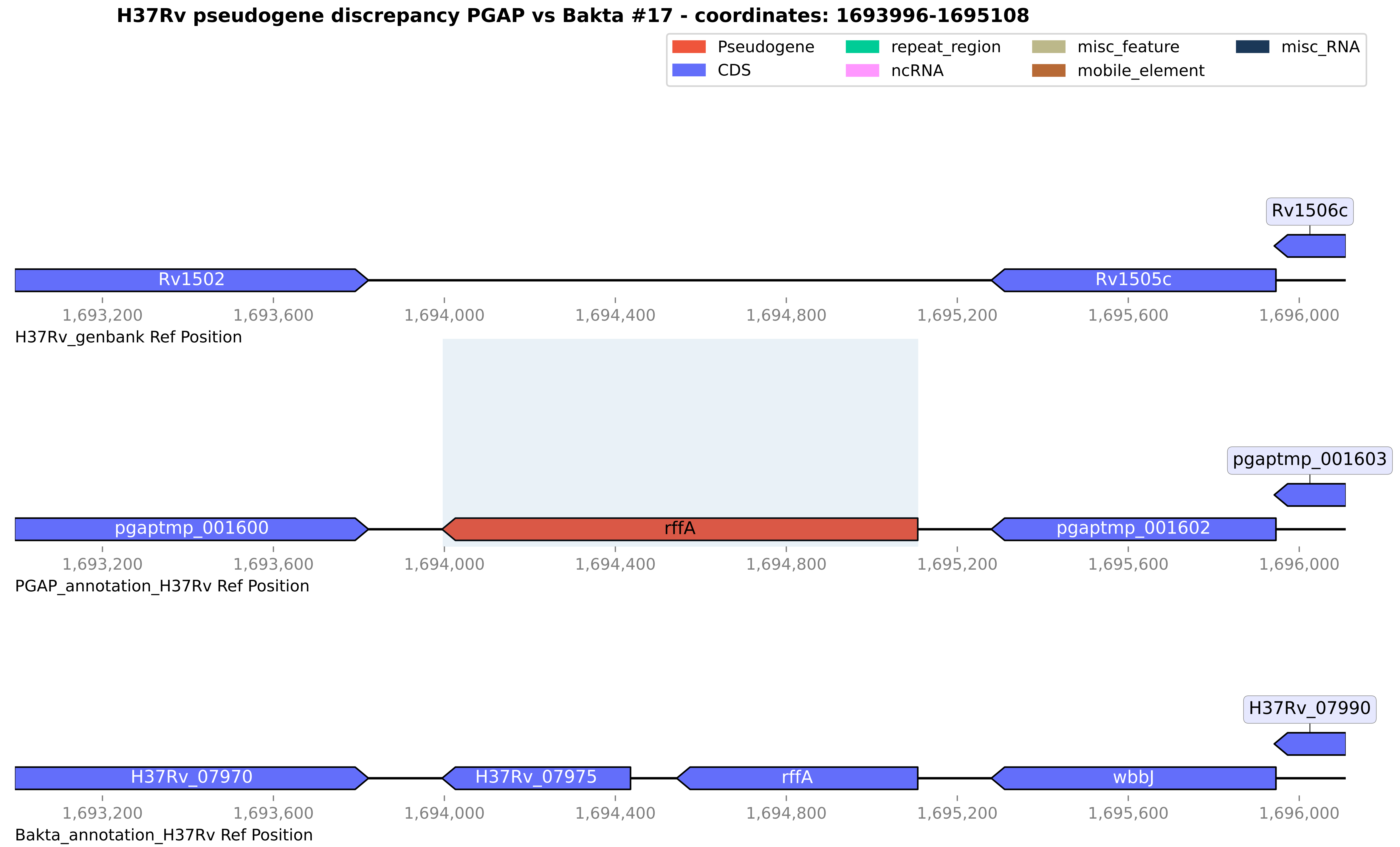

H37Rv pseudogene discrepancy PGAP vs Bakta #17 - coordinates: 1693996-1695108
Pseudogene
CDS
repeat_region
ncRNA
misc_feature
mobile_element
misc_RNA
Rv1506c
Rv1502
Rv1505c
1,693,200
1,693,600
1,694,000
1,694,400
1,694,800
1,695,200
1,695,600
1,696,000
H37Rv_genbank Ref Position
pgaptmp_001603
pgaptmp_001600
rffA
pgaptmp_001602
1,693,200
1,693,600
1,694,000
1,694,400
1,694,800
1,695,200
1,695,600
1,696,000
PGAP_annotation_H37Rv Ref Position
H37Rv_07990
H37Rv_07970
H37Rv_07975
rffA
wbbJ
1,693,200
1,693,600
1,694,000
1,694,400
1,694,800
1,695,200
1,695,600
1,696,000
Bakta_annotation_H37Rv Ref Position

#### H37Rv pseudogene discrepancy PGAP vs Bakta #18 - coordinates: 1753606-1755431

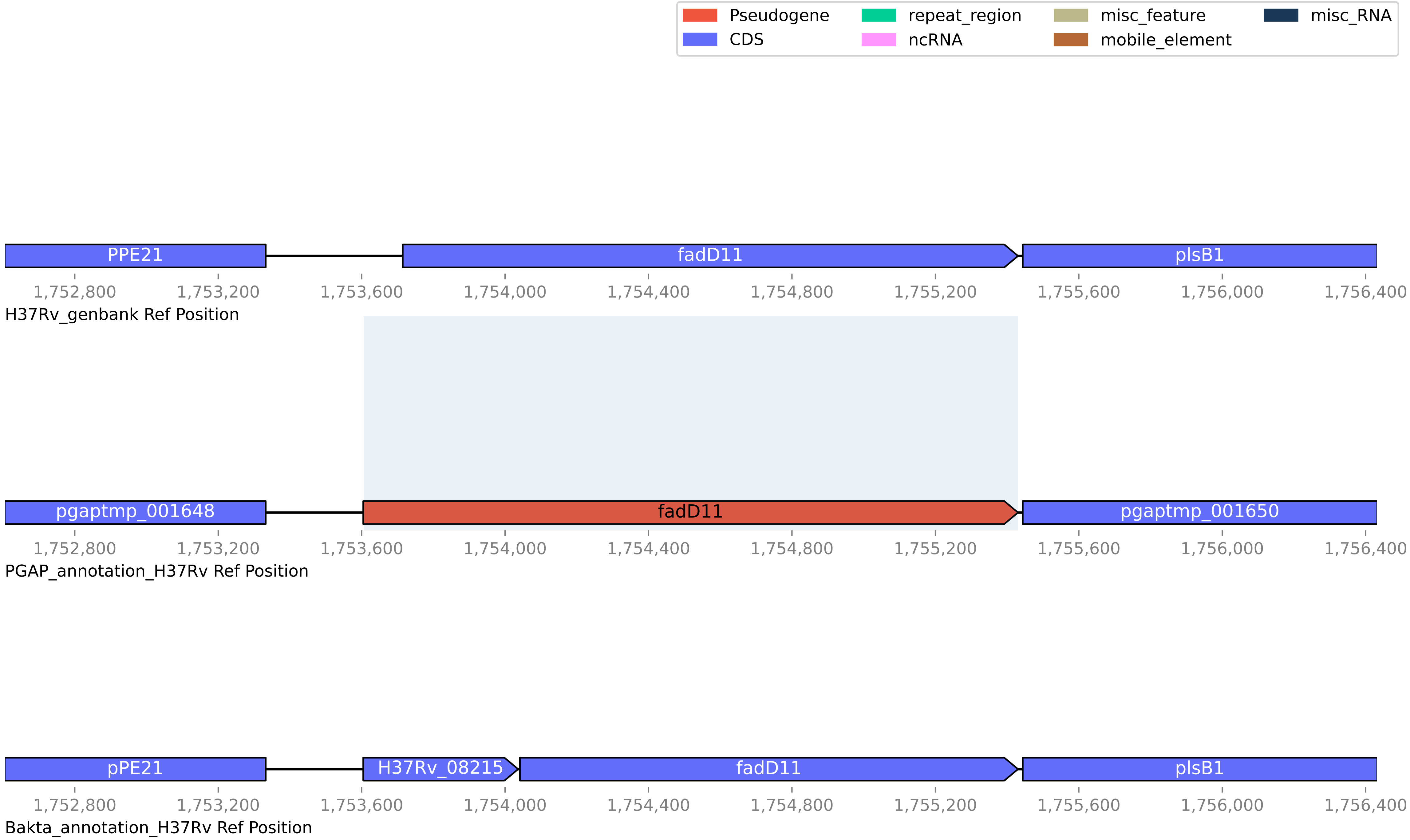

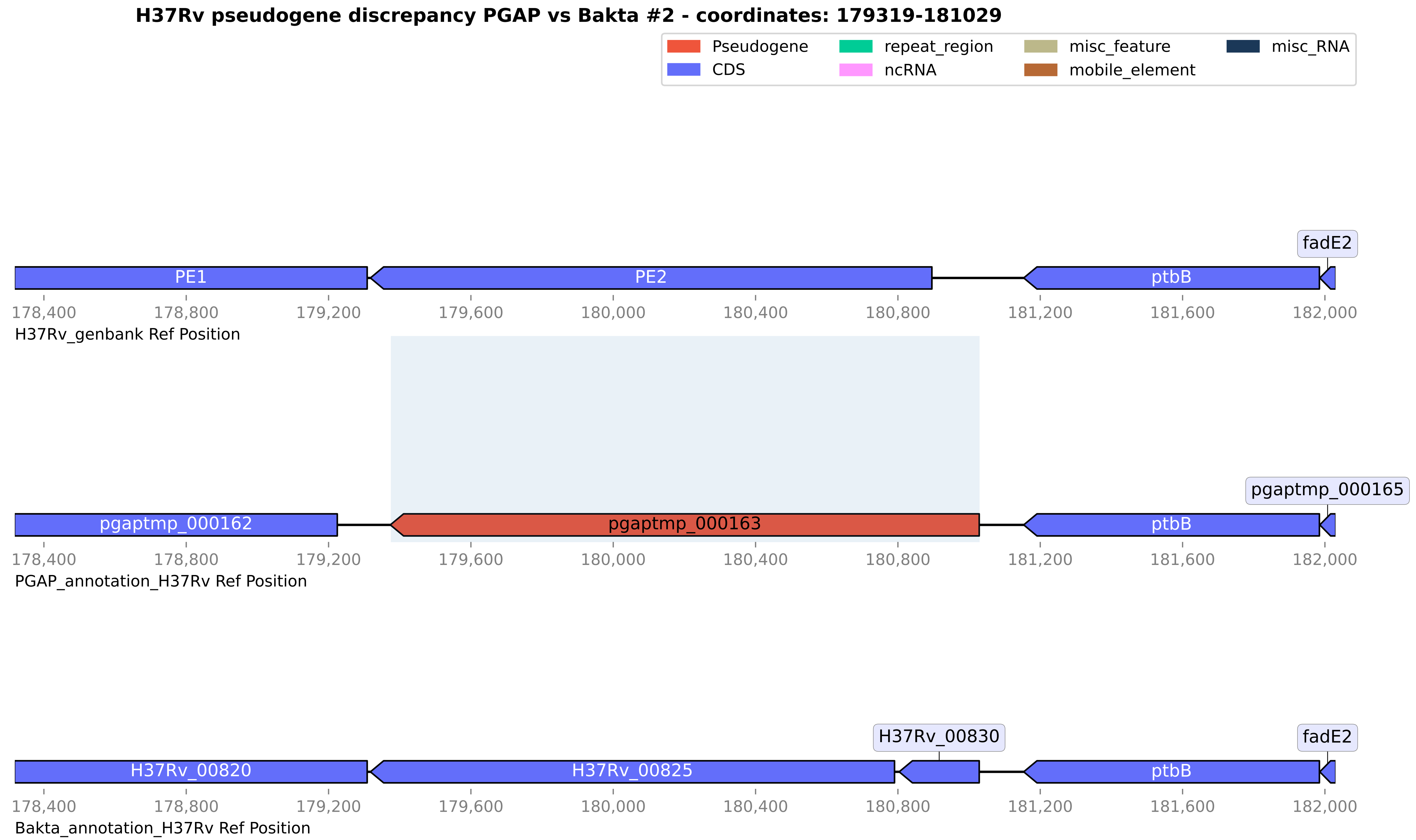
H37Rv pseudogene discrepancy PGAP vs Bakta #2 - coordinates: 179319-181029
Pseudogene
CDS
repeat_region
ncRNA
misc_feature
mobile_element
misc_RNA
fadE2
PE1
PE2
ptbB
178,400
178,800
179,200
179,600
180,000
180,400
180,800
181,200
181,600
182,000
H37Rv_genbank Ref Position
pgaptmp_000165
pgaptmp_000162
pgaptmp_000163
ptbB
PGAP_annotation_H37Rv Ref Position
H37Rv_00830
fadE2
H37Rv_00820
H37Rv_00825
ptbB
Bakta_annotation_H37Rv Ref Position

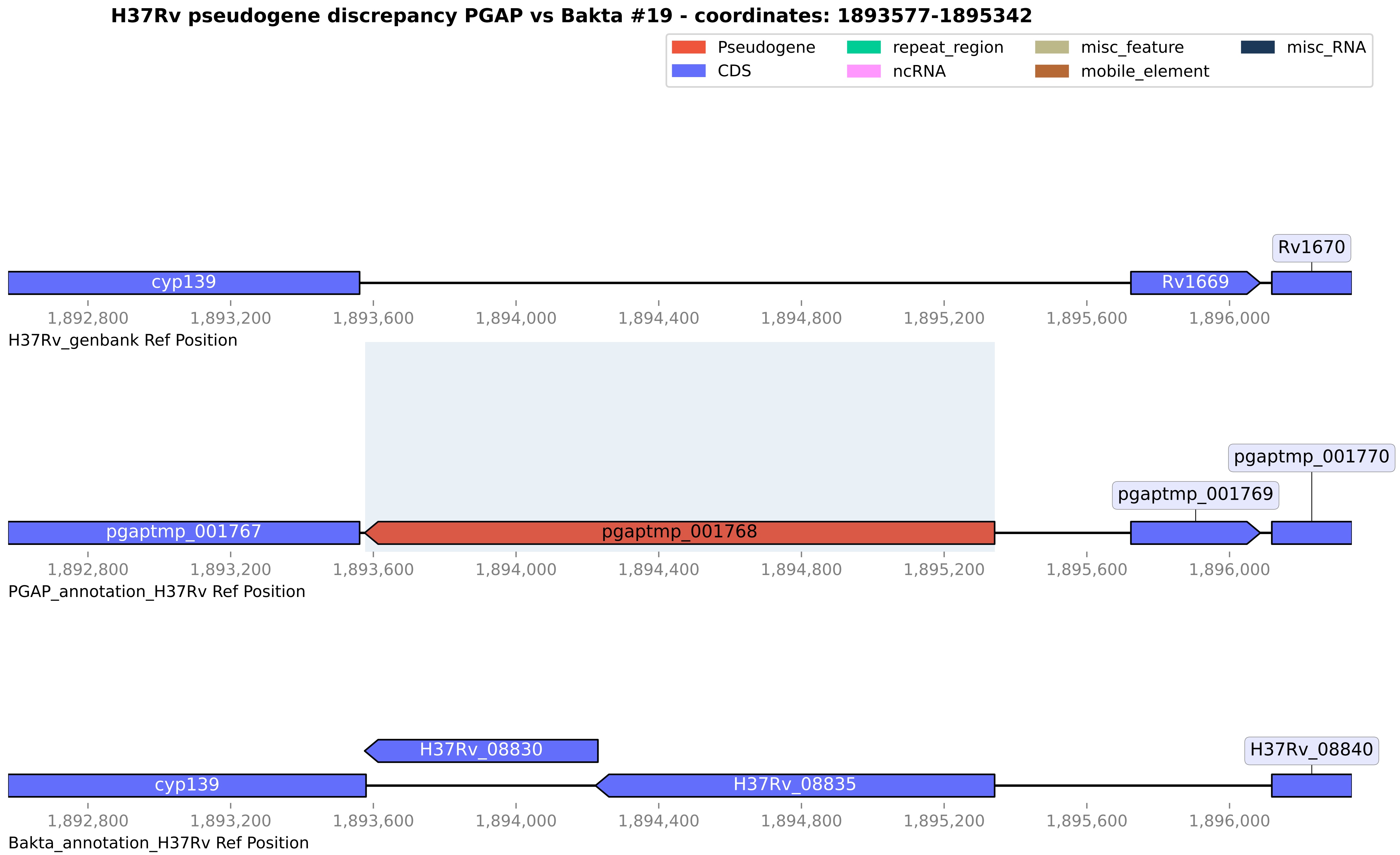
H37Rv pseudogene discrepancy PGAP vs Bakta #19 - coordinates: 1893577-1895342
Pseudogene
CDS
repeat_region
ncRNA
misc_feature
mobile_element
misc_RNA
cyp139
Rv1669
Rv1670
1,892,800
1,893,200
1,893,600
1,894,000
1,894,400
1,894,800
1,895,200
1,895,600
1,896,000
H37Rv_genbank Ref Position
pgaptmp_001767
pgaptmp_001768
pgaptmp_001769
pgaptmp_001770
PGAP_annotation_H37Rv Ref Position
H37Rv_08830
cyp139
H37Rv_08835
H37Rv_08840
Bakta_annotation_H37Rv Ref Position

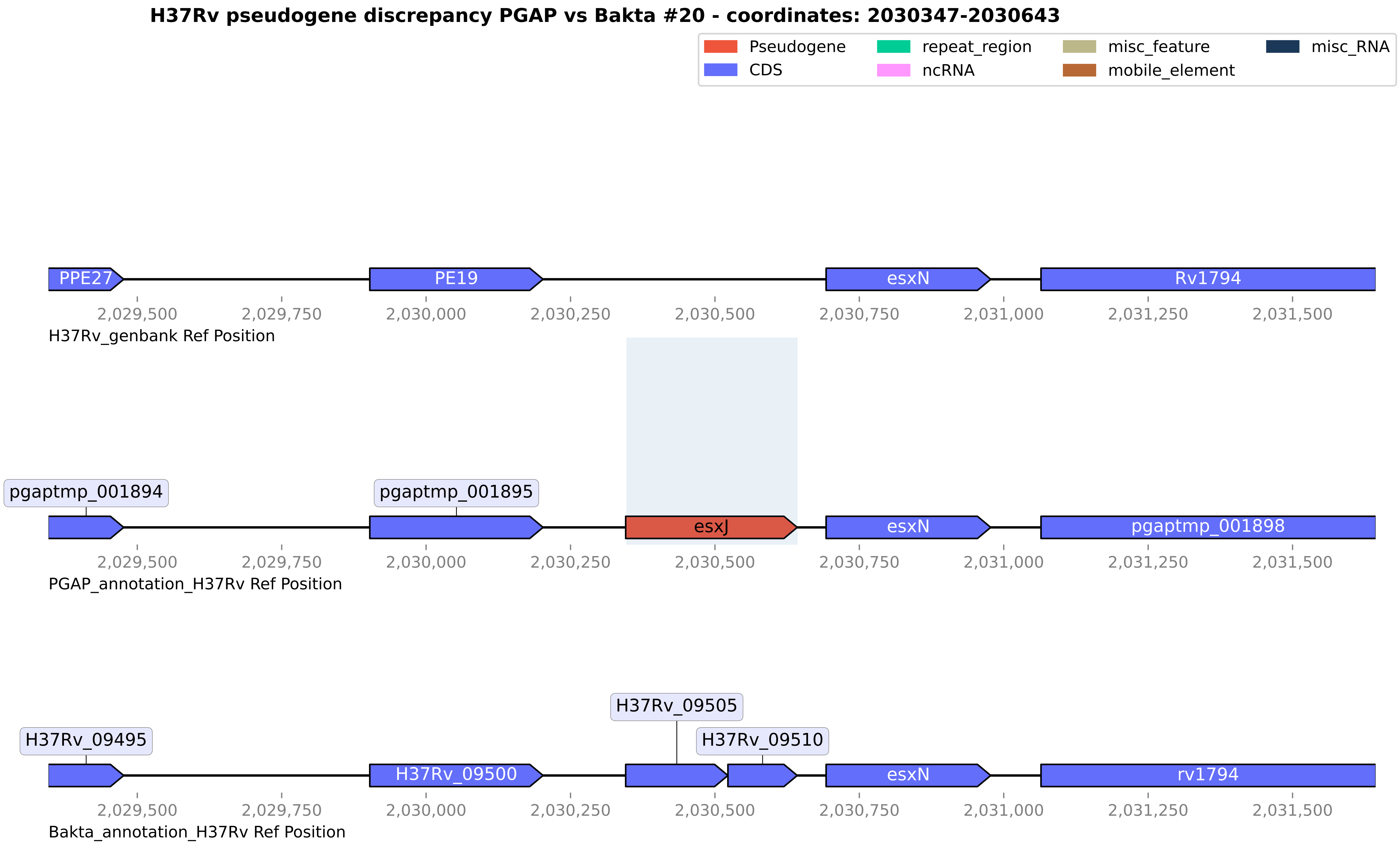
H37Rv pseudogene discrepancy PGAP vs Bakta #20 - coordinates: 2030347-2030643
Pseudogene
CDS
repeat_region
ncRNA
misc_feature
mobile_element
misc_RNA
PPE27
PE19
esxN
Rv1794
2,029,500
2,029,750
2,030,000
2,030,250
2,030,500
2,030,750
2,031,000
2,031,250
2,031,500
H37Rv_genbank Ref Position
pgaptmp_001894
pgaptmp_001895
esxJ
esxN
pgaptmp_001898
PGAP_annotation_H37Rv Ref Position
H37Rv_09495
H37Rv_09500
H37Rv_09505
H37Rv_09510
esxN
rv1794
Bakta_annotation_H37Rv Ref Position

#### H37Rv pseudogene discrepancy PGAP vs Bakta #21 - coordinates: 2138174-2139017

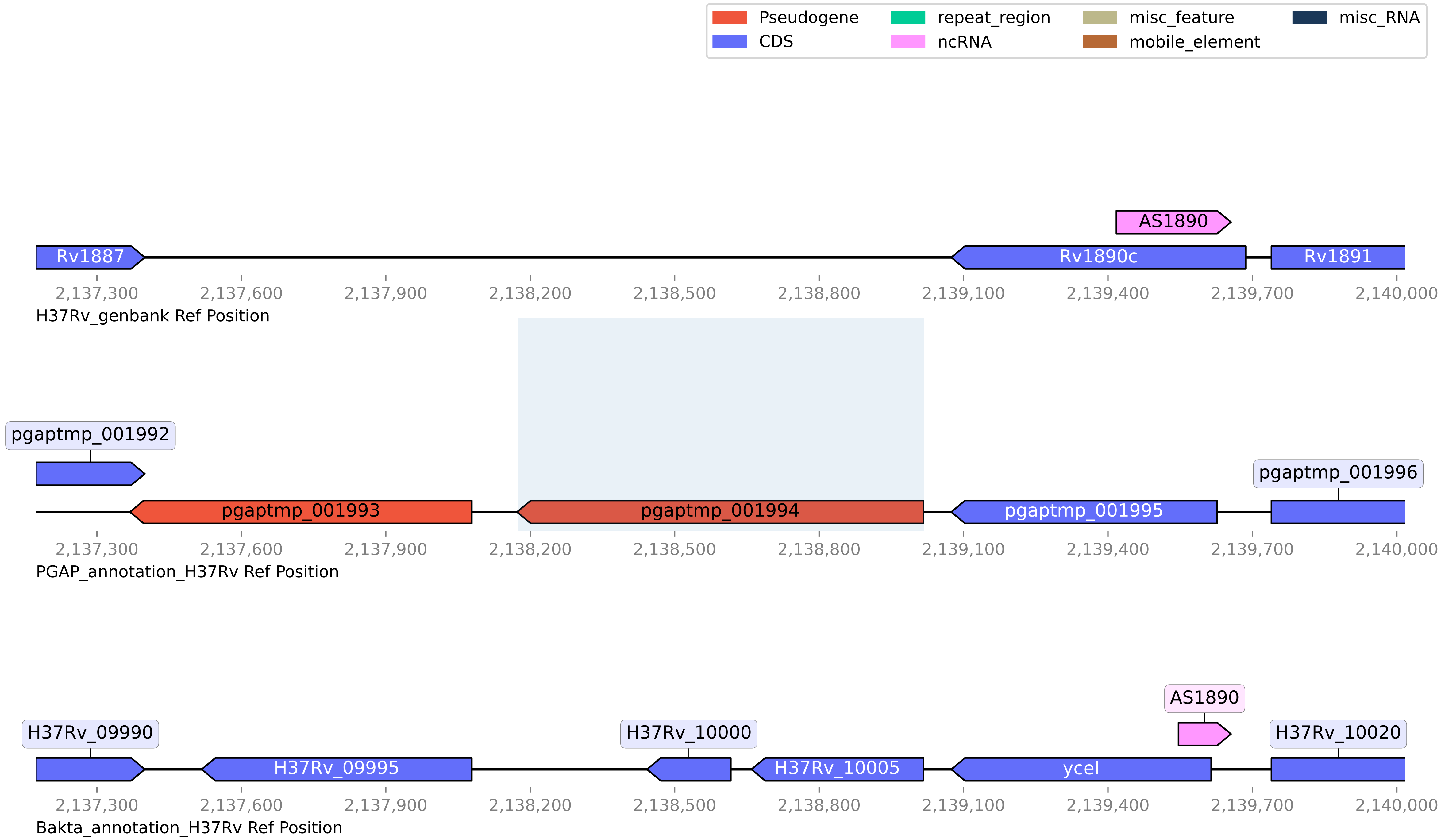

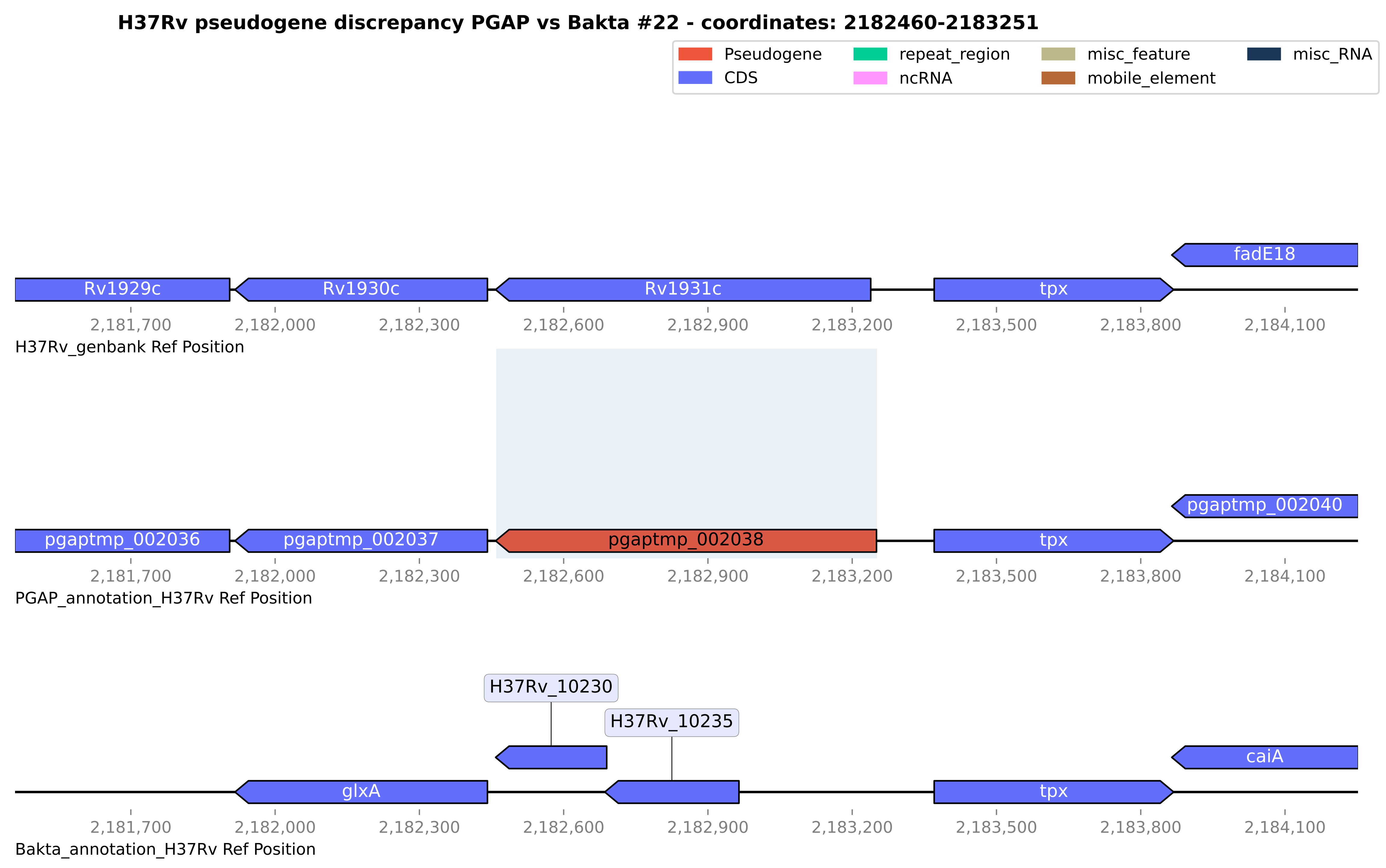
H37Rv pseudogene discrepancy PGAP vs Bakta #22 - coordinates: 2182460-2183251
Pseudogene
CDS
repeat_region
ncRNA
misc_feature
mobile_element
misc_RNA
fadE18
Rv1929c
Rv1930c
Rv1931c
tpx
2,181,700
2,182,000
2,182,300
2,182,600
2,182,900
2,183,200
2,183,500
2,183,800
2,184,100
H37Rv_genbank Ref Position
pgaptmp_002040
pgaptmp_002036
pgaptmp_002037
pgaptmp_002038
tpx
PGAP_annotation_H37Rv Ref Position
H37Rv_10230
H37Rv_10235
caiA
glxA
tpx
Bakta_annotation_H37Rv Ref Position

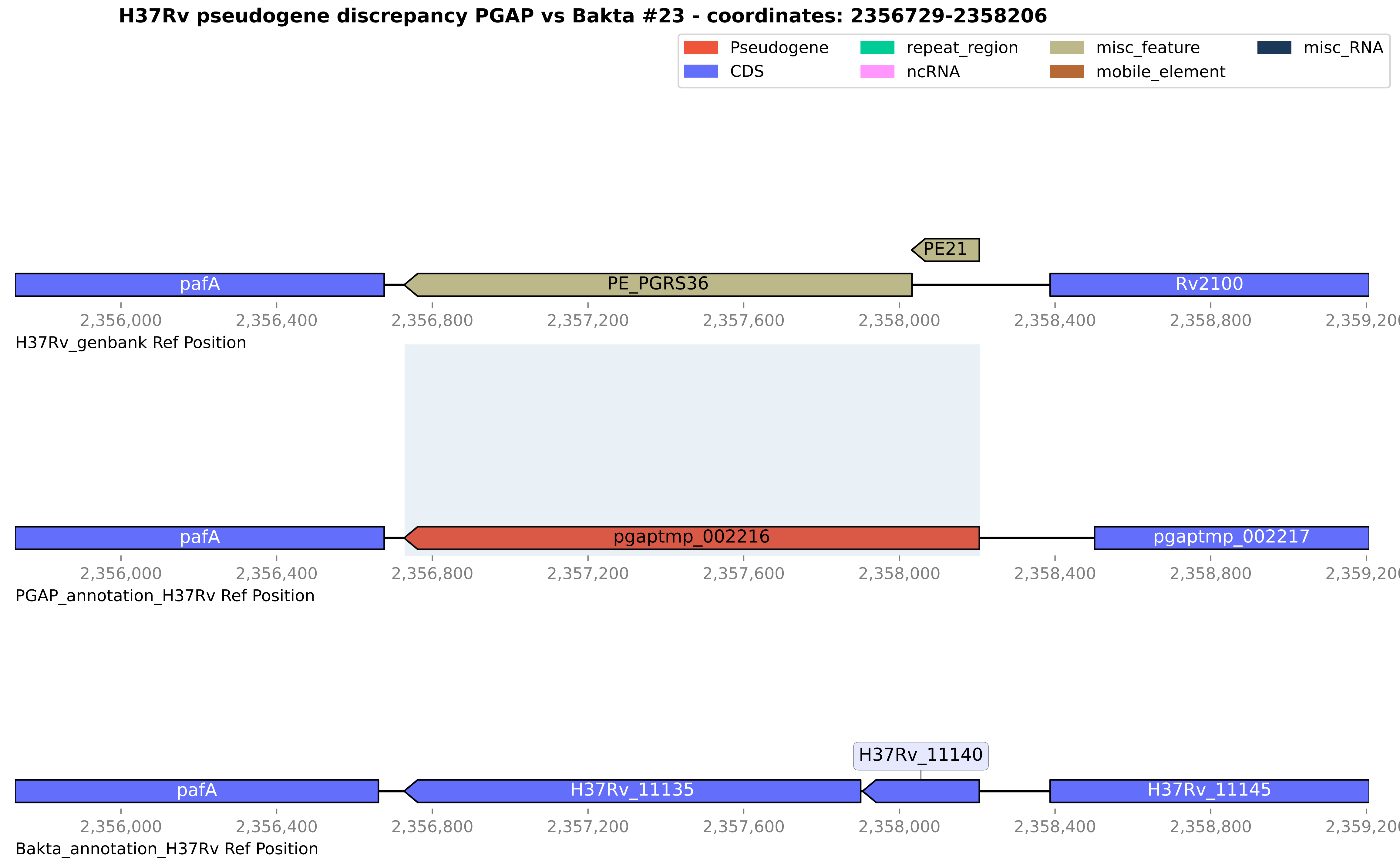
H37Rv pseudogene discrepancy PGAP vs Bakta #23 - coordinates: 2356729-2358206
Pseudogene
CDS
repeat_region
ncRNA
misc_feature
mobile_element
misc_RNA
PE21
pafA
PE_PGRS36
Rv2100
2,356,000
2,356,400
2,356,800
2,357,200
2,357,600
2,358,000
2,358,400
2,358,800
2,359,200
H37Rv_genbank Ref Position
pafA
pgaptmp_002216
pgaptmp_002217
PGAP_annotation_H37Rv Ref Position
H37Rv_11140
pafA
H37Rv_11135
H37Rv_11145
Bakta_annotation_H37Rv Ref Position

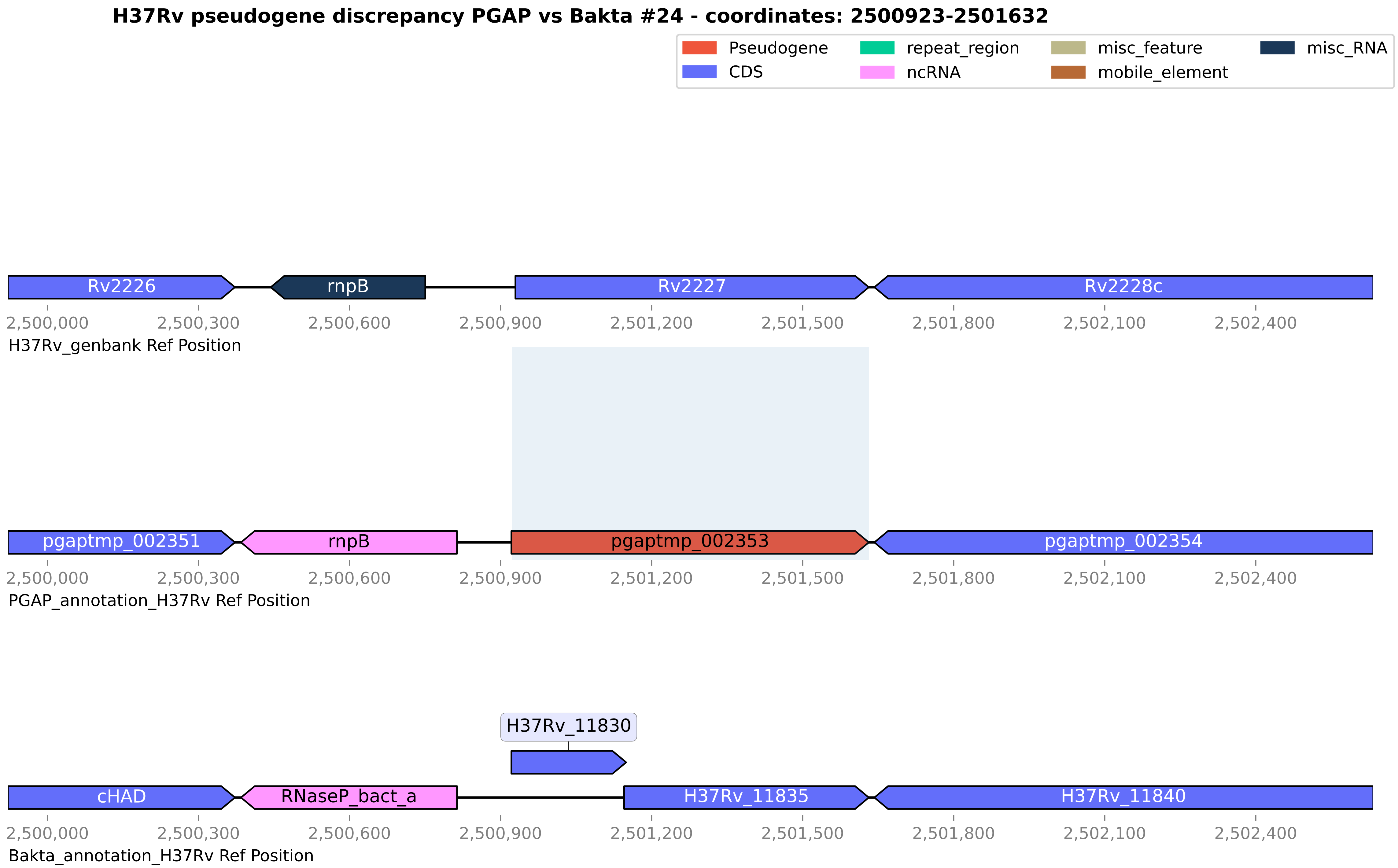

H37Rv pseudogene discrepancy PGAP vs Bakta #24 - coordinates: 2500923-2501632
Pseudogene
CDS
repeat_region
ncRNA
misc_feature
mobile_element
misc_RNA
Rv2226
rnpB
Rv2227
Rv2228c
2,500,000
2,500,300
2,500,600
2,500,900
2,501,200
2,501,500
2,501,800
2,502,100
2,502,400
H37Rv_genbank Ref Position
pgaptmp_002351
rnpB
pgaptmp_002353
pgaptmp_002354
PGAP_annotation_H37Rv Ref Position
H37Rv_11830
cHAD
RNaseP_bact_a
H37Rv_11835
H37Rv_11840
Bakta_annotation_H37Rv Ref Position

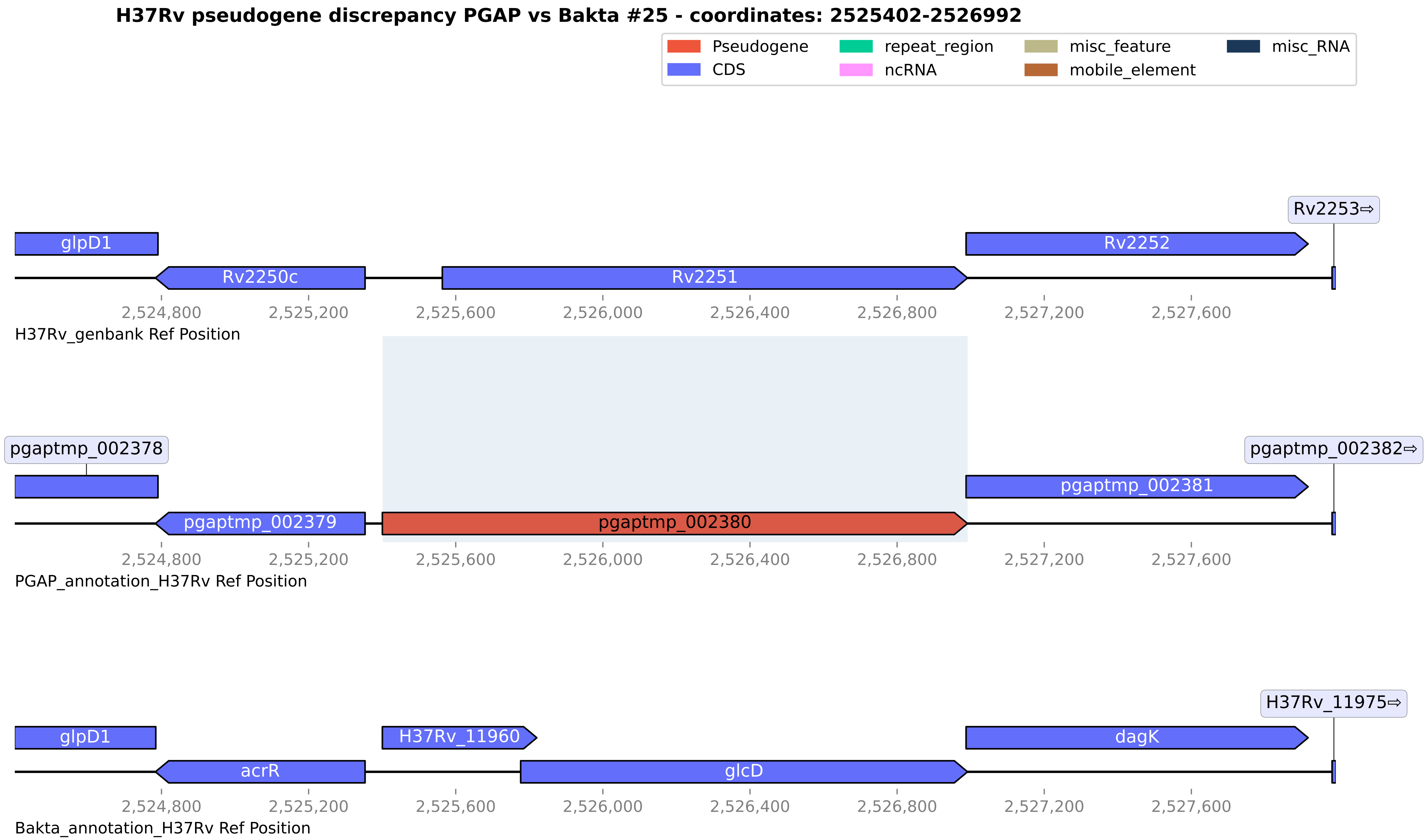

H37Rv pseudogene discrepancy PGAP vs Bakta #25 - coordinates: 2525402-2526992
Pseudogene
CDS
repeat_region
ncRNA
misc_feature
mobile_element
misc_RNA
Rv2253⇨
glpD1
Rv2252
Rv2250c
Rv2251
2,524,800
2,525,200
2,525,600
2,526,000
2,526,400
2,526,800
2,527,200
2,527,600
H37Rv_genbank Ref Position
pgaptmp_002378
pgaptmp_002382⇨
pgaptmp_002381
pgaptmp_002379
pgaptmp_002380
2,524,800
2,525,200
2,525,600
2,526,000
2,526,400
2,526,800
2,527,200
2,527,600
PGAP_annotation_H37Rv Ref Position
H37Rv_11975⇨
glpD1
H37Rv_11960
dagK
acrR
glcD
2,524,800
2,525,200
2,525,600
2,526,000
2,526,400
2,526,800
2,527,200
2,527,600
Bakta_annotation_H37Rv Ref Position

**H37Rv pseudogene discrepancy PGAP vs Bakta #26 - coordinates: 2534042-2535552**

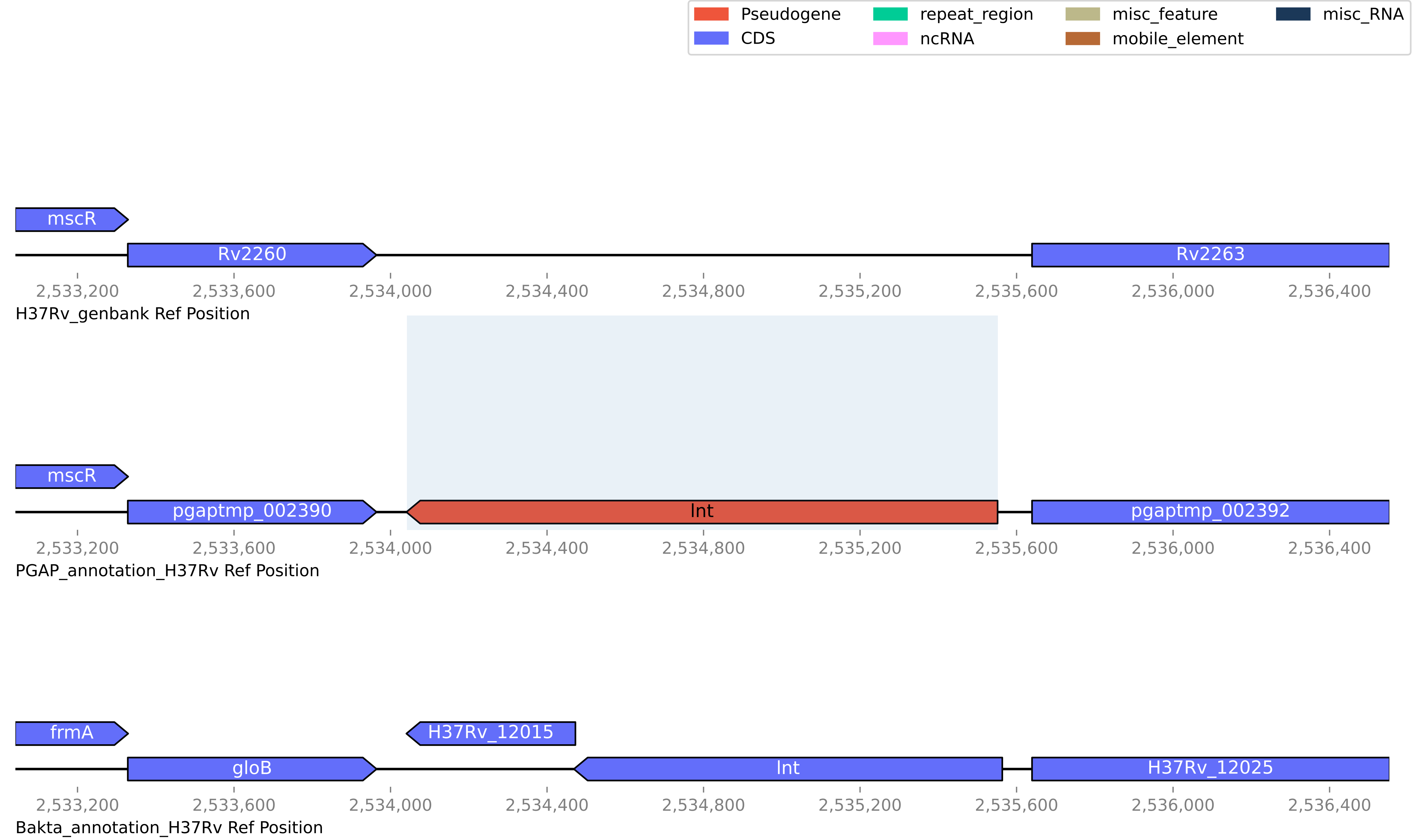

#### H37Rv pseudogene discrepancy PGAP vs Bakta #27 - coordinates: 2881409-2882147

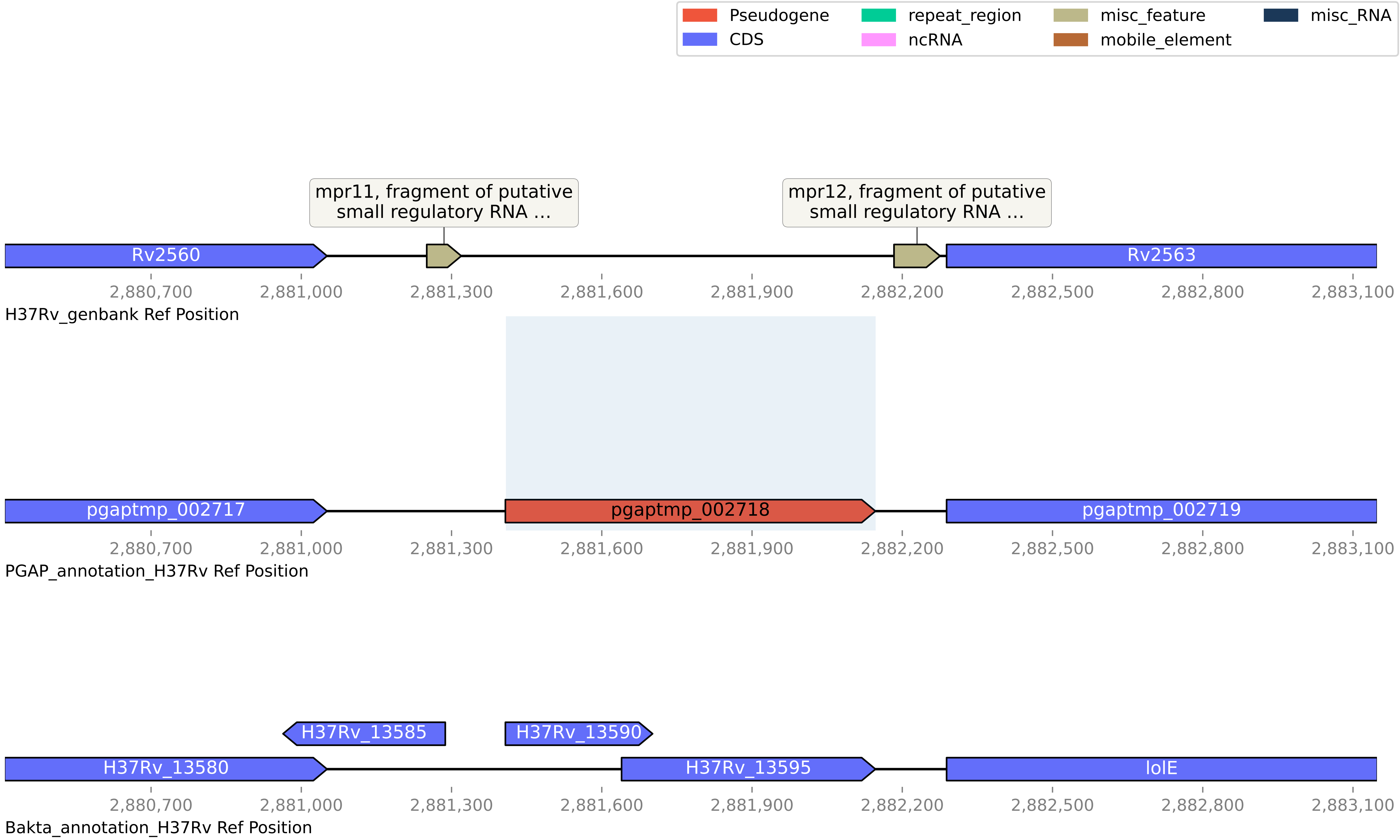

#### H37Rv pseudogene discrepancy PGAP vs Bakta #28 - coordinates: 3291503-3297819

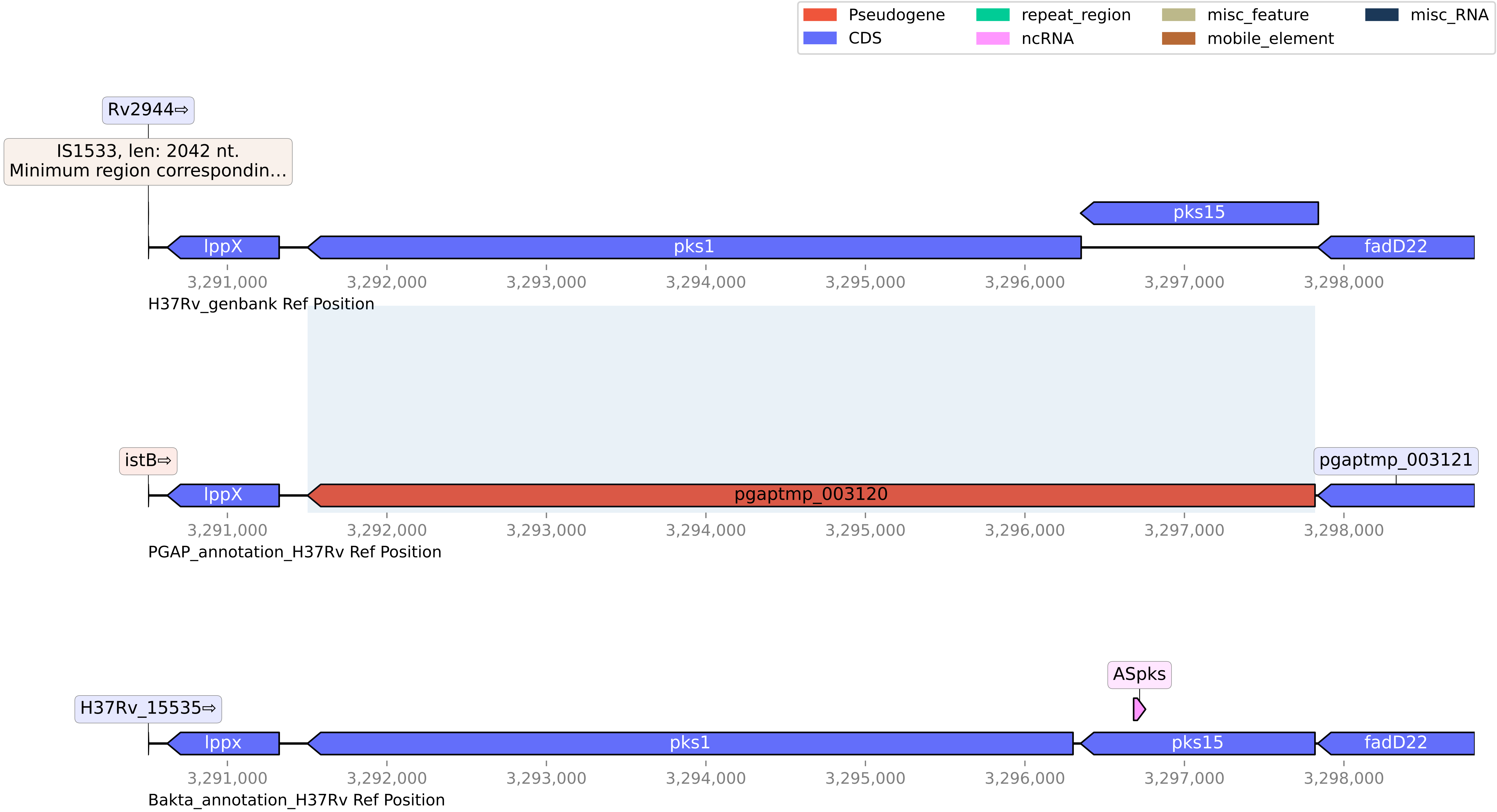

**H37Rv pseudogene discrepancy PGAP vs Bakta #29 - coordinates: 3329949-3331612**

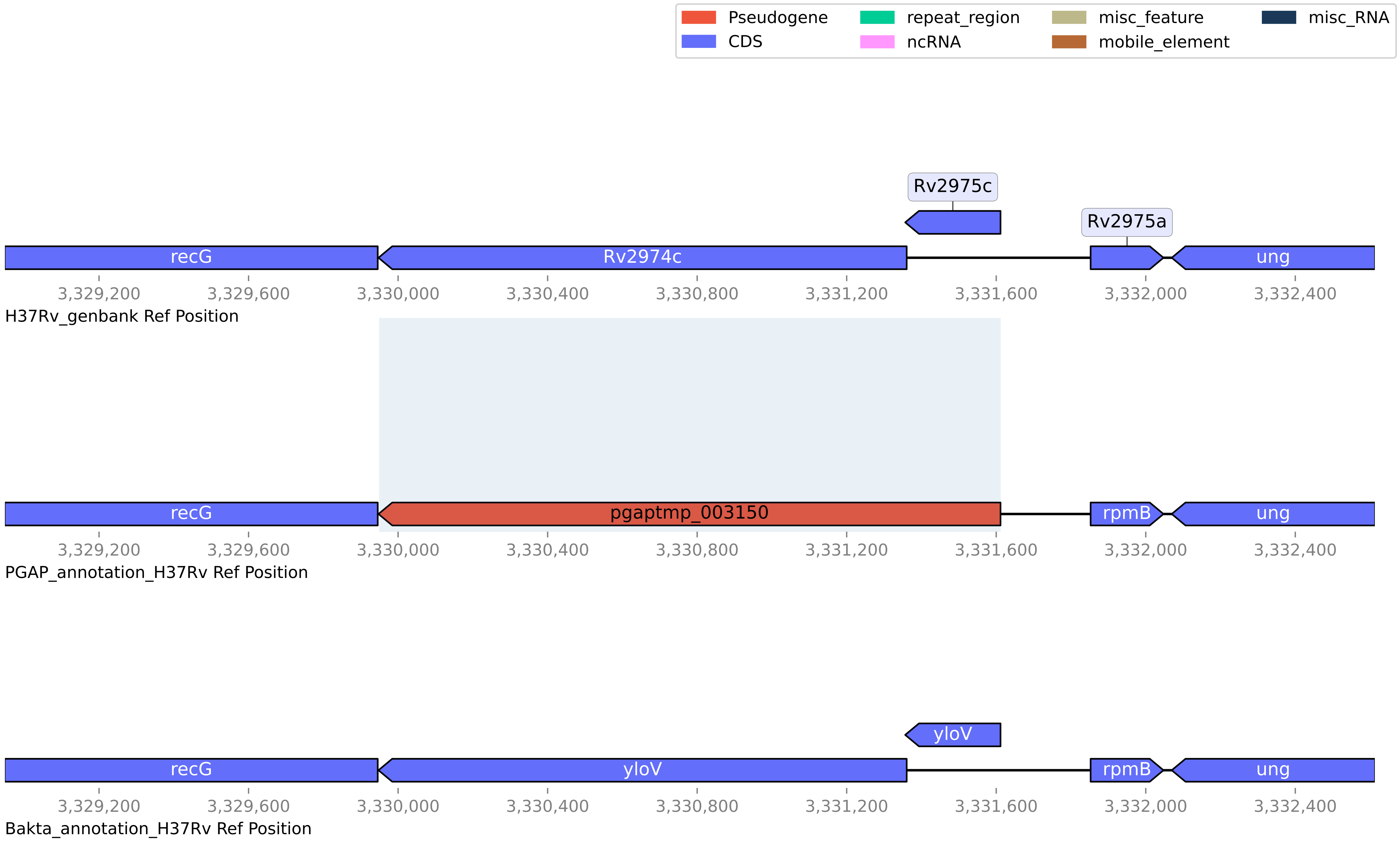

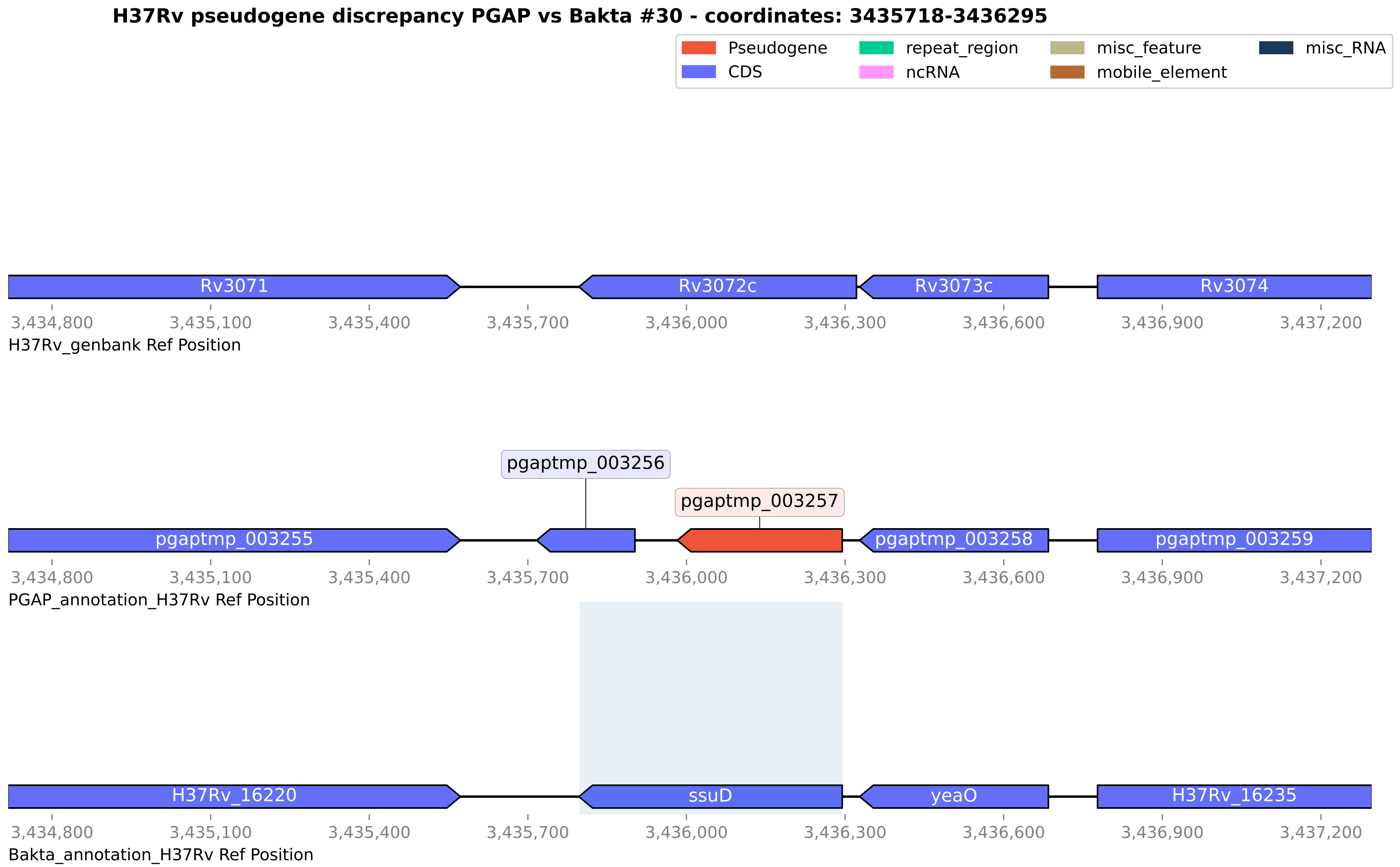
H37Rv pseudogene discrepancy PGAP vs Bakta #30 - coordinates: 3435718-3436295
Pseudogene
CDS
repeat_region
ncRNA
misc_feature
mobile_element
misc_RNA
Rv3071
Rv3072c
Rv3073c
Rv3074
3,434,800
3,435,100
3,435,400
3,435,700
3,436,000
3,436,300
3,436,600
3,436,900
3,437,200
H37Rv_genbank Ref Position
pgaptmp_003256
pgaptmp_003257
pgaptmp_003255
pgaptmp_003258
pgaptmp_003259
PGAP_annotation_H37Rv Ref Position
H37Rv_16220
ssuD
yeaO
H37Rv_16235
Bakta_annotation_H37Rv Ref Position

#### H37Rv pseudogene discrepancy PGAP vs Bakta #31 - coordinates: 3609781-3611189

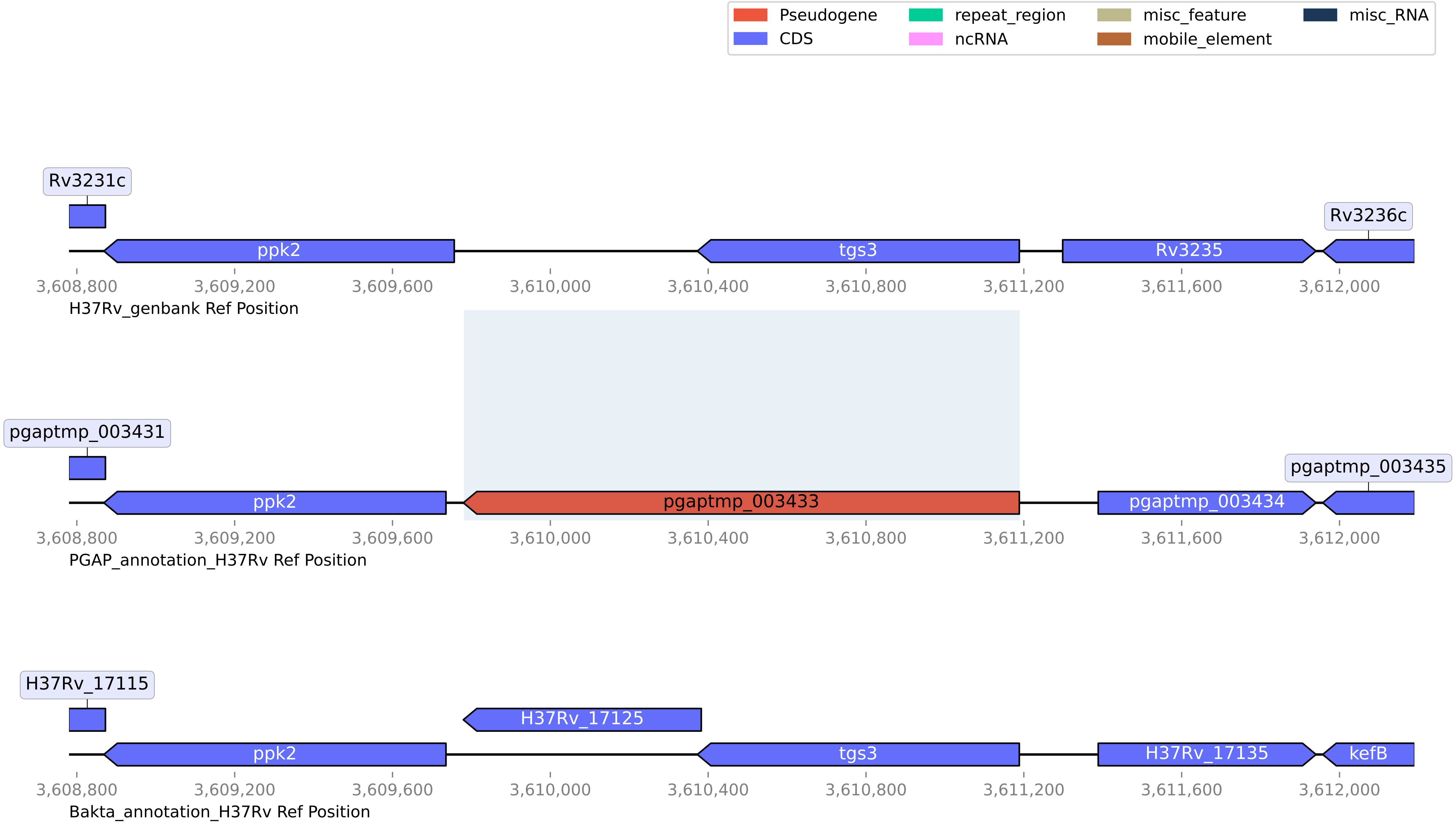

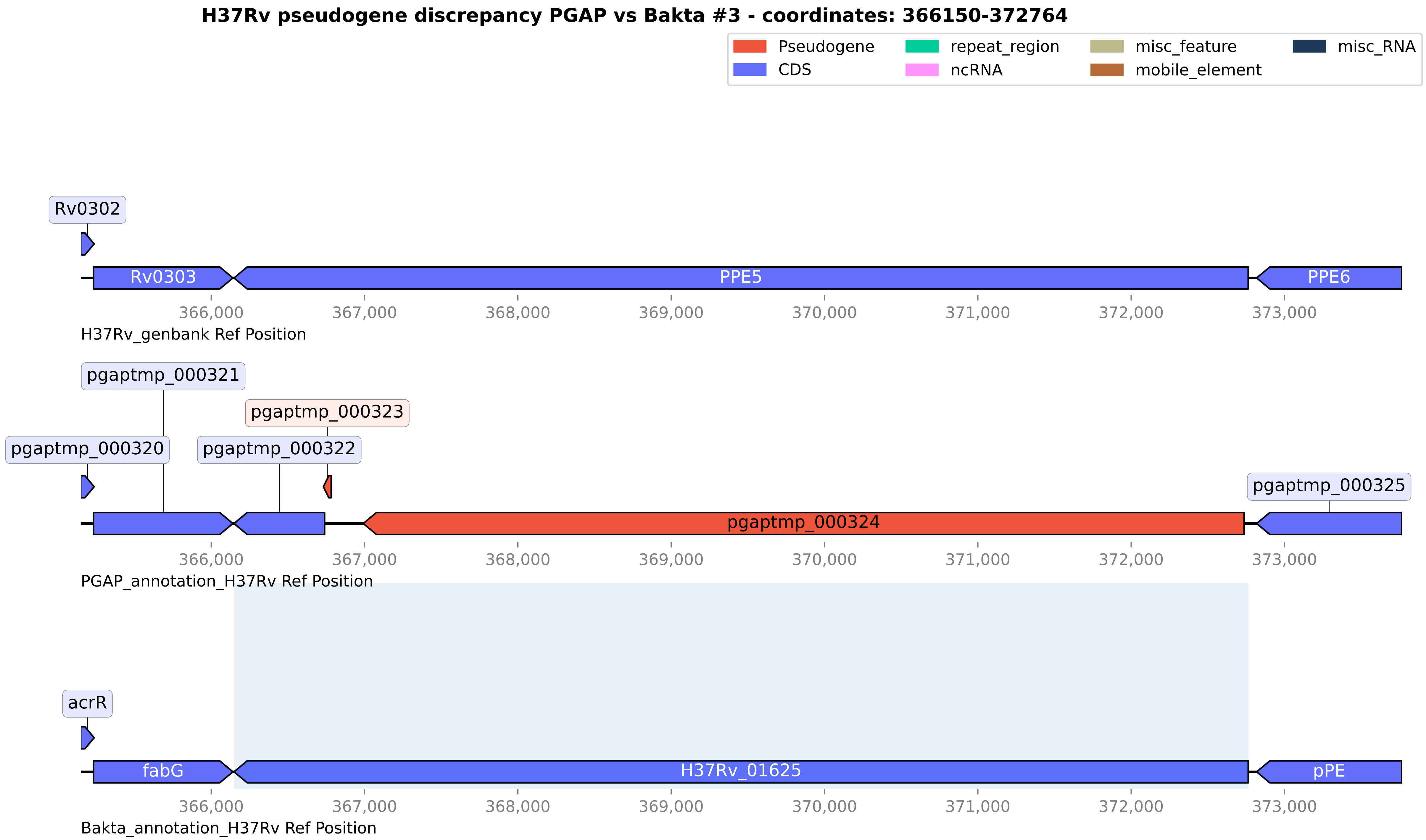

H37Rv pseudogene discrepancy PGAP vs Bakta #3 - coordinates: 366150-372764
Pseudogene
CDS
repeat_region
ncRNA
misc_feature
mobile_element
misc_RNA
Rv0302
Rv0303
PPE5
PPE6
366,000
367,000
368,000
369,000
370,000
371,000
372,000
373,000
H37Rv_genbank Ref Position
pgaptmp_000321
pgaptmp_000323
pgaptmp_000320
pgaptmp_000322
pgaptmp_000325
pgaptmp_000324
PGAP_annotation_H37Rv Ref Position
acrR
fabG
H37Rv_01625
pPE
Bakta_annotation_H37Rv Ref Position

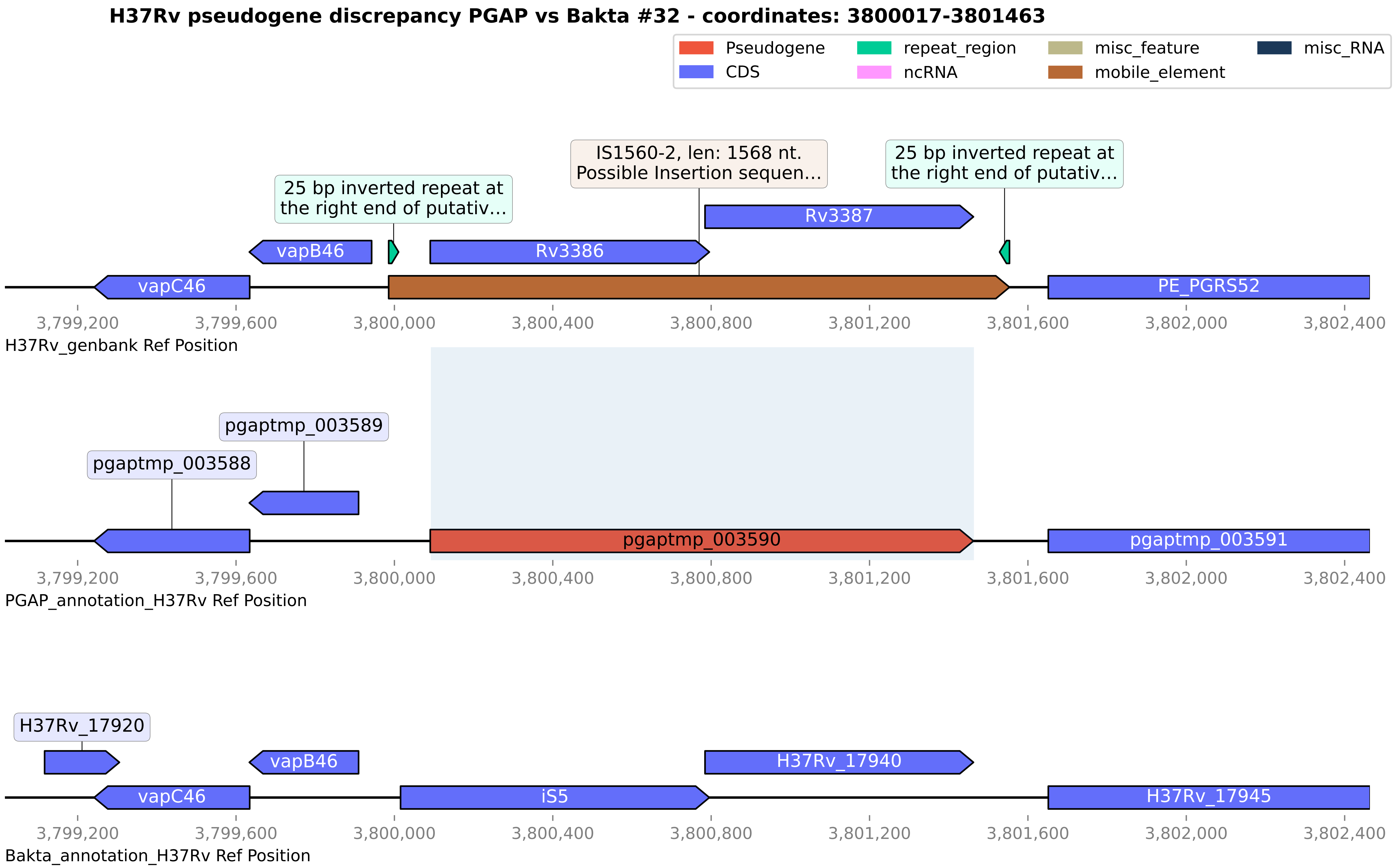

H37Rv pseudogene discrepancy PGAP vs Bakta #32 - coordinates: 3800017-3801463
Pseudogene
CDS
repeat_region
ncRNA
misc_feature
mobile_element
misc_RNA
25 bp inverted repeat at the right end of putativ...
IS1560-2, len: 1568 nt. Possible Insertion sequen...
25 bp inverted repeat at the right end of putativ...
Rv3387
vapB46
Rv3386
vapC46
PE_PGRS52
3,799,200
3,799,600
3,800,000
3,800,400
3,800,800
3,801,200
3,801,600
3,802,000
3,802,400
H37Rv_genbank Ref Position
pgaptmp_003589
pgaptmp_003588
pgaptmp_003590
pgaptmp_003591
PGAP_annotation_H37Rv Ref Position
H37Rv_17920
vapB46
H37Rv_17940
vapC46
iS5
H37Rv_17945
Bakta_annotation_H37Rv Ref Position

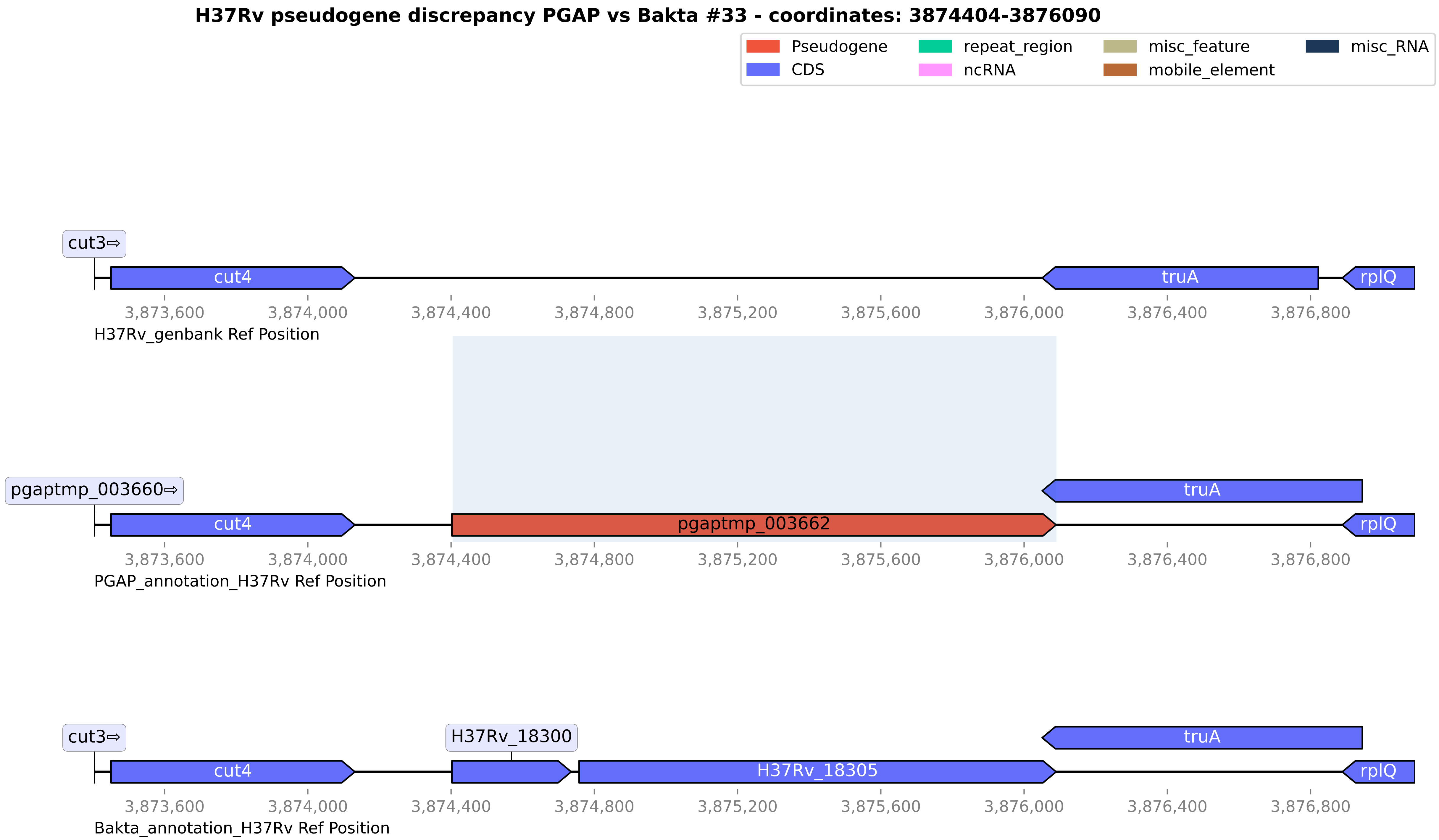
H37Rv pseudogene discrepancy PGAP vs Bakta #33 - coordinates: 3874404-3876090
Pseudogene
CDS
repeat_region
ncRNA
misc_feature
mobile_element
misc_RNA
cut3⇨
cut4
truA
rplQ
3,873,600
3,874,000
3,874,400
3,874,800
3,875,200
3,875,600
3,876,000
3,876,400
3,876,800
H37Rv_genbank Ref Position
pgaptmp_003660⇨
pgaptmp_003662
PGAP_annotation_H37Rv Ref Position
H37Rv_18300
H37Rv_18305
Bakta_annotation_H37Rv Ref Position

**H37Rv pseudogene discrepancy PGAP vs Bakta #34 - coordinates: 4075752-4076984**

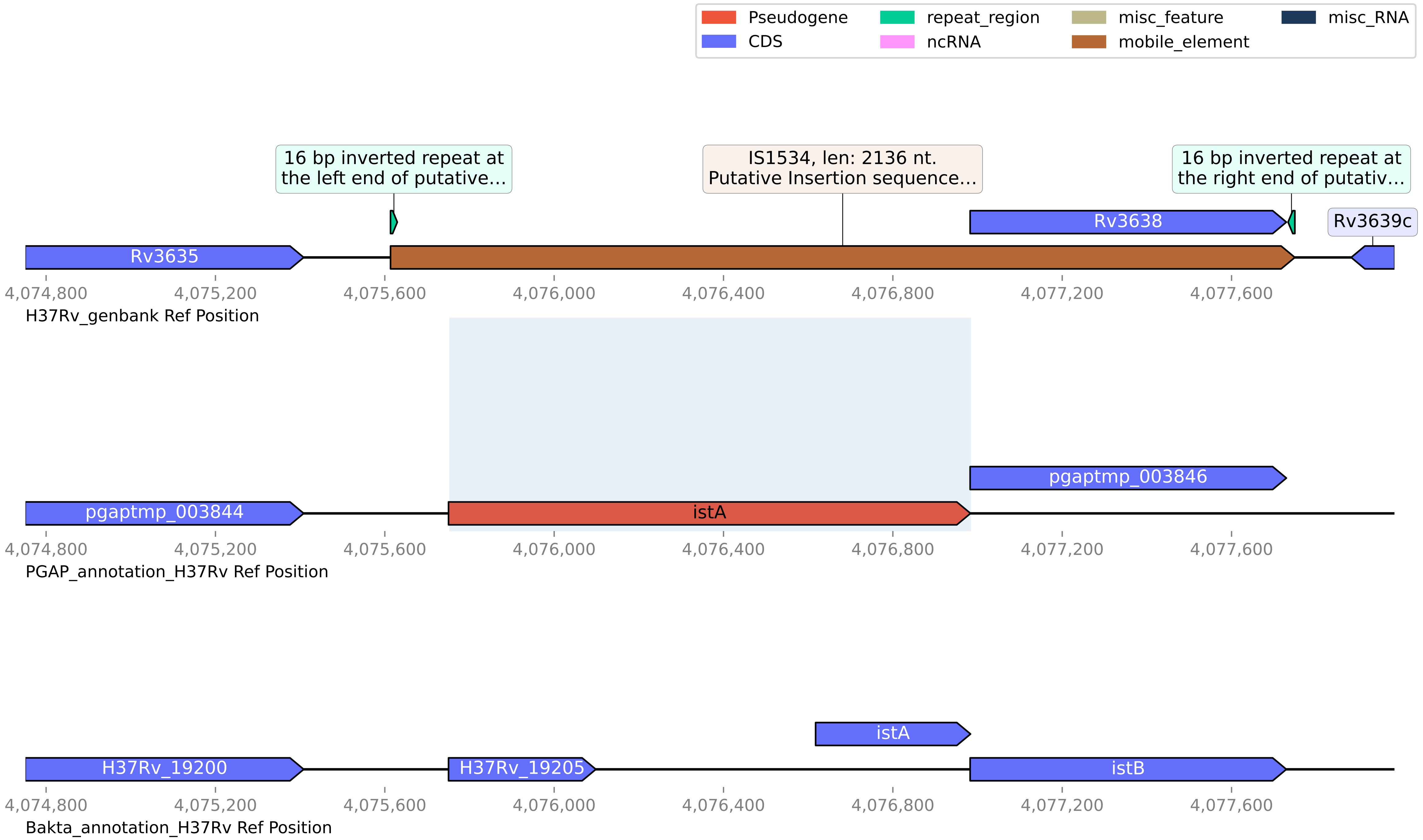

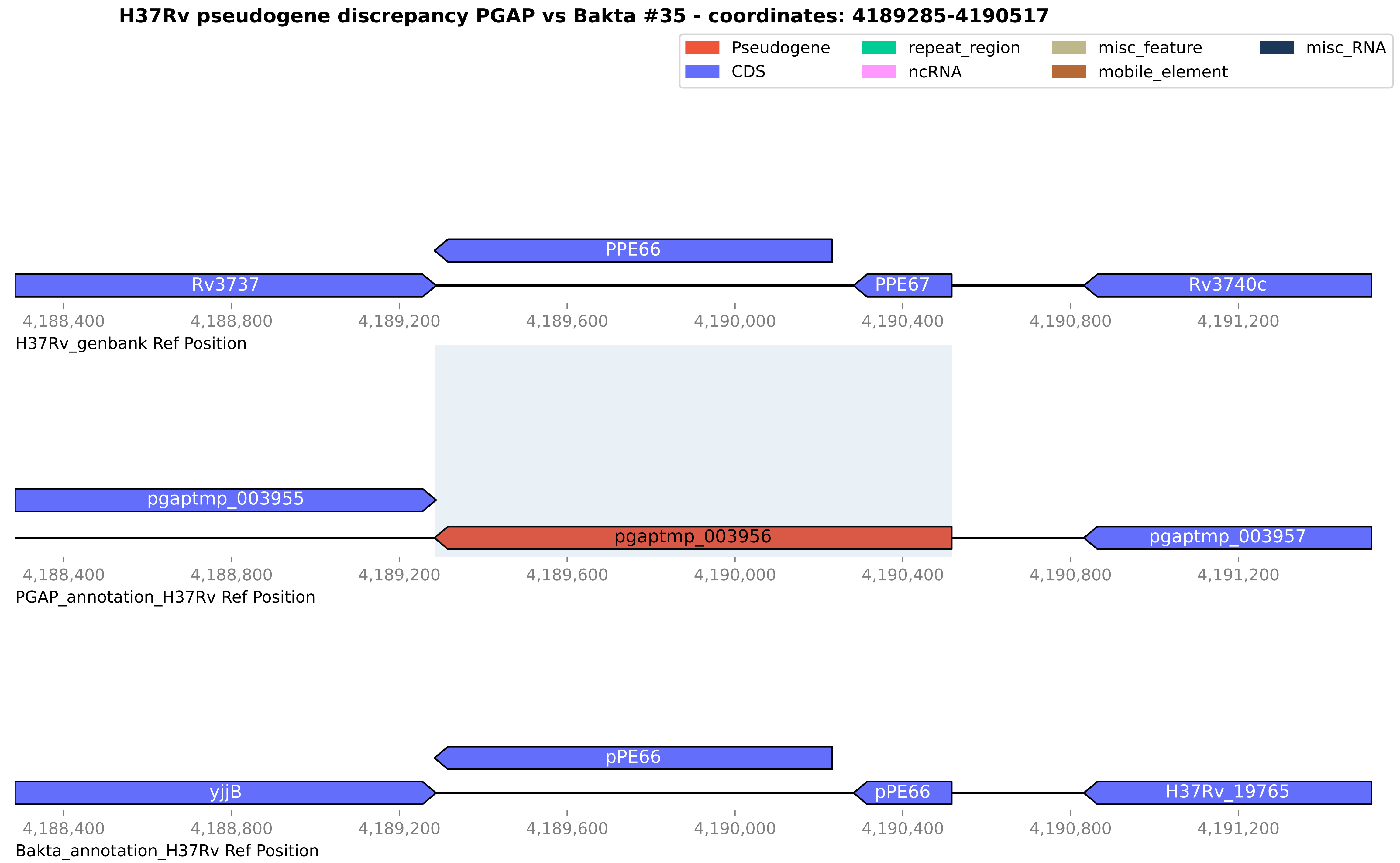

H37Rv pseudogene discrepancy PGAP vs Bakta #35 - coordinates: 4189285-4190517
Pseudogene
CDS
repeat_region
ncRNA
misc_feature
mobile_element
misc_RNA
PPE66
Rv3737
PPE67
Rv3740c
4,188,400
4,188,800
4,189,200
4,189,600
4,190,000
4,190,400
4,190,800
4,191,200
H37Rv_genbank Ref Position
pgaptmp_003955
pgaptmp_003956
pgaptmp_003957
PGAP_annotation_H37Rv Ref Position
pPE66
yjjB
pPE66
H37Rv_19765
Bakta_annotation_H37Rv Ref Position

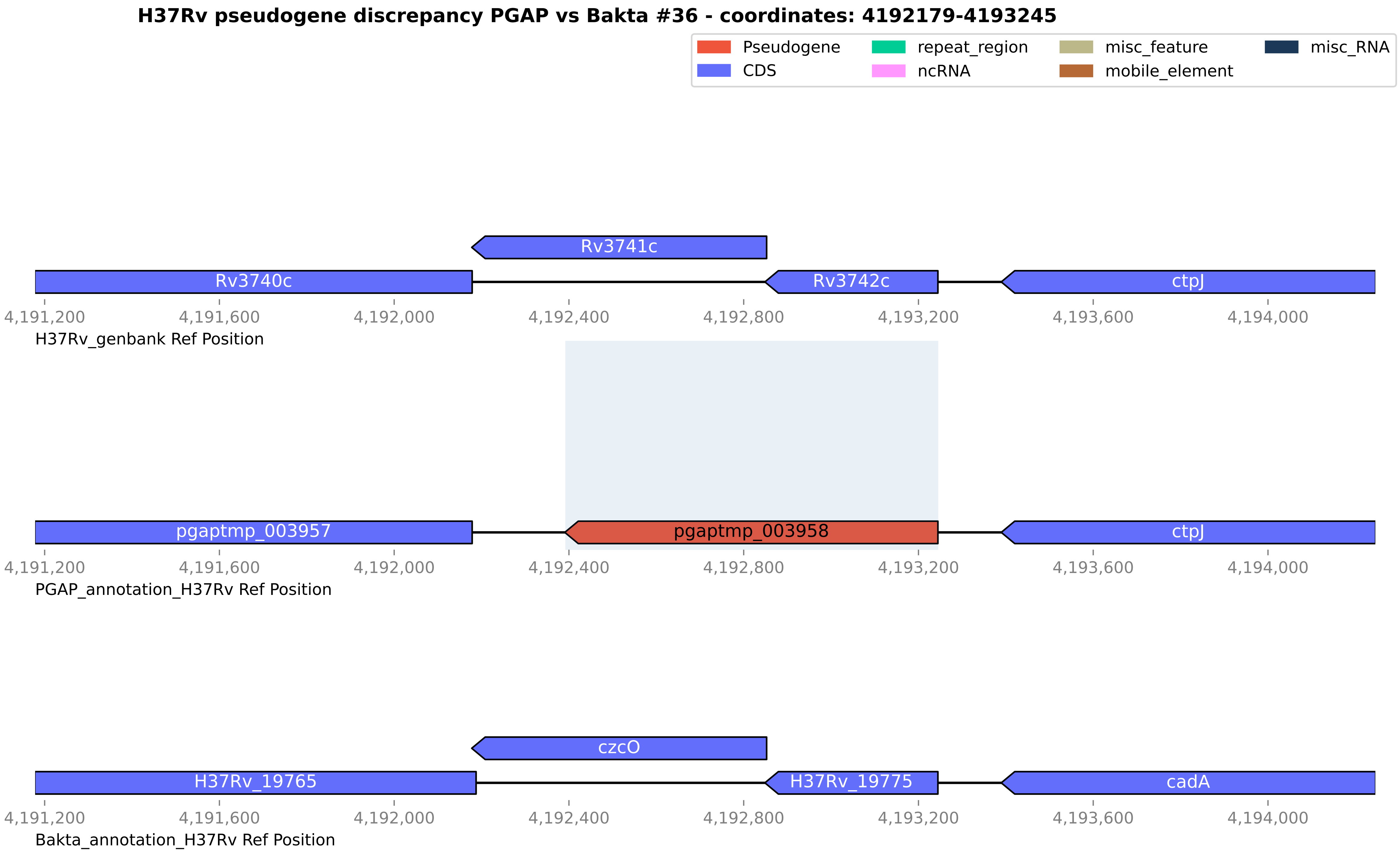
H37Rv pseudogene discrepancy PGAP vs Bakta #36 - coordinates: 4192179-4193245
Pseudogene
CDS
repeat_region
ncRNA
misc_feature
mobile_element
misc_RNA
Rv3741c
Rv3740c
Rv3742c
ctpJ
4,191,200
4,191,600
4,192,000
4,192,400
4,192,800
4,193,200
4,193,600
4,194,000
H37Rv_genbank Ref Position
pgaptmp_003957
pgaptmp_003958
ctpJ
PGAP_annotation_H37Rv Ref Position
czcO
H37Rv_19765
H37Rv_19775
cadA
Bakta_annotation_H37Rv Ref Position

#### H37Rv pseudogene discrepancy PGAP vs Bakta #37 - coordinates: 4215881-4216295

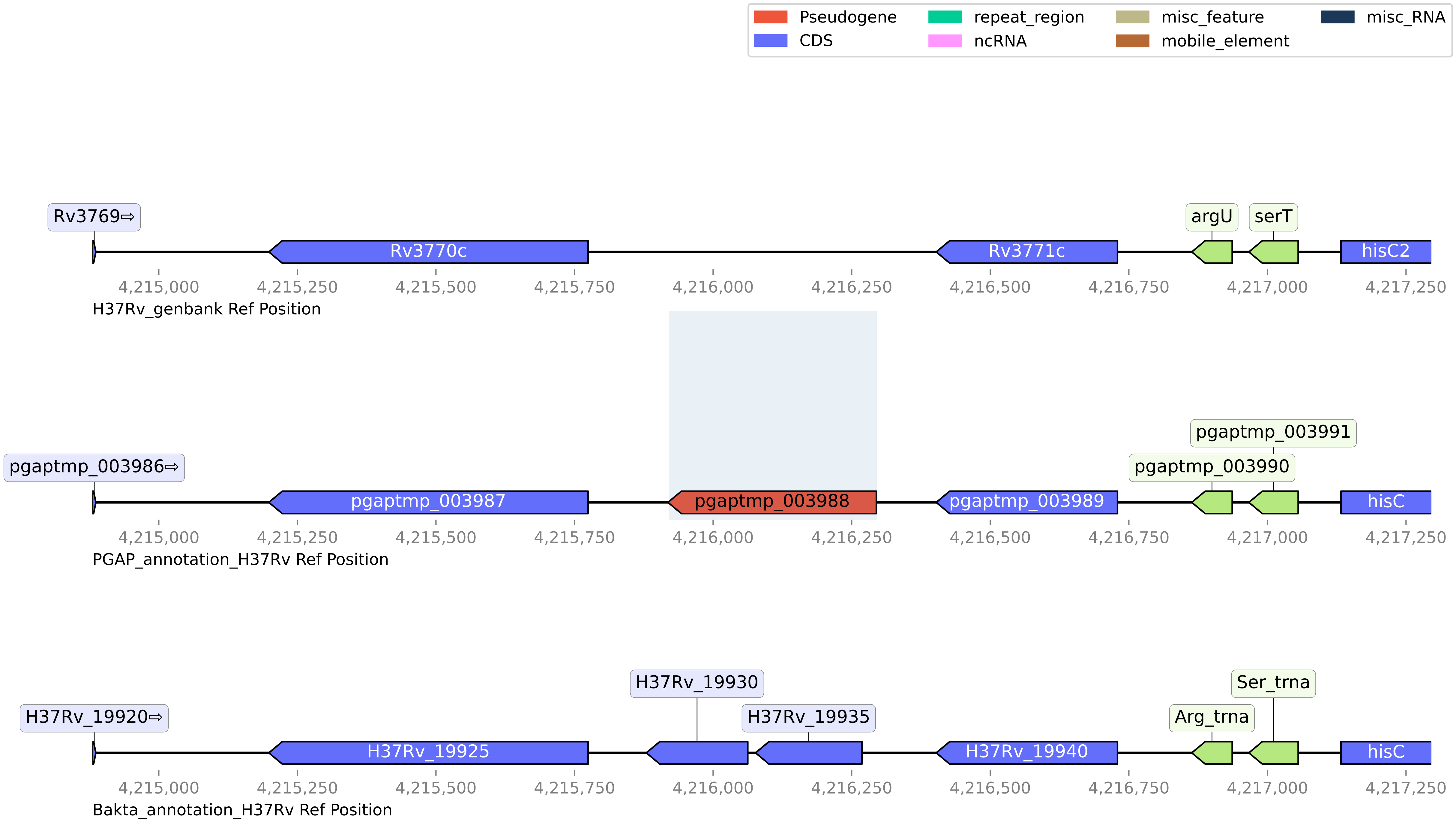

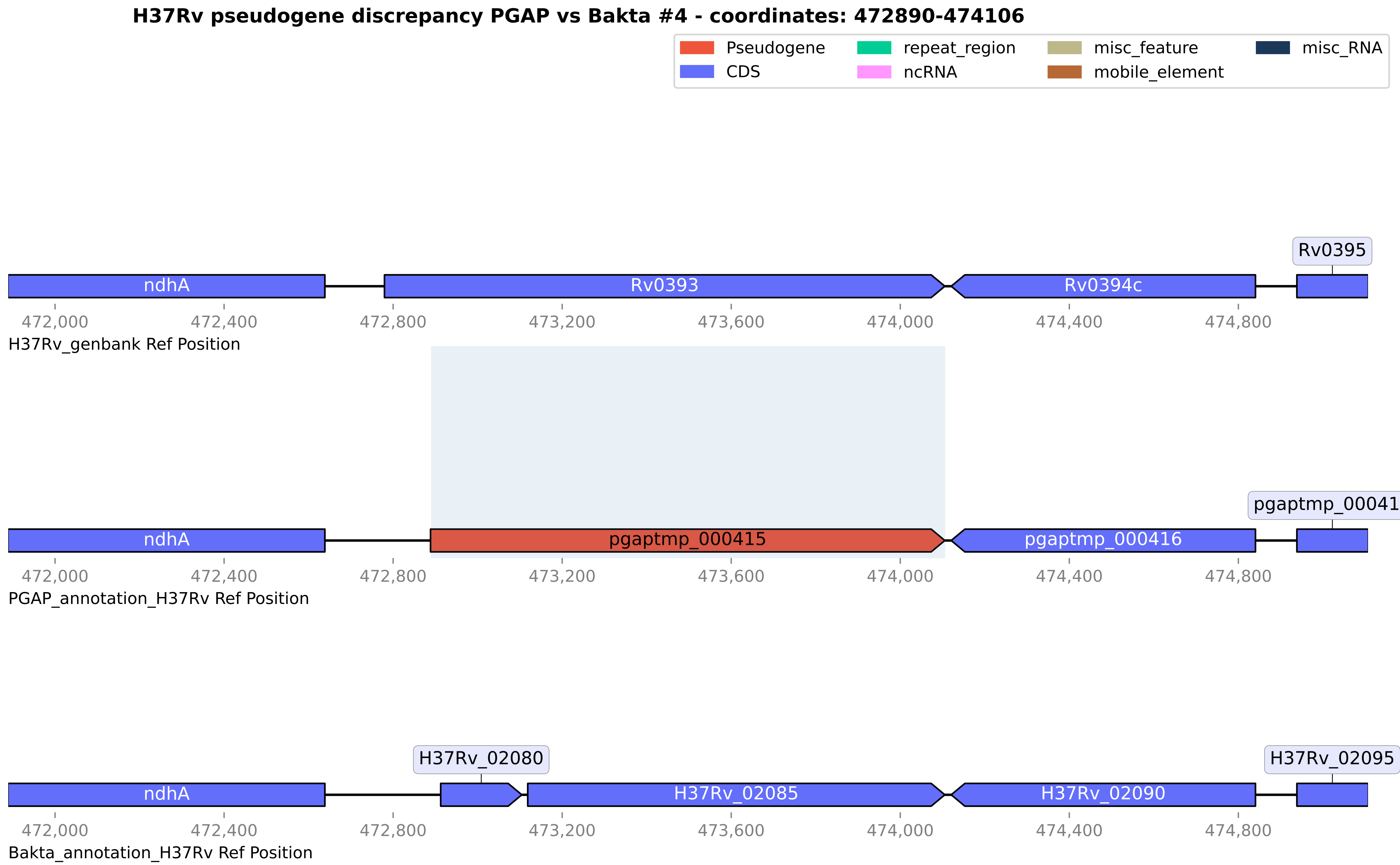

H37Rv pseudogene discrepancy PGAP vs Bakta #4 - coordinates: 472890-474106
Pseudogene
CDS
repeat_region
ncRNA
misc_feature
mobile_element
misc_RNA
Rv0395
ndhA
Rv0393
Rv0394c
472,000
472,400
472,800
473,200
473,600
474,000
474,400
474,800
H37Rv_genbank Ref Position
pgaptmp_000417
ndhA
pgaptmp_000415
pgaptmp_000416
PGAP_annotation_H37Rv Ref Position
H37Rv_02080
H37Rv_02095
ndhA
H37Rv_02085
H37Rv_02090
Bakta_annotation_H37Rv Ref Position

**H37Rv pseudogene discrepancy PGAP vs Bakta #5 - coordinates: 688032-689062**

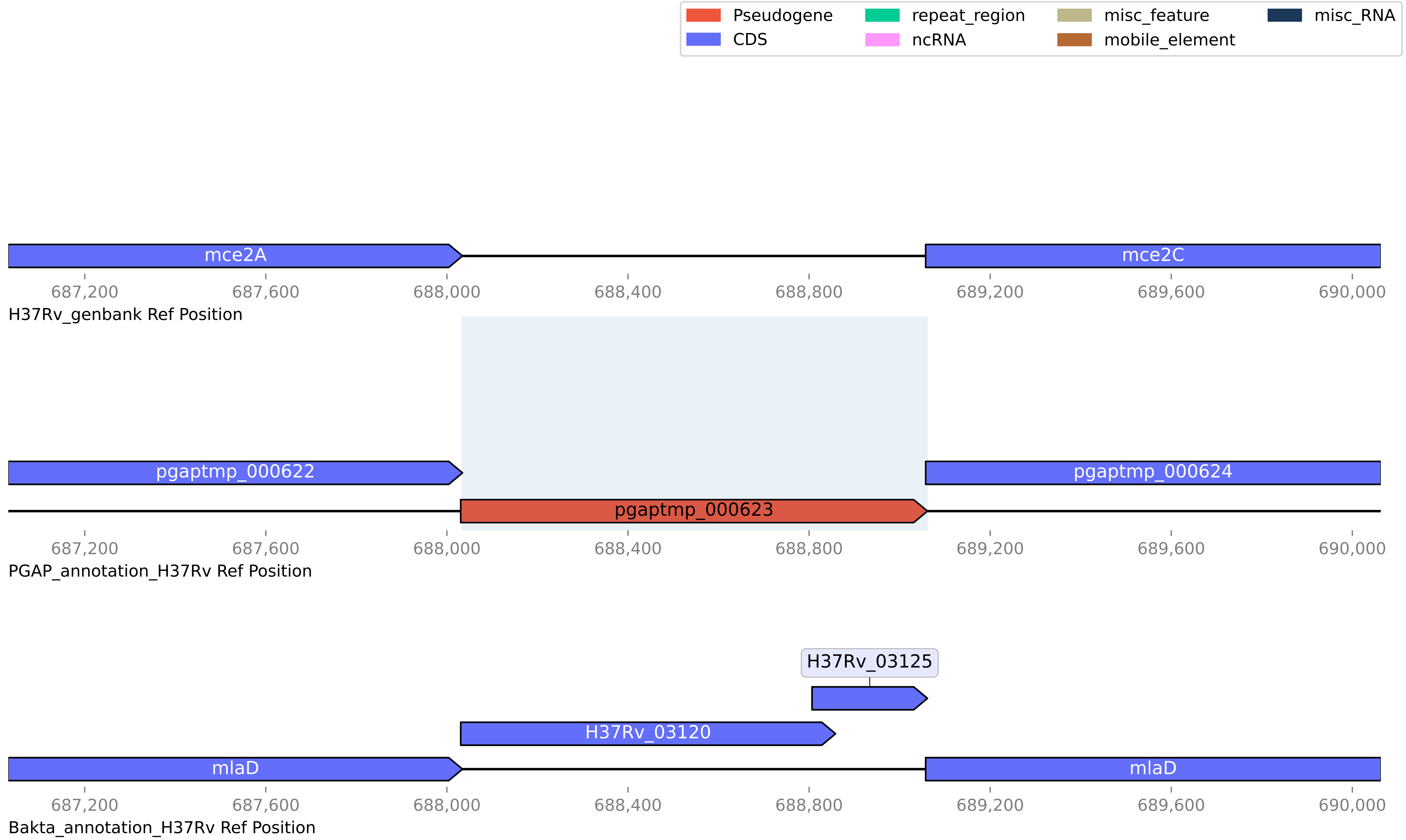

#### H37Rv pseudogene discrepancy PGAP vs Bakta #6 - coordinates: 711536-712719

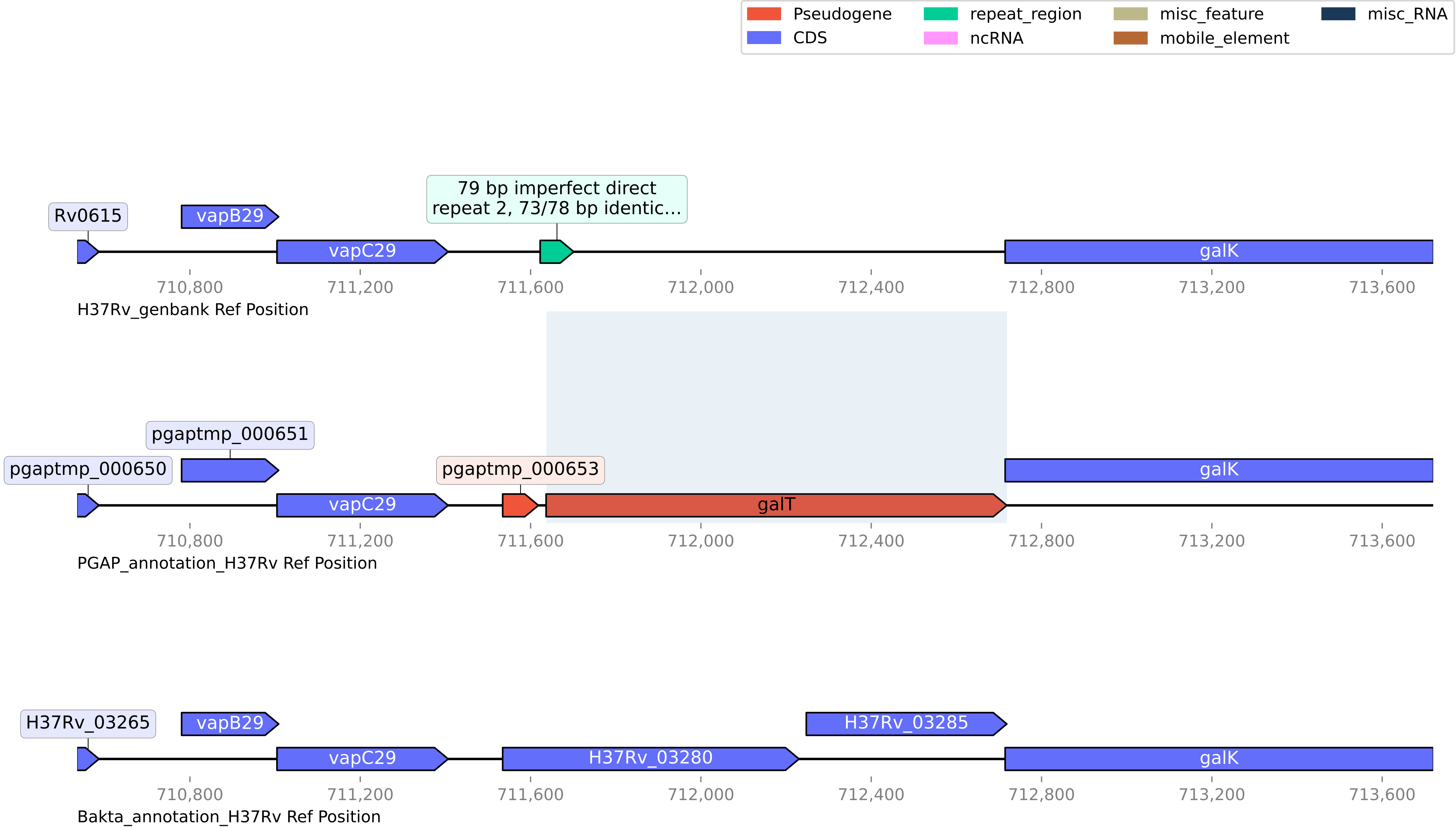

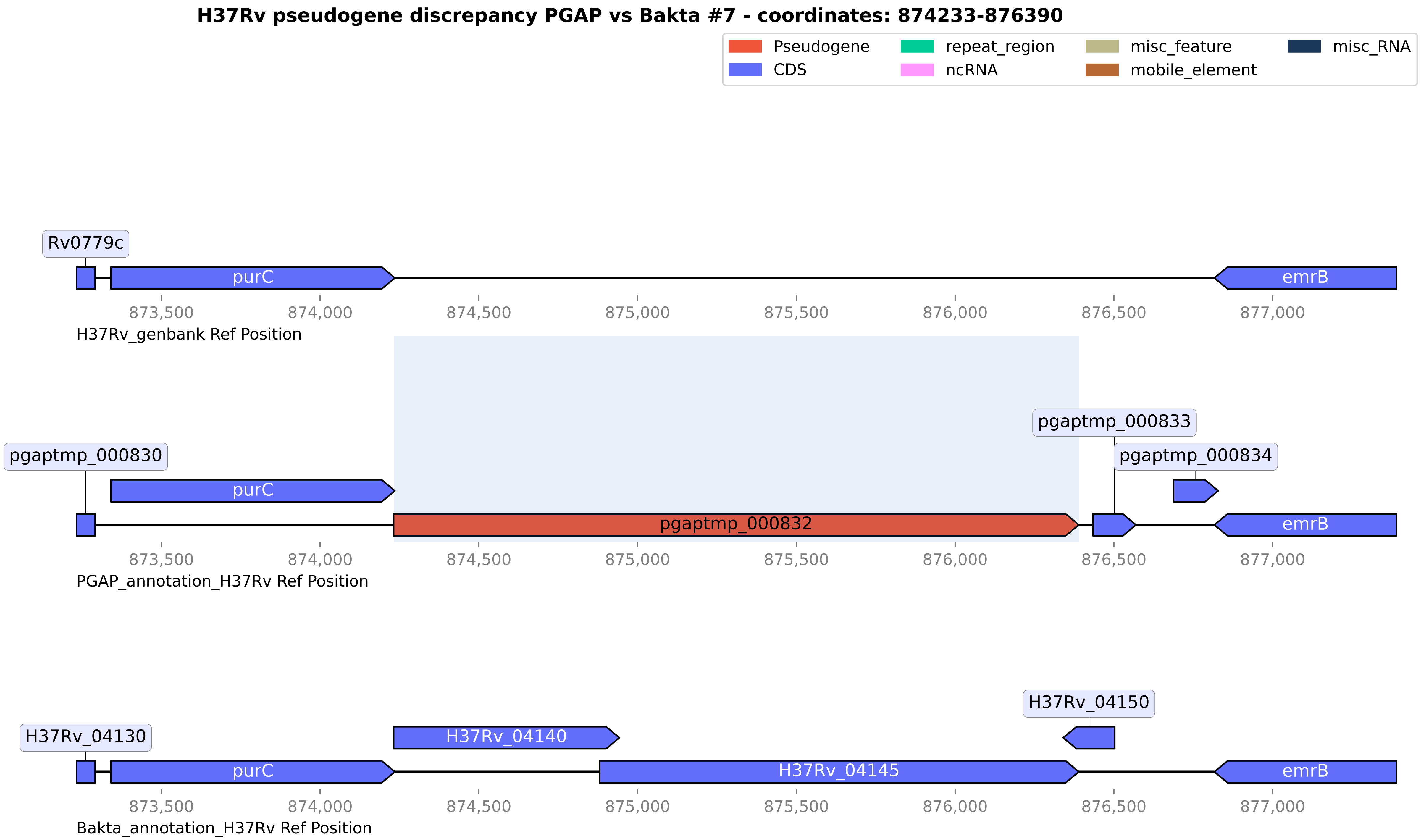
H37Rv pseudogene discrepancy PGAP vs Bakta #7 - coordinates: 874233-876390
Pseudogene
CDS
repeat_region
ncRNA
misc_feature
mobile_element
misc_RNA
Rv0779c
purC
emrB
873,500
874,000
874,500
875,000
875,500
876,000
876,500
877,000
H37Rv_genbank Ref Position
pgaptmp_000830
pgaptmp_000833
pgaptmp_000834
purC
pgaptmp_000832
emrB
PGAP_annotation_H37Rv Ref Position
H37Rv_04130
H37Rv_04140
H37Rv_04150
purC
H37Rv_04145
emrB
Bakta_annotation_H37Rv Ref Position
